# SOI: Robust identification of orthologous synteny with the *Orthology Index* and broad applications in evolutionary genomics

**DOI:** 10.1101/2024.08.22.609065

**Authors:** Ren-Gang Zhang, Hong-Yun Shang, Richard Ian Milne, Fabricio Almeida-Silva, Hengchi Chen, Min-Jie Zhou, Heng Shu, Kai-Hua Jia, Yves Van de Peer, Yong-Peng Ma

## Abstract

With the explosive growth of whole-genome datasets, the accurate detection of orthologous synteny has become crucial for the reconstruction of evolutionary history. However, the currently available methods for the identification of orthologous synteny have great limitations: the methods are difficult to scale with varying polyploidy histories, and the accurate removal of out-paralogy is challenging. In this study, we developed a scalable and robust approach, the *Orthology Index* (*OI*), to identify orthologous synteny. Our evaluation of a large-scale dataset with diverse polyploidization events demonstrated that the technique is highly reliable. This discovery highlights *OI* as a potentially unified criterion for the identification of orthologous synteny, and this is further validated using simulation-based benchmarks. In addition, we explore its broad applications in reconstructing the evolutionary histories of plant genomes, including inference of polyploidy, identification of reticulation, and phylogenomics. In conclusion, *OI* offers a robust, interpretable, and scalable approach for identifying orthologous synteny, significantly enhancing our analytical prowess in plant evolutionary genomics.

## Introduction

Reconstruction of the evolutionary histories of organisms, including inference of the tree/network of life, identification of polyploidy events, and placement of these events on the tree/network, usually relies on the orthologous relationships at gene, chromosomal block, chromosome, subgenome and/or whole genome scales. In general, the greater the number of reliably orthologous genes or loci are involved in the reconstruction of evolutionary history, the more confidence can be placed in the reconstruction. Indeed, phylogenomic reconstructions using genome-scale data have become the gold standard for understanding the evolution of lineages in the tree of life (1). Orthologous synteny has been established as a proxy to generate a maximum number of reliable orthologs, from the chromosomal to the subgenomic/genomic scales, for example, in the reconstruction of evolutionary histories for the octoploid strawberry (2) and the major angiosperm clades (3). The identification of orthologous relationships is also vital for other synteny-based analyses, such as inference of polyploidy (4) and subgenome phasing (2). However, accurate identification of orthologous synteny is still challenging. This is especially true in plants, where recurrent whole-genome duplication (WGD, also known as polyploidization) events have produced substantial numbers of syntenic paralogs. This can significantly complicate the inference of the orthology (1,5) and may mislead the reconstruction of evolutionary history. For example, the overlooked orthologous relationships in the synteny between two *Papaver* species resulted in an incorrect interpretation of the history of polyploidy in these species (6–8).

To date, two main strategies have been employed to identify orthologous synteny. One strategy (the “criterion strategy”) is to filter the detected synteny using certain criteria, which are usually specific to each case and are therefore not scalable for large-scale datasets. One widely-used criterion for this is the synonymous substitution rate (*Ks*). Because the WGD event (which produces paralogs) and speciation event (which produces orthologs) occurred at different times in the past, *Ks* between syntenic gene pairs can be used to differentiate the ages of orthologous and paralogous synteny (9,10). However, such events occur at different times in different lineages, and substitution rates also vary between lineages, both of which cause *Ks* values to vary case by case. Hence, *Ks*-based methods are not always effective for distinguishing syntenic blocks from different evolutionary events (11), and cannot be universally applied to the large-scale automated identification of orthologous syntenic blocks. Another criterion, *homo*, has been proposed for use with WGDI, to extract the best homology (orthology) of syntenic blocks (12). Unfortunately, this criterion relies on a parameter, *multiple*, to define the top number of hits as best hits (12), but *multiple* cannot be universally applied to different cases with different syntenic depth ratios. A further tool, QUOTA-ALIGN (11) was developed to screen orthologous syntenic blocks under given constraints on syntenic depths (*QUOTA*). However, the prior *QUOTA* criterion must be set by users, and will differ between cases with different evolutionary histories, thus it is also challenging to robustly distinguish synteny blocks from different events using QUOTA-ALIGN. For example, it is necessary to set *QUOTA* = 1:1 for *Arabidopsis thaliana* and *A. lyrata*, but to set *QUOTA* = 4:2 for *A. thaliana* and poplar (11). This is also similarly required when setting the *multiple* parameter in WGDI (-c option).

An alternative strategy (the “pre-inferred strategy”) for the identification of orthologous synteny is the use of pre-inferred orthologs to call synteny, with tools such as MCScanX_h (13). This strategy is scalable for large-scale datasets. However, hidden out-paralogs (i.e. false positives in ortholog inference) in the pre-inferred orthologs may result in out-paralogous syntenic blocks that need to be further removed with the above mentioned “criterion” strategy. Our previous work, such as that on *Salix* (14) and *Populus* (9), faced this problem. Moreover, the presence of hidden orthologs (i.e. false negatives in ortholog inference) can reduce the efficiency of the subsequent detection of orthologous synteny in practice.

To address these issues and to screen orthologous synteny robustly for large-scale genomic datasets, we developed a scalable approach called the *Orthology Index* (*OI*), which describes the proportion of collinear gene pairs that are pre-inferred as orthologs (see Methods). We evaluated the efficacy of *OI* using a large-scale dataset comprising 91 well-documented cases with diverse WGD and speciation events and a simulated benchmarking dataset. Our results demonstrated the high robustness and accuracy of *OI*, which outperformed the existing methods. We then explored its broad usefulness in evolutionary inference in analyses of polyploidy, reticulation and phylogenomics. We finally integrated the index into a toolkit (freely available from https://github.com/zhangrengang/SOI) to facilitate its use.

## Methods

### Empirical data collection and pre-processing

The genomic data were obtained from public databases or from the corresponding authors, as detailed in **Table S2**.

An all-versus-all BLAST search of protein sequences for each species was conducted pairwise using DIAMOND v0.9.24 (15). Orthologous relationships were inferred using OrthoFinder v2.3.1 (Emms and Kelly 2019) (parameters: -M msa). Syntenic/collinear blocks were identified with the ‘-icl’ option of WGDI (12) v0.6.2 (default parameters). The synonymous substitution rate (*Ks*) was calculated for homologous gene pairs using the ‘-ks’ option of WGDI (default parameters).

### Definition of the *Orthology Index*

We propose an index, named the *Orthology Index*, to distinguish the syntenic blocks of orthology from out-paralogy. The index (*OI*) is defined as:

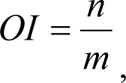

where *m* represents the total number of collinear gene pairs in a collinearity block, and *n* denotes the number of collinear gene pairs pre-inferred as orthologs. Thus, the index represents the proportion of orthologous syntenic gene pairs within a syntenic block. It ranges from 0 to 1 and can differentiate orthology from out-paralogy. Employment of the index is expected to result in a polarized pattern, where orthologous synteny is expected to result in a high *OI* value (with an expectation of 1, relying on the recall of pre-inferred orthologs), whereas out-paralogous synteny is expected to result in a relatively low *OI* value (with an expectation of 0, relying on the precision of pre-inferred orthologs). From our experience with many different cases (Figs. S1–90), the major peak with the highest index can be considered to be orthologous in principle.

### Implementation of the *Orthology Index*

Using this index as a foundation, we developed a user-friendly all-in-one toolkit (called SOI) for visualization and downstream analyses. The subcommand ‘dotplot’ enables visualization and evaluation of synteny, with the dots colored by the *OI* or *K*s values. The subcommand ‘filter’ retrieves orthologous blocks by discarding all blocks with less than a default *OI* value of 0.6. However, users can also apply a stringent standard (higher *OI* and block length cutoffs) to obtain long, highly credible blocks. The subcommand ‘cluster’ groups orthologous syntenic genes into syntenic orthogroups (SOGs) by constructing an orthologous syntenic graph and applying the Markov Cluster (MCL) algorithm (16) to perform graph clustering with breaking weak links and bridging unexpectedly disrupted links. The clustering algorithm is widely employed by popular tools of orthogroup clustering, such as OrthoFinder2 (17), SonicParanoid2 (18), and OrthoMCL (19). The subcommand ‘outgroup’ retrieves syntenic orthologs from outgroups that lack WGDs shared with ingroups. The subcommand ‘phylo’ reconstructs multi-copy or single-copy gene trees, by aligning protein sequences with MAFFT v7.481 (20), converting protein alignments to codon alignments with PAL2NAL v14 (21), trimming alignments with trimAl v1.2 (22) (parameter: - automated1) and reconstructing maximum-likelihood trees with IQ-TREE v2.2.0.3 (23). These gene trees serve as input to infer a species tree with the coalescence-based method ASTRAL-Pro v1.10.1.3 (24). The default threshold for missing taxa is set to 40 %, according to an evaluation for this parameter (25).

This tool is implemented in Python3 and supports synteny outputs from state-of-the-art synteny detectors, including MCscan/JCVI (26), MCscanX (13) and WGDI (12), as well as orthology outputs from OrthoFinder2 (17), Broccoli (27), SonicParanoid2 (18), Proteinortho6 (28), InParanoid (29), OrthoMCL (19) and other tools upon request. These orthology inference methods have competitive accuracy (18,27,30) and thereby should produce similar *OI* values. The tool can be easily installed using the conda environment or the Apptainer/Singularity container system (31). Since both synteny and orthology analyses are generally standard procedures for a genomics project, the *OI* tool can be seamlessly integrated into these and their downstream pipelines. The source code is accessible on GitHub (https://github.com/zhangrengang/SOI).

### Simulation-based benchmarks

To validate the feasibility and robustness of *OI*, we performed simulation-based benchmarks. We simulated a shared-WGD scenario similar to the Salicaceae case (Fig. 1A) using Zombi (32). The input species tree was “(((A1:0.2, B1:0.2):Δ*T*, (A2:0.2, B2:0.2):Δ*T*):0.2, O:Δ*T*+0.4):0.1”, where A1 and A2, and B1 and B2 is the two subgenomes of taxa A and B, respectively, and were treated as pseduo-species, and O is an outgroup species, while Δ*T* varied from 0.01 to 1 (scaled in time unit of substitutions per site). 5000 coding sequences were output for each simulation with other parameters (e.g. those regarding substitutions and chromosomal rearrangements) being default (see https://github.com/AADavin/Zombi/tree/master/Parameters/). Then the simulated three-taxon genomic data were input to the aforementioned OrthoFinder2–WGDI–SOI pipeline to calculate the *Ks* and *OI* values and to identify syntenic orthologs between taxa A and B. The true positives (TP), false positives (FP), and false negatives (FN) were counted from the results, and the precision (*P* = TP / (TP + FP)), recall (*R* = TP / (TP + FN)), and F1-score (2 × *P* × *R* / (*P* + *R*)) were calculated. The simulation was independently replicated 50 times for each Δ*T* condition.

**Fig. 1.**
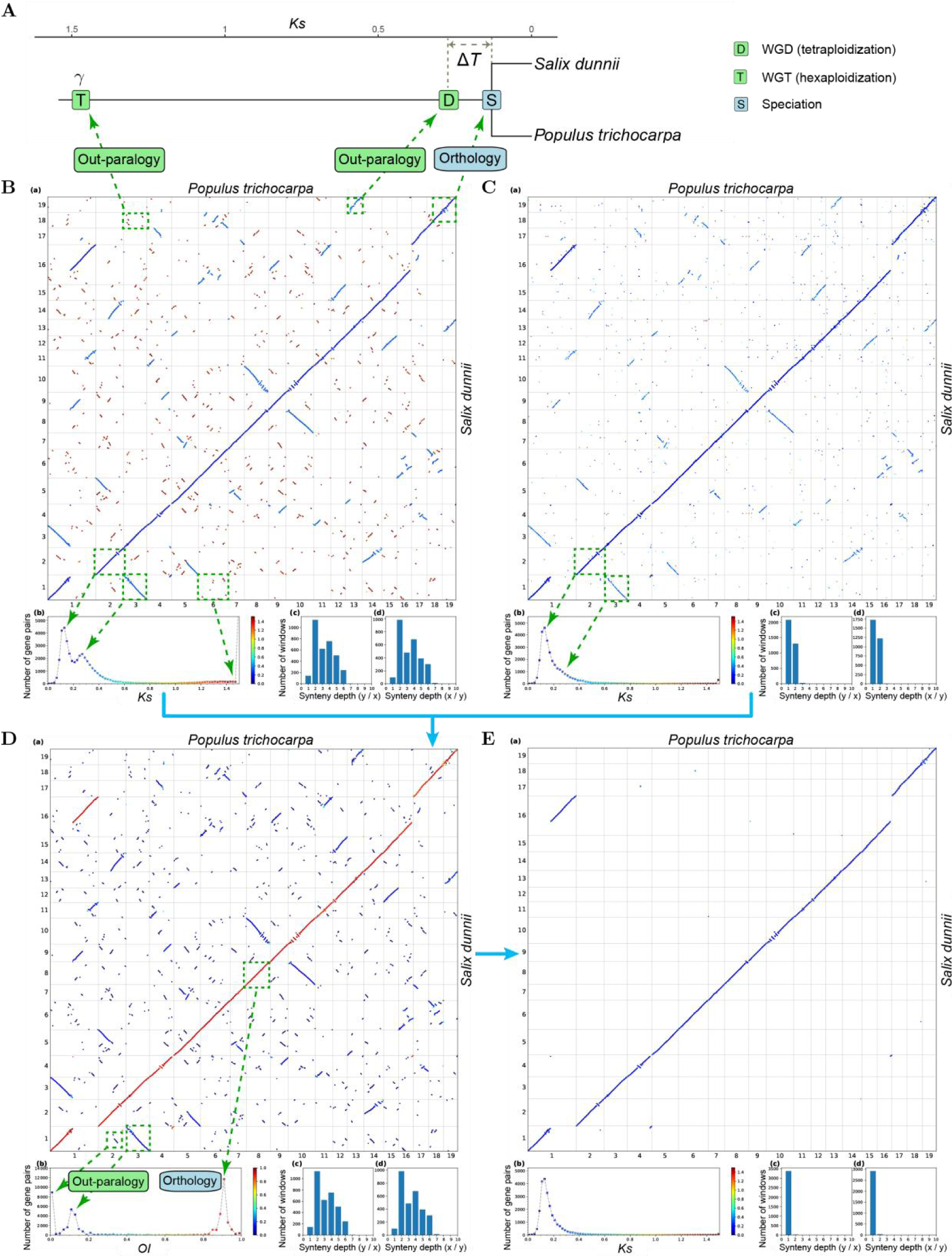
The illustration of the *Orthology Index* when identifying orthologous synteny for the typical Salicaceae case. **A**) Schematic of the evolutionary history of the poplar and willow genomes, adapted from the literature (33–36). **B**) *Ks*-colored dot plots showing synteny detected by WGDI, with an observable distinction between the three categories of syntenic blocks derived from three evolutionary events (three peaks: *Ks* ≈ 1.5, *Ks* ≈ 0.27, and *Ks* ≈ 0.13). **C**) *Ks*-colored dot plots illustrating the orthology as inferred by OrthoFinder2, with an observable high proportion (∼15 %) of hidden out-paralogs (*Ks* ≈ 0.27). **D**) *Orthology Index* (*OI*)-colored dot plots: integrating synteny (B) and orthology (C), showing polarized distinction of the three categories of syntenic blocks (three peaks: *OI* ≈ 0, *OI* ≈ 0.1, and *OI* ≈ 0.9). **E**) *Ks*-colored dot plots of synteny after application of an *OI* cutoff of 0.6, with clean one-to-one orthology as expected from the evolutionary history. B–E are plotted using the ‘dotplot’ subcommand with four subplots: a) dot plots colored by *Ks* or *OI* (x-axis and y-axis, chromosomes of the two genomes; a dot indicates a homologous gene pair between the two genomes), b) histogram of *Ks* or *OI* (x-axis, *Ks* or *OI*; y-axis, number of homologous gene pairs), using the same color map as the dot plots, c-d) synteny depth (relative ploidy) derived from 50-gene windows (x-axis, synteny depth; y-axis, number of windows). Examples of the syntenic blocks from three evolutionary events [referred to as WGT-SBs (*Ks* ≈ 1.5, *OI* ≈ 0), WGD-SBs (*Ks* ≈ 0.27, *OI* ≈ 0.1), and S-SBs (*Ks* ≈ 0.13, *OI* ≈ 0.9)] are highlighted with dashed squares. These are associated with the evolutionary events and peaks of *Ks* or *OI*, indicated by arrows, and labeled as ‘Out-paralogy’ or ‘Orthology’. Additional cases illustrating other lineages can be found in **Figs. S1–90** (summarized in **Table S1**).

## Results and Discussion

### Identification of orthologous synteny using the Orthology Index

We aimed to create a method that would robustly distinguish orthologous synteny from out-paralogy for genomes of any two species. The genomes should have shared a whole genome duplication event (WGD, or polyploidization; producing out-paralogy) and a speciation event (producing orthology), and these events should have occurred within a certain time period (denoted as Δ*T*, **Fig. 1A**). To evaluate our approach, we selected ninety well-documented plant species pairs, with each pair being descended from a common ancestor that had undergone at least one polyploidy event (**Table S1**, **Figs. S1-90**), along with the typical poplar–willow (Salicaceae) case (**Fig. 1**).

**Fig. 1** shows an analysis for the Salicaceae case, represented by *Populus trichocarpa* and *Salix dunnii*. The two genera speciated ∼50 million years ago (Mya) (33,34), after a WGD event in their common ancestor ∼60 Mya (33–36) (**Fig. 1A**). Additionally, both genera also share a much earlier paleohexaploidization event (the γ event, whole genome triplication or WGT) in the common ancestor of core eudicots ∼120 Mya (37), making three detectable evolutionary events in total (**Fig. 1A**).

With the “criterion” strategy using the collinearity detector WGDI (12), syntenic blocks derived from these three evolutionary events could be identified and distinguished visually using their distinct synonymous substitution rate (*Ks*) values and fragmentation extent (the more ancient the event, the more fragmented the blocks or greater the chromosomal rearrangements, and the higher the *Ks* value) (**Fig. 1B**). These three categories of syntenic blocks were denoted WGT-SBs (*Ks* ≈ 1.5, most fragmented, out-paralogous), WGD-SBs (*Ks* ≈ 0.27, moderately fragmented, out-paralogous), and S-SBs (*Ks* ≈ 0.13, least fragmented, orthologous) (**Fig. 1B**) according to the evolutionary events (i.e. WGT, WGD and speciation) (**Fig. 1A**). However, the recent two *Ks* peaks derived from WGD-SBs (out-paralogy) and S-SBs (orthology) largely overlapped (**Fig. 1B**) and were difficult to split with a hard cutoff. This is likely to be because the evolutionary rates in different genes also vary, resulting in similar accumulated substitutions for genes from different events. Indeed, only a few empirical cases (e.g. **Figs. S4, 30**) can employ a hard *Ks* cutoff to distinguish orthology and out-paralogy. For some extreme cases where the Δ*T* was quite small, *Ks* peaks from two events (shared WGD and speciation) overlapped completely, and showed only one peak (e.g. **Figs. S1, 8, 16, 34**) with no distinction between orthology and out-paralogy.

With the “pre-inferred” strategy using the orthology inference method, OrthoFinder2 (17), the peak (*Ks* ≈ 0.27) of out-paralogs from WGD-SBs was significantly reduced, while out-paralogs from the older WGT event (WGT-SBs, *Ks* ≈ 0.13) were almost entirely eliminated (**Fig. 1C**). This suggests that the level of accuracy of orthology inference by OrthoFinder2 (∼ 85 % orthologs located in the orthologous chromosomal segments for the Salicaceae case) is relatively high. Nevertheless, in the Salicaceae case, this method inferred a substantial number of “orthologous” genes (∼ 15 %) that exhibit synteny in the out-paralogous blocks (*Ks* ≈ 0.27; **Fig. 1C**), suggesting that they are hidden paralogs and would introduce out-paralogous synteny if they were imported into tools such as MCScanX_h (13). This problem was also observed in numerous instances where the Δ*T* was not substantial (e.g. **Figs. S1, 5, 6, 8, 10**), which could be attributed to systematic errors or biased gene loss, and which could have a detrimental effect on downstream analyses, such as phylogenomics (1).

Consequently, we combined the algorithmic advances of the two methods described above by introducing a straightforward index, referred to as the *Orthology Index* (*OI*), to determine the orthology of a syntenic block. *OI* represents the proportion of gene pairs that are pre-inferred to be orthologs in a syntenic block. Thus, orthologous synteny expects an *OI* of 1 (i.e., all syntenic gene pairs are inferred to be orthologs), while out-paralogous synteny expects an *OI* of 0 (i.e., none of the syntenic gene pairs are inferred to be orthologs). However, for real datasets, due to the limited recall and precision from orthology inference methods, the *OI* distribution could be biased from the expectations. For the Salicaceae case, the WGT-SBs and WGD-SBs (both out-paralogous but with different ages) exhibit an *OI* of nearly 0 and approximately 0.1, respectively, whereas the S-SBs (orthologous) display an *OI* peak of approximately 0.9 (**Fig. 1D**), close to the expectations. The *OI* distribution reflected the same orthologous and out-paralogous relationships as *Ks* in this case, but was able to much better distinguish between them than was *Ks* (**Fig. 1 B** and **D**). Remarkably, the *OI* peaks derived from orthology and out-paralogy did not overlap, but showed a polarized pattern with a very clear, wide dividing range (*OI* = ∼0.3–0.7) between the orthology and out-paralogy peaks (**Fig. 1D**). Therefore, in this case, the *OI* was easily able to distinguish orthology from out-paralogy, with a hard cutoff between 0.3 and 0.7 (**Fig. 1D**).

We next applied the index analysis to ninety study cases with diverse polyploidy histories and varying Δ*T* (**Table S1, Figs. S1-90**). We found that there were also clear divides around *OI* = ∼0.5–0.6 in nearly all instances (although minor noise around *OI* = ∼0.5–0.6 can be observed in some cases) (**Figs. 2A, S1-90D**), suggesting that a potentially unified *OI* criterion could be employed for the identification of orthologous synteny. The *OI* cutoff was further adjusted to 0.6 empirically to identify orthologous synteny according to the *OI* distributions in these cases (**Figs. 1D and 2A, Figs. S1**–**90D**), in order to balance the visible false positives (out-paralogy mis-identified as orthology) and false negatives (excessive removal of true orthologous synteny). Interestingly, we observed that the index performed exceptionally and consistently well in identifying orthologous synteny in these instances (**Fig. 1E, Figs. S1–90E**).

**Fig. 2.**
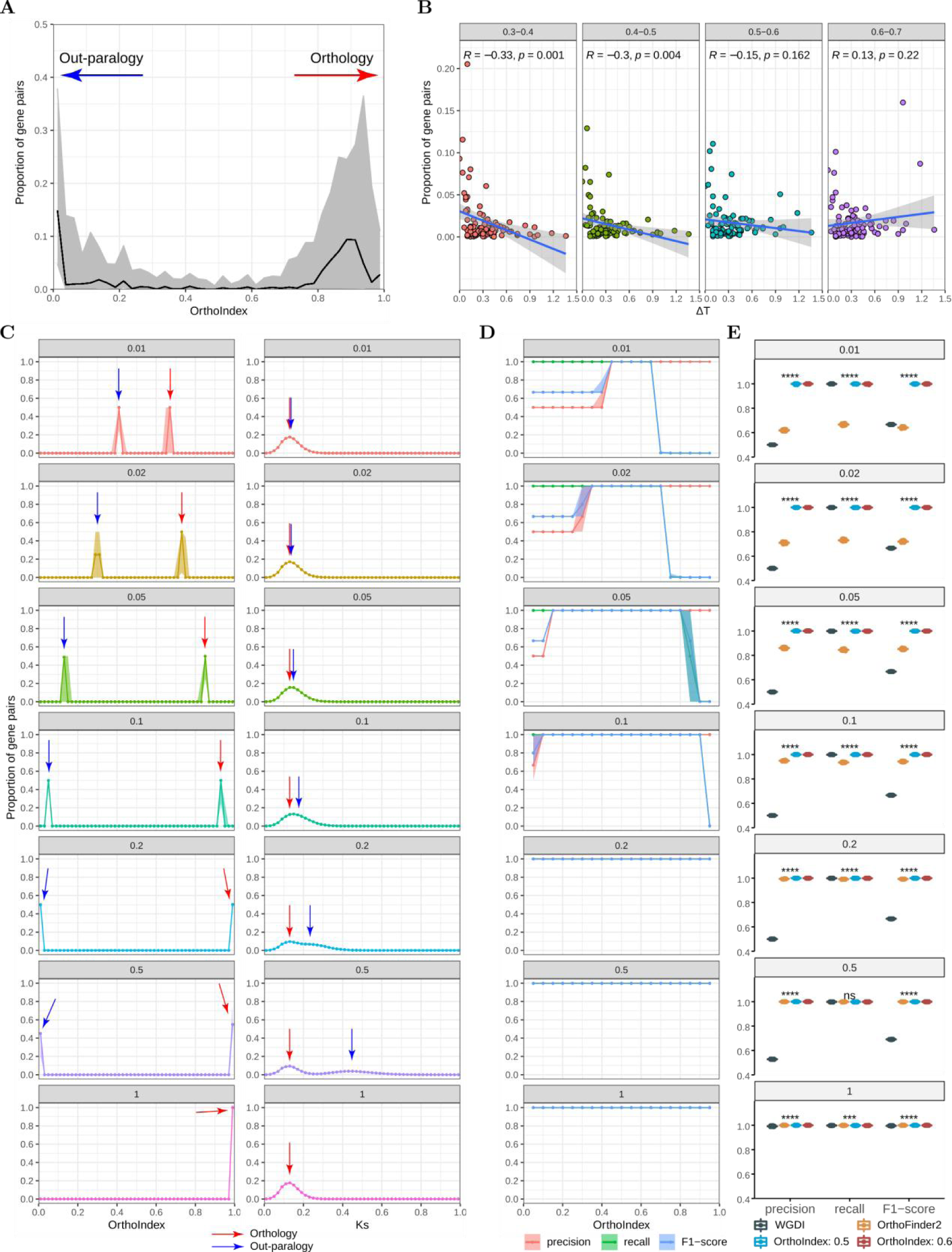
The performance of the *Orthology Index* in identifying orthologous synteny in empirical and simulated datasets. **A)** Summary of *OI* distributions in the 91 empirical test cases. The black line and gray shadow represent the median and percentile-based 95 % confidence interval (CI) values, respectively. **B)** The correlation between Δ*T* and the noise around *OI* = 0.5 in the empirical datasets. The Δ*T* values were roughly obtained from the *Ks* peak values of shared WGD and speciation events. The noise was indicated by the accumulated proportion of syntenic gene pairs within 0.3–0.4, 0.4–0.5, 0.5–0.6 and 0.6–0.7 intervals of the *OI*. **C)** Comparisons of inter-genomic *Ks* and *OI* distributions in the simulated datasets at different values of Δ*T* (Δ*T* ɛ {0.01, 0.02, 0.05, 0.1, 0.2, 0.5, 1}, in time units of expected substitutions per site). The line/point and shadow represent the median and 95 % CI values from 50 repeated simulations, respectively. **D)** Comparisons of the precision, recall and F1 scores of orthology identification using different *OI* cutoffs (0.05–0.95) from the simulated benchmarks. The line/point and shadow represent the median and percentile-based 95 % CI values from 50 repeated simulations, respectively. **E)** Comparisons of the precision, recall and F1 score of orthology identification using WGDI, OrthoFinder2 and *OI* with cutoffs 0.5 and 0.6, based on the simulated benchmarks. The boxplot represents the values from 50 repeated simulations. ns, *P* > 0.05; ***, *P* <= 0.001; ****, *P* <= 0.0001; Kruskal-Wallis test.

For example, in the Salicaceae case, a cutoff value of *OI* = 0.6 resulted in retrieval of a clean 1:1 orthology relationship without visible false positives or false negatives (**Fig. 1E**), as expected. In this case, 12.2 % of the syntenic blocks and 47.6 % of the syntenic pairs were retained. Of the retained syntenic gene pairs, 90.4 % were pre-inferred as orthologs using OrthoFinder2, while the remaining 9.6 % were presumed to be orthologs but were not identified using OrthoFinder2. Moreover, although some visible false positives or false negatives were observed in a few study cases, there were many fewer false positives than when using the above two strategies (**Figs. S50, 68, 74, 86E**). This finding highlights the efficiency and robustness of *OI* as a unified criterion for identifying orthologous synteny.

To validate the feasibility and robustness of *OI*, we further performed benchmarking by simulating a shared-WGD scenario similar to that of the Salicaceae case (**Fig. 1A**). Δ*T* was correlated to the unexpected noises around *OI* = 0.5 in the empirical datasets (**Fig. 2B**), and was therefore assumed to be a key parameter affecting the accuracy in identifying orthologous synteny. With a varied Δ*T* (Δ*T* ɛ {0.01, 0.02, 0.05, 0.1, 0.2, 0.5, 1}, in a time unit of expected substitutions per site), *OI* consistently showed much higher ability to distinguish between orthology and out-paralogy than did *Ks* (**Fig. 2C**). At values of Δ*T* between 0.01 and 0.1, *Ks* values from both orthology and out-paralogy showed only one peak without any observable distinction. When Δ*T* reached 0.2, the two *Ks* peaks still overlapped significantly and could not be efficiently distinguished. In striking contrast, *OI* showed clear dividing ranges around *OI* = ∼0.5–0.6 for all values of Δ*T* (**Fig. 2C**). By applying a hard cutoff of *OI* around ∼0.5–0.6, both the precision and recall consistently approached to 1.0 under different values of Δ*T* (**Fig. 2D**). Moreover, when Δ*T* was between 0.01 and 0.05, where OrthoFinder2 indicated a relatively low accuracy in ortholog inference (precision = 0.62–0.85, recall = 0.66–0.83, and F1 score = 0.64–0.84), the *OI* method with a hard cutoff (cutoff = 0.5 or 0.6) highly improved both the precision (0.992–1.0) and recall (0.992–1.0) of ortholog detection (**Fig. 2D–E**). This suggests that *OI* could significantly improve ortholog inference under these conditions and could therefore facilitate syntelog-based analyses, such as phylogenomics and pangenomics. Overall, these simulation-based benchmarking results were highly consistent with those above based on empirical datasets, demonstrating that the *OI* method outperforms the existing methods (*Ks*-based and “pre-inferred”) and is able to produce accurate results when identifying orthologous synteny.

*OI* can therefore be applied in automated pipelines for large-scale datasets (ranging from dozens to hundreds of genomes at present) with a unified threshold of *OI*. We integrated the *OI* method into a toolkit (https://github.com/zhangrengang/SOI) to identify orthologous synteny and to facilitate downstream applications.

### Applications of *OI* in polyploidy inference

Polyploidization (or WGD) has been identified as a critical mechanism in eukaryotic evolution and is pervasive in ancient and recent plant lineages (38). Both synteny and *Ks-*based methods have been widely used in the inference of polyploidy, including identification of occurrence and resulting ploidy and the phylogenetic placement of WGDs (12,39). However, inter-species orthologous synteny patterns have been previously overlooked in polyploidy inference, potentially leading to misunderstandings regarding the evolutionary history of WGD events (see discussions in (40)). We argue that distinguishing synteny resulting from orthology from that resulting from out-paralogy is not only straightforward but also vital for the inference of the evolutionary history of the polyploidy. We illustrate a straightforward model to explain the process of polyploidy inference using the patterns of orthologous synteny (**Fig. 3A**). When two genomes exhibit a 2:2 ratio of synteny depth, there are two main hypotheses to consider: either the genomes share a tetraploidization event (WGD) that occurred in a common ancestor, or they each underwent a lineage-specific tetraploidization event (WGD) independently, after they diverged (**Fig. 3A**). The *OI* offers a straightforward and visual method to test these two hypotheses with separated orthology and out-paralogy. If the genomes display a clear 1:1 orthology + 1:1 out-paralogy, similar to the Salicaceae case (**Fig. 1D**; ignoring the very ancient out-paralogy from the γ event), the first hypothesis is supported (**Fig. 3A, left panel**). In contrast, if they exhibit a 2:2 orthology, the second hypothesis is supported (**Fig. 3A, right panel**). Applying this to 90 known cases (**Figs. S1–90**) suggests that these inferences of polyploidy history from orthologous synteny patterns are reasonable and accurate.

**Fig. 3.**
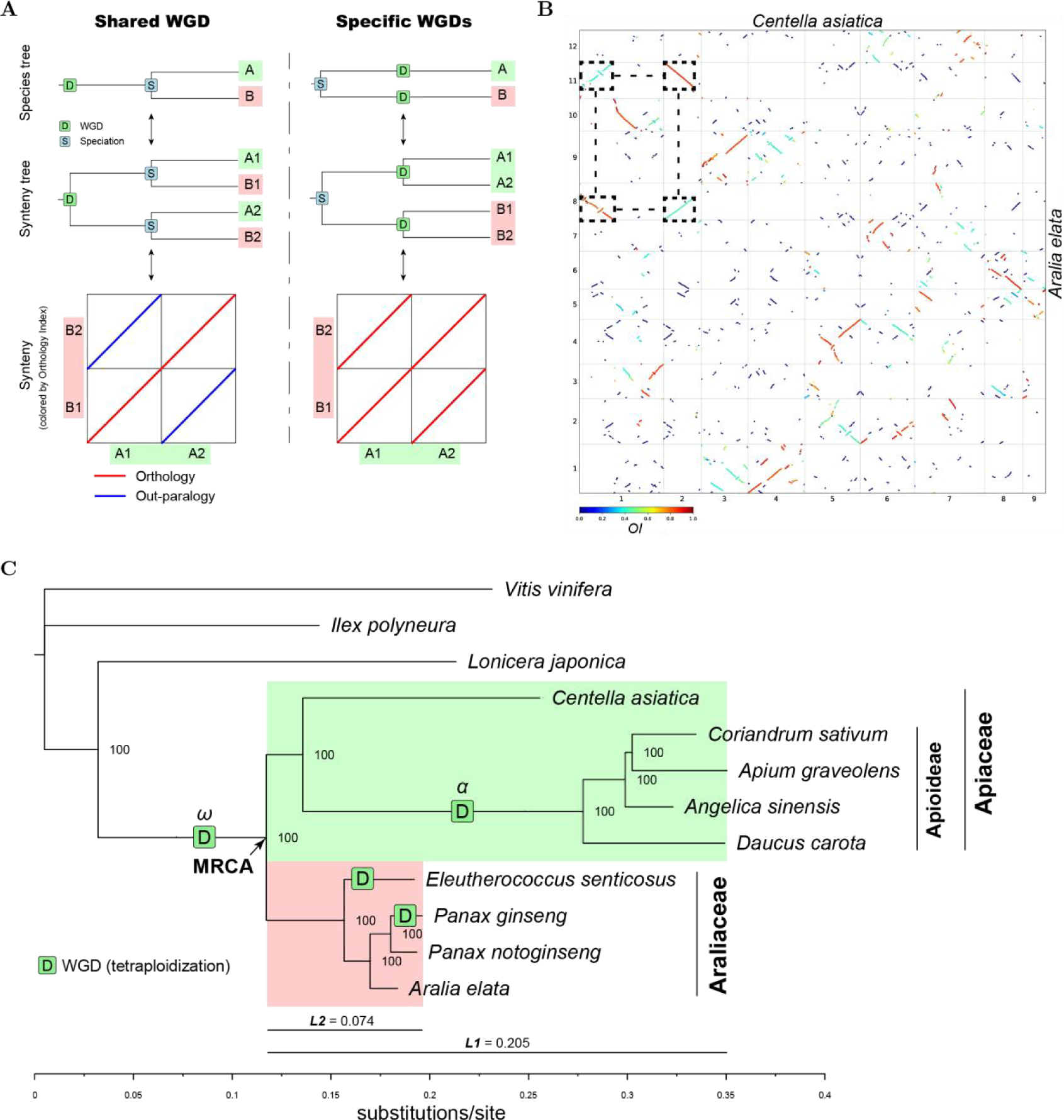
Inference of polyploidy in Apiales (Apiaceae + Araliaceae) genomes using the *Orthology Index*. **A)** Schematic illustration of determination of shared or lineage-specific polyploidy events using the orthologous synteny patterns identified with the *Orthology Index*. Despite a similar 2:2 ratio of synteny depth, the two scenarios have distinct patterns of orthologous synteny (1:1 orthology vs. 2:2 orthology). The labels A and B indicate two species, and A1, A2 and B1, B2 indicate duplicated chromosomes or blocks from WGD event(s). **B)** *Orthology Index*-colored dot plots indicating orthologous and out-paralogous synteny between the genomes of *Centella asiatica* (Apiaceae) and *Aralia elata* (Araliaceae). A typical 1:1 orthology + 1:1 out-paralogy synteny pattern is highlighted by the dashed squares. **C)** Phylogeny reconstructed from the genomes of certain species in the Apiaceae and the Araliaceae (Apiales) with labels indicating polyploidization events. *L1* and *L2* represent the average branch length (substitution rate) of the Apiaceae and the Araliaceae, respectively, from their most recent common ancestor (MRCA). Numbers at the nodes denote bootstrap values. The maximum-likelihood phylogenetic tree was reconstructed using IQ-TREE2, based on concatenated codon alignments of 2,363 single-copy genes (with at most 20 % taxa missing). Additional evidence supporting the inferred polyploidization events can be found in **Figs. S91**–**97**.

Apiaceae and Araliaceae (both within Apiales) provide the perfect test case for this. Previous synteny and *Ks-*based analysis indicates either a WGD event in their common ancestor (41), or independent WGD events in the two families (42–44). Therefore, we reexamined the polyploidy of the available genomes from the two families using the newly developed *OI* method.

We first determined the occurrence of polyploidization and the resulting ploidy using the orthologous synteny patterns and a summarized term, relative ploidy (*p*, i.e, the overall orthologous synteny depth relative to a given reference genome), for the Apiales genomes, referring to the *Vitis vinifera* (Vitaceae) genome, which has not undergone lineage-specific polyploidy since the paleohexaploid γ event shared by the core eudicots (37). Specifically, *p* > 1 indicates one or more independent polyploidization events since the divergence with the reference genome, which resulted in a ploidy of 2*p* for a haploid genome assembly (e.g. *p =* 2 suggesting tetraploidy and *p =* 3 suggesting hexaploidy). Using *Centella asiatica* (Apiaceae) and *Aralia elata* (Araliaceae) as representatives, we observed a clear 2:1 orthologous synteny depth (*p =* 2) with the *Vitis vinifera* (Vitaceae) genome (**Fig. S91**), suggesting that both species have undergone a tetraploidization event following their divergence from Vitaceae, consistent with previous research (43,44). However, upon comparison of the two genomes, a clear 1:1 orthology (*p =* 1) + 1:1 out-paralogy pattern was highlighted by the *OI* (**Fig. 3B**), aligning with the first model (**Fig. 3A, left panel**) and thus supporting the hypothesis of shared WGD (ω, **Fig. 3C**) in their common ancestor (see Song et al (41)). This inference was corroborated with macro-synteny phylogenies (**Fig. S92**), demonstrating the reliability of this technique. Similar patterns were also observed in other combinations of genomes from the two families (e.g. *C. asiatica* : *Eleutherococcus senticosus* = 1:2 orthology) (**Figs. S93**–**94**), further supporting the inference. Notably, we observed a considerable disparity in branch length (or substitution rate) between the two families from their most recent common ancestor (MRCA), with the mean branch length of Apiaceae (*L1* = 0.205) nearly three times than that of Araliaceae (*L2* = 0.074) (**Fig. 3C**). This discrepancy might explain the unreasonable phylogenetic placements drawn from the traditional *Ks*-based methods (42–44), which assumed equal substitution rates for different lineages when estimating the relative timing of polyploidization and speciation events. In fact, the assumption of equal substitution rates is often false or uncertain in practice.

Furthermore, we deduced additional lineage-specific polyploidy events within the two families using the *OI*-based approach **(Fig 3C)**. We observed that all studied genomes from species in the Apioideae subfamily (including *Daucus carota*, *Angelica sinensis*, *Apium graveolens* and *Coriandrum sativum*) exhibit a 2:1 orthologous synteny depth (*p =* 2) when compared to the *C. asiatica* genome (**Fig. S95**), suggesting that they each experienced a tetraploidization event subsequent to their divergence from *C. asiatica*. Moreover, any two genomes within this subfamily display distinct 1:1 orthology (*p =* 1) + 1:1 out-paralogy patterns (**Figs. S56, 96**), suggesting a shared tetraploidization event (α), which is in line with previous research (41,44). Thus the α event was placed between the stem and crown of the Apioideae subfamily (**Fig. 3C**). In addition, employing the *OI*, we also inferred one species-specific tetraploidization event each in the *Panax ginseng* and *Eleutherococcus senticosus* genomes (**Fig. 3C, Figs. S57–59, 97**), consistent with some previous research (43,44) but in conflict with the inference of two independent WGD events in *P. ginseng* in Song et al (41), which may be misled by other types of gene duplications (non-WGD).

In summary, each previous inference for polyploidy in the Apiales genomes (41–44) was partly correct but did not represent the whole picture. *OI* can now unravel the history of polyploidy with strong evidence of orthologous synteny patterns. These findings suggest that misinterpretation may be difficult to avoid if inter-genomic orthologous synteny patterns are not considered, potentially leading to the confusion with out-paralogs and other types of gene duplications. Traditional *Ks*-based methods can be only used with great uncertainty in the identification of occurrence and phylogenetic placement of WGDs (45), while traditional synteny-based methods can not be used in phylogenetic placement of WGDs when showing potential out-paralogous synteny. In contrast, orthologous synteny inferred using the *OI,* which combines both synteny and orthology information, is straightforward to use in the identification of the occurrence of polyploidization and the resulting ploidy, as well as in the high-confidence phylogenetic placement of WGDs.

### Applications in identification of reticulation

Reticulation, driven by allopolyploidization or hybridization, is a significant factor in eukaryotic evolution, yielding novel phenotypes that facilitate ecological diversification and the occupation of new niches (46). Numerous genomes originating from recent reticulation events have been documented (summarized in Jia et al (47)), encompassing critical cereals (48), fruits (49), vegetables (50), trees (9) and fish (51). We evaluated the application of the *Orthology Index* in several representative, well-documented cases, ranging from simple to complex evolutionary scenarios (**Fig. 4**).

**Fig. 4.**
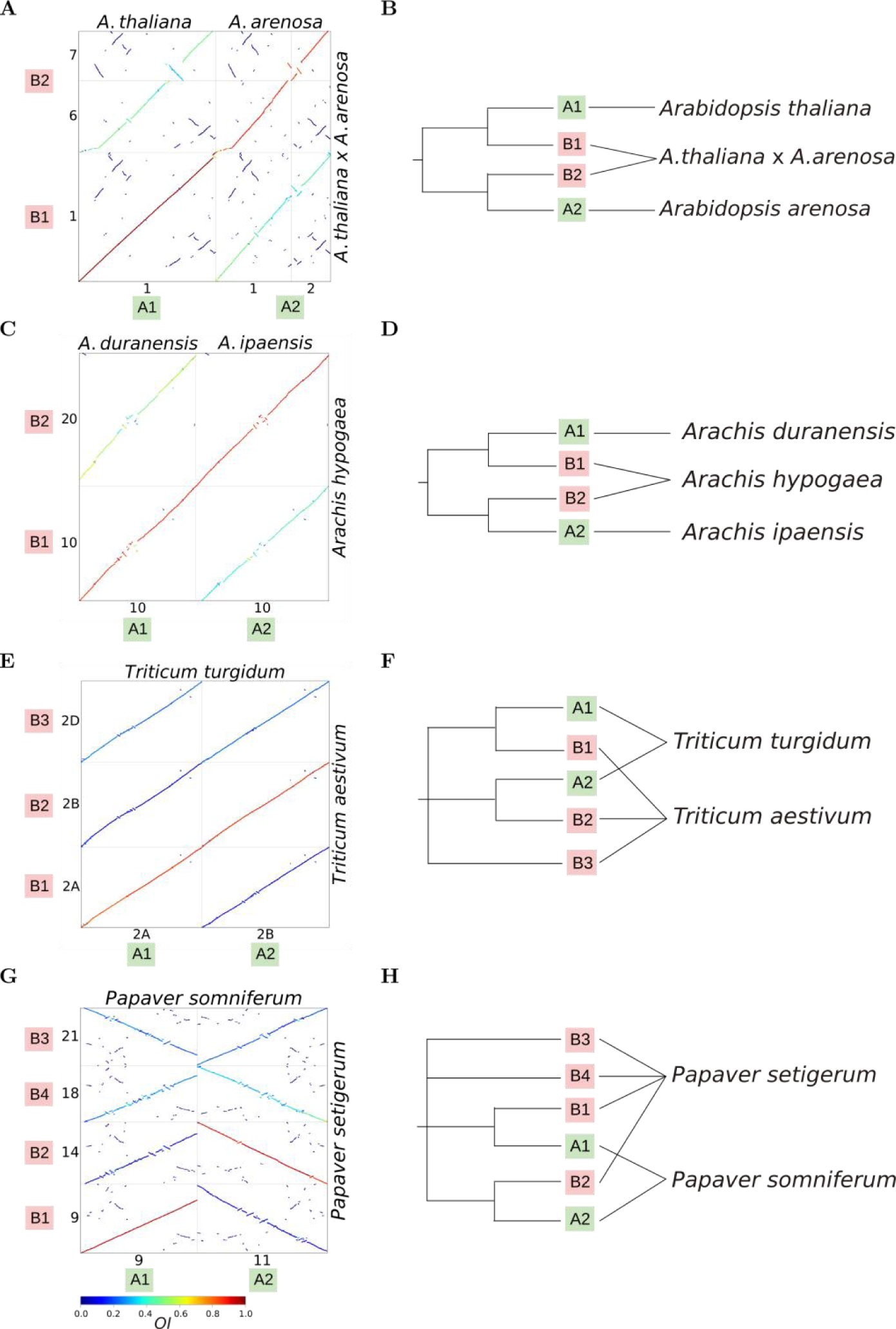
Examples of reticulation inferences based on the *Orthology Index*. **A**–**B**) *Orthology Index*-colored dot plots (**A**) of the genomes of *Arabidopsis thaliana* + *Arabidopsis arenosa* and their hybrid *Arabidopsis thaliana* × *Arabidopsis arenosa*, and the inference (**B**; in dendrogram form) from the orthologous relationships. **C**–**D**) *Orthology Index*-colored dot plots (**C**) of the genomes of the tretaploid *Arachis hypogaea* and its diploid progenitors, and the inference (**D**; in dendrogram form) from the orthologous relationships. **E**–**F**) *Orthology Index*-colored dot plots (**E**) of the genomes of the hexaploid *T. aestivum* and its intermediate tretaploid *T. turgidum*, and the inference (**F**; in dendrogram form) from the orthologous relationships. **G**–**H**) *Orthology Index*-colored dot plots (**G**) of the genomes of the neo-octoploid *P. setigerum* and its intermediate tretaploid *P. somniferum*, and the inference (**H**; in dendrogram form) from the orthologous relationships. Only one set of representative homoeologous chromosomes is shown in the dot plots; dot plots with the full set of chromosomes can be found in **Figs. S98-101**.

The cases of *Arabidopsis* (52) and *Arachis* (53) represent simple hybridization or allopolyploidization scenarios. From the *OI*-colored dot plots, the two subgenomes of both the hybrid (*Arabidopsis thaliana × Arabidopsis renosa*) and tetraploid (*Arachis hypogaea*) show clear and separate orthologous relationships with their diploid progenitors (**Fig. 4A and 4C, Fig. S98-99**). Based on the orthologous relationships revealed by the *OI*, the hybridization events can be easily inferred with straightforward visualization (**Fig. 4B and 4D**).

The orthologous relationships between the genomes in the complicated polyploid species complexes *Triticum* and *Papaver* are also clear from the *OI*-colored dot plots (**Fig. 4E and 4G, Fig. S100–101**). The two subgenomes of the tetraploid *T. turgidum* are orthologous to the two subgenomes of the hexaploid *T. aestivum* (**Fig. 4E, Fig. S100**), leading the inference that *T. turgidum* is the intermediate tetraploid progenitor of the allohexaploid *T. aestivum* (**Fig. 4F**), in line with previous results (48). Similarly, a reticulate allopolyploidization origin in two *Papaver* genomes (**Fig. 4H**) can be inferred from the *OI*-colored dot plots (**Fig. 4G, Fig. S101**). This inference agrees with our previous work (7); however, the phylogenetic relationships between homoeologous subgenomes cannot be resolved directly from the *OI*-colored dot plots and require further evidence, such as chromosome or subgenome-scale phylogenies (7).

Therefore, we conclude that reticulate speciation with intact subgenomic structure can be simply and directly inferred from the *OI*-colored dot plots. Although phylogeny-based methods could provide a deeper insight into the reticulation (2), the *OI* method is more straightforward and allows visualization that at least suggests a hypothesis to test further. Indeed, the straightforward visualization of the patterns of orthologous synteny could make it easy to recognize the signals of reticulation and thus to reasonablely elucidate the evolutionary history (e.g. (6–8)).

### Applications in phylogenomics

Accurate inference of orthology plays a crucial role in the estimation of species trees, and the pseudo-orthologs (i.e. hidden paralogs) derived from WGD and gene loss can mislead species tree inference greatly under some circumstances (54). As demonstrated above (**Fig. 1–2, Figs. S1–90**), the *OI* exhibited a high level of accuracy in identifying syntenic orthologs (or syntelogs) and can therefore minimize the detrimental influence of pseudo-orthologs. Consequently, the resulting knowledge of syntenic orthologs can be directly applied to species tree reconstruction. We used the example of the core eudicots to showcase this application. It is accepted that all core eudicots share a paleohexaploidization event (γ event, WGT) around 120 Mya, whereas no two orders within the core eudicots share an additional polyploidy event (38). Therefore, it is appropriate to use the *OI* to remove the out-paralogy produced from the γ event, and to better resolve the phylogenetic relationships among the orders of the core eudicots, many of which remain poorly resolved, such as the Celastrales–Oxalidales–Malpighiales (COM) clade (55). Here we utilized the genome-scale syntenic orthologs inferred by the *OI* to reconstruct a backbone phylogeny of the core eudicots, aiming to minimize the detrimental influence of the γ event.

We compiled a high-quality genomic dataset that covers 28 (70 %) of the 40 orders and 98 (33 %) of the 298 families of core eudicots treated in APG IV (55). We then applied the *OI* to this dataset to identify syntenic orthologs. This resulted in the identification of 54,322 syntenic orthogroups (SOGs). After filtering, 12,277 multi-copy and 5,154 single-copy SOGs were retrieved, allowing for up to 40 % missing taxa (**Fig. 5A**). This imbalance between the numbers of multi-copy and single-copy SOGs is attributable to lineage-specific polyploidy events within the core eudicots. As a result of these polyploidy events, the occupancy of single-copy SOGs showed significant decreases in species with high relative ploidy (i.e. orthologous syntenic depth relative to the *Vitis vinifera* genome) (**Fig 5B**). Nevertheless, the order-level species tree topologies based on the two gene sets were identical and both were strongly supported with high posterior probabilities (**Fig. 5C)**, although the tree based on multi-copy SOGs was more robust with equal or higher posterior probabilities at nearly all nodes (**Fig. 5C)** and there were slight differences in the positions of a few of the species (**Figs. S102–103**).

**Fig. 5.**
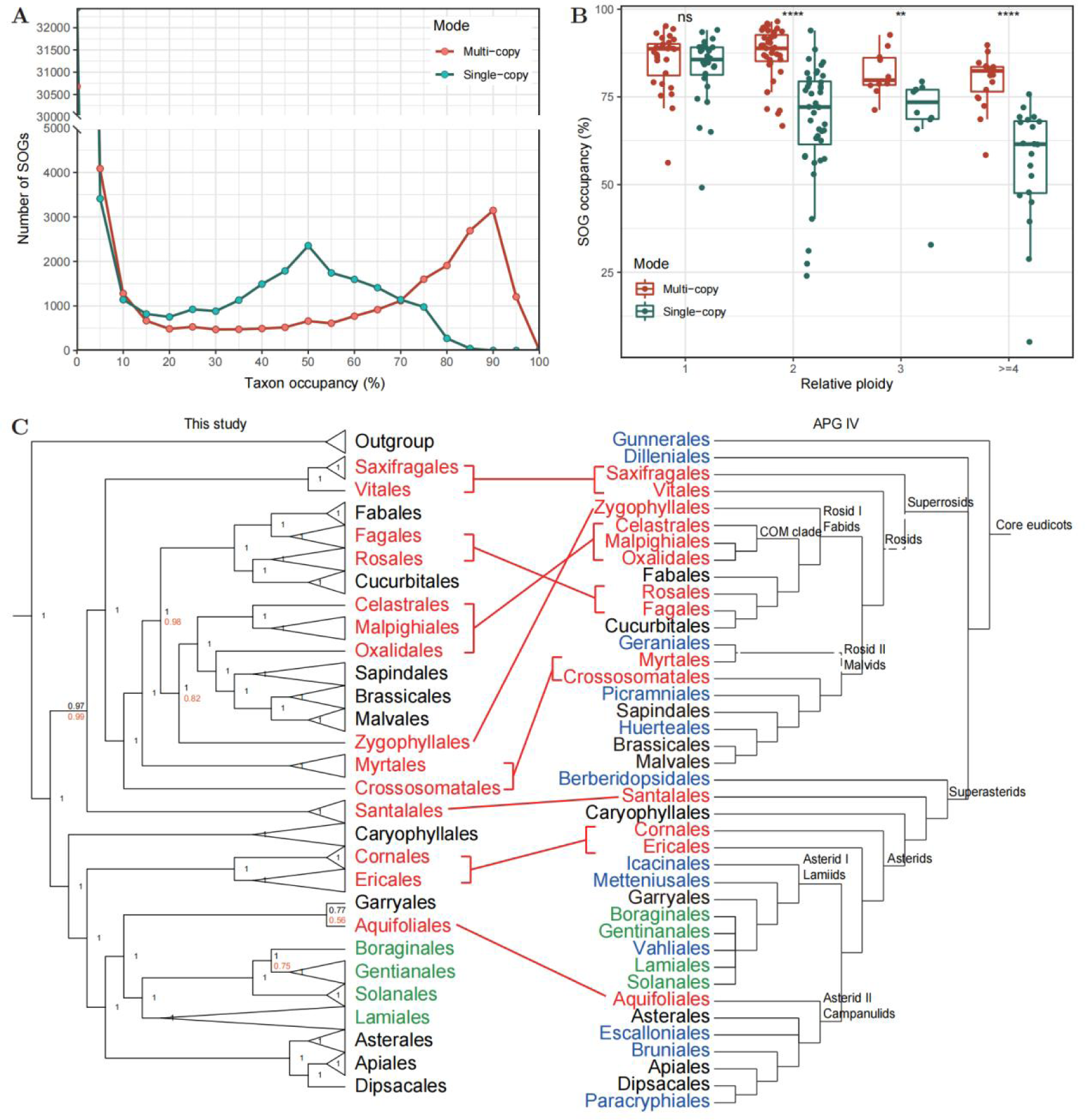
An example (core eudicots) of phylogenomics based on the *Orthology Index*. **A**) The number of multi-copy and single-copy syntenic orthogroups (SOGs) with different taxon occupancy. **B**) The SOG occupancy in species with different relative ploidy (i.e. orthologous syntenic depth to the grape genome) with at most 40 % taxa missing. Each point represents one species. ns, *P* > 0.05; **, *P* <= 0.01; ****, *P* <= 0.0001; Wilcoxon test. **C**) Phylogenetic relationships within the core eudicots reconstructed in this study versus in APG IV. Conflicting positions are marked in red; unresolved relationships in APG IV are marked in green, and orders not covered in this study are marked in blue. The numbers at the nodes are posterior probabilities from ASTRAL, with the black representing those from the multi-copy SOGs and orange representing those from the single-copy SOGs (omitted for equal values). Further details of the two trees reconstructed in this study can be found in **Figs. S102–103**.

We observed large incongruences between our phylogeny and the APG IV (55), as nearly half of the orders covered in this study have inconsistent phylogenetic positions (**Fig. 5C)**. For example, in our analysis, the COM clade was not monophyletic and was placed with the malvids, in contrast to that from APG IV (**Fig. 5C)**. The incongruences are likely to be due to the fact that the APG IV analysis was conducted using a very limited number of phylogenetic markers (55). In contrast, recent phylogenomics/phylotranscriptomic studies are consistent with most of our findings (56–63). For example, the phylogenetic positions of the Fagales, Rosales, Celastrales– Oxalidales–Malpighiales, Myrtales, Cornales and Ericales in our study (**Fig. 5C)** are consistent with the genome-scale phylogenomics based on coalescent-based analysis of 482 single-copy nuclear orthologous sequences (60). In addition, the phylogenetic positions of the Santalales (sister to the superrosids), Aquifoliales (sister to the Garryales), and Crossosomatales (sister to the fabids + remaining malvids) in our study (**Fig. 5C)** are consistent with the phylogenetic inference based on coalescent tree analysis of 410 single-copy gene families extracted from transcriptome and genome data (57).

Our results further provide interesting phylogenomic insights into the core eudicots. In our study, the Vitales was found to be sister to the Saxifragales with high support (**Fig. 5C)**, and this clade was sister to the remaining rosids (**Fig. 5C)**. These data are inconsistent with the hypothesis that the Vitales are sister to the Saxifragales and all other taxa in the superrosids clade (56–58,60,61,63). We also found that the Zygophyllales were sister to the malvids clade with high support (**Fig. 5C)**, inconsistent with its placement as sister of the Myrtales (57,61,63). These discrepancies can be mainly attributed to incomplete lineage sorting and ancient hybridization (64). Considering that our analyses involved ten thousand syntenic orthologous gene families, many more than than the hundreds or thousands of loci used in previous studies (56–63), and also that we have minimized the detrimental effects of the shared γ event, our results could represent a more accurate depiction of the real tree of life of the core eudicots than the other mentioned studies. Previous studies have used ortholog-based methods, which are prone to misidentify out-paralogs as orthologs, and we argue that our orthologous synteny-based method should be emphasized in plant phylogenomic studies. Nevertheless, our method is limited by taxon sampling (lack of high-quality genomes for some orders and families). However, this is expected to be resolved in the near future with the continuing development and effort in the field of genome sequencing (65).

### Limitations

The *Orthology Index* may not perform well in extremely complex cases. For instance, when Δ*T* is notably small (e.g. radiation following polyploidization in a few generations), similar to other methods (e.g. *Ks-*based), *OI* may also find it difficult to distinguish between out-paralogy and orthology (for a difficult case also see Fig. S68). Therefore, although we set a unified *OI* cutoff (0.6) in our pipeline, users should manually inspect the results (mainly dot plots) for confirmation, and the extremely complex cases showing unexpected patterns should be investigated on a case-by-case basis. This method is also limited in some scenarios where orthology inference and/or synteny detection is limited. For example, synteny is known not to be conserved in distantly-related or fast-evolving lineages (66), suggesting that this method should not be applied in cases without conserved synteny (e.g. angiosperms–ferns). Additionally, fragmented assemblies, as well as mis-assemblies, can disrupt synteny and subsequently reduce the efficiency of the method. However, this is likely to cease to be a concern in the near future with the fast development of sequencing and assembly techniques.

## Conclusions

In summary, we present a human-interpretable and machine-actionable approach to distinguish orthology from out-paralogy for syntenic blocks. The approach can identify orthologous synteny robustly, as validated with nearly 100 representative cases. We have demonstrated the broad and valuable applications of this approach to the reconstruction of evolutionary history in plant genomes, including reconstruction of the tree/network of life, and identification of and placing of polyploidy events on the tree/network. This approach will extend our analytical capacity in evolutionary genomics and might reduce the misleading data generated using some traditional methods.

## Code availability

The codes of the SOI tool and pipeline, and typical examples can be found on GitHub (https://github.com/zhangrengang/SOI and https://github.com/zhangrengang/evolution_example).

## Data availability

The codon alignments and gene trees of the core eudicots are available from Figshare (https://doi.org/10.6084/m9.figshare.24174930).

## Acknowledgments

We thank Professors Ying-Xiong Qiu, Xiao-Ming Song, Quan-Jun Hu, Tao Zhou, and Zhao-Ying Liu for generously sharing their genomic data, and the other researchers who have already released their genomic data publicly.

## Author contributions

RGZ and YPM conceived and designed the study; RGZ, HYS, MJZ and HS programmed and/or tested the code; RGZ collected and analyzed the data; RGZ and HYS prepared figures; RGZ, KHJ and HYS drafted the manuscript; YPM, RIM, FA, HC, KHJ and YVdP revised the manuscript; all authors approved the final manuscript.

## Supplementary Figures and Tables

**Figs. S1**–**90.** *Orthology Index* in the identification of orthologous synteny in species X and species Y. Refer to Fig. 1 for detailed descriptions.

**Fig. S91.** *Orthology Index*-colored dot plots showing orthologous syntenic relationships between *Centella asiatica* : *Vitis vinifera* (2:1 orthologous synteny depth ratio), *Aralia elata* : *Vitis vinifera* (2:1), *Angelica sinensis* : *Vitis vinifera* (4:1), and *Panax ginseng* : *Vitis vinifera* (4:1).

**Fig. S92.** Macro-synteny phylogenies inferred based on concatenated 1:2:2 (*Vitis vinifera* : *Aralia* elata : *Centella* asiatica) orthologous syntenic genes. The leaf labels on the trees are chromosome numbers. *n*, number of orthologous syntenic genes involved. Numbers at nodes represent the bootstrap values (percentage).

**Fig. S93.** *Orthology Index*-colored dot plots showing orthologous syntenic relationships between *Centella* asiatica and the Araliaceae (*Centella asiatica* : *Aralia* elata = 1:1, *Centella asiatica* : *Eleutherococcus senticosus* = 1:2, *Centella asiatica* : *Panax ginseng* = 1:2, *Centella asiatica* : *Panax notoginseng* = 1:1).

**Fig. S94.** *Orthology Index*-colored dot plots showing orthologous syntenic relationships between *Aralia* elata and the Apioideae (all 1:2).

**Fig. S95.** *Orthology Index*-colored dot plots showing orthologous syntenic relationships between *Centella asiatica* and the Apioideae (all 1:2).

**Fig. S96.** *Orthology Index*-colored dot plots showing orthologous syntenic relationships within the Apioideae (all 1:1).

**Fig. S97.** *Orthology Index*-colored dot plots showing orthologous syntenic relationships within the Araliaceae (*Panax ginseng* : *Panax notoginseng* = 2:1, *Aralia elata* : *Panax notoginseng* = 1:1, *Eleutherococcus senticosus* : *Panax notoginseng* = 2:1). The other orthologous syntenic depth ratios (*Aralia elata* : *Eleutherococcus senticosus* = 1:2, *Aralia elata* : *Panax ginseng* = 1:2, *Eleutherococcus senticosus* : *Panax ginseng* = 2:2) can be found in Fig. S57-59.

**Fig. S98.** *Orthology Index*-colored dot plots showing orthologous syntenic relationships between *Arabidopsis thaliana × A. arenosa* and *A. thaliana* + *A. arenosa*.

**Fig. S99.** *Orthology Index*-colored dot plots showing orthologous syntenic relationships between *Arachis hypogaea* and *A. duranensis* + *A. ipaensis*.

**Fig. S100.** *Orthology Index*-colored dot plots showing orthologous syntenic relationships between *Triticum turgidum* and *T. aestivum*.

**Fig. S101.** *Orthology Index*-colored dot plots showing orthologous syntenic relationships between *Papaver somniferum* and *P. setigerum*.

**Fig. S102.** Phylogenetic relationships within the core eudicots based on 12,277 multi-copy SOGs. The numbers at the nodes are posterior probabilities from ASTRAL. Bar, 3.0 coalescent units.

**Fig. S103.** Phylogenetic relationships within the core eudicots based on 5,154 single-copy SOGs. The numbers at the nodes are posterior probabilities from ASTRAL. Bar, 3.0 coalescent units.

**Table S1.** Cases with shared polyploidization event(s) used in this study.

**Table S2.** Genomic data used in this study (accessed before 2023-06-29).

