## Supplemental Tables for "SOI: Robust identification of orthologous synteny with the *Orthology Index* and broad applications in evolutionary genomics"

Ren-Gang Zhang *et al.*

**This PDF file includes:**

Figs. S1 to S103
Tables S1 to S2
References (S1 to S206)

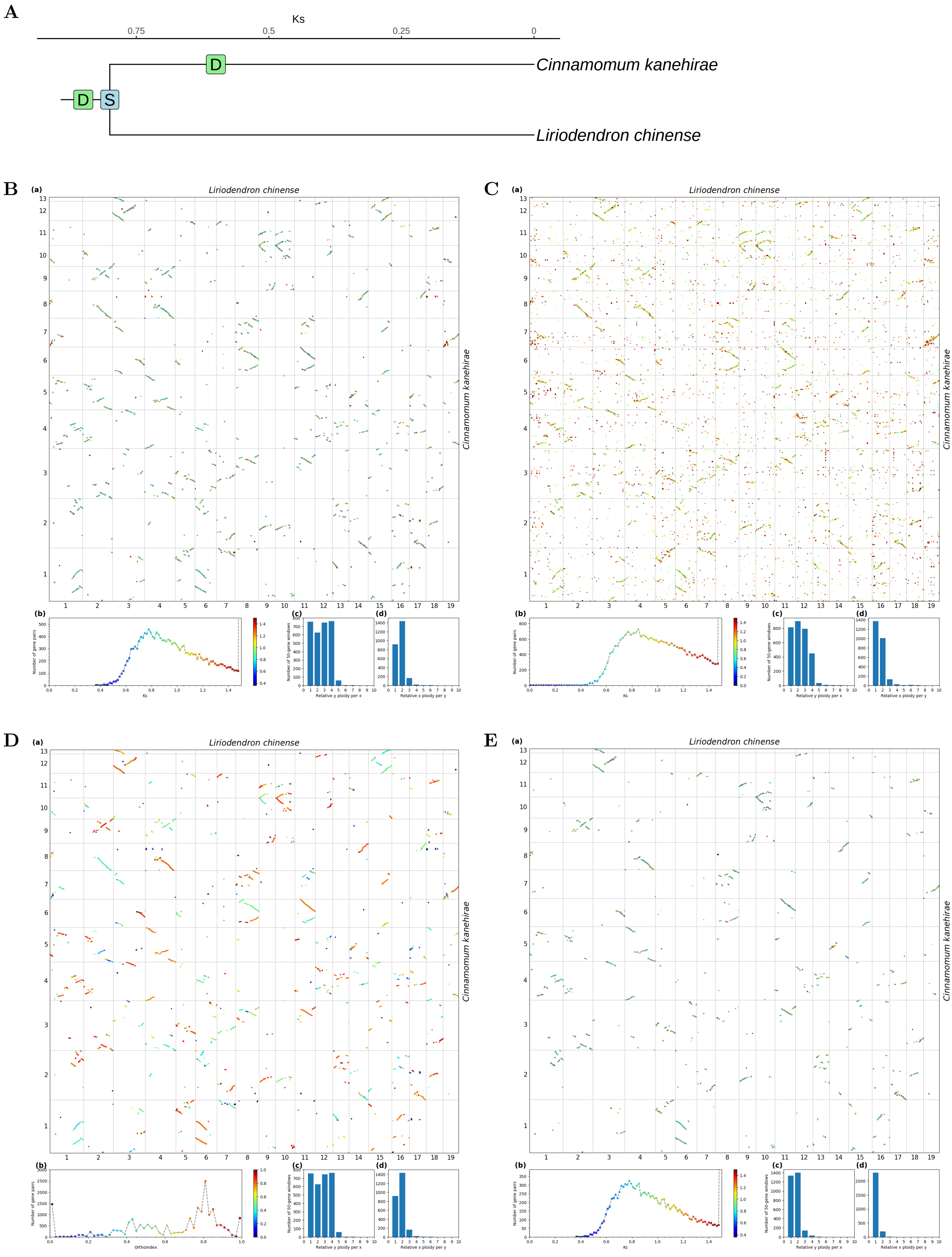


**Fig S1.** ***Orthology Index* in the identification of orthologous synteny in *Liriodendron chinense* and *Cinnamomum kanehirae*.** Refer to **Fig.1** for detailed descriptions.

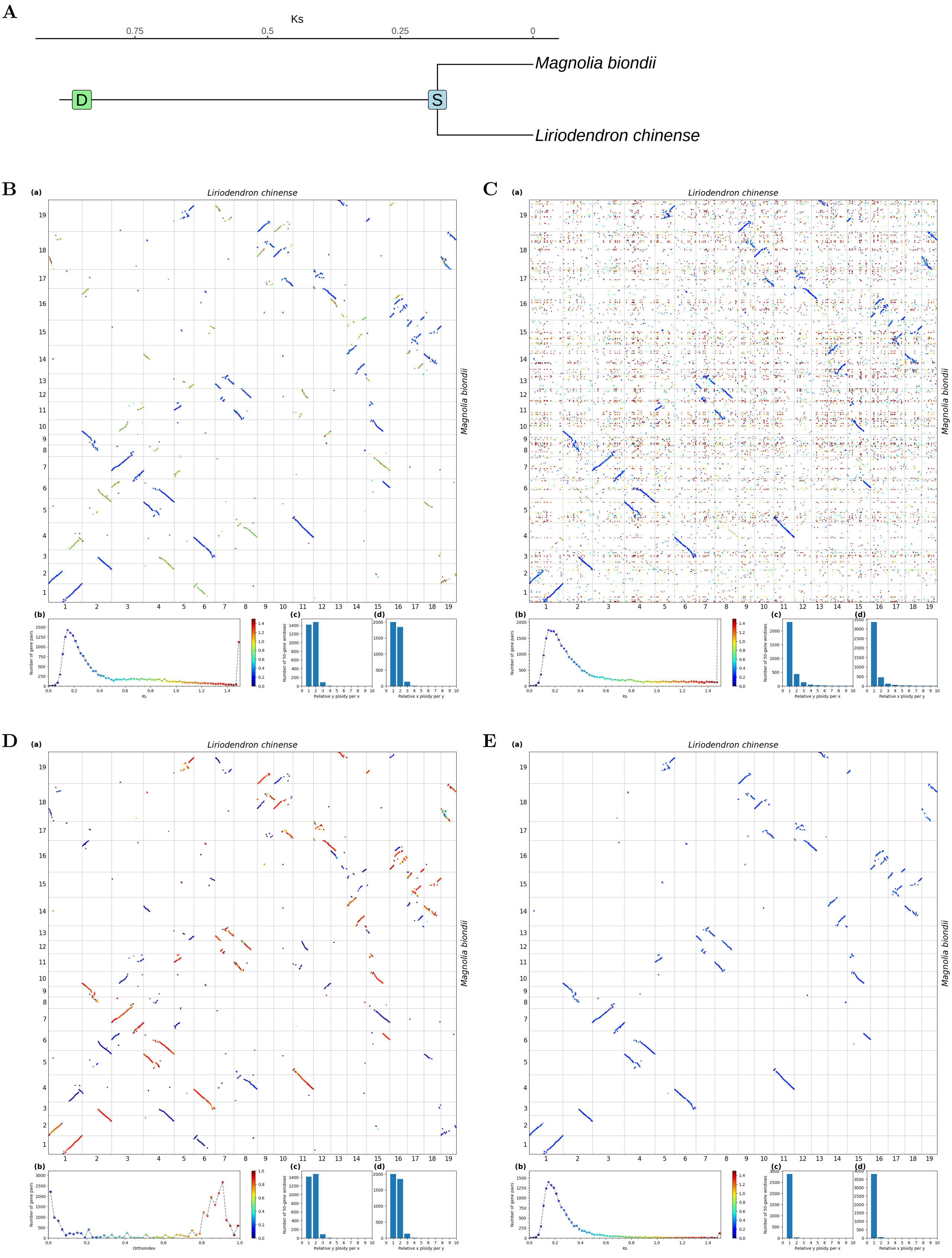


**Fig S2.** ***Orthology Index*** **in the identification of orthologous synteny in** ***Liriodendron chinense*** **and** ***Magnolia biondii.*** Refer to **Fig.1** for detailed descriptions.

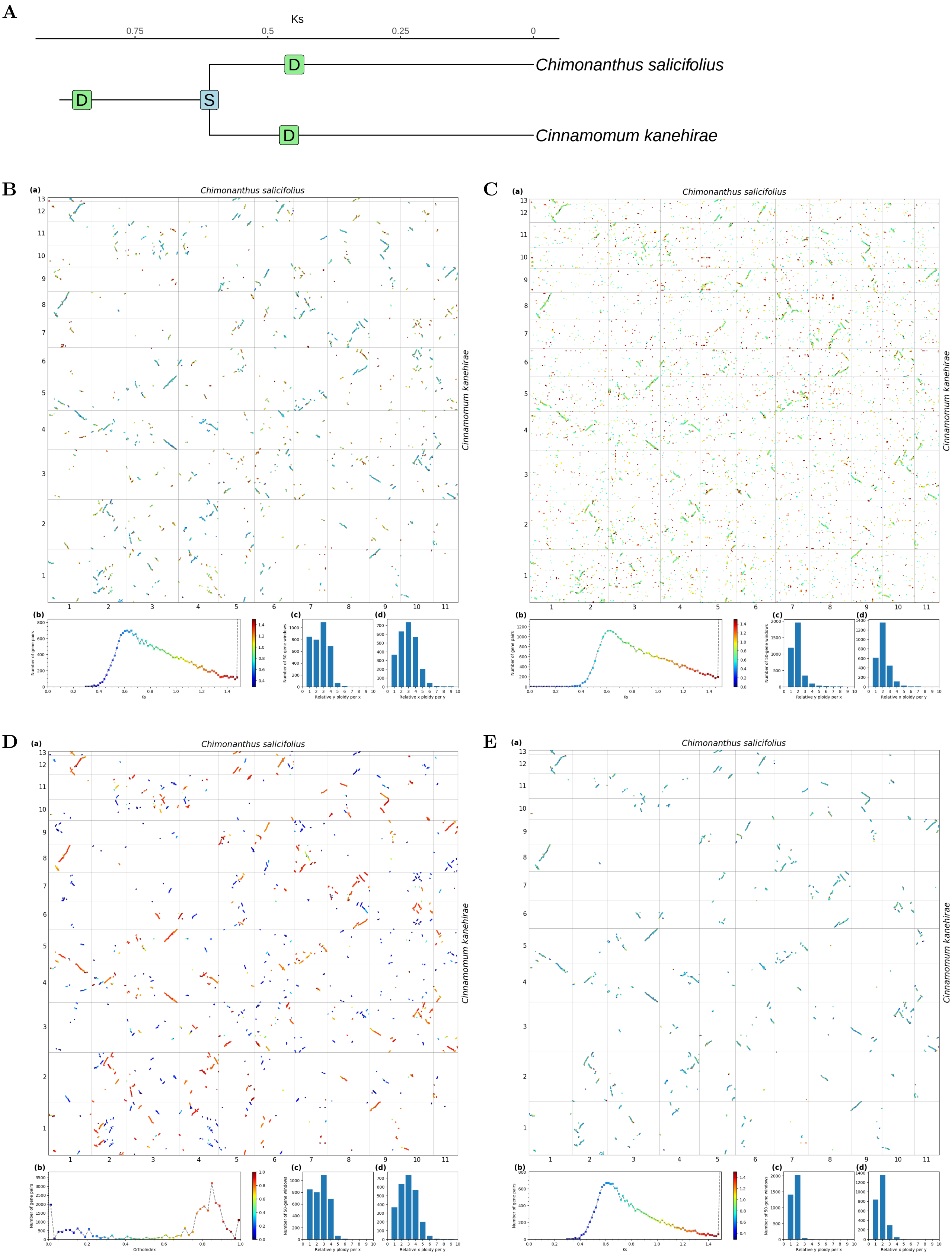


**Fig S3.** ***Orthology Index*** **in the identification of orthologous synteny in** ***Cinnamomum kanehirae*** **and** ***Chimonanthus salicifolius.*** Refer to **Fig.1** for detailed descriptions.

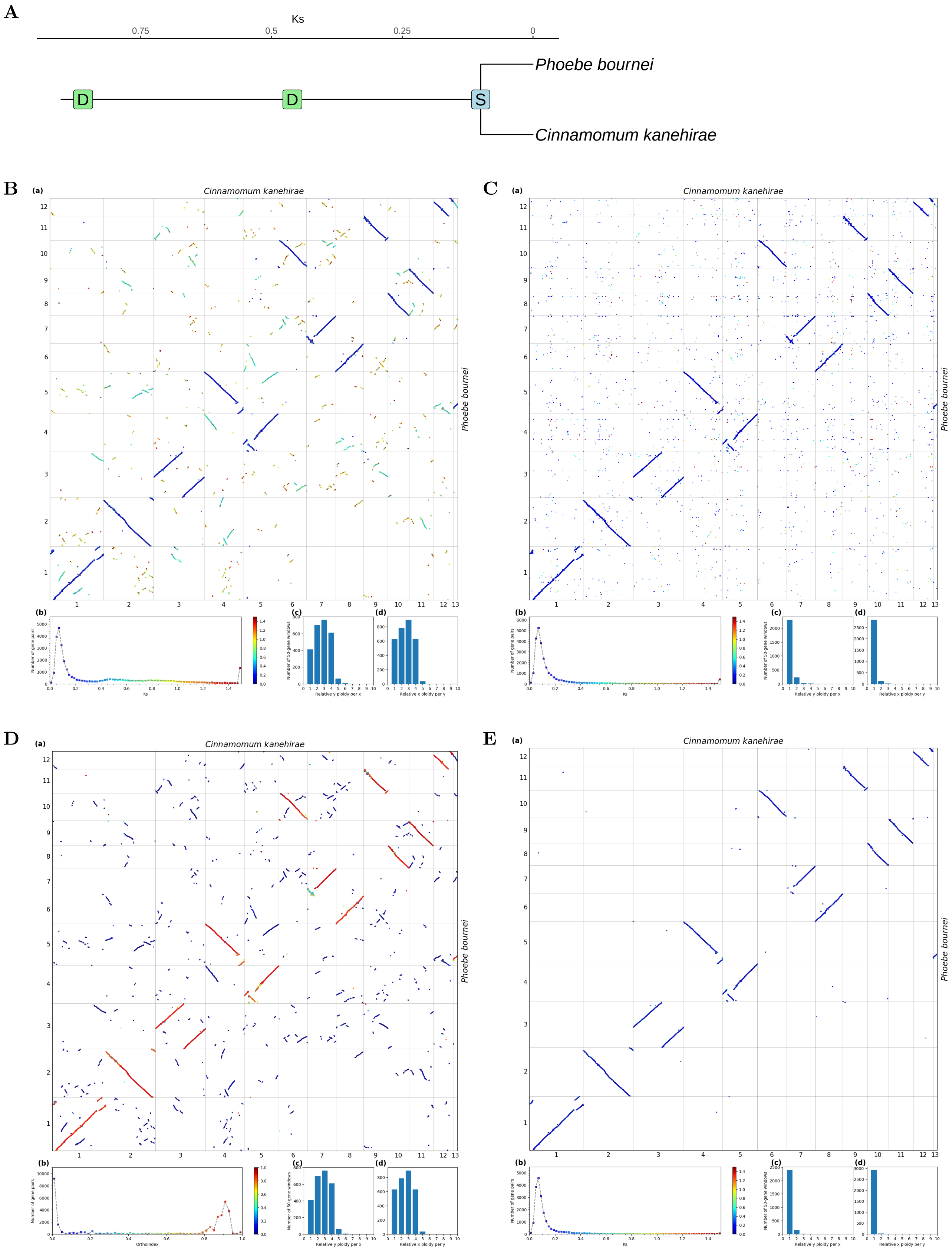


**Fig S4.** ***Orthology Index* in the identification of orthologous synteny in *Cinnamomum kanehirae* and *Phoebe bournei*.** Refer to **Fig.1** for detailed descriptions.

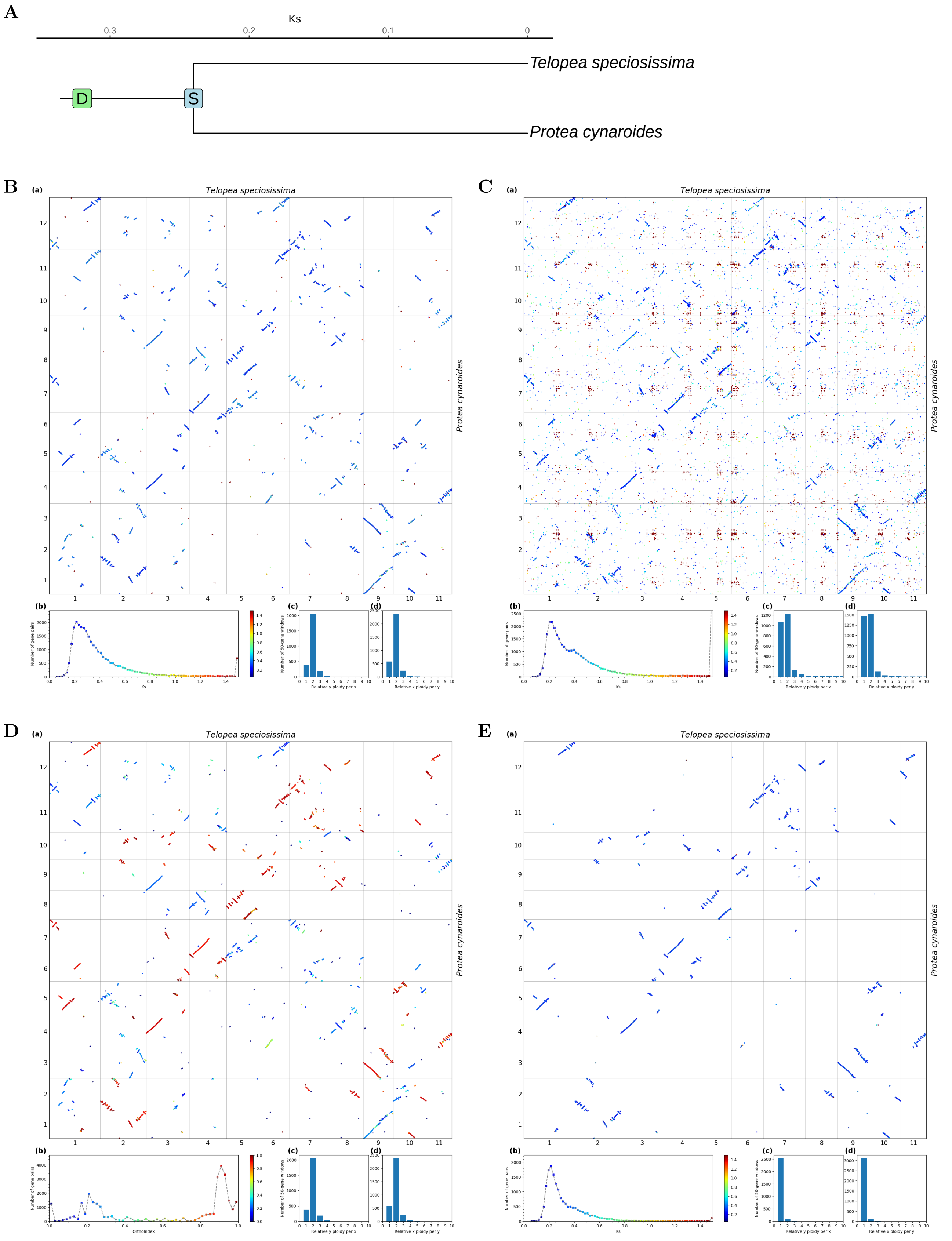


**Fig S5.** ***Orthology Index* in the identification of orthologous synteny in *Protea cynaroides* and *Telopea speciosissima*.** Refer to **Fig.1** for detailed descriptions.

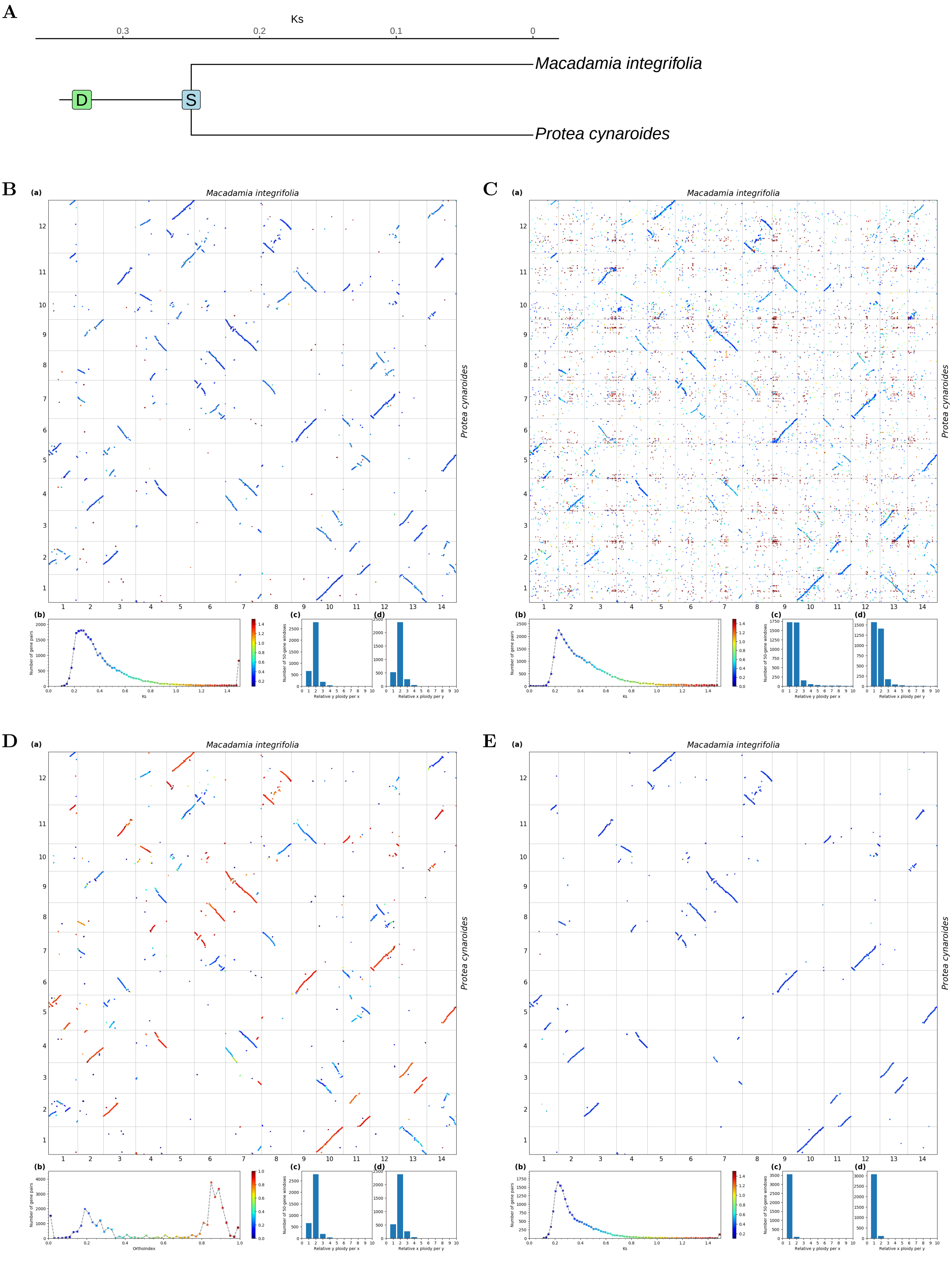


**Fig S6.** ***Orthology Index* in the identification of orthologous synteny in *Protea cynaroides* and *Macadamia integrifolia*.** Refer to **Fig.1** for detailed descriptions.

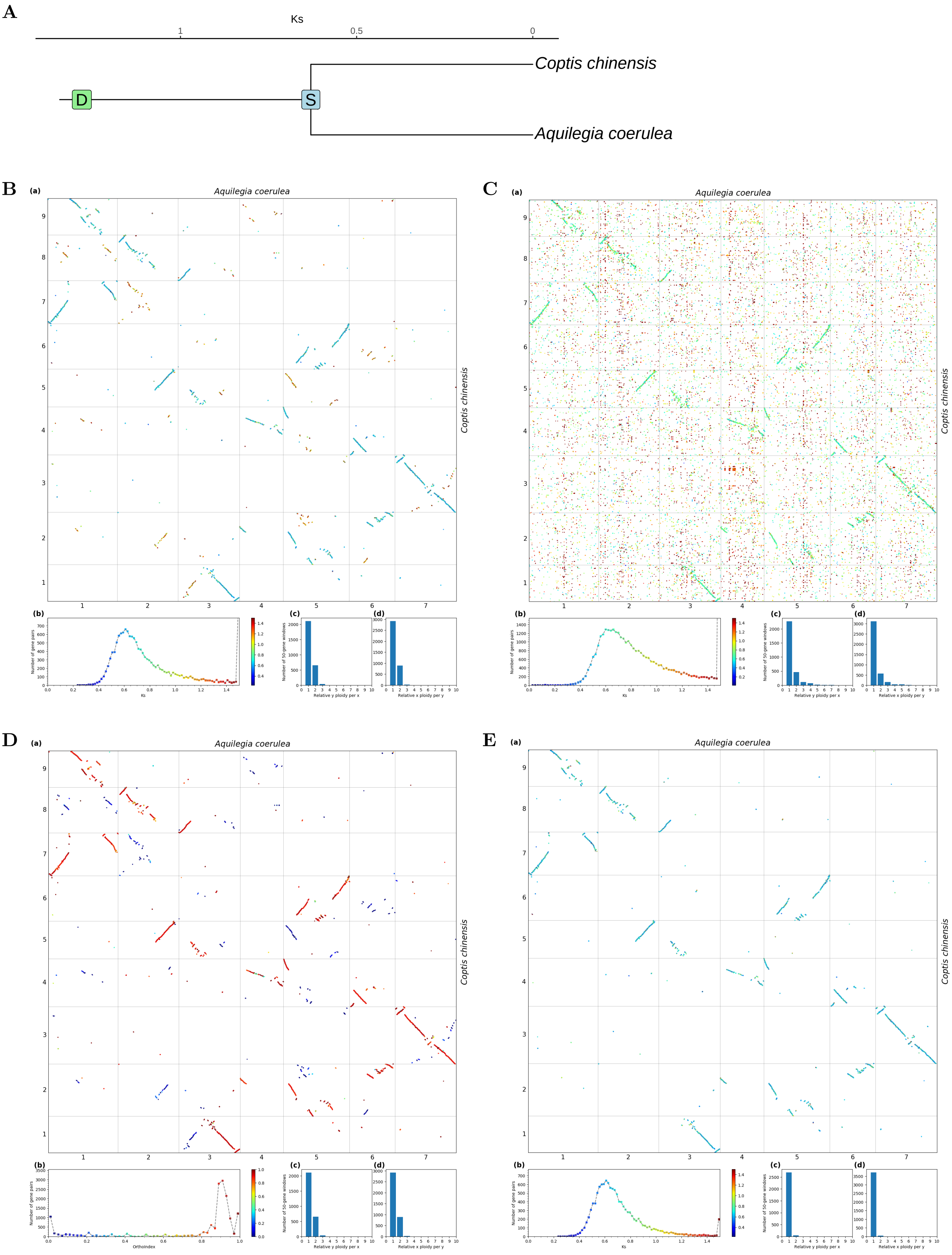


**Fig S7.** ***Orthology Index*** **in the identification of orthologous synteny in *Aquilegia coerulea* and *Coptis chinensis*.** Refer to **Fig.1** for detailed descriptions.

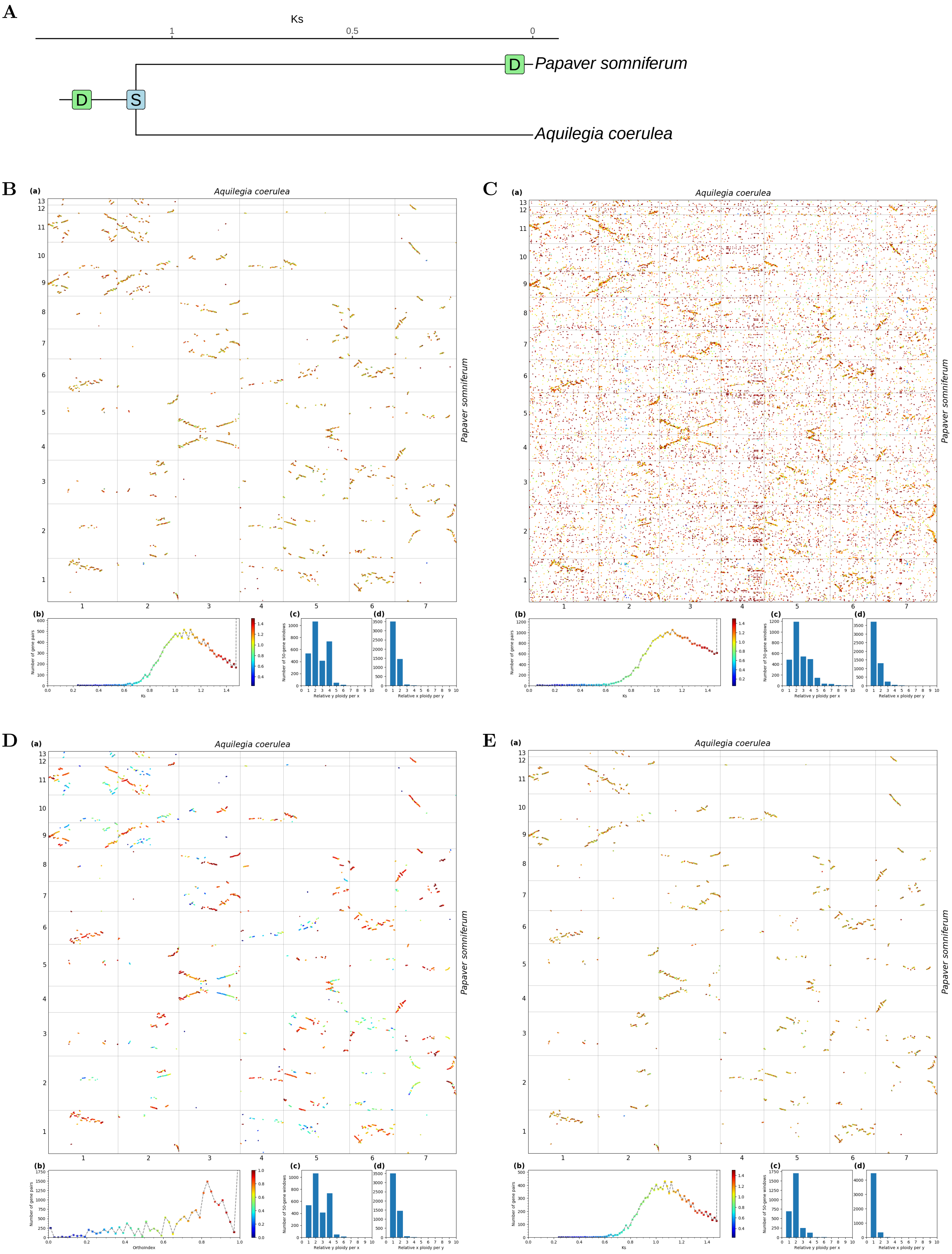


**Fig S8.** ***Orthology Index*** **in the identification of orthologous synteny in** ***Aquilegia coerulea*** **and** ***Papaver somniferum*****.** Refer to **Fig.1** for detailed descriptions.

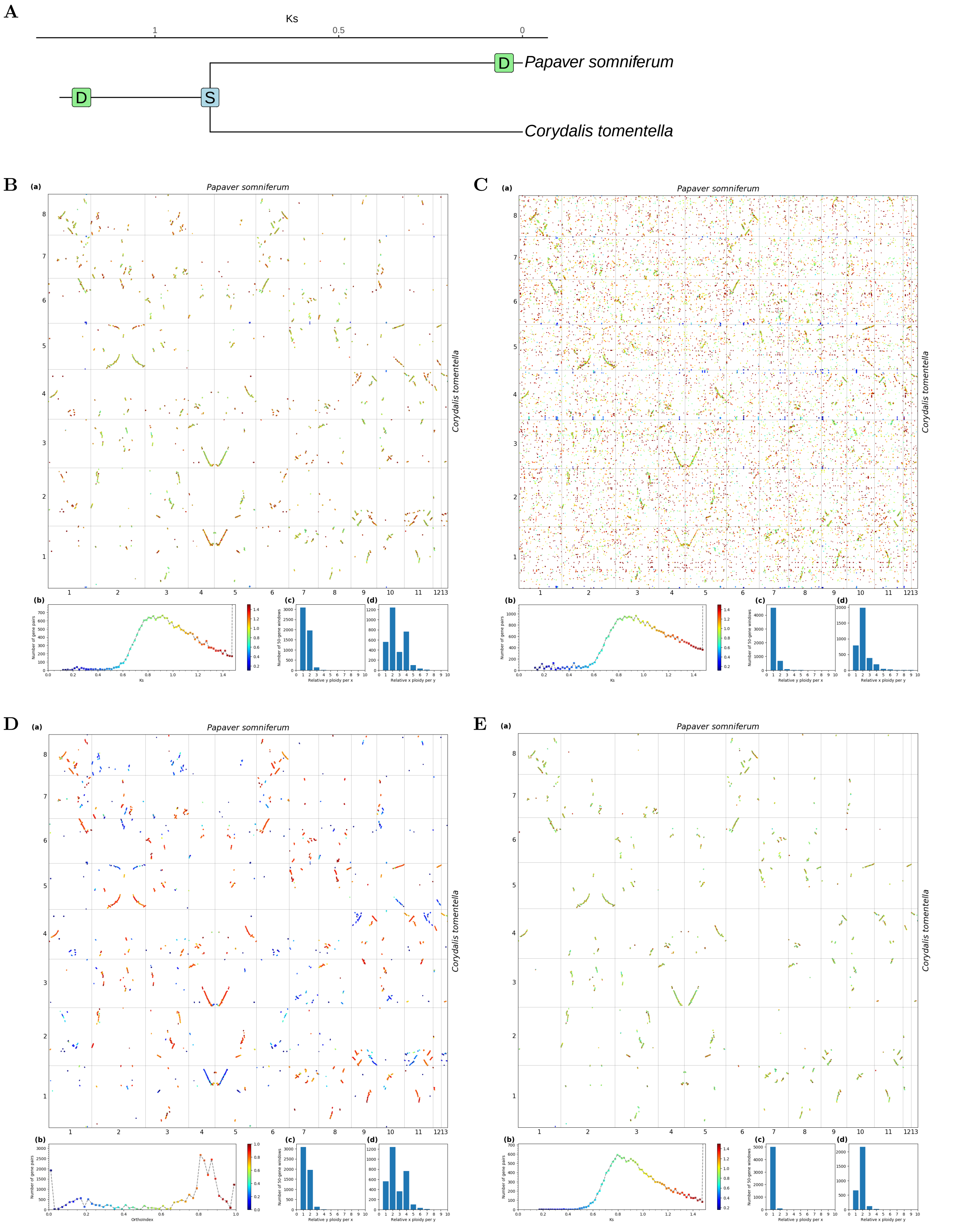


**Fig S9.** ***Orthology Index*** **in the identification of orthologous synteny in** ***Corydalis tomentella*** **and** ***Papaver somniferum*****.** Refer to **Fig.1** for detailed descriptions.

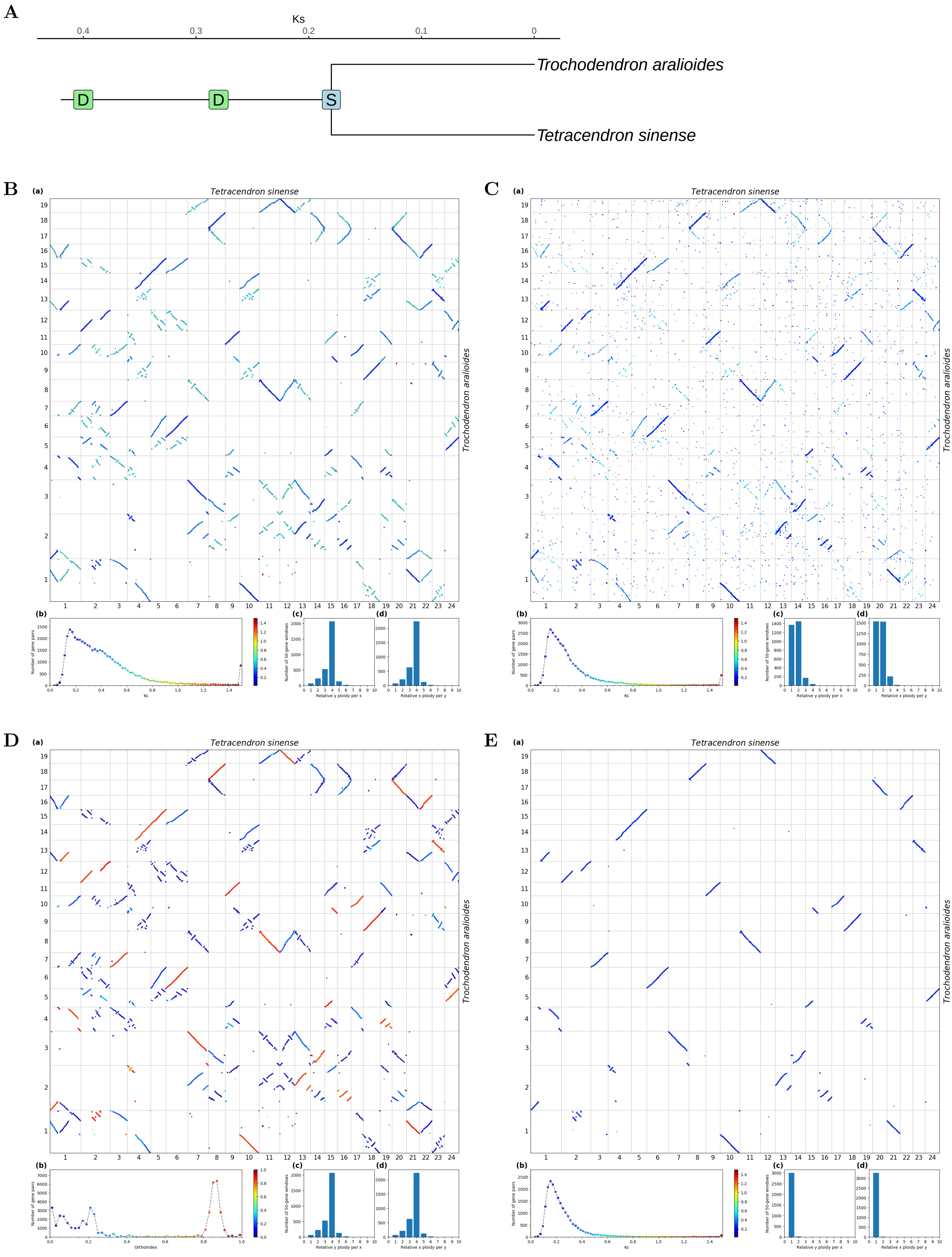


**Fig S10.** ***Orthology Index*** **in the identification of orthologous synteny in** ***Tetracendron sinense*** **and** ***Trochodendron aralioides*****.** Refer to **Fig.1** for detailed descriptions.

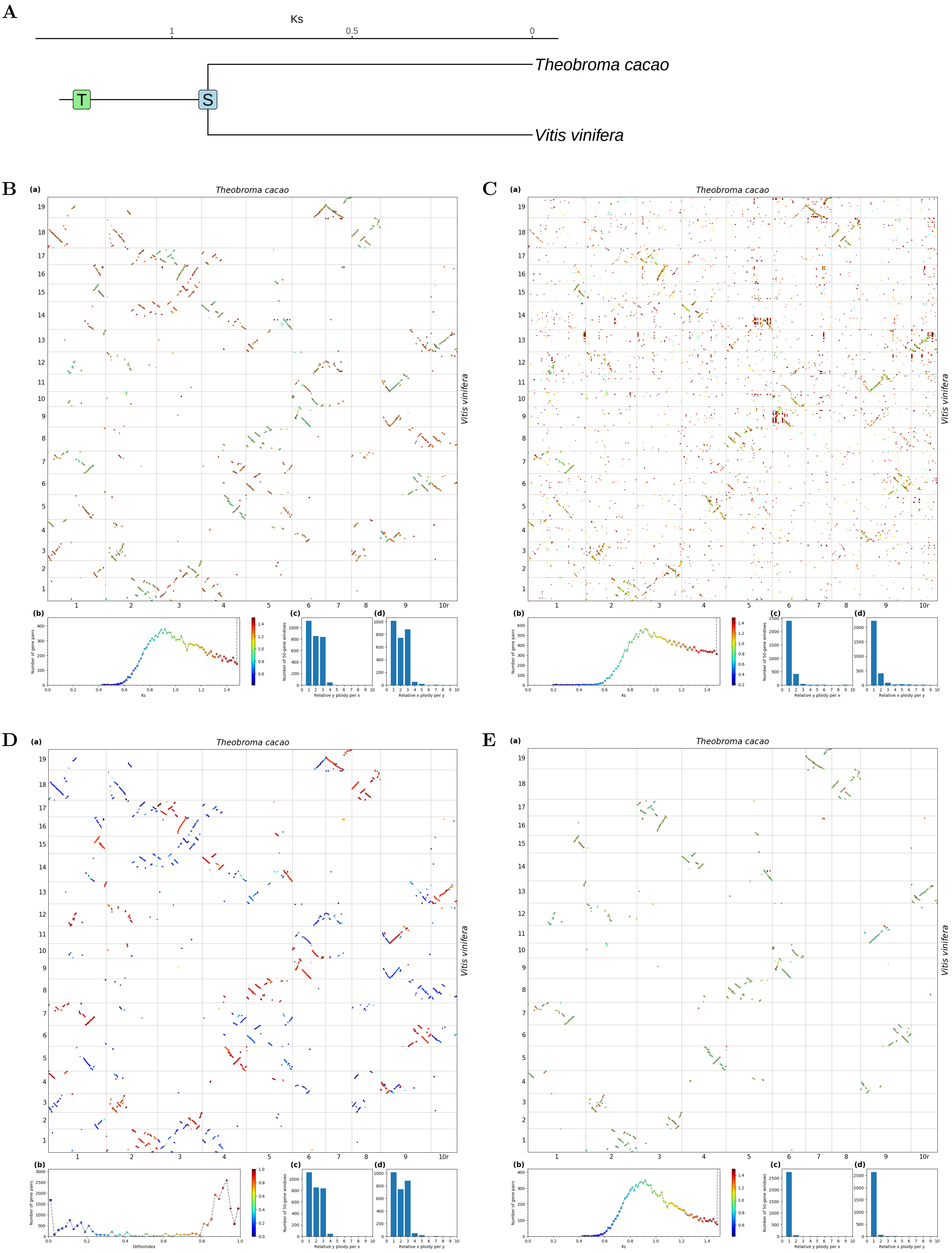


**Fig S11.** ***Orthology Index* in the identification of orthologous synteny in *Vitis vinifera* and *Theobroma cacao*.** Refer to **Fig.1** for detailed descriptions.

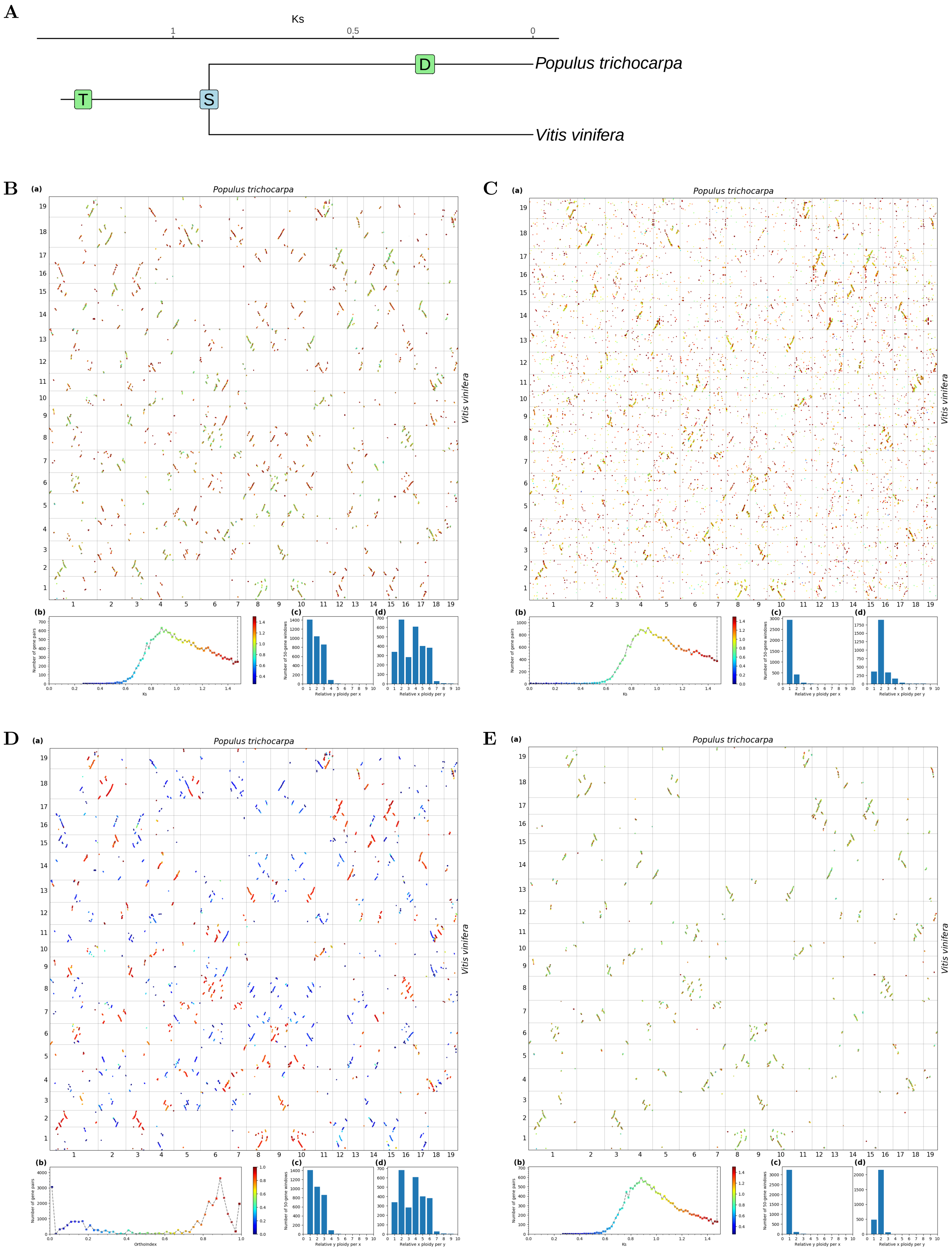


**Fig S12.** ***Orthology Index* in the identification of orthologous synteny in *Vitis vinifera* and *Populus trichocarpa*.** Refer to **Fig.1** for detailed descriptions.

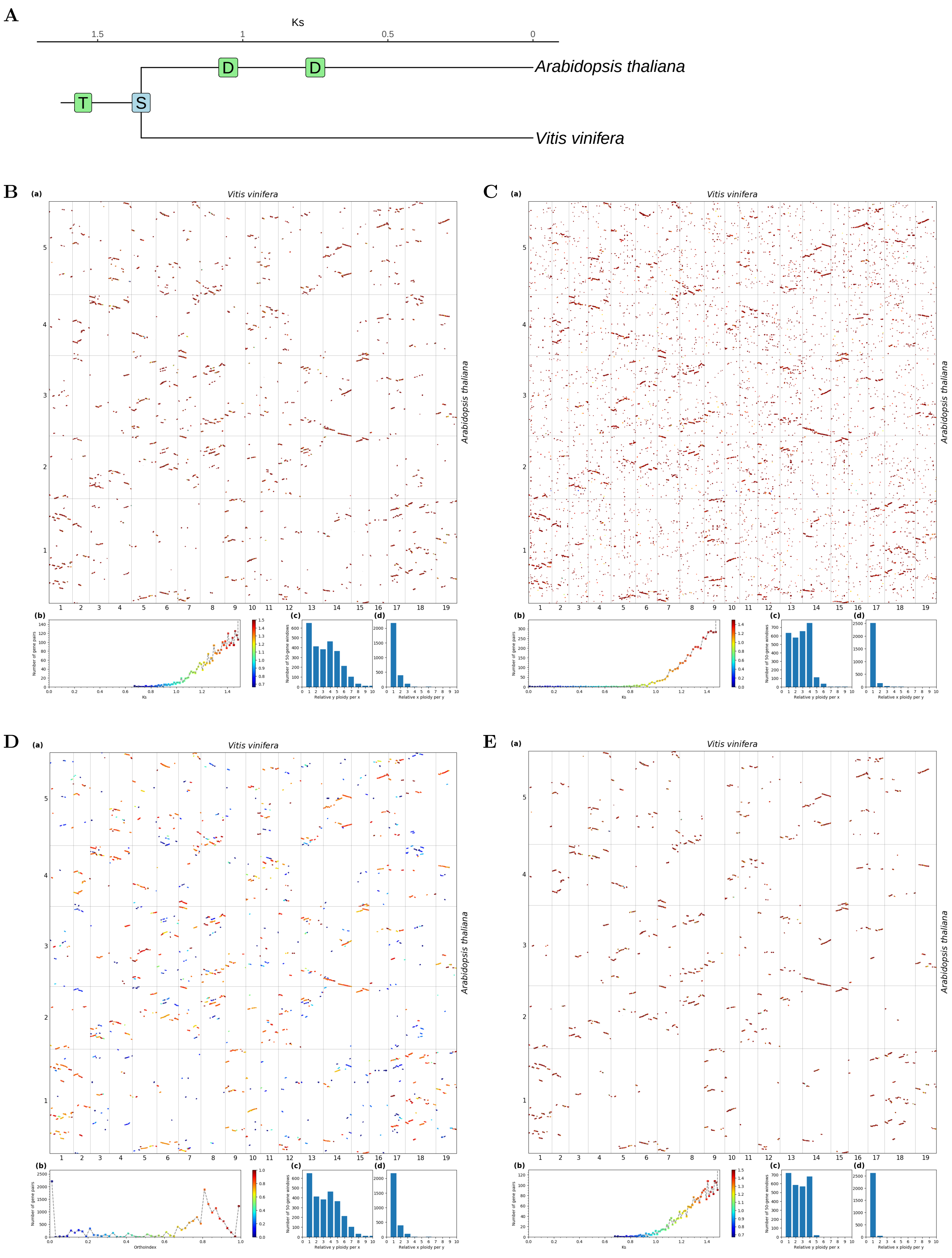


**Fig S13.** ***Orthology Index*** **in the identification of orthologous synteny in** ***Vitis vinifera*** **and** ***Arabidopsis thaliana.*** Refer to **Fig.1** for detailed descriptions.

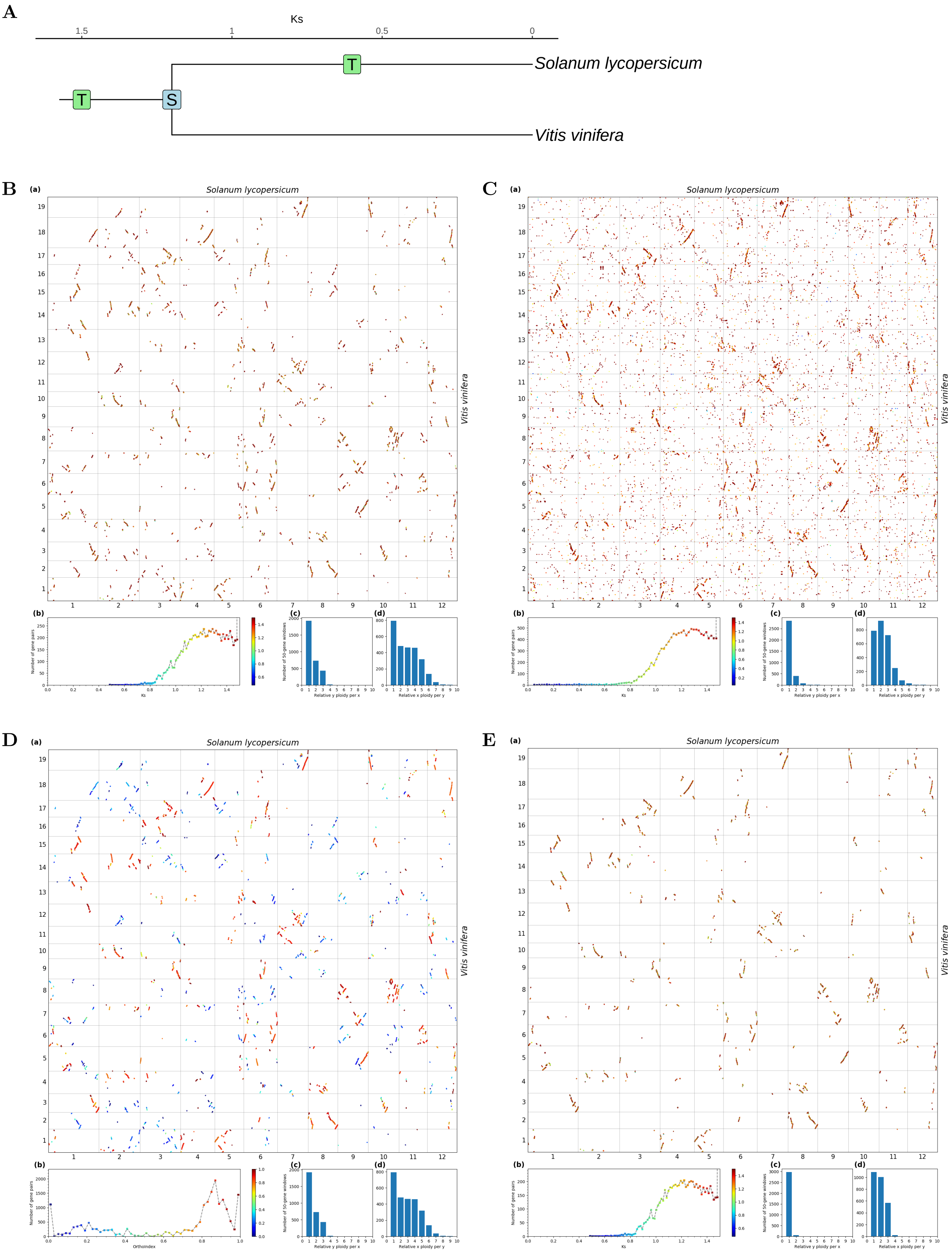


**Fig S14.** ***Orthology Index* in the identification of orthologous synteny in *Vitis vinifera* and *Solanum lycopersicum*.** Refer to **Fig.1** for detailed descriptions.

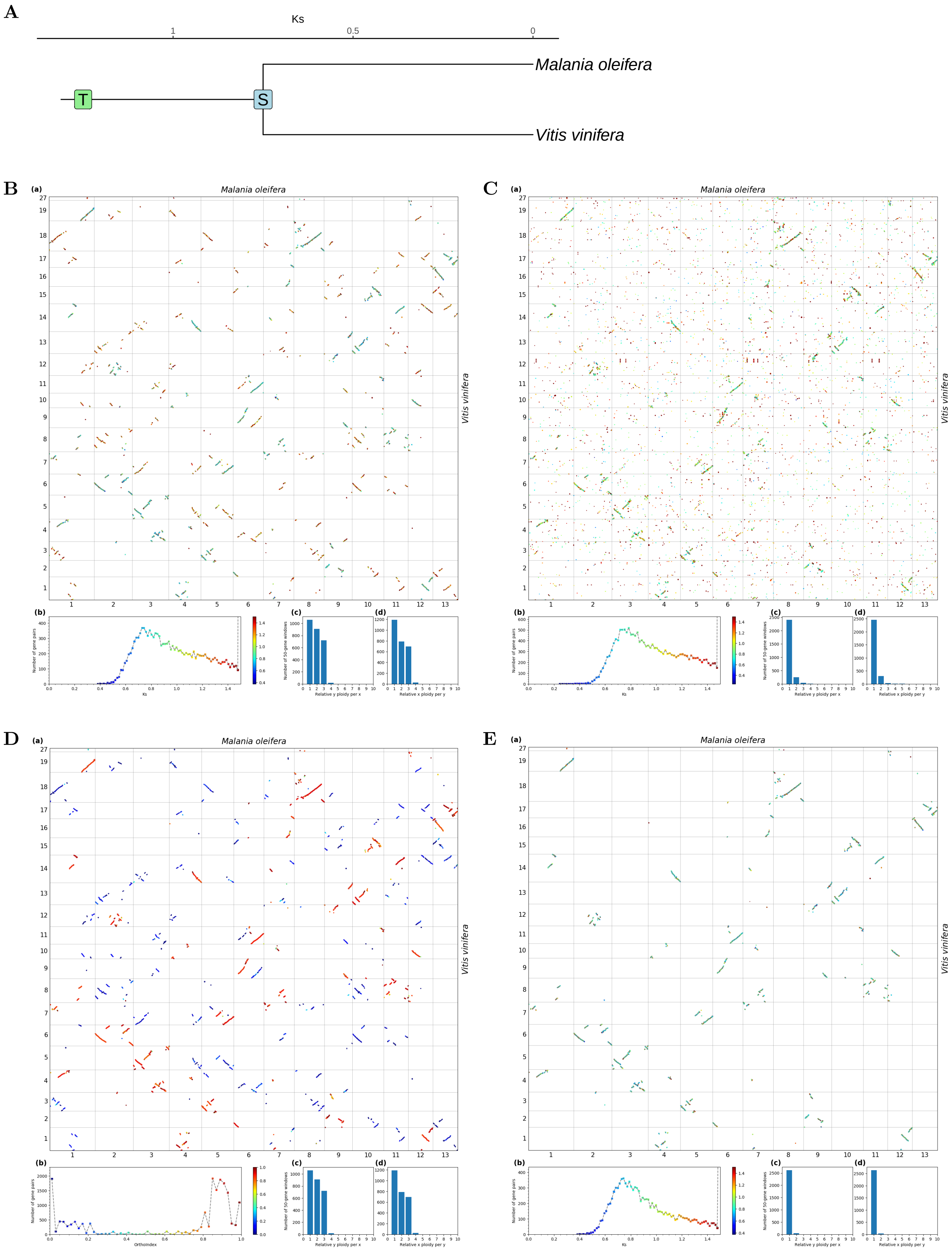


**Fig S15.** ***Orthology Index*** **in the identification of orthologous synteny in** ***Vitis vinifera*** **and** ***Malania oleifera*****.** Refer to **Fig.1** for detailed descriptions.

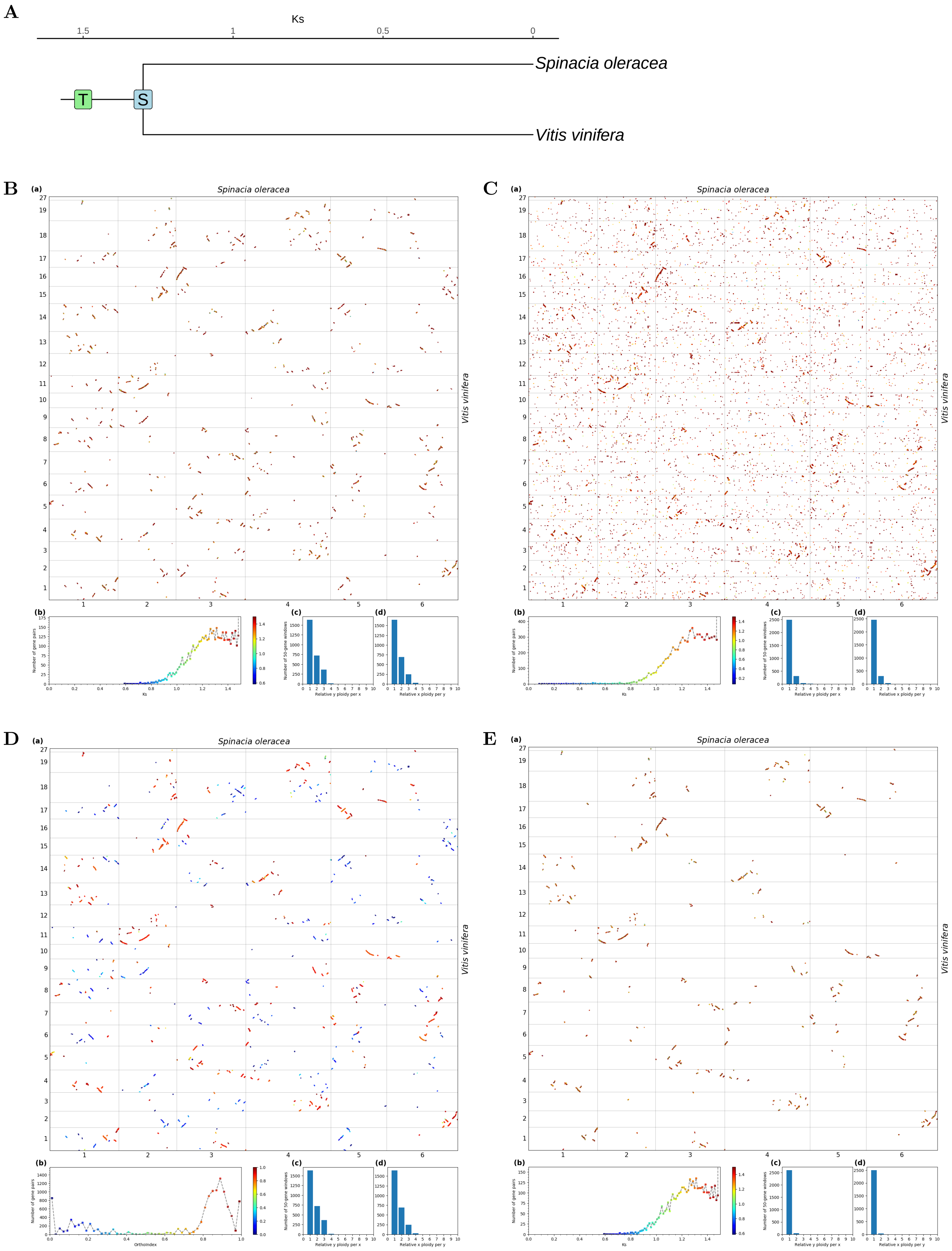


**Fig S16.** ***Orthology Index*** **in the identification of orthologous synteny in** ***Vitis vinifera*** **and** ***Spinacia oleracea.*** Refer to **Fig.1** for detailed descriptions.

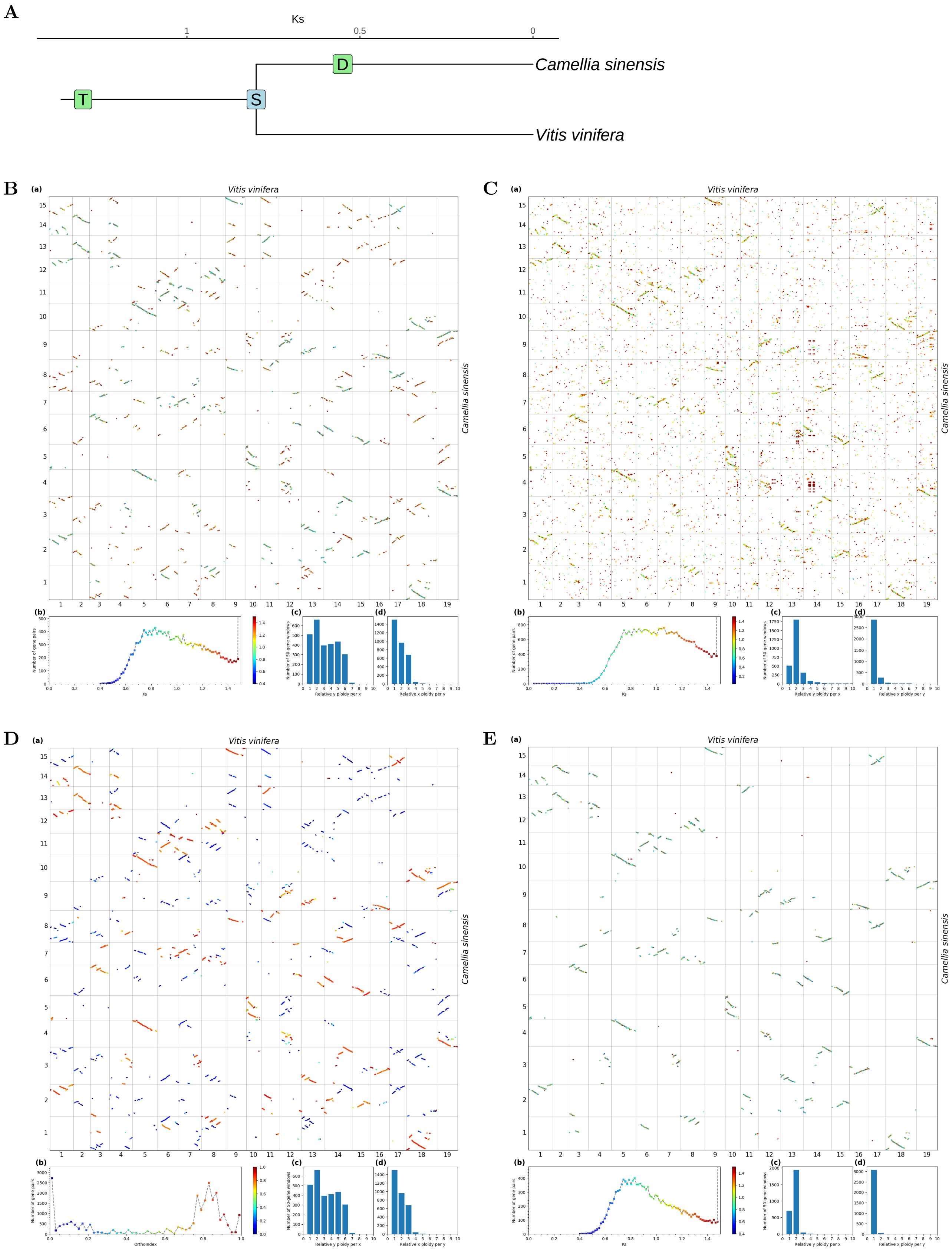


**Fig S17.** ***Orthology Index* in the identification of orthologous synteny in *Vitis vinifera* and *Camellia sinensis*.** Refer to **Fig.1** for detailed descriptions.

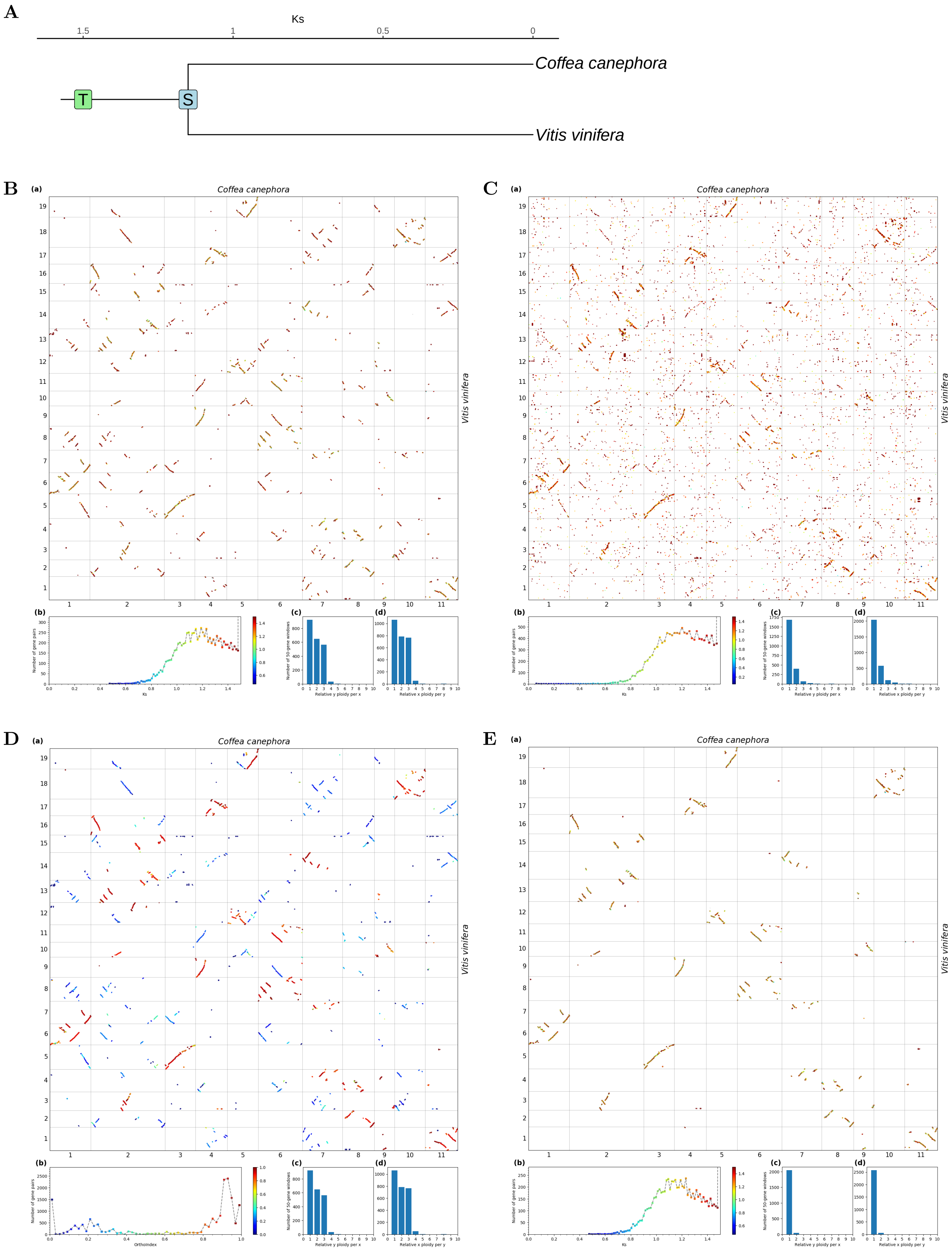


**Fig S18.** ***Orthology Index*** **in the identification of orthologous synteny in** ***Vitis vinifera*** **and** ***Coffea canephora.*** Refer to **Fig.1** for detailed descriptions.

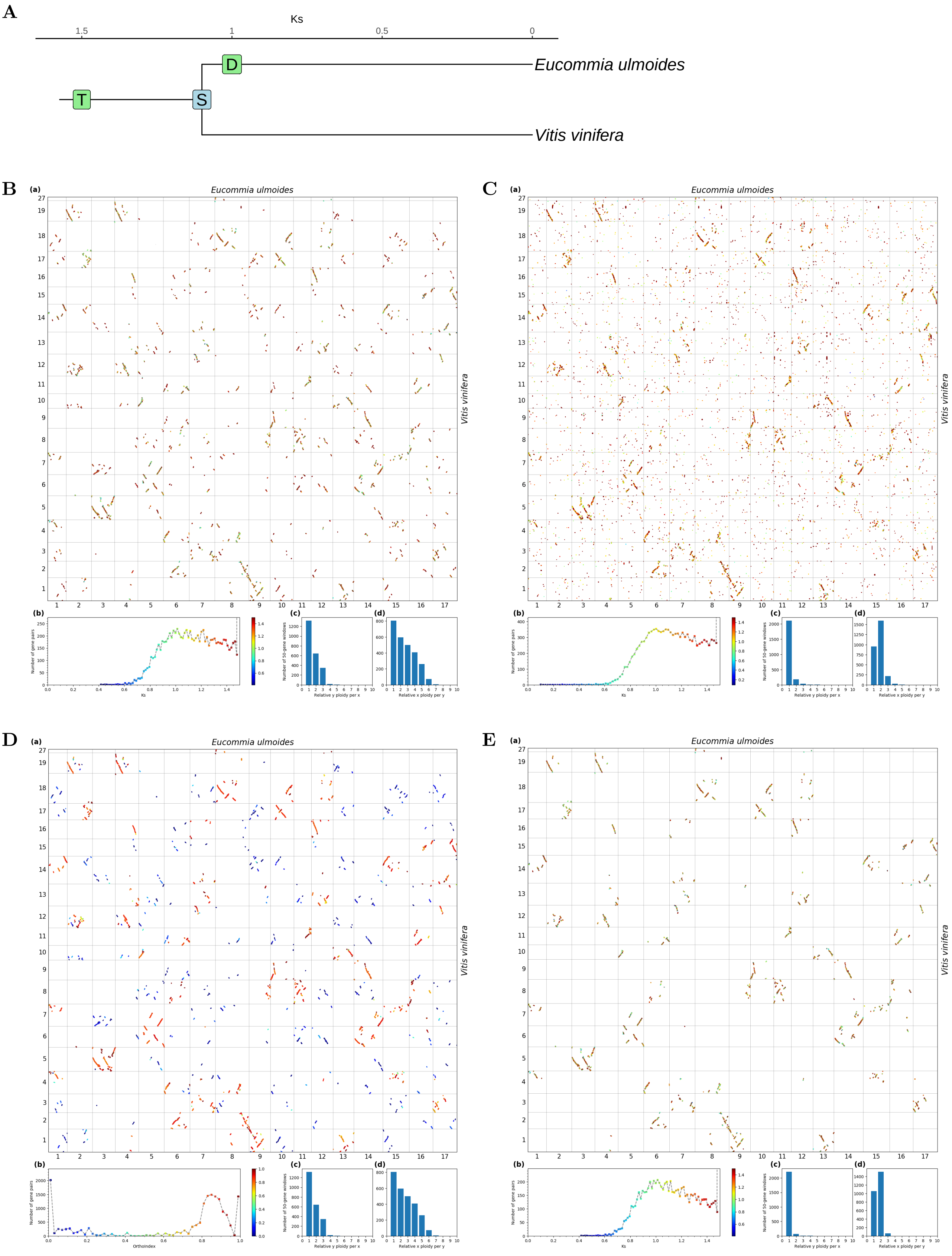


**Fig S19.** ***Orthology Index*** **in the identification of orthologous synteny in** ***Vitis vinifera*** **and** ***Eucommia ulmoides*.** Refer to **Fig.1** for detailed descriptions.

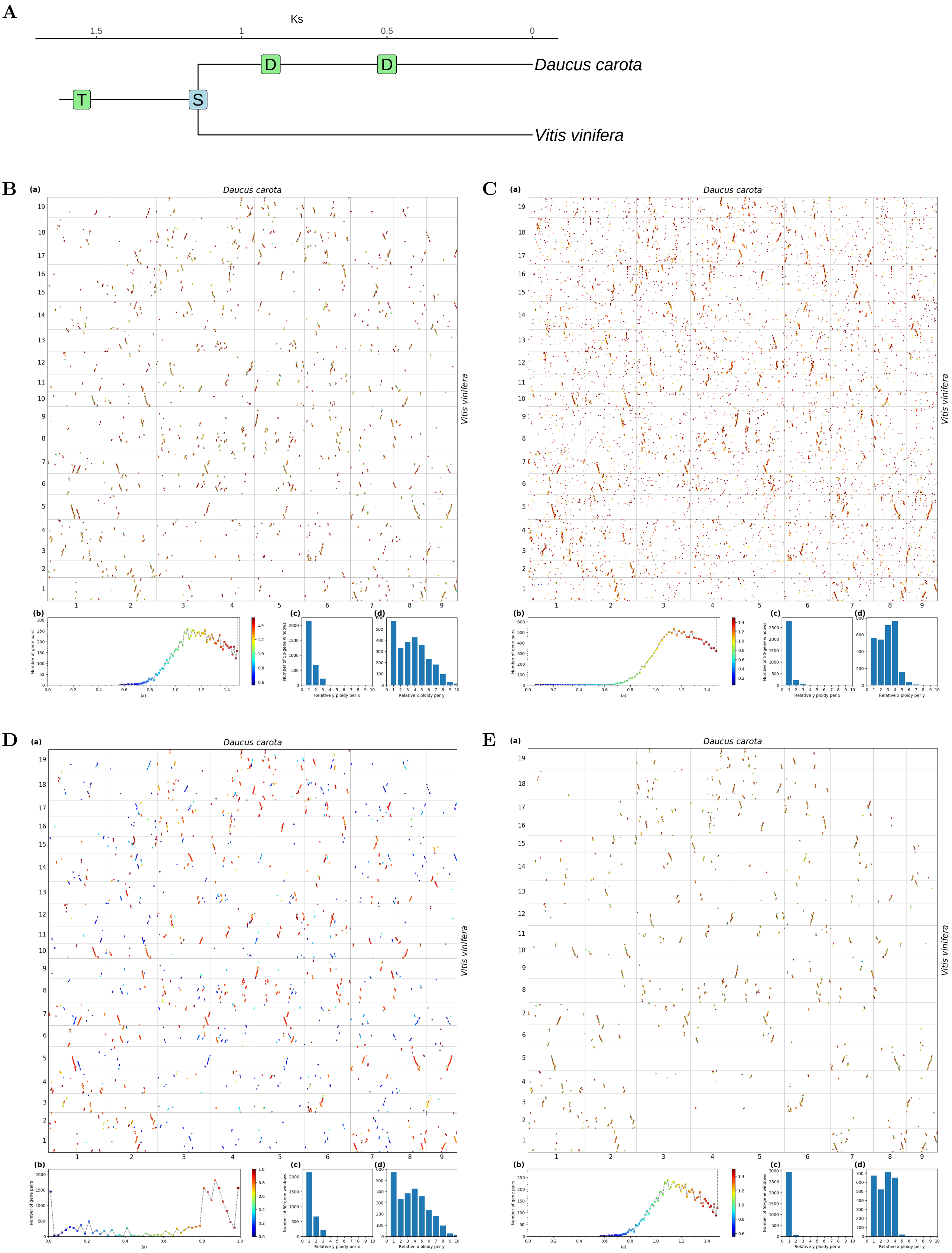


**Fig S20.** ***Orthology Index* in the identification of orthologous synteny in *Vitis vinifera* and *Daucus carota*.** Refer to **Fig.1** for detailed descriptions.

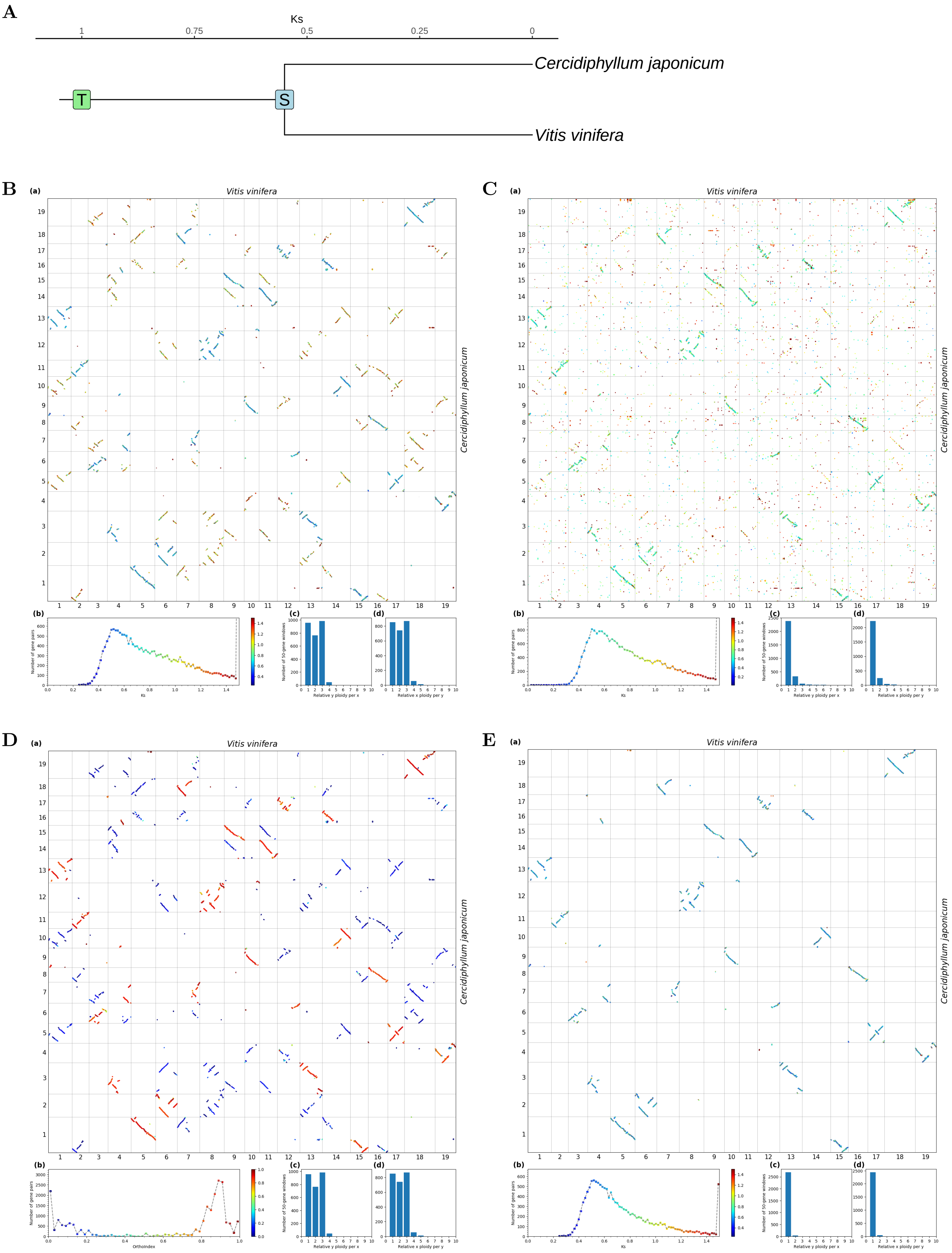


**Fig S21.** ***Orthology Index*** **in the identification of orthologous synteny in** ***Vitis vinifera*** **and** ***Cercidiphyllum japonicum.*** Refer to **Fig.1** for detailed descriptions.

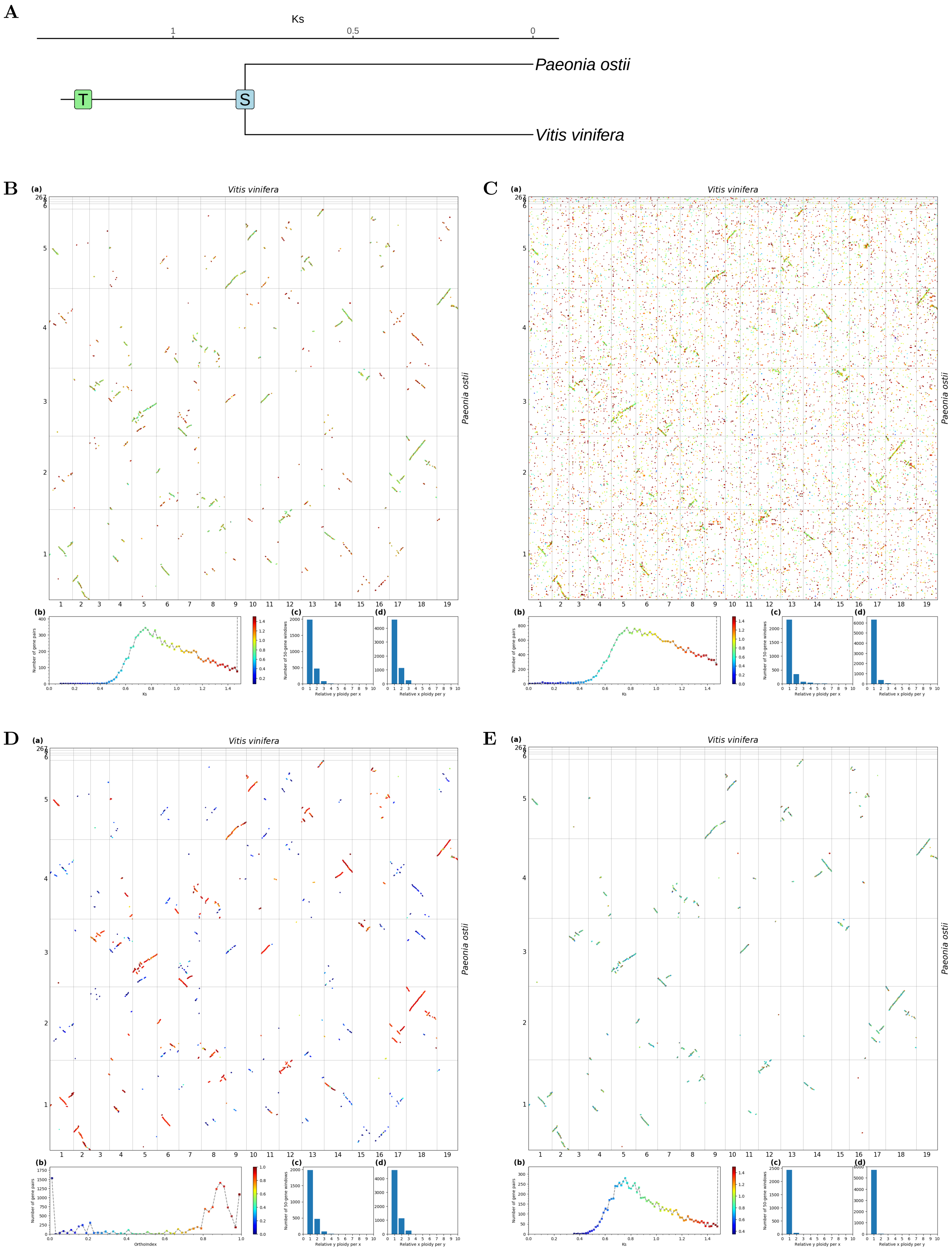


**Fig S22.** ***Orthology Index* in the identification of orthologous synteny in *Vitis vinifera* and *Paeonia ostii*.** Refer to **Fig.1** for detailed descriptions.

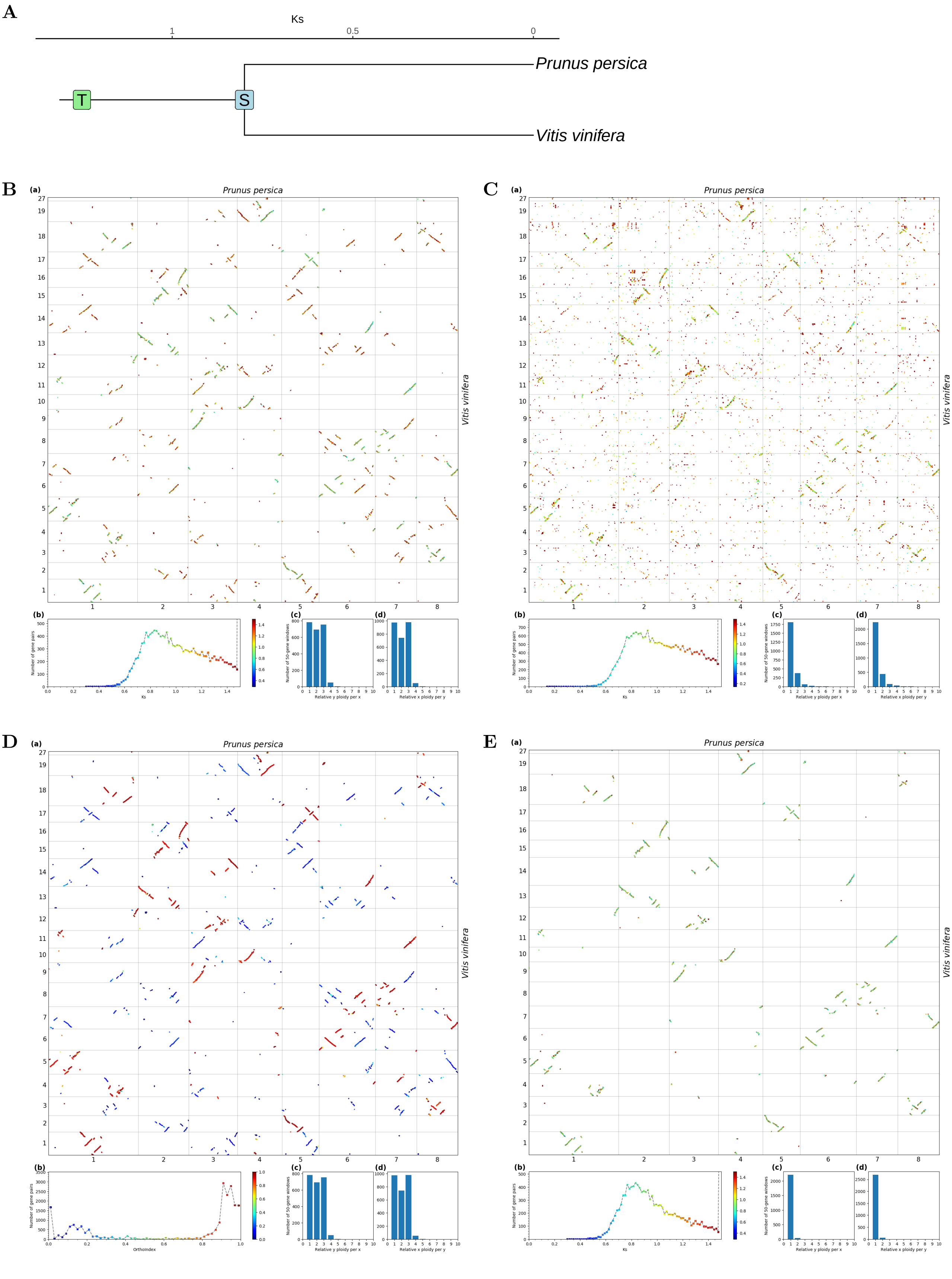


**Fig S23.** ***Orthology Index*** **in the identification of orthologous synteny in** ***Vitis vinifera*** **and** ***Prunus persica.*** Refer to **Fig.1** for detailed descriptions.

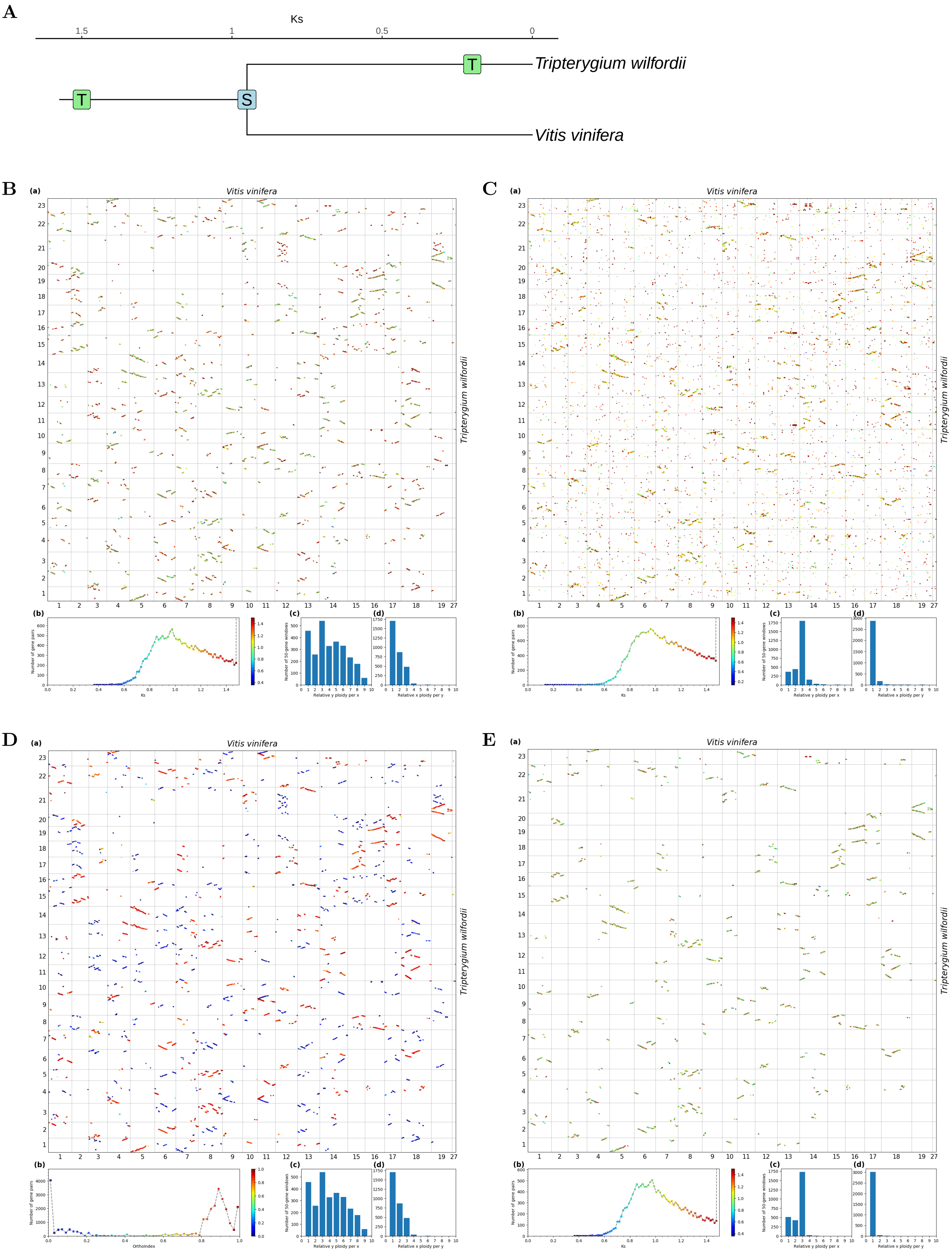

**Fig S24.** ***Orthology Index*** **in the identification of orthologous synteny in** ***Vitis vinifera*** **and** ***Tripterygium wilfordii.*** Refer to **Fig.1** for detailed descriptions.

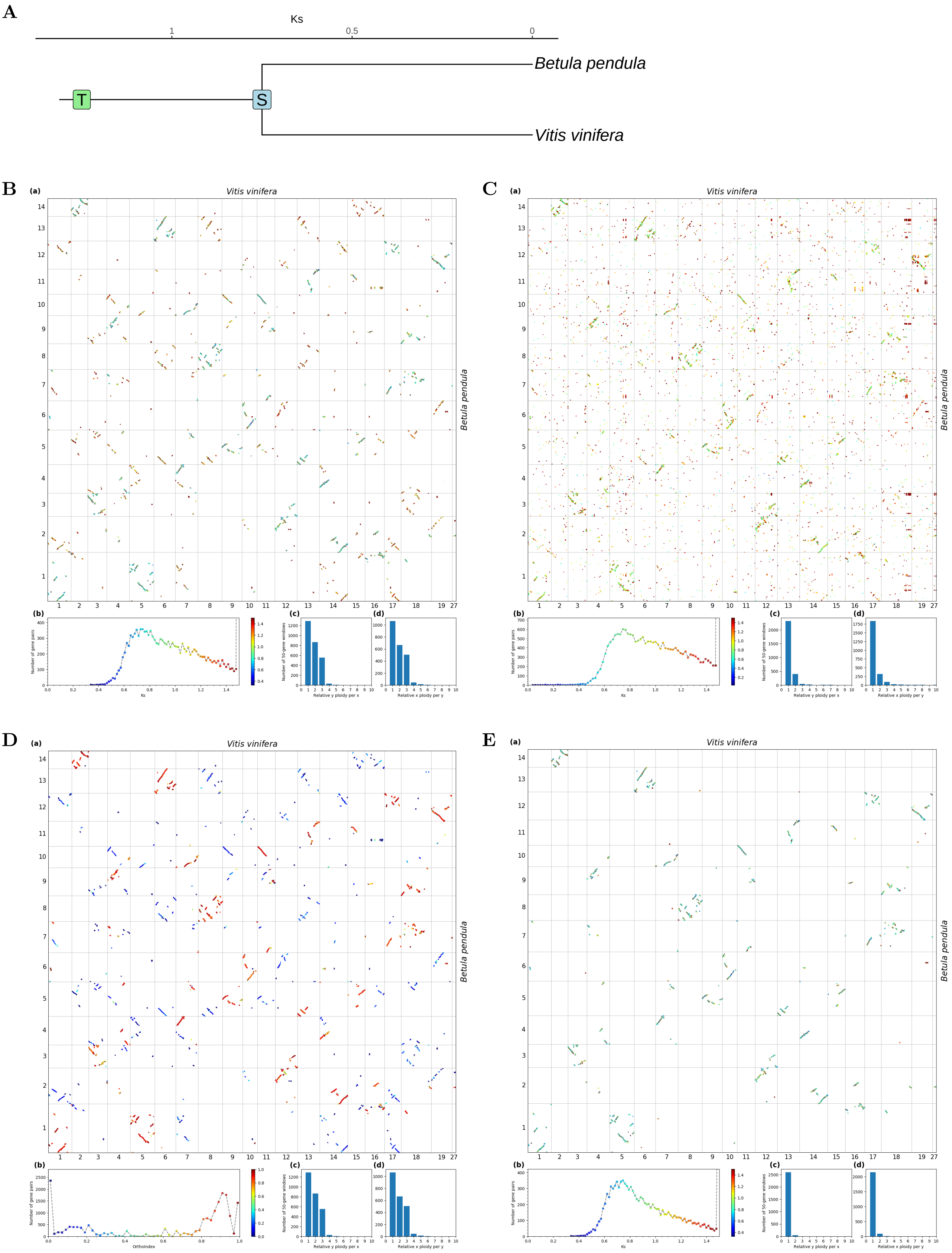


**Fig S25.** ***Orthology Index*** **in the identification of orthologous synteny in** ***Vitis vinifera*** **and** ***Betula pendula.*** Refer to **Fig.1** for detailed descriptions.

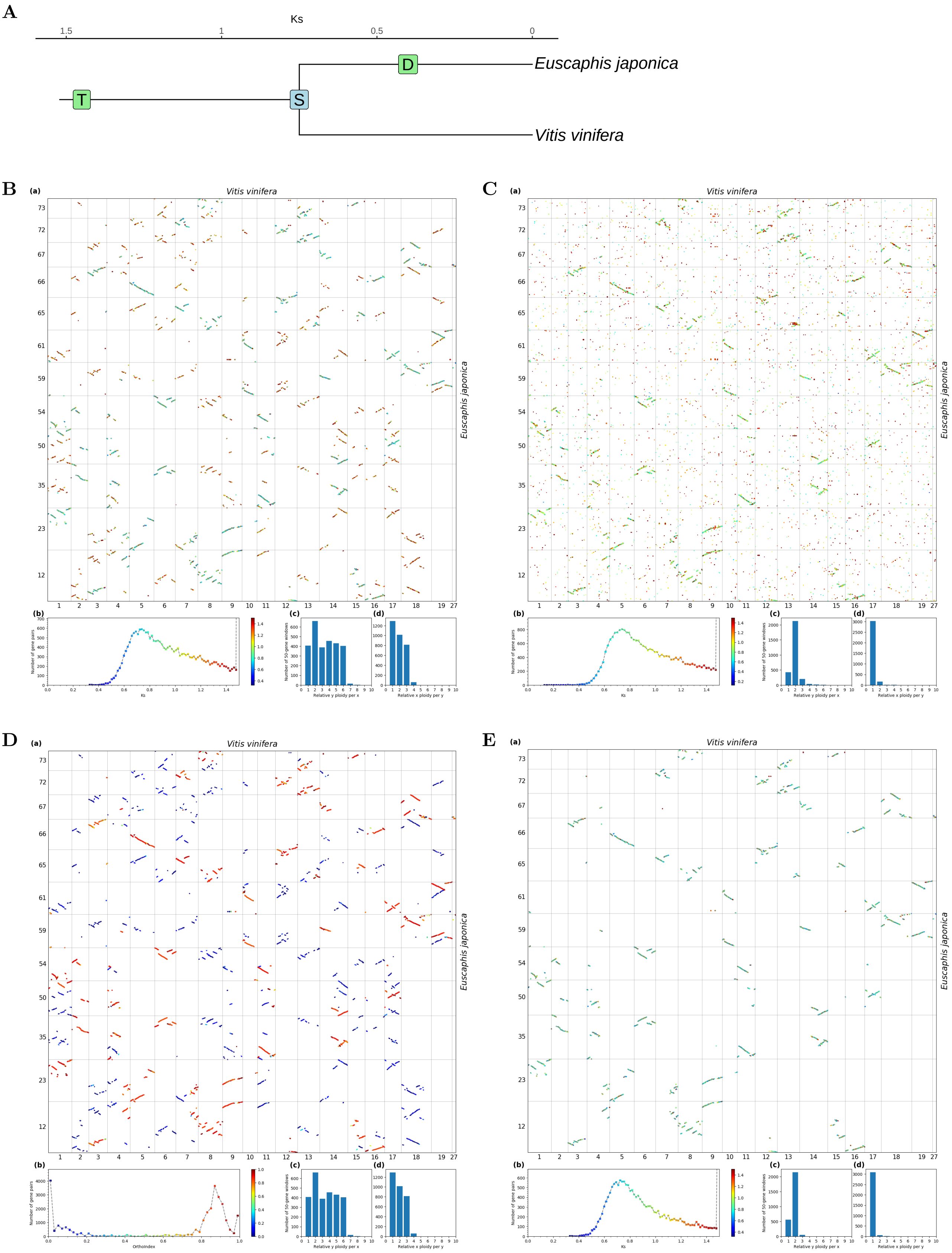


**Fig S26.** ***Orthology Index*** **in the identification of orthologous synteny in** ***Vitis vinifera*** **and** ***Euscaphis japonica*****.** Refer to **Fig.1** for detailed descriptions.

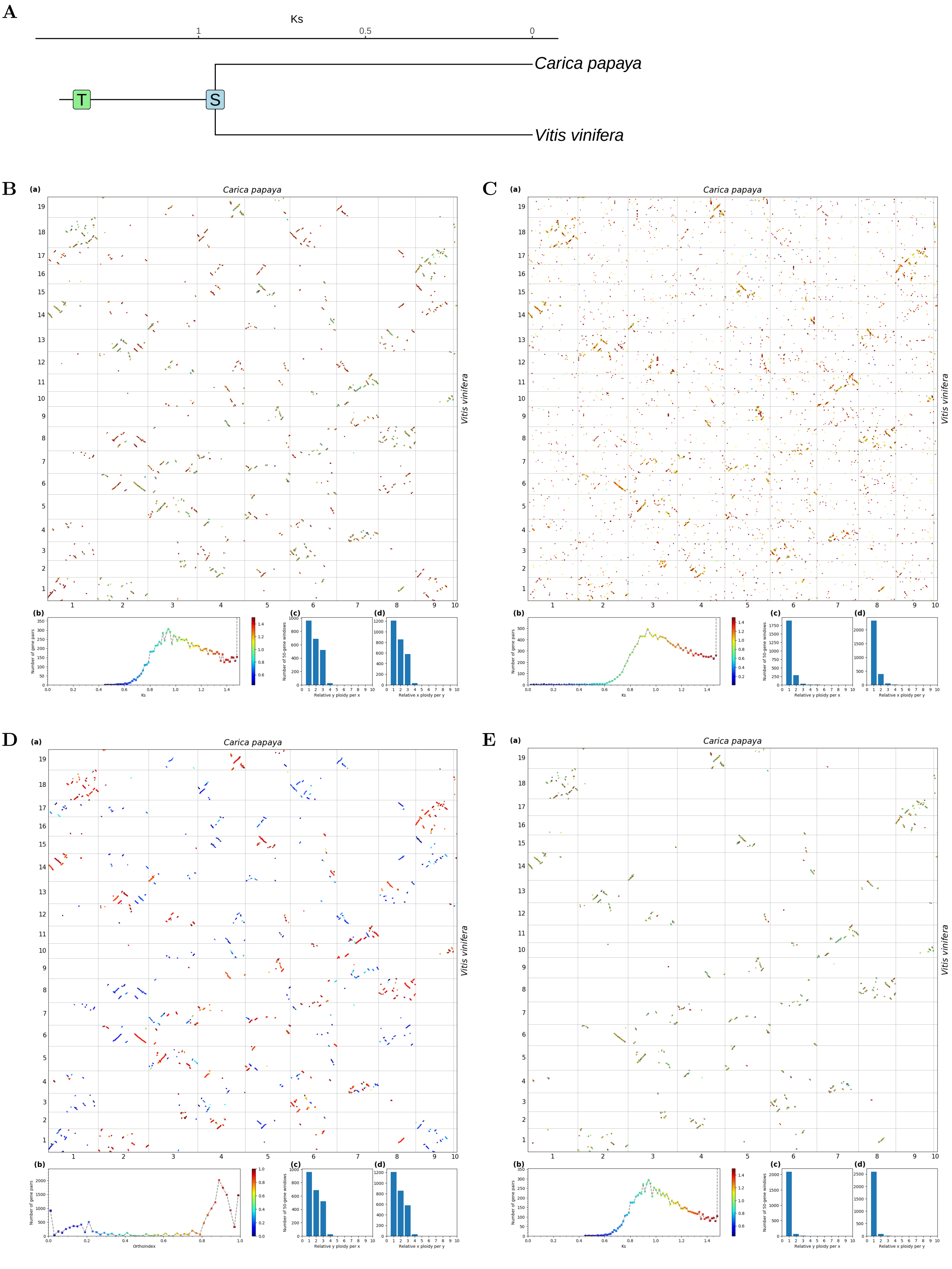


**Fig S27.** ***Orthology Index* in the identification of orthologous synteny in *Vitis vinifera* and *Carica papaya*.** Refer to **Fig.1** for detailed descriptions.

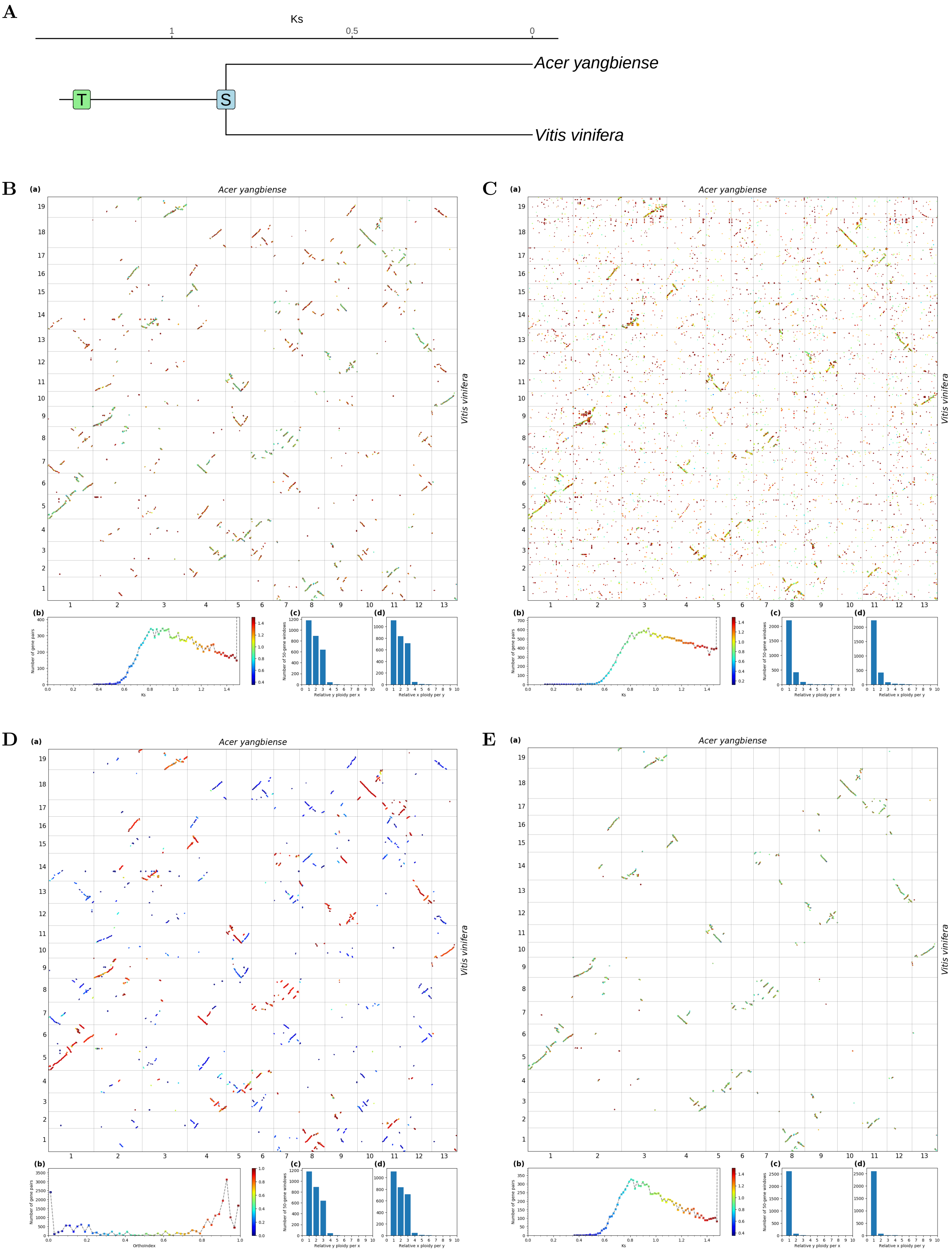


**Fig S28.** ***Orthology Index*** **in the identification of orthologous synteny in** ***Vitis vinifera*** **and** ***Acer yangbiense*****.** Refer to **Fig.1** for detailed descriptions.

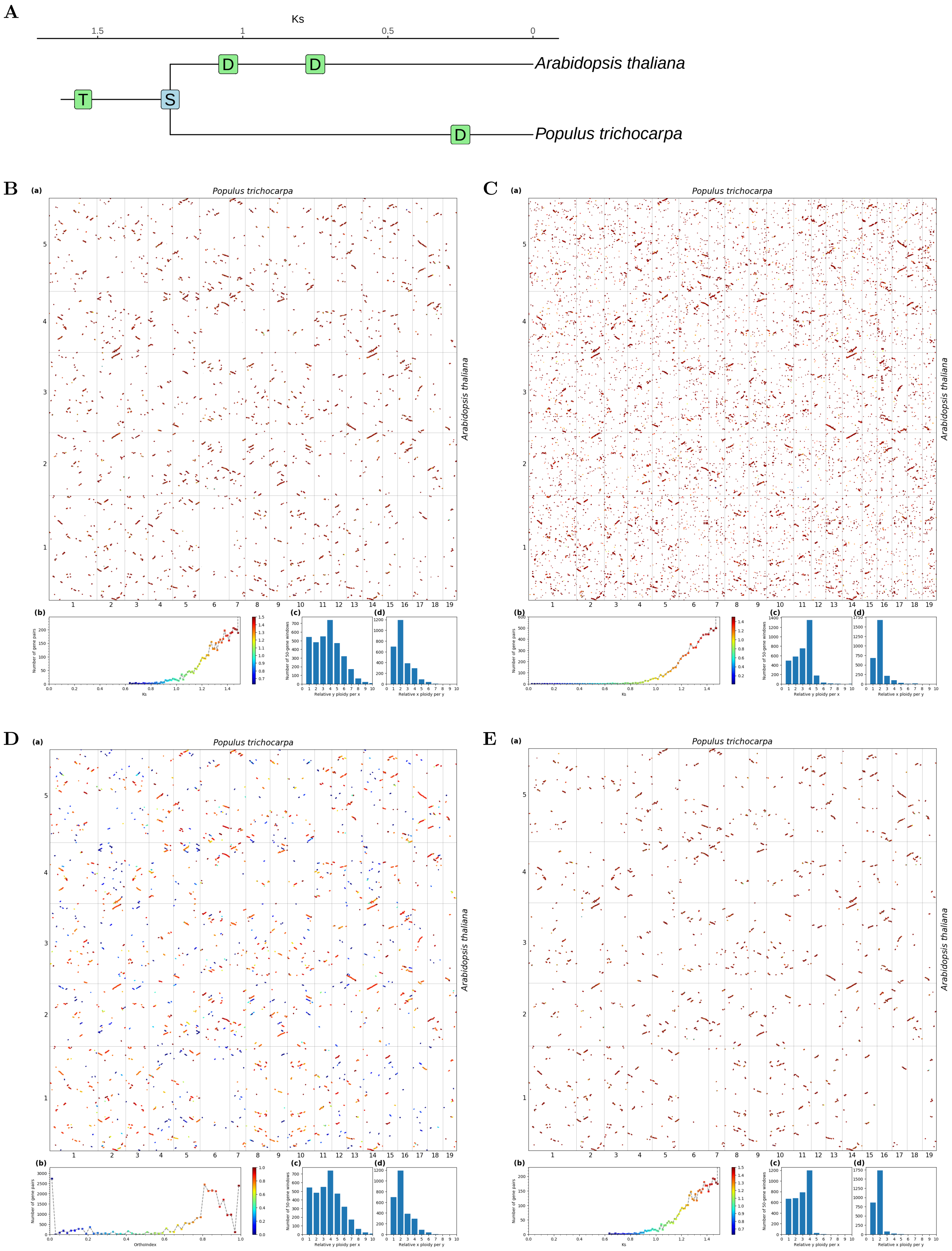


**Fig S29.** ***Orthology Index*** **in the identification of orthologous synteny in** ***Populus trichocarpa*** **and** ***Arabidopsis thaliana*****.** Refer to **Fig.1** for detailed descriptions.

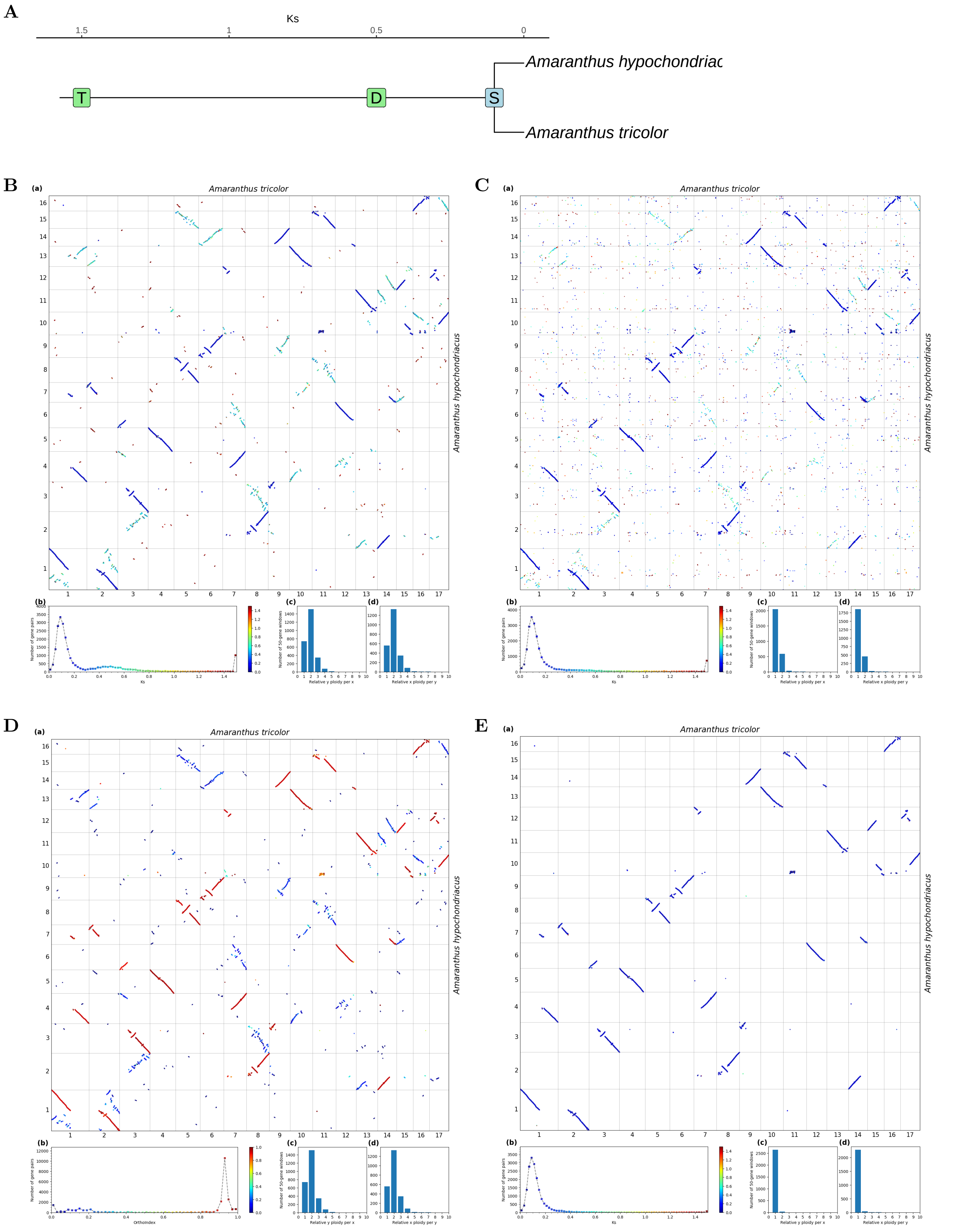


**Fig S30.** ***Orthology Index*** **in the identification of orthologous synteny in** ***Amaranthus tricolor*** **and** ***Amaranthus hypochondriacus*****.** Refer to **Fig.1** for detailed descriptions.

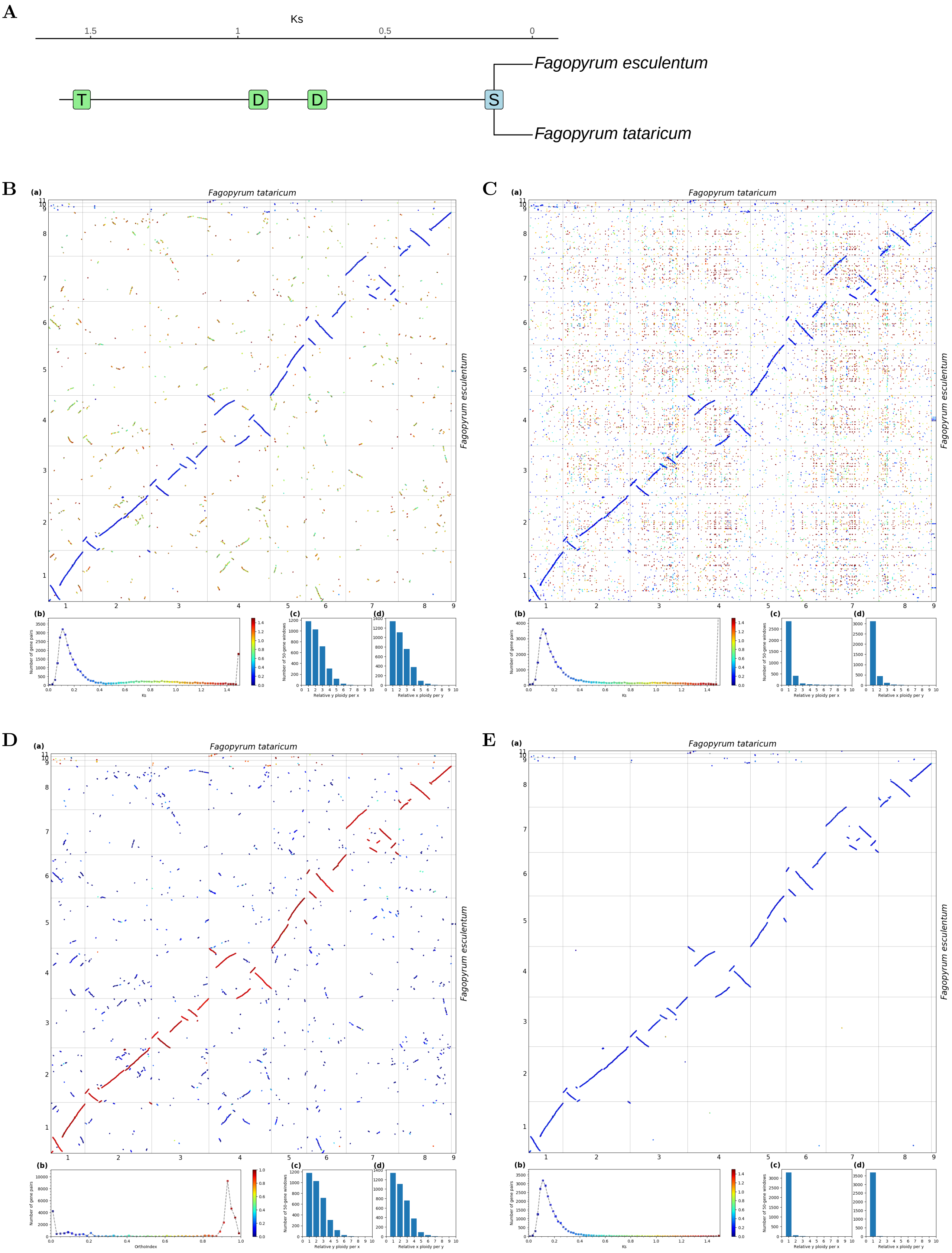


**Fig S31.** ***Orthology Index*** **in the identification of orthologous synteny in** ***Fagopyrum tataricum*** **and** ***Fagopyrum esculentum.*** Refer to **Fig.1** for detailed descriptions.

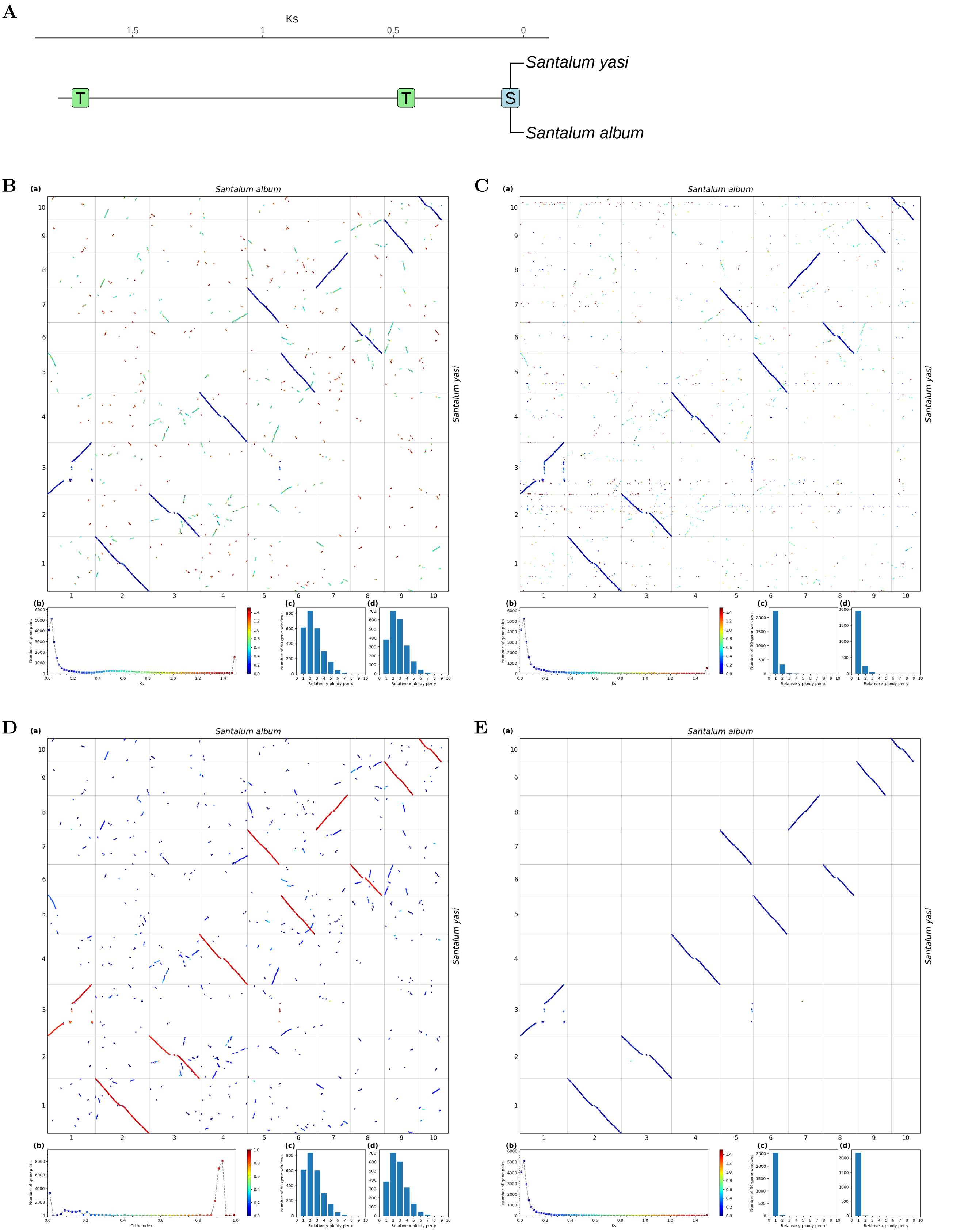


**Fig S32.** ***Orthology Index*** **in the identification of orthologous synteny in** ***Santalum album*** **and** ***Santalum yasi*****.** Refer to **Fig.1** for detailed descriptions.

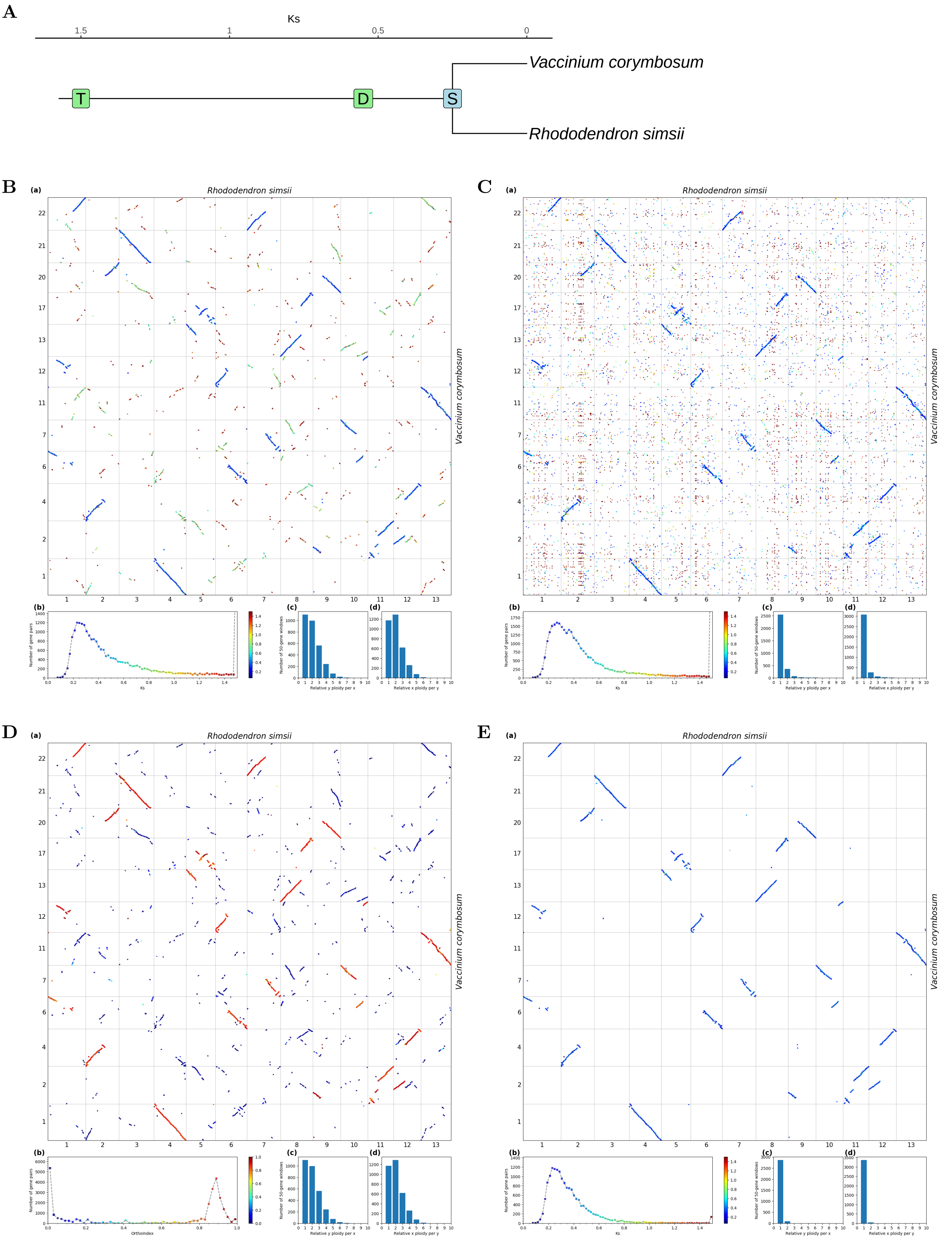


**Fig S33.** ***Orthology Index*** **in the identification of orthologous synteny in** ***Rhododendron simsii*** **and** ***Vaccinium corymbosum.*** Refer to **Fig.1** for detailed descriptions.

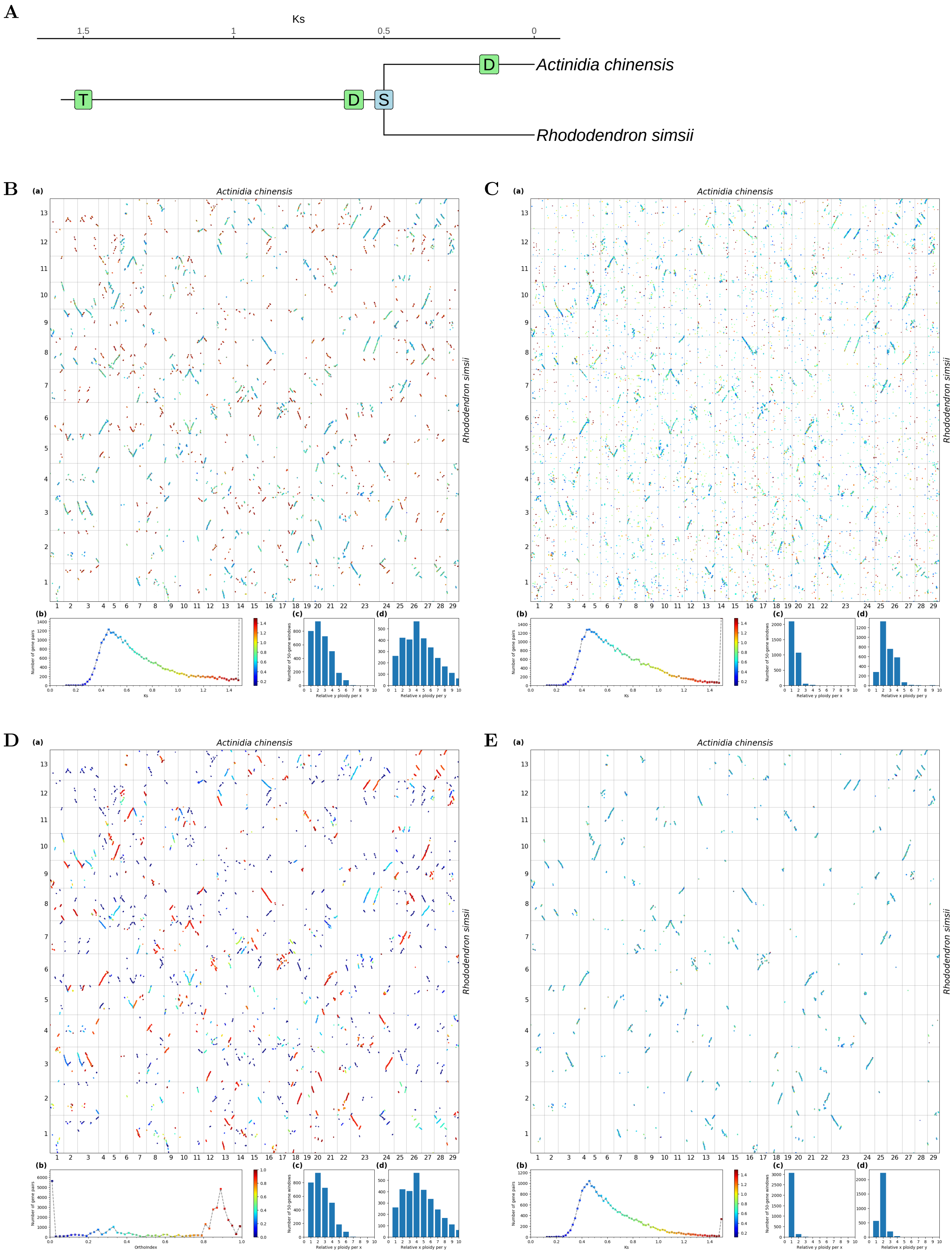


**Fig S34.** ***Orthology Index* in the identification of orthologous synteny in *Rhododendron simsii* and *Actinidia chinensis*.** Refer to **Fig.1** for detailed descriptions.

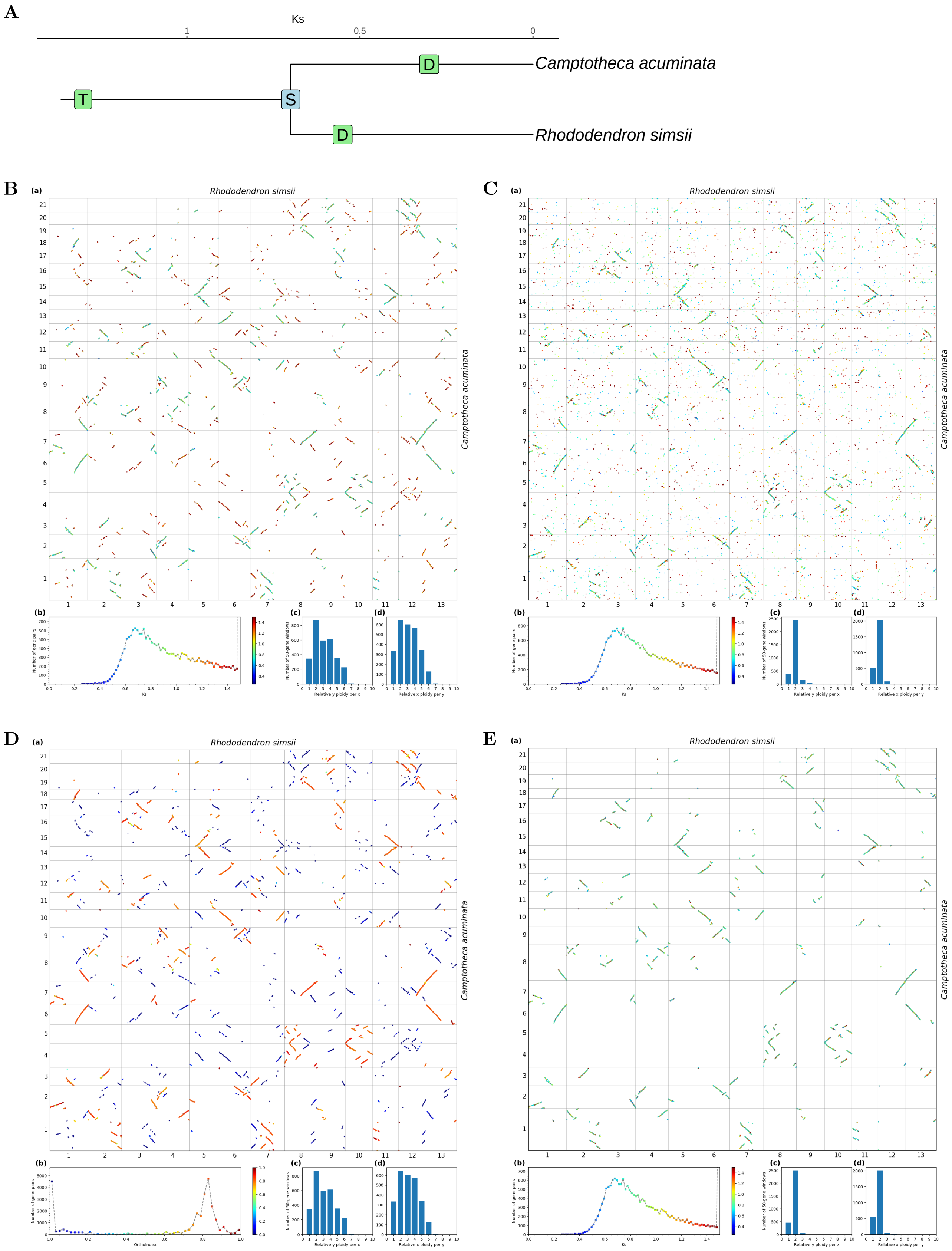


**Fig S35.** ***Orthology Index* in the identification of orthologous synteny in *Rhododendron simsii* and *Camptotheca acuminata*.** Refer to **Fig.1** for detailed descriptions.

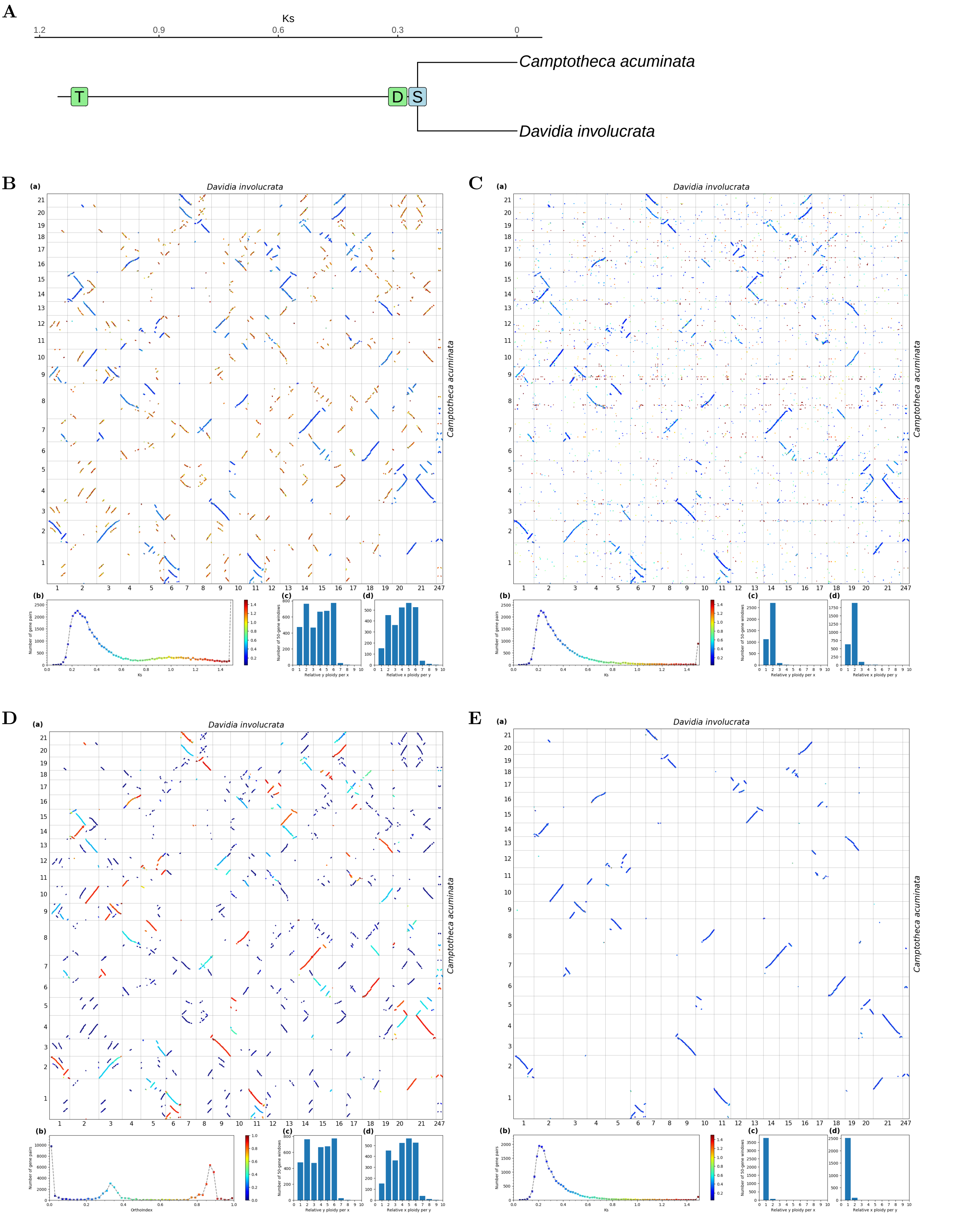


**Fig S36.** ***Orthology Index*** **in the identification of orthologous synteny in** ***Davidia involucrata*** **and** ***Camptotheca acuminata*****.** Refer to **Fig.1** for detailed descriptions.

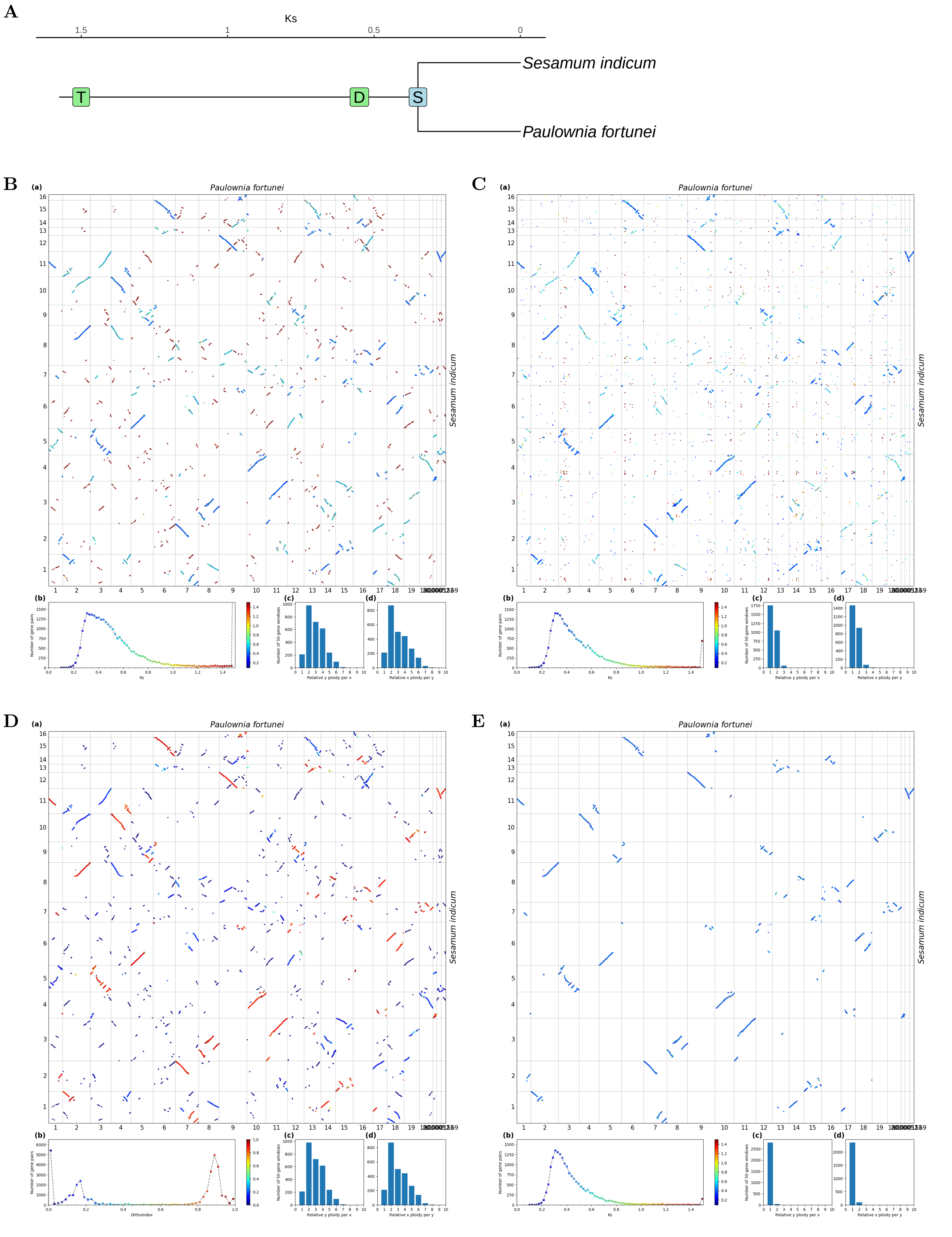


**Fig S37.** ***Orthology Index* in the identification of orthologous synteny in *Paulownia fortunei* and *Sesamum indicum*.** Refer to **Fig.1** for detailed descriptions.

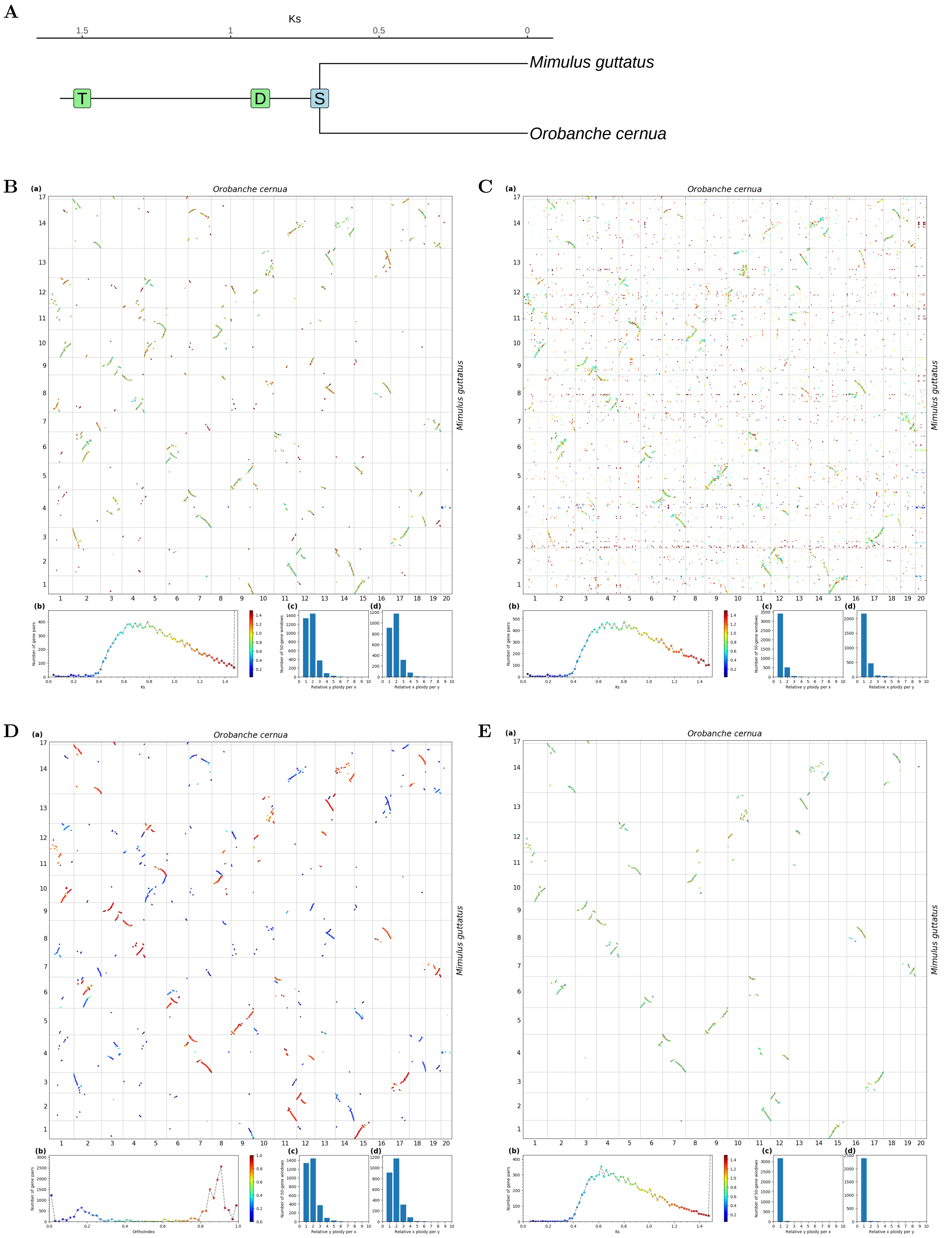


**Fig S38.** ***Orthology Index* in the identification of orthologous synteny in *Orobanche cernua* and *Mimulus guttatus*.** Refer to **Fig.1** for detailed descriptions.

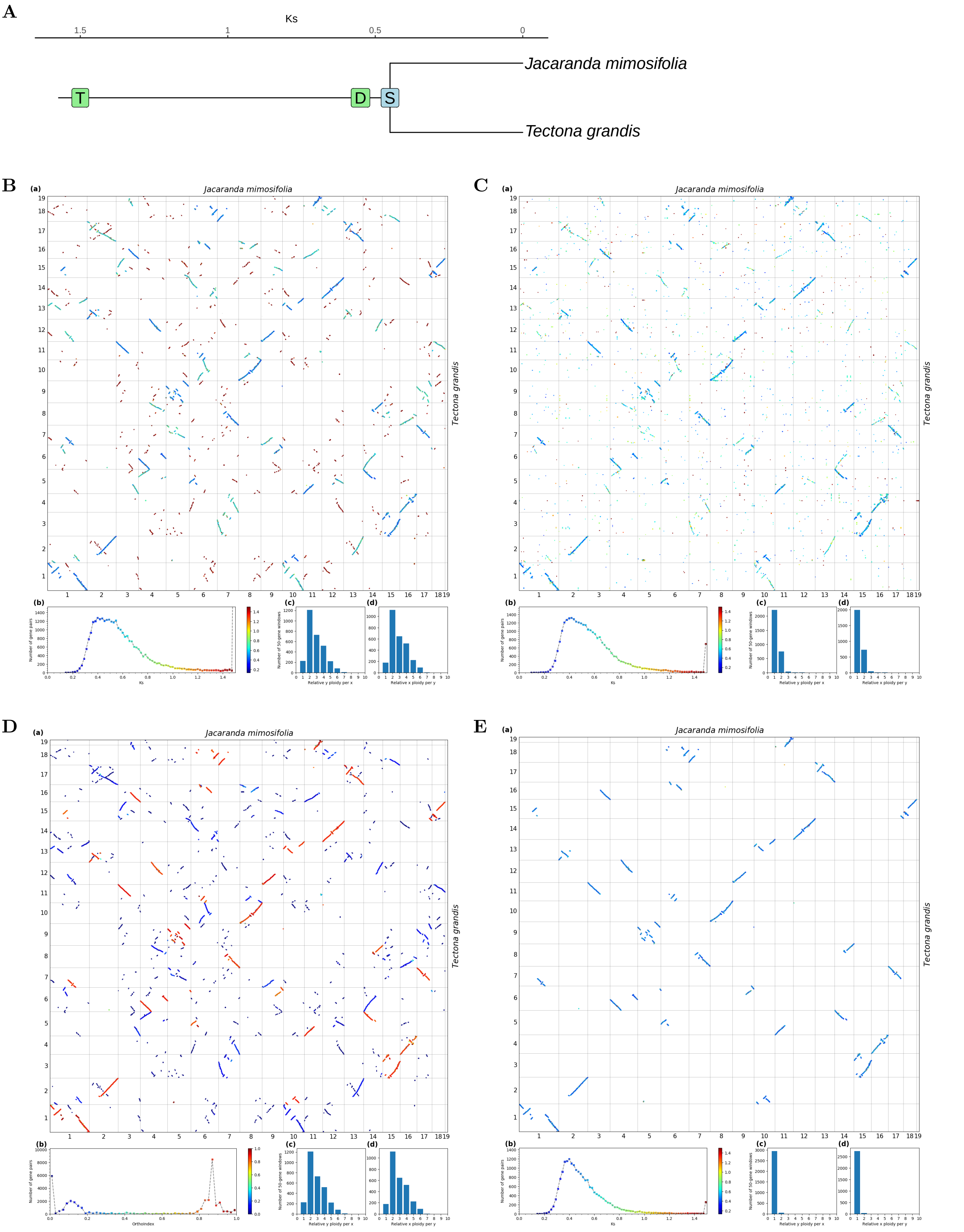


**Fig S39.** ***Orthology Index*** **in the identification of orthologous synteny in** ***Tectona grandis*** **and** ***Jacaranda mimosifolia.*** Refer to **Fig.1** for detailed descriptions.

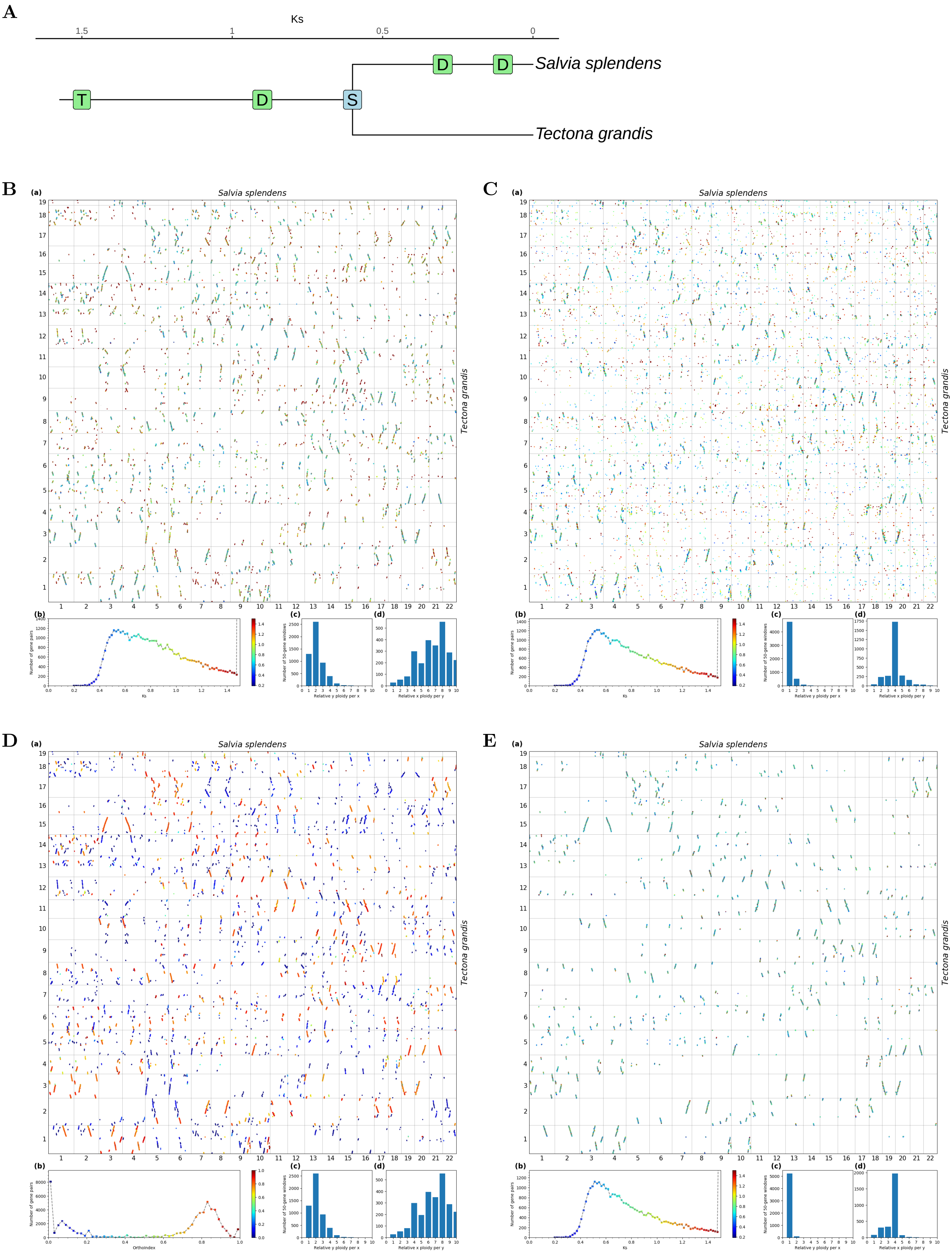


**Fig S40.** ***Orthology Index*** **in the identification of orthologous synteny in** ***Tectona grandis*** **and** ***Salvia splendens*****.** Refer to **Fig.1** for detailed descriptions.

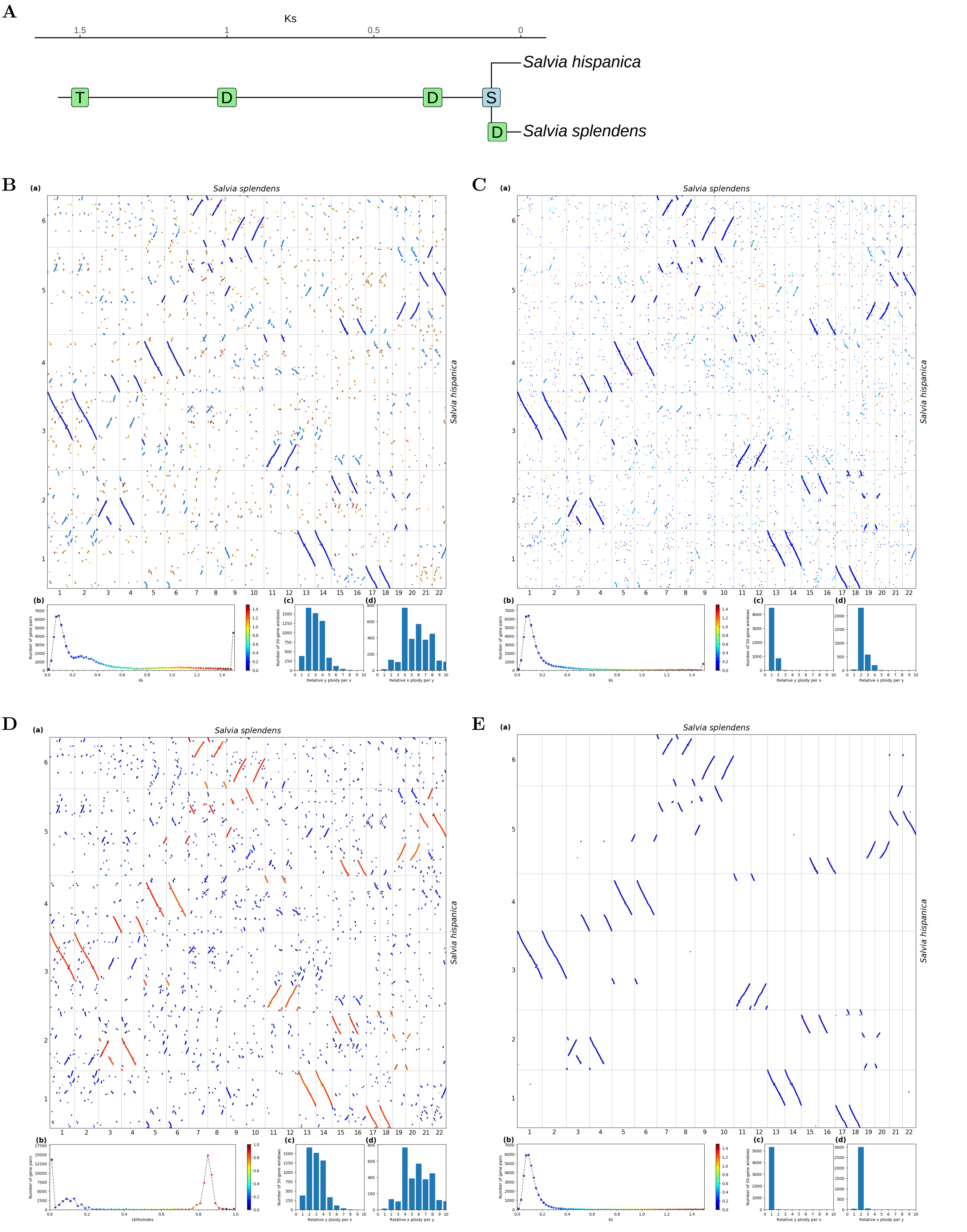


**Fig S41.** ***Orthology Index*** **in the identification of orthologous synteny in** ***Salvia splendens*** **and** ***Salvia hispanica.*** Refer to **Fig.1** for detailed descriptions.

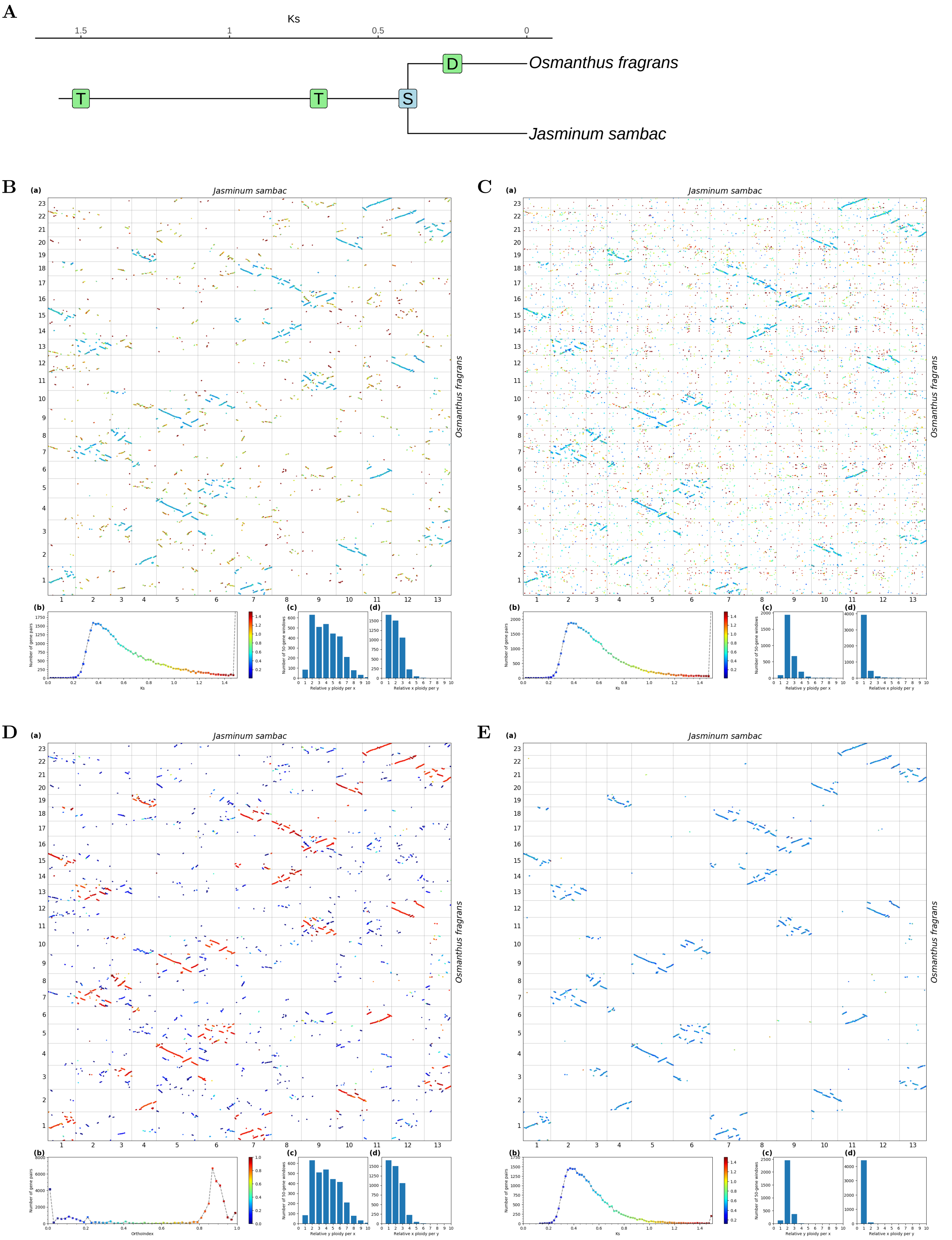


**Fig S42.** ***Orthology Index*** **in the identification of orthologous synteny in** ***Jasminum sambac*** **and** ***Osmanthus fragrans.*** Refer to **Fig.1** for detailed descriptions.

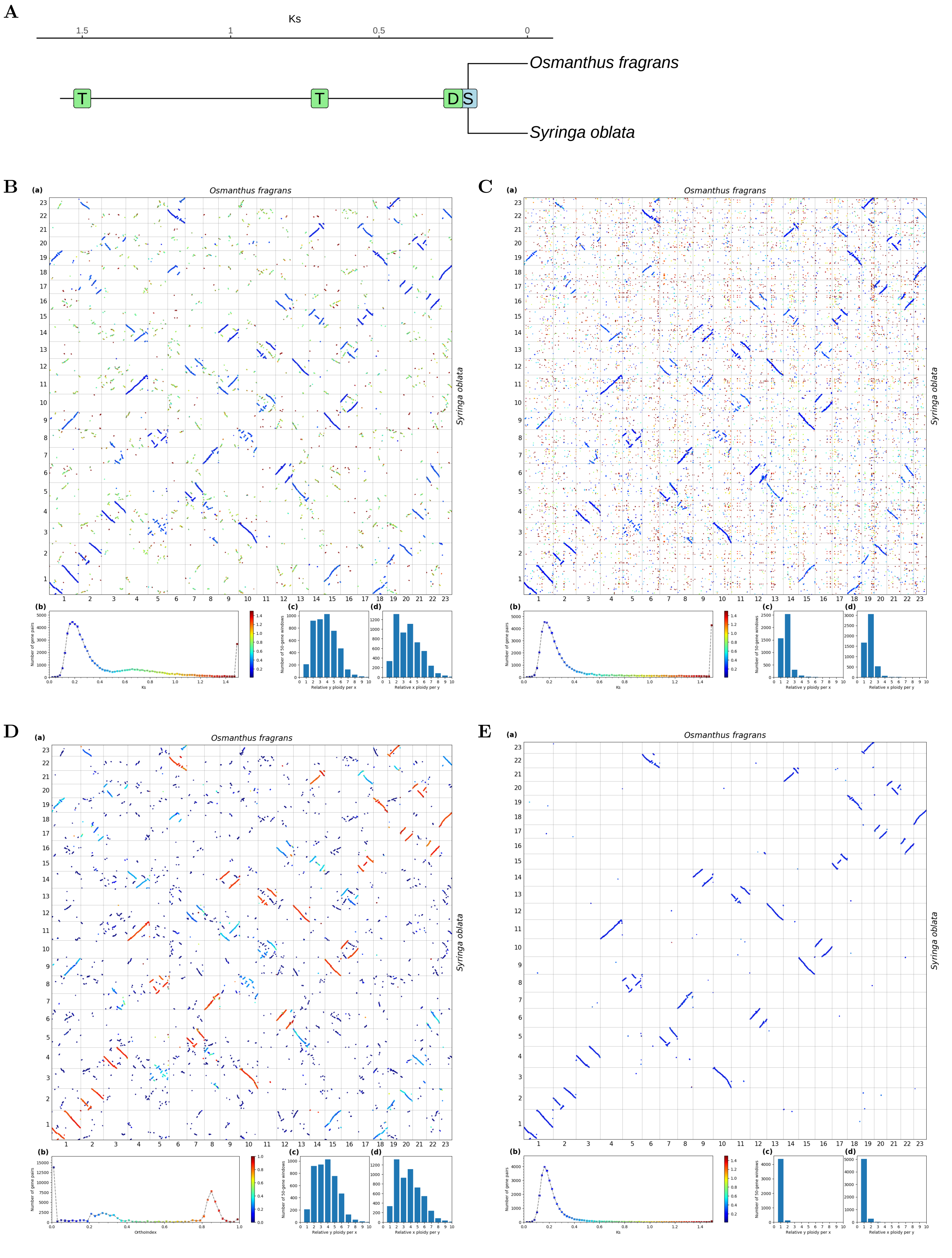


**Fig S43.** ***Orthology Index*** **in the identification of orthologous synteny in** ***Syringa oblata*** **and** ***Osmanthus fragrans.*** Refer to **Fig.1** for detailed descriptions.

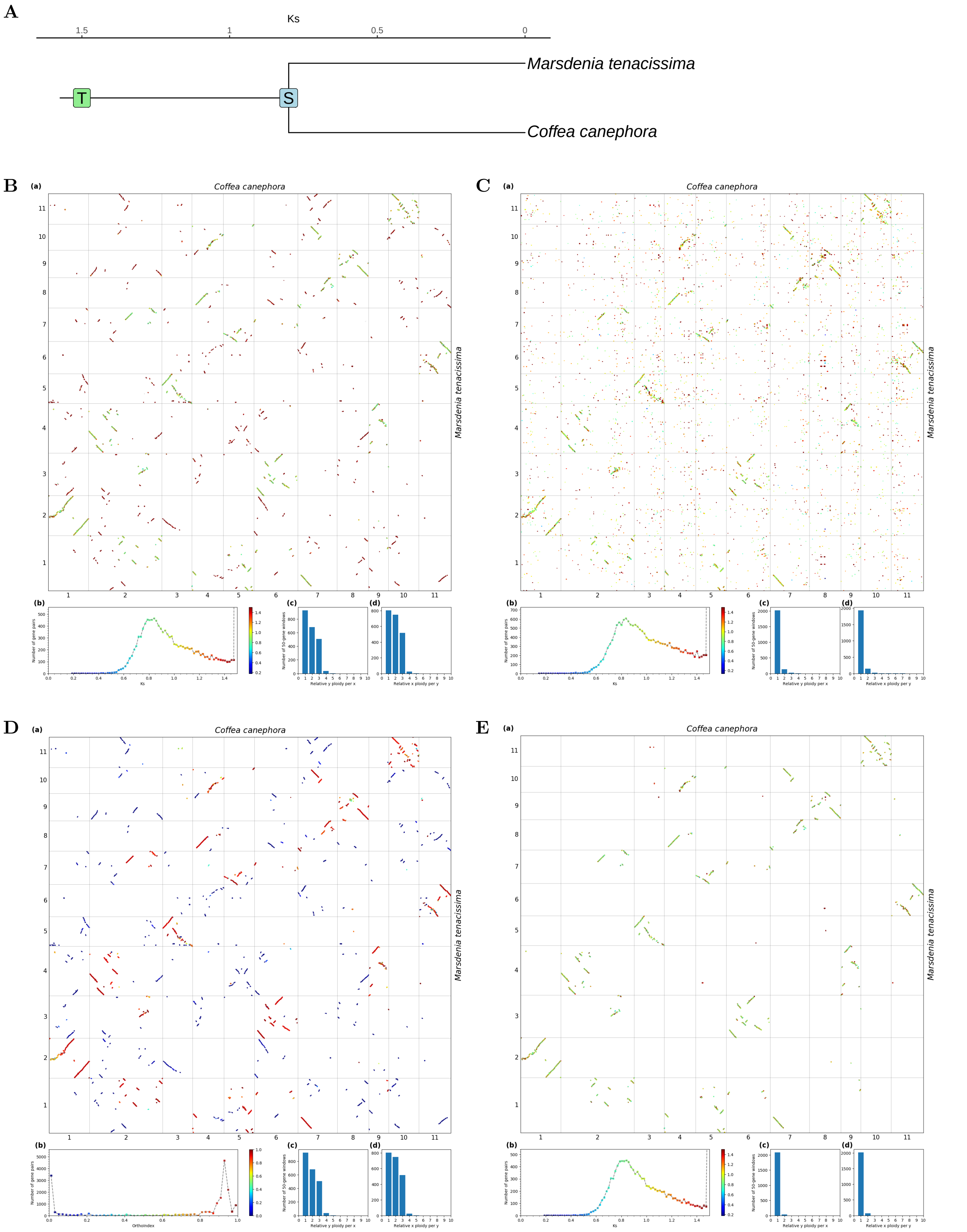


**Fig S44.** ***Orthology Index* in the identification of orthologous synteny in *Coffea canephora* and *Marsdenia tenacissima*.** Refer to **Fig.1** for detailed descriptions.

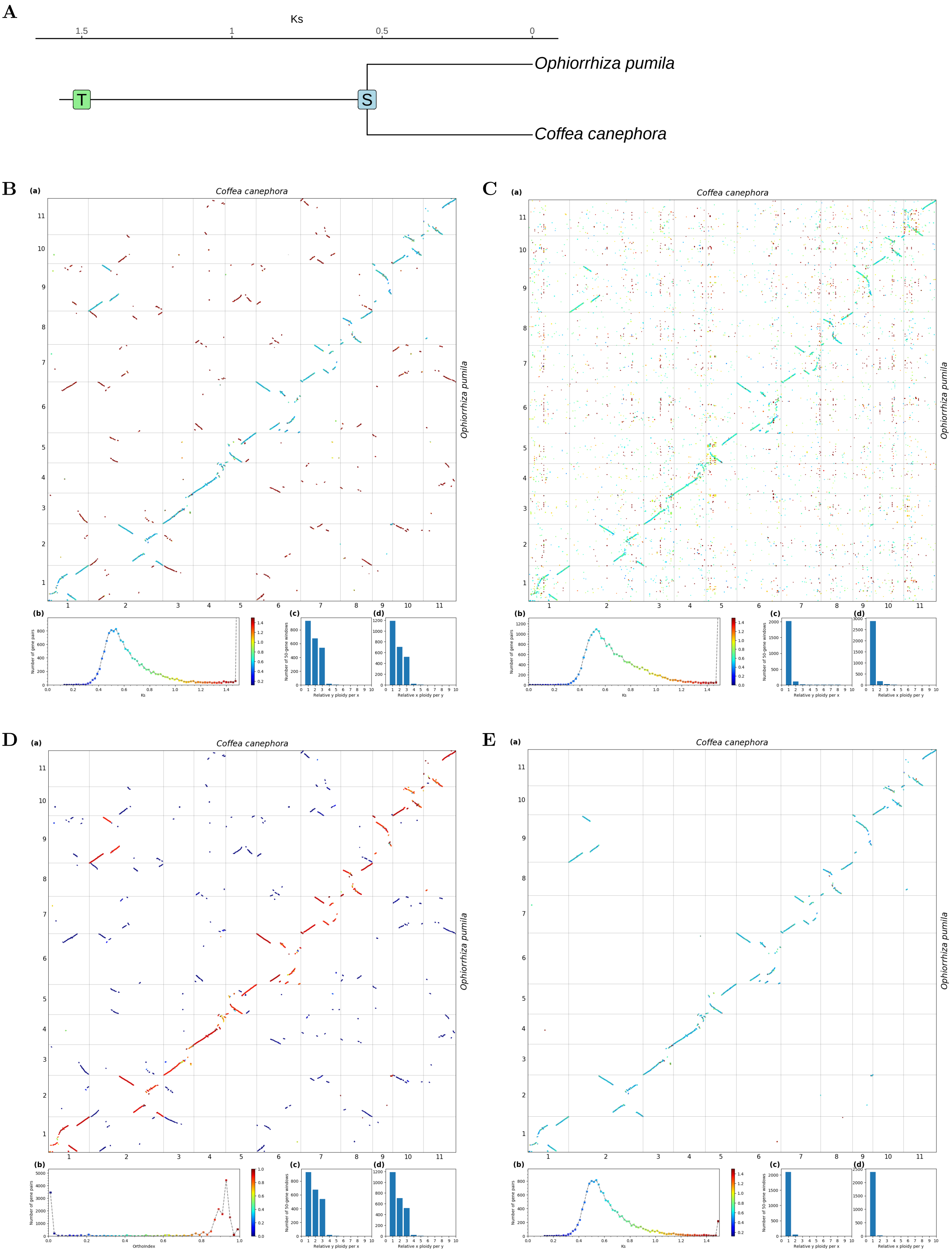


**Fig S45.** ***Orthology Index*** **in the identification of orthologous synteny in** ***Coffea canephora*** **and** ***Ophiorrhiza pumila.*** Refer to **Fig.1** for detailed descriptions.

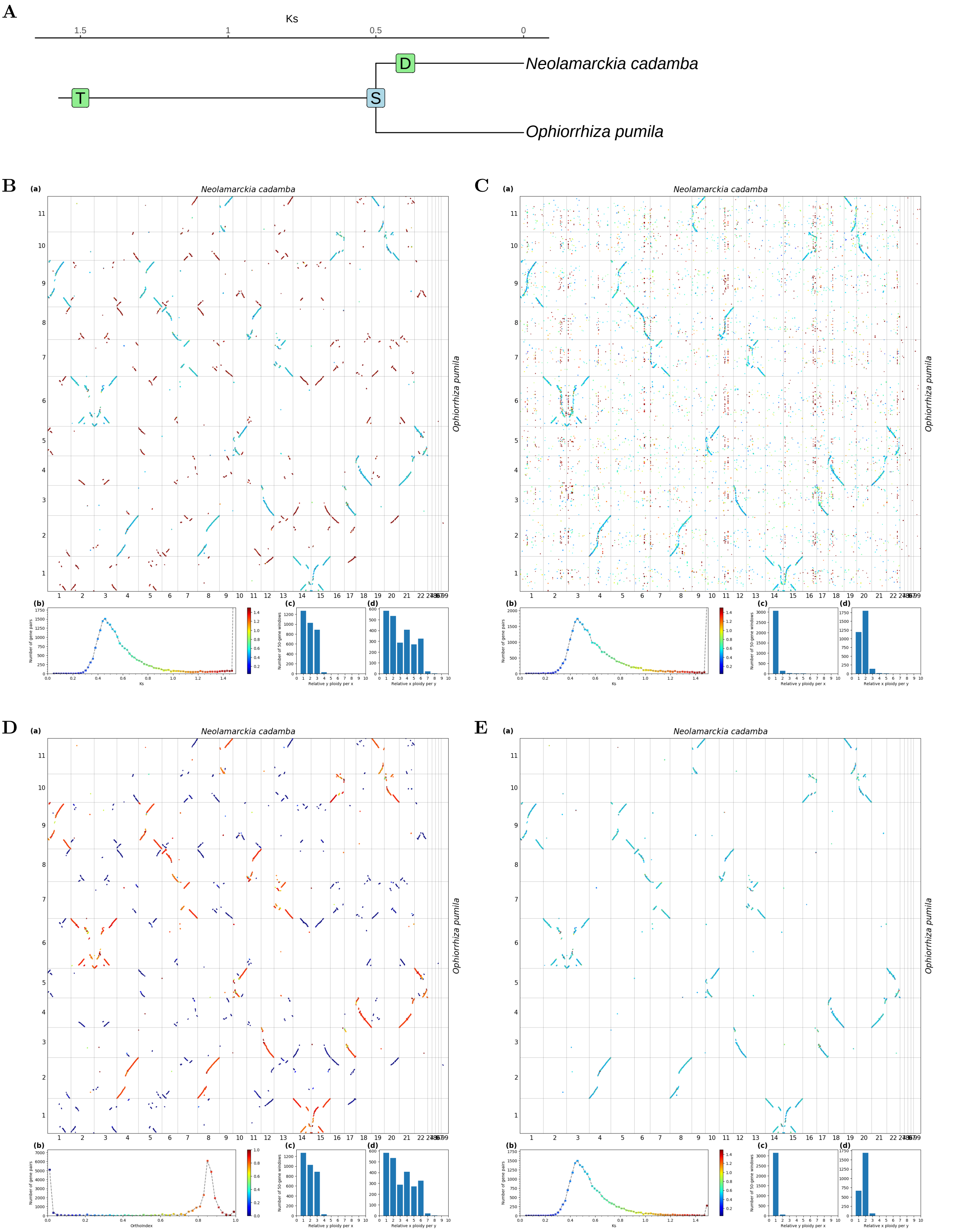


**Fig S46.** ***Orthology Index*** **in the identification of orthologous synteny in** ***Ophiorrhiza pumila*** **and** ***Neolamarckia cadamba.*** Refer to **Fig.1** for detailed descriptions.

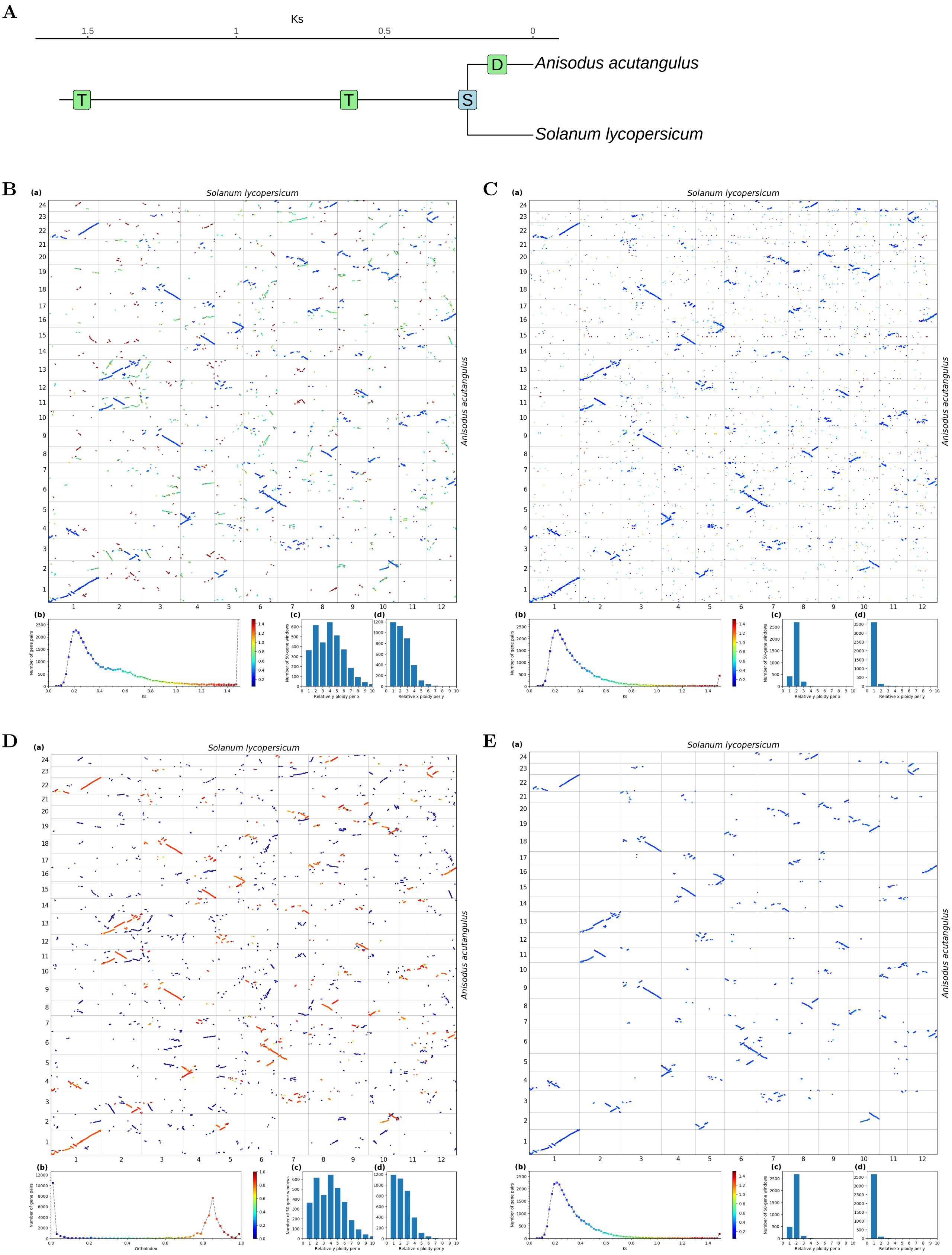


**Fig S47.** ***Orthology Index*** **in the identification of orthologous synteny in** ***Solanum lycopersicum*** **and** ***Anisodus acutangulus.*** Refer to **Fig.1** for detailed descriptions.

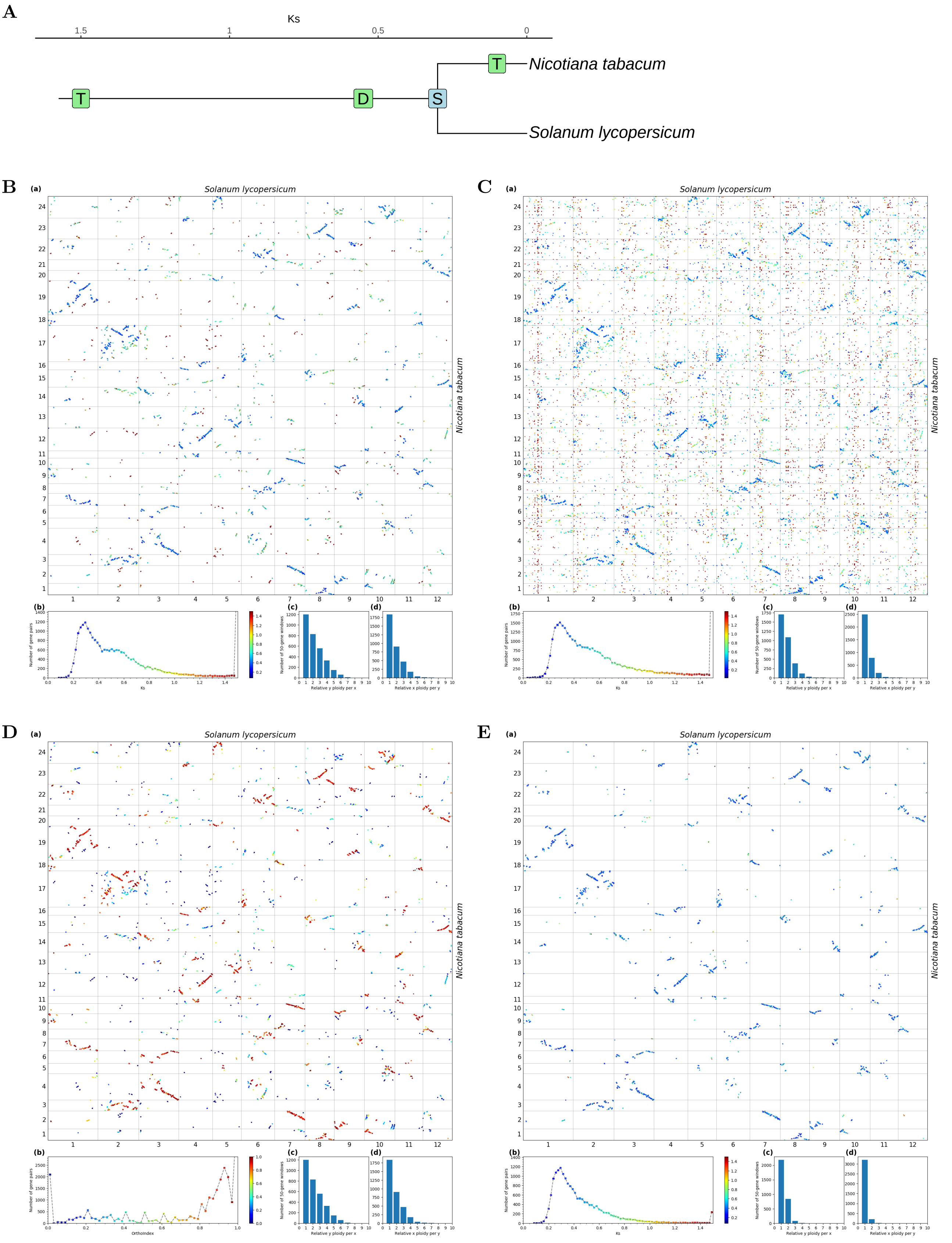


**Fig S48.** ***Orthology Index*** **in the identification of orthologous synteny in** ***Solanum lycopersicum*** **and** ***Nicotiana tabacum.*** Refer to **Fig.1** for detailed descriptions.

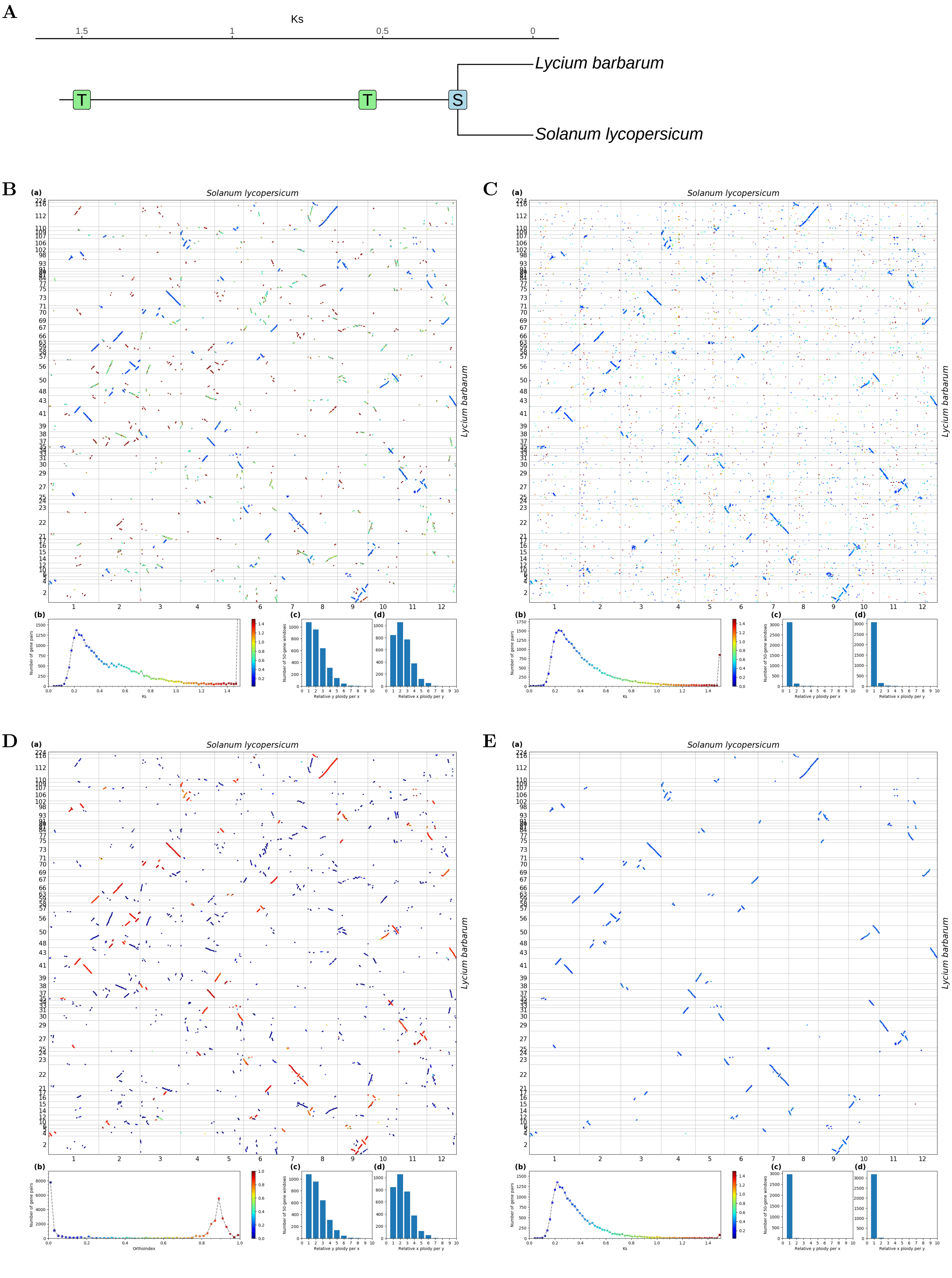


**Fig S49.** ***Orthology Index*** **in the identification of orthologous synteny in** ***Solanum lycopersicum*** **and** ***Lycium barbarum*****.** Refer to **Fig.1** for detailed descriptions.

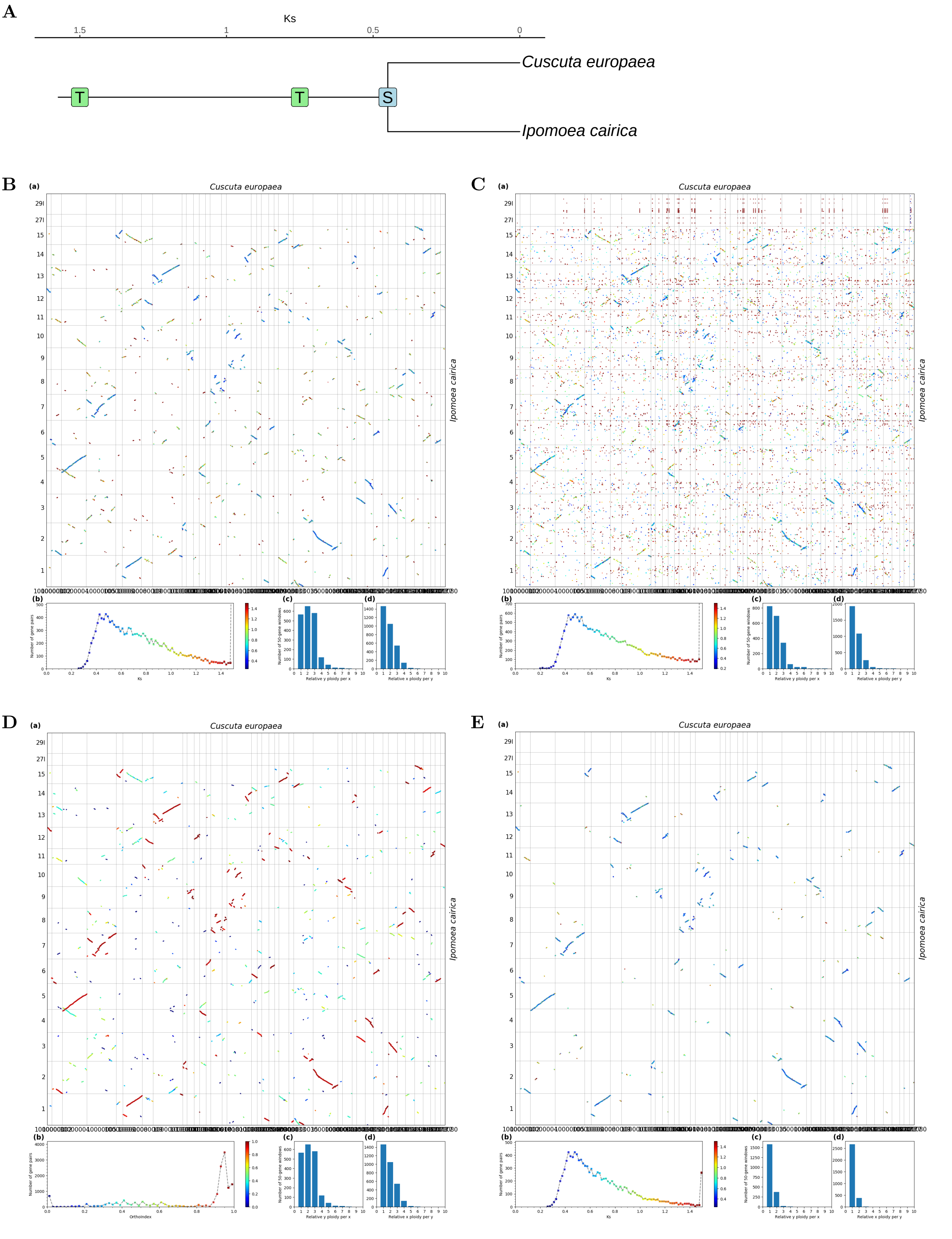


**Fig S50.** ***Orthology Index* in the identification of orthologous synteny in *Ipomoea cairica* and *Cuscuta europaea*.** Refer to **Fig.1** for detailed descriptions.

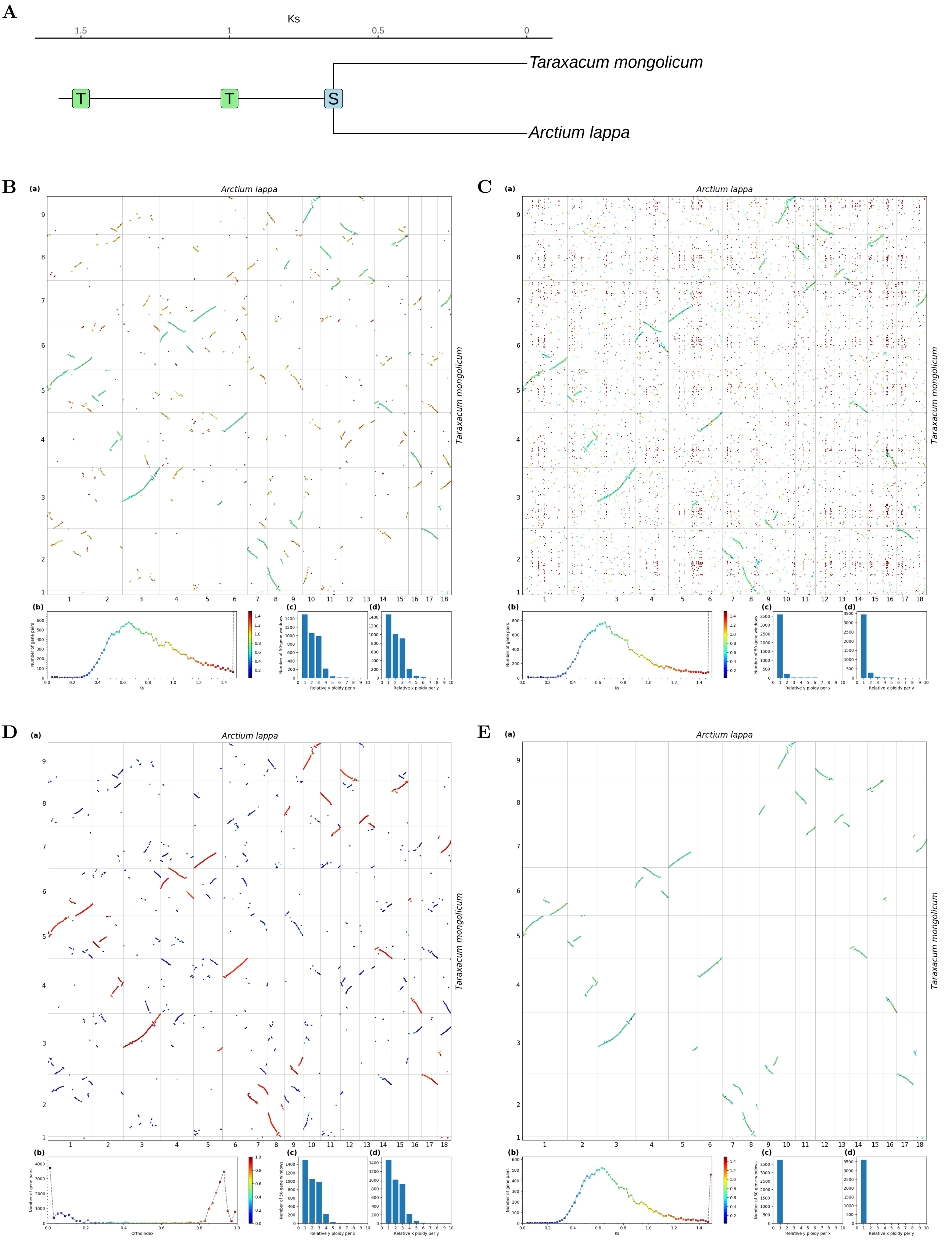


**Fig S51.** ***Orthology Index*** **in the identification of orthologous synteny in** ***Arctium lappa*** **and** ***Taraxacum mongolicum*.** Refer to **Fig.1** for detailed descriptions.

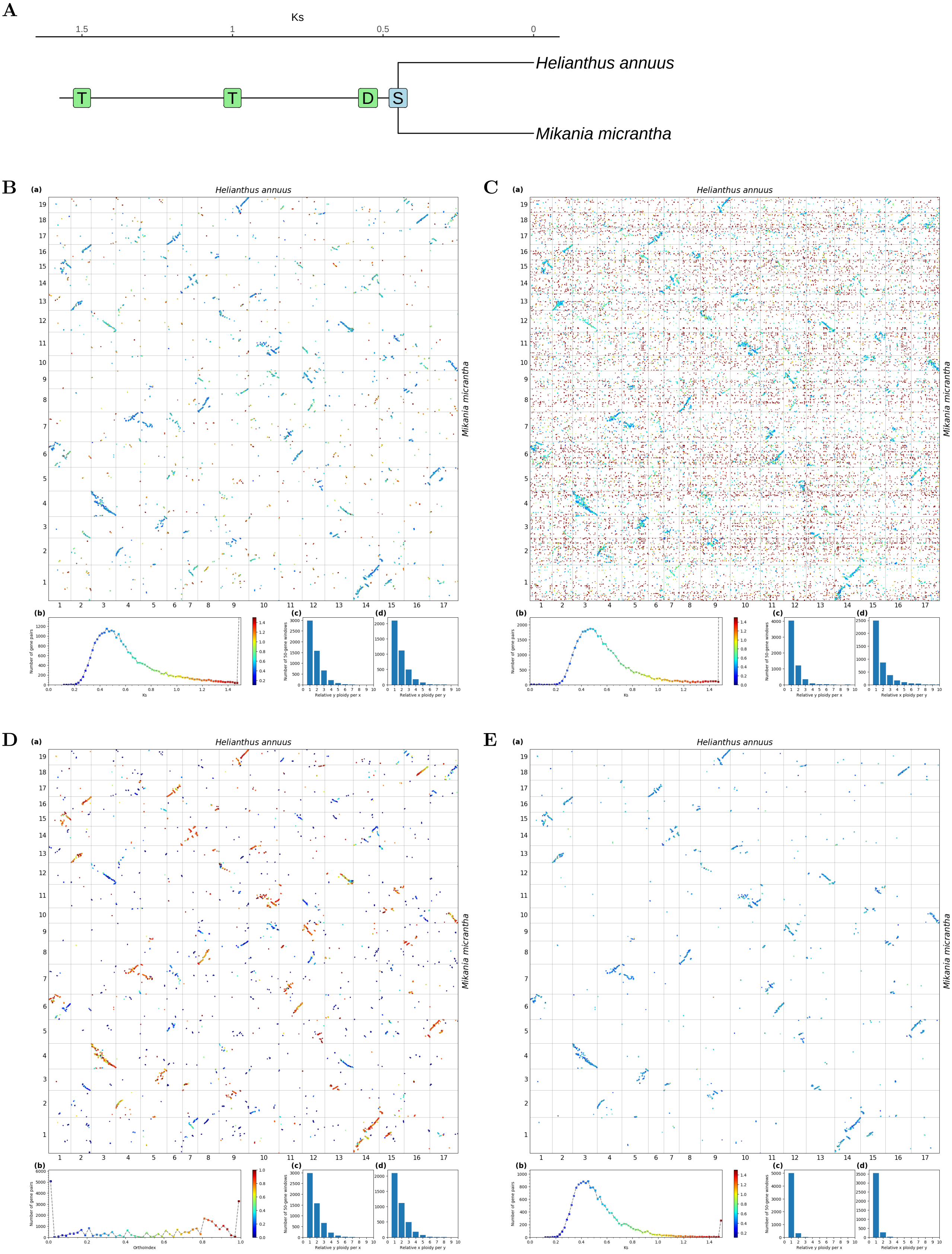


**Fig S52.** ***Orthology Index*** **in the identification of orthologous synteny in** ***Mikania micrantha*** **and** ***Helianthus annuus*****.** Refer to **Fig.1** for detailed descriptions.

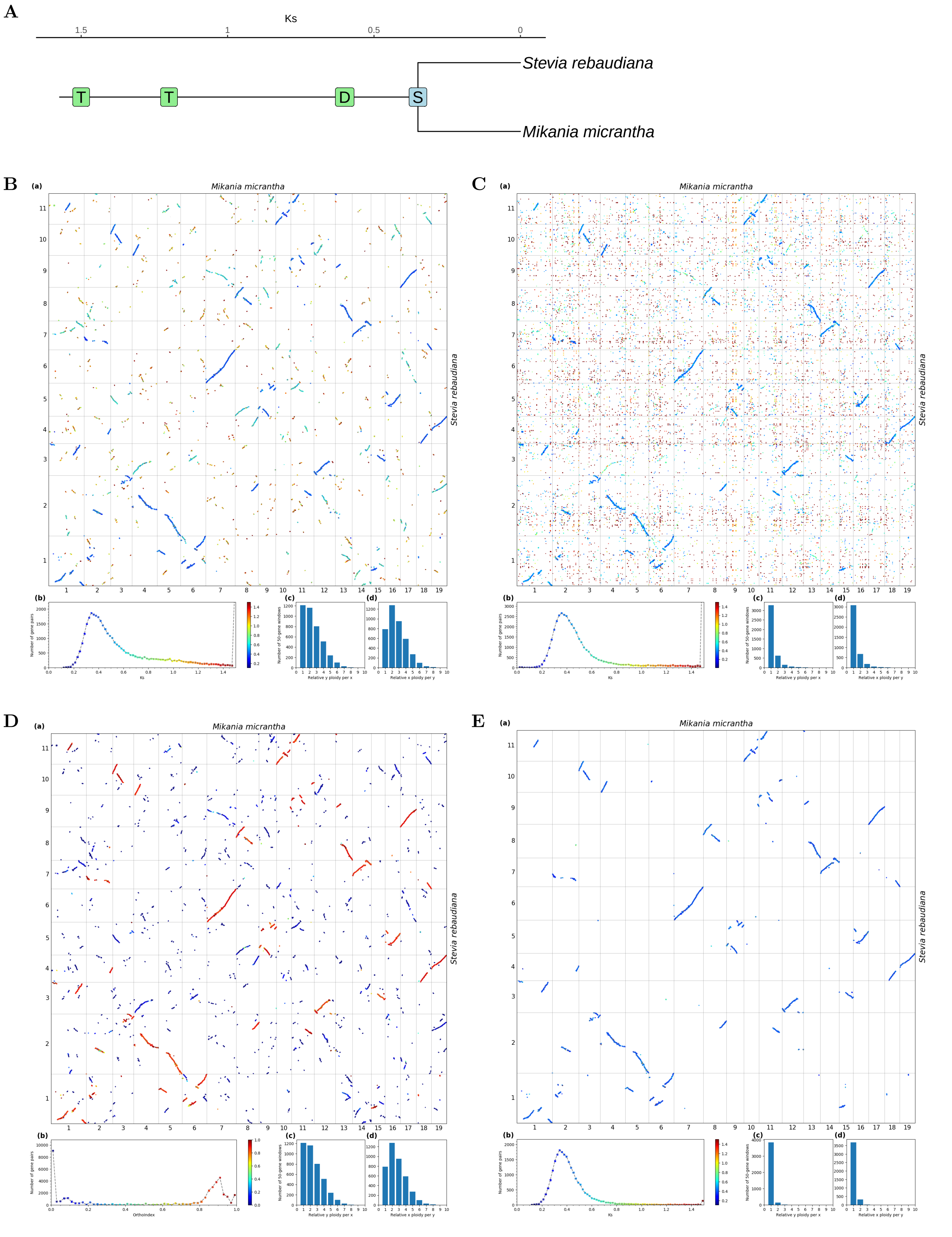


**Fig S53.** ***Orthology Index*** **in the identification of orthologous synteny in** ***Mikania micrantha*** **and** ***Stevia rebaudiana.*** Refer to **Fig.1** for detailed descriptions.

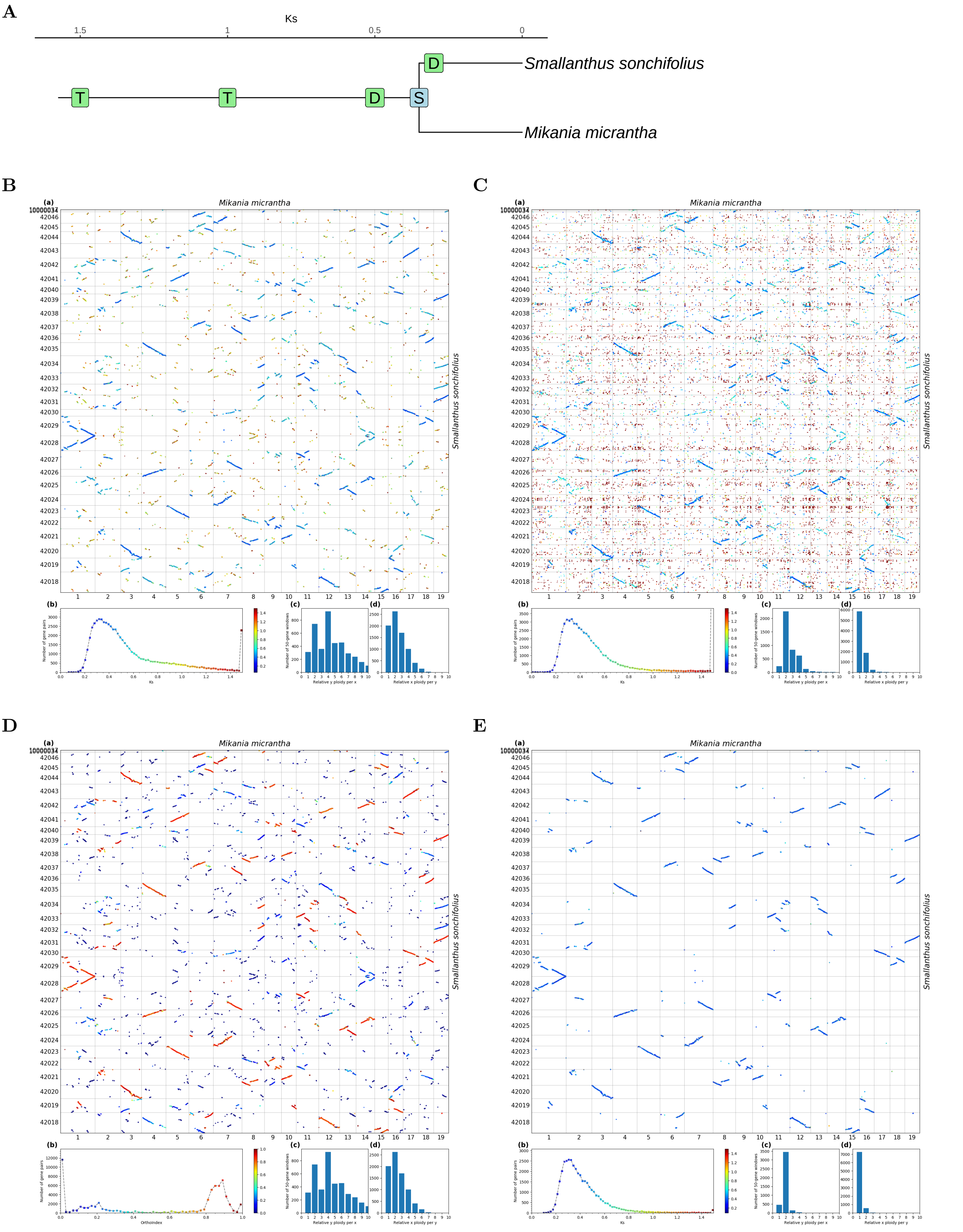


**Fig S54.** ***Orthology Index* in the identification of orthologous synteny in *Mikania micrantha* and *Smallanthus sonchifolius*.** Refer to **Fig.1** for detailed descriptions.

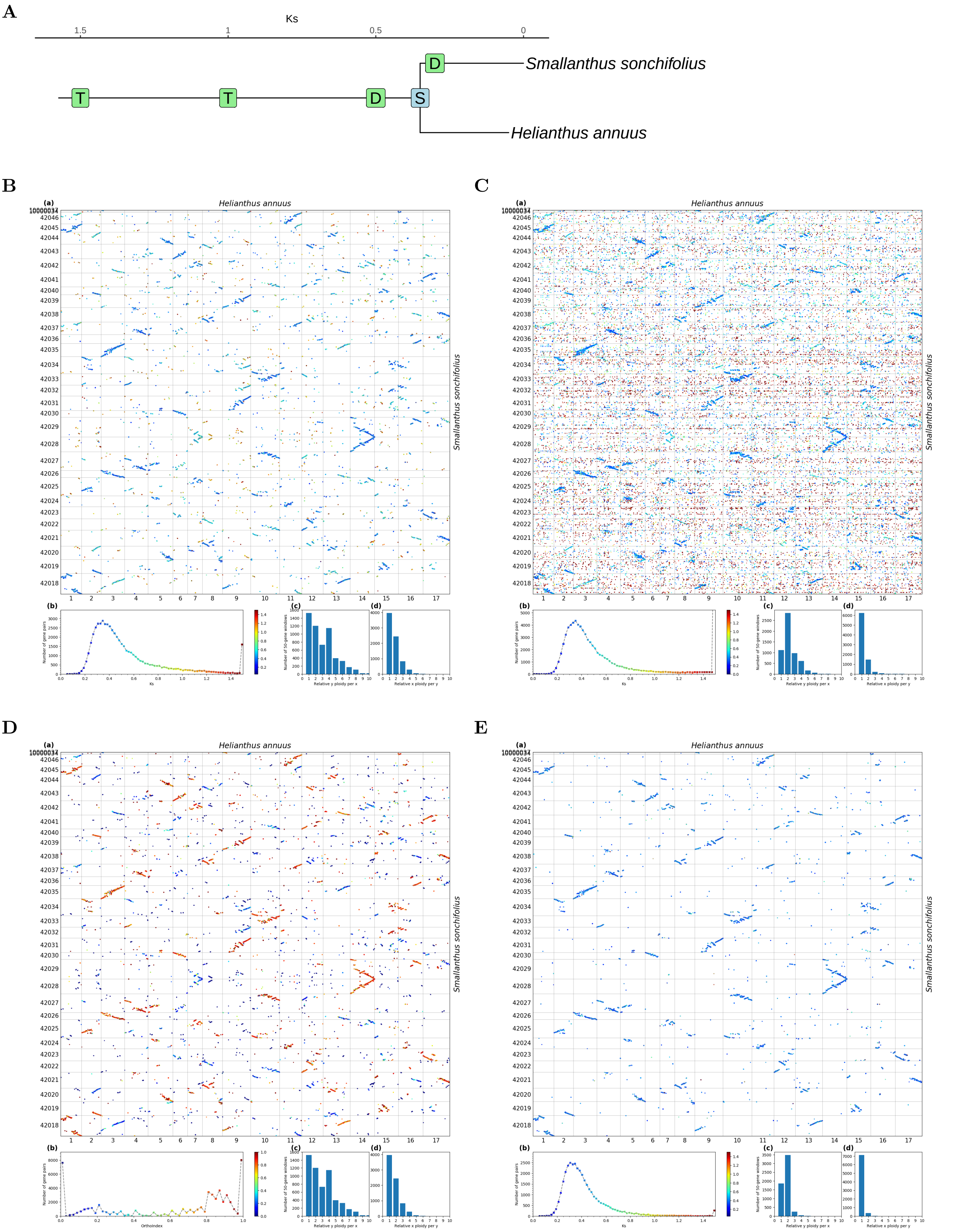


**Fig S55.** ***Orthology Index*** **in the identification of orthologous synteny in** ***Helianthus annuus*** **and** ***Smallanthus sonchifolius.*** Refer to **Fig.1** for detailed descriptions.

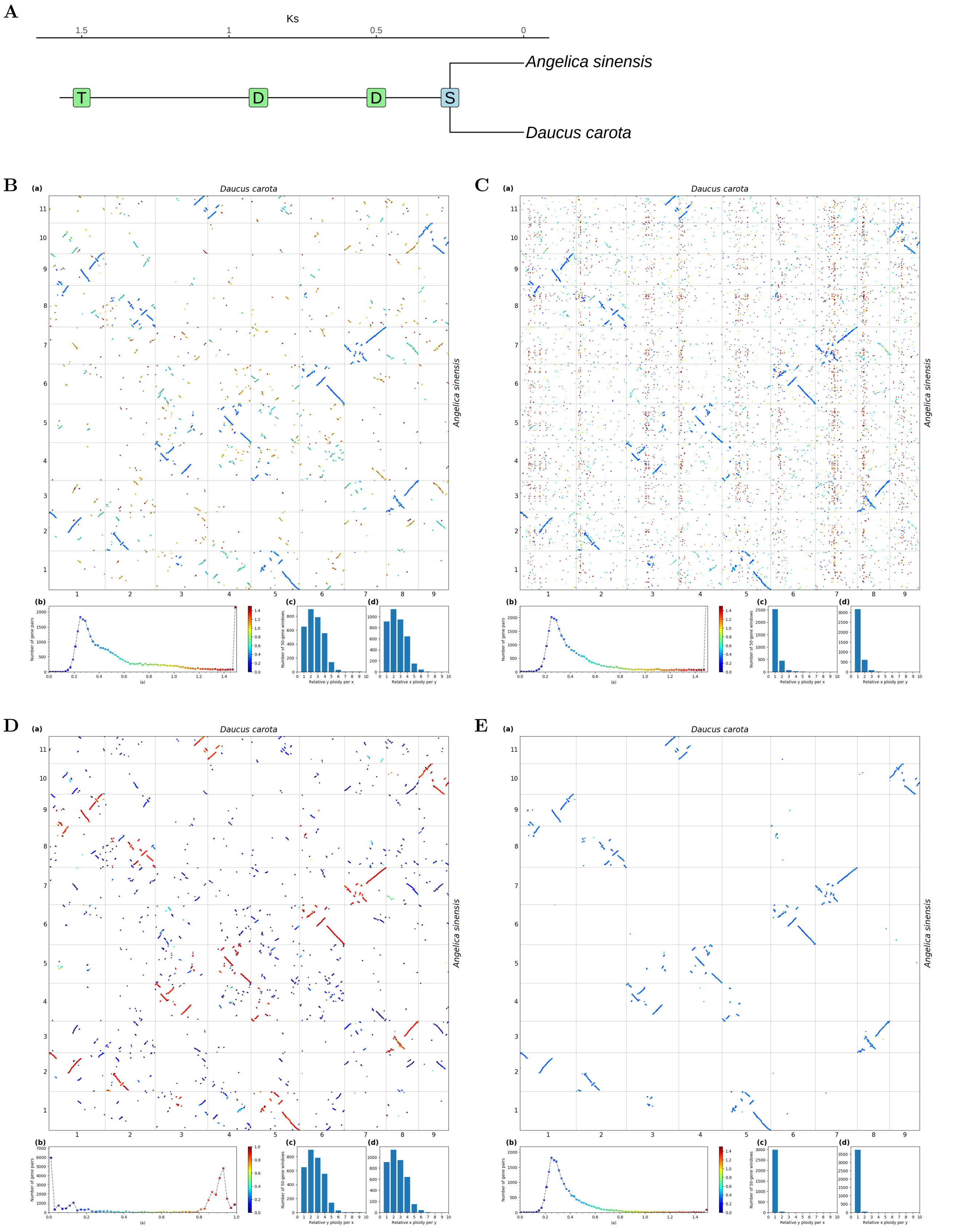


**Fig S56.** ***Orthology Index*** **in the identification of orthologous synteny in** ***Daucus carota*** **and** ***Angelica sinensis.*** Refer to **Fig.1** for detailed descriptions.

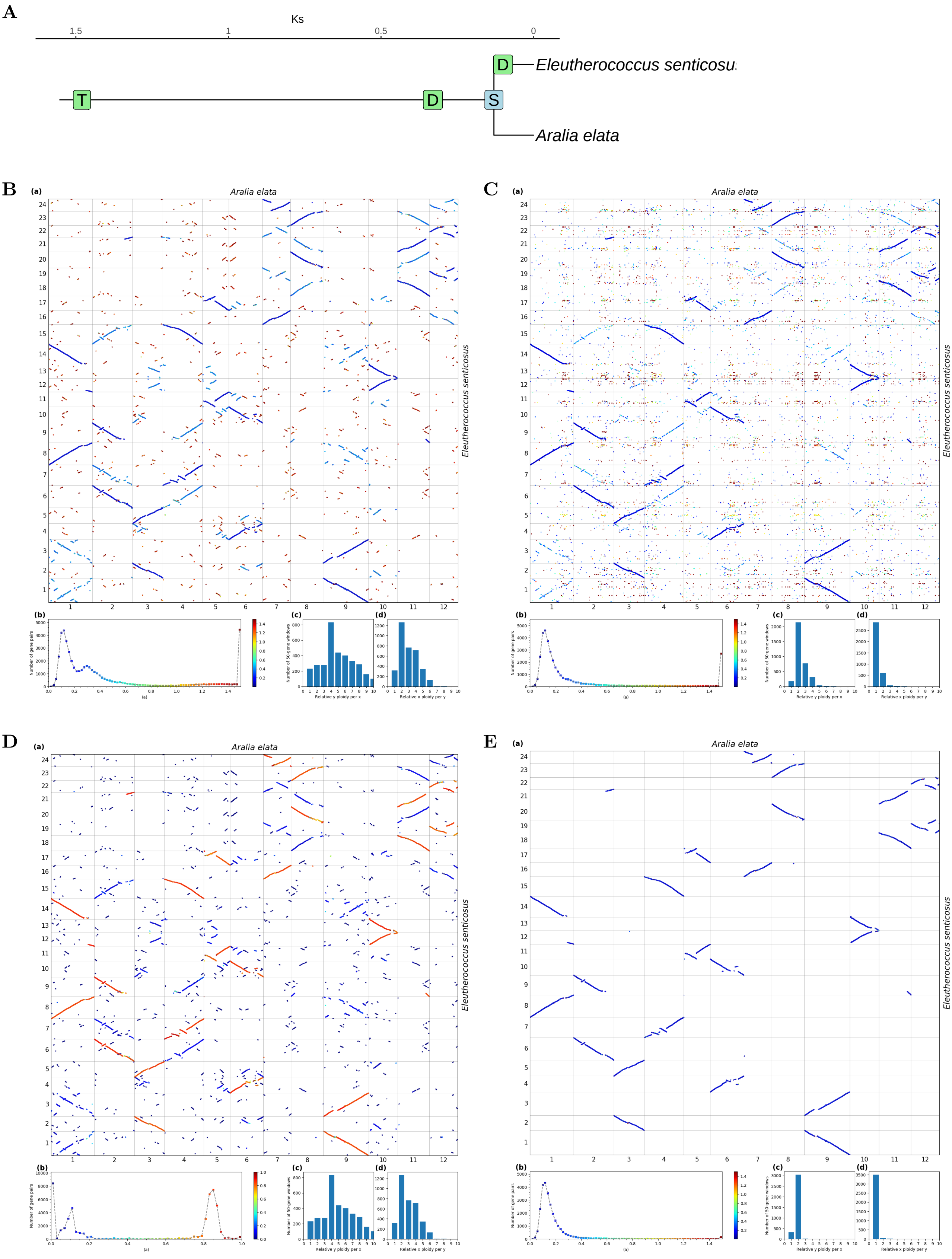

**Fig S57.** ***Orthology Index*** **in the identification of orthologous synteny in *Aralia elata* and *Eleutherococcus senticosus*.** Refer to **Fig.1** for detailed descriptions.

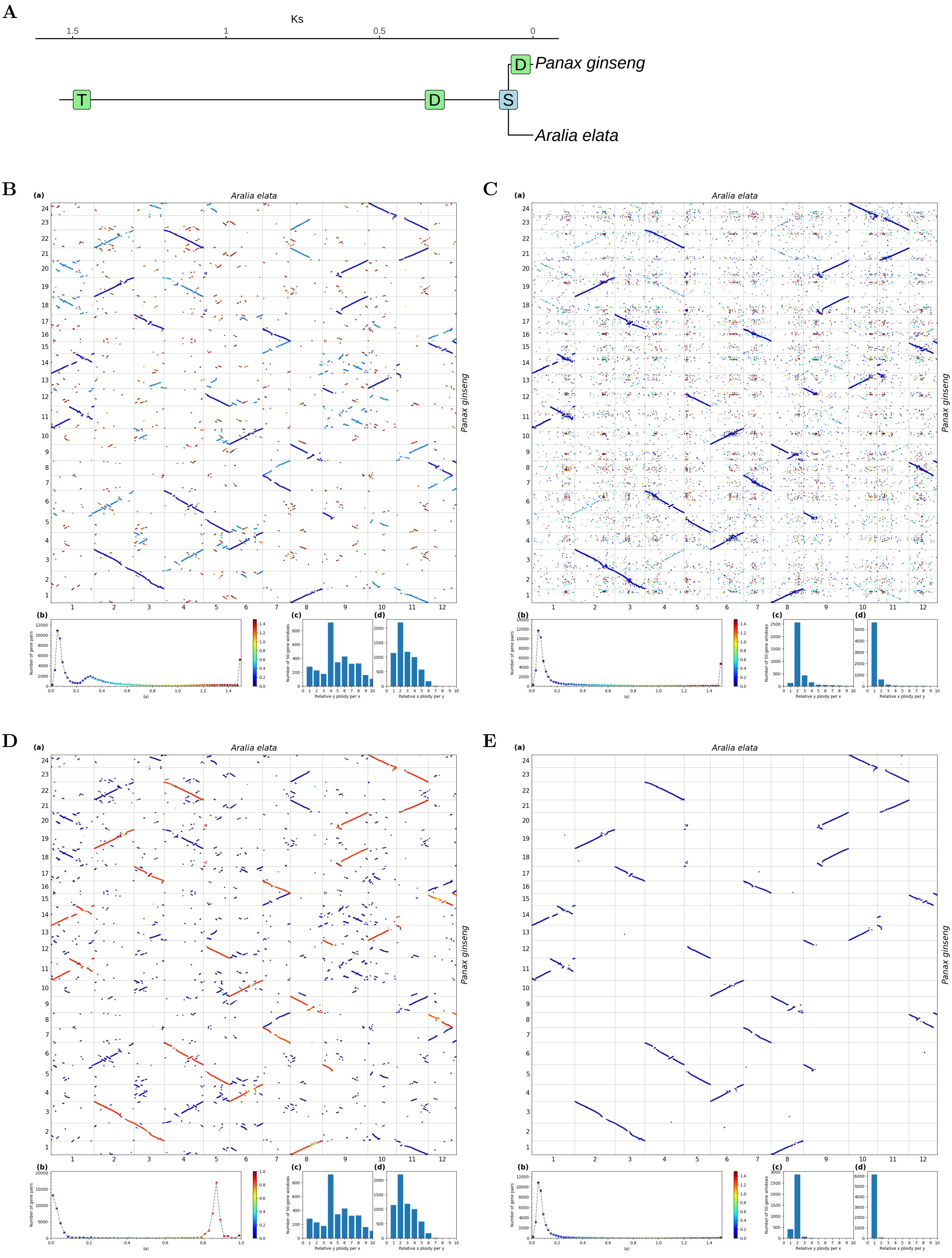


**Fig S58.** ***Orthology Index* in the identification of orthologous synteny in *Aralia elata* and *Panax ginseng*.** Refer to **Fig.1** for detailed descriptions.

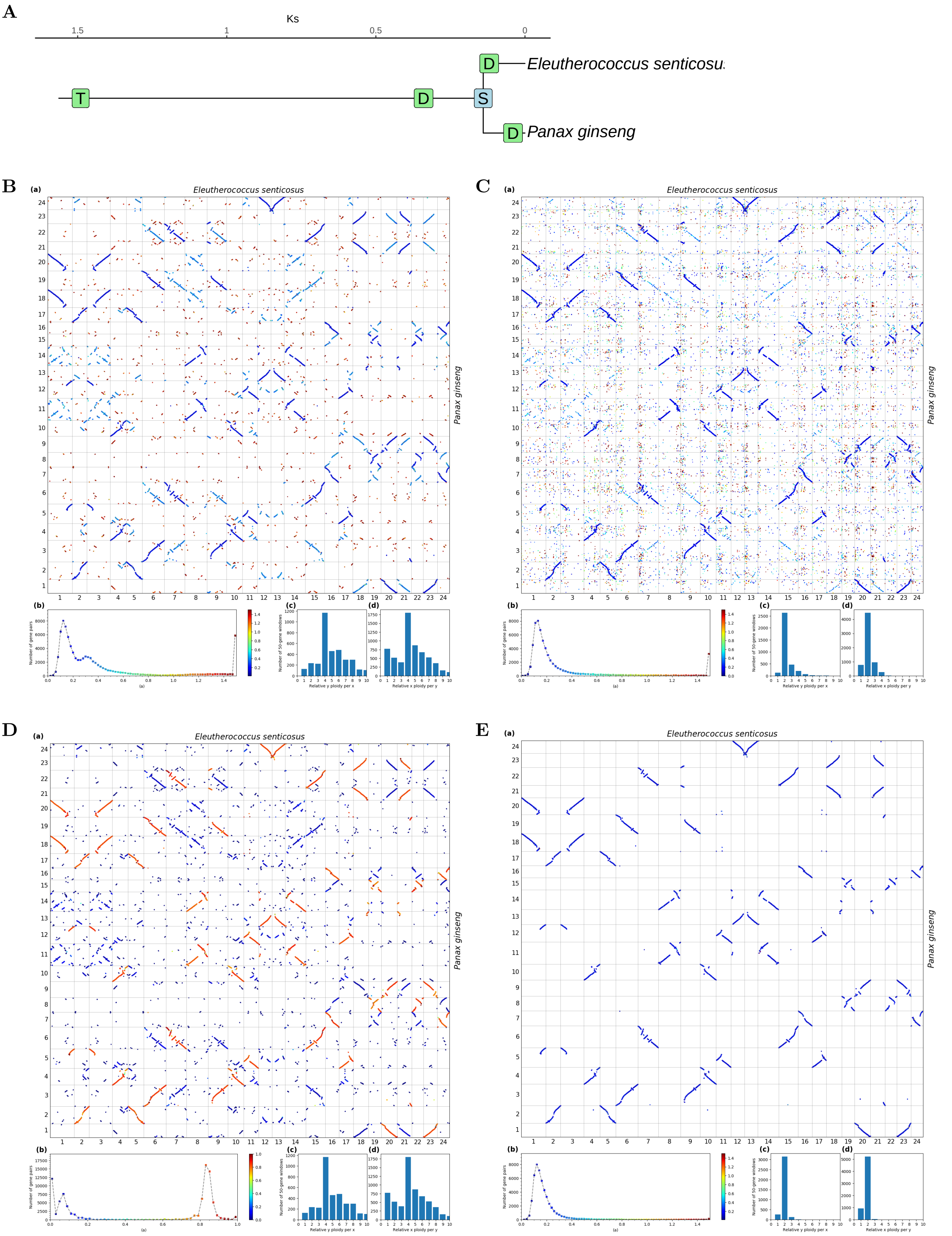


**Fig S59.** ***Orthology Index*** **in the identification of orthologous synteny in** ***Panax ginseng*** **and** ***Eleutherococcus senticosus.*** Refer to **Fig.1** for detailed descriptions.

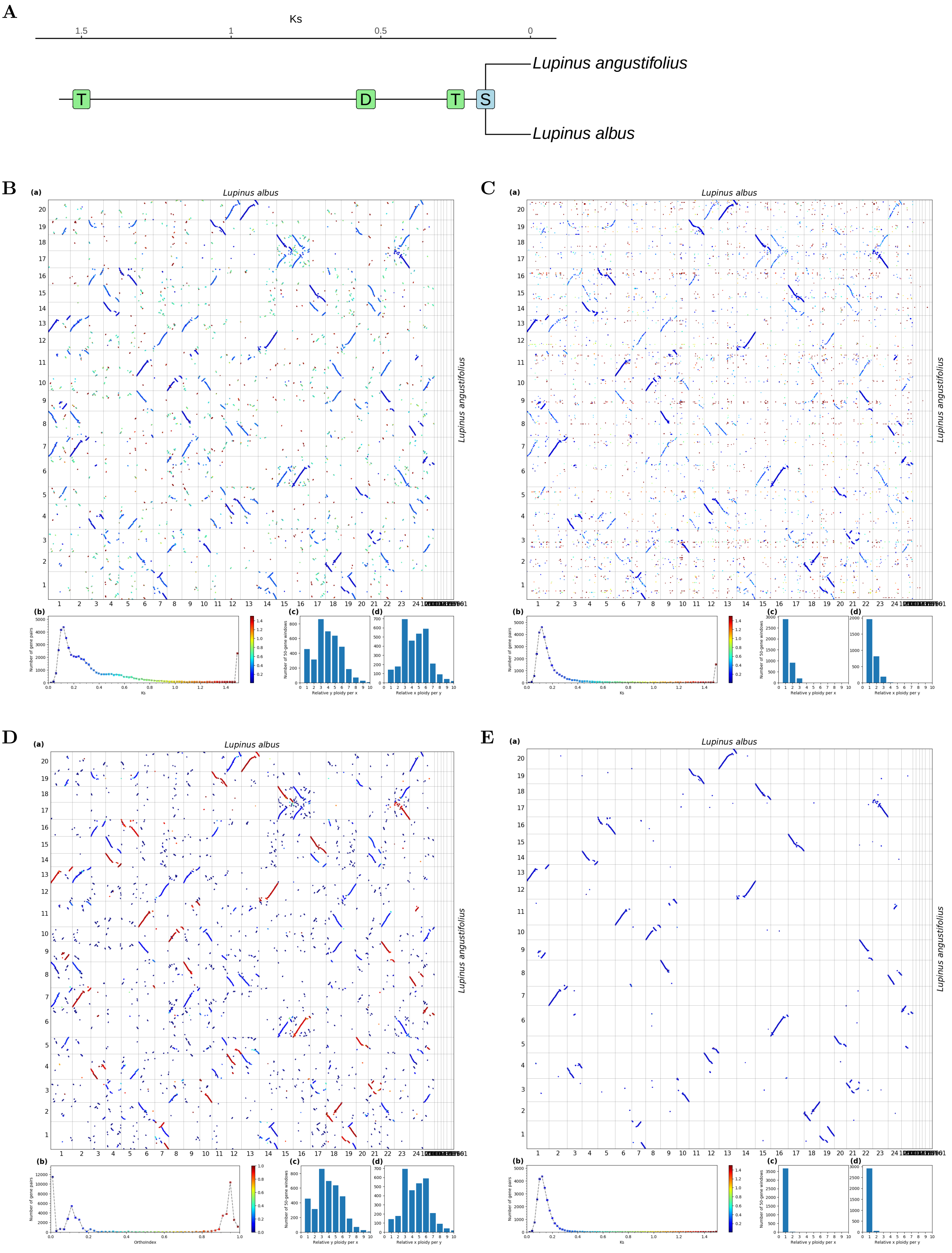


**Fig S60.** ***Orthology Index*** **in the identification of orthologous synteny in** ***Lupinus albus*** **and** ***Lupinus angustifolius.*** Refer to **Fig.1** for detailed descriptions.

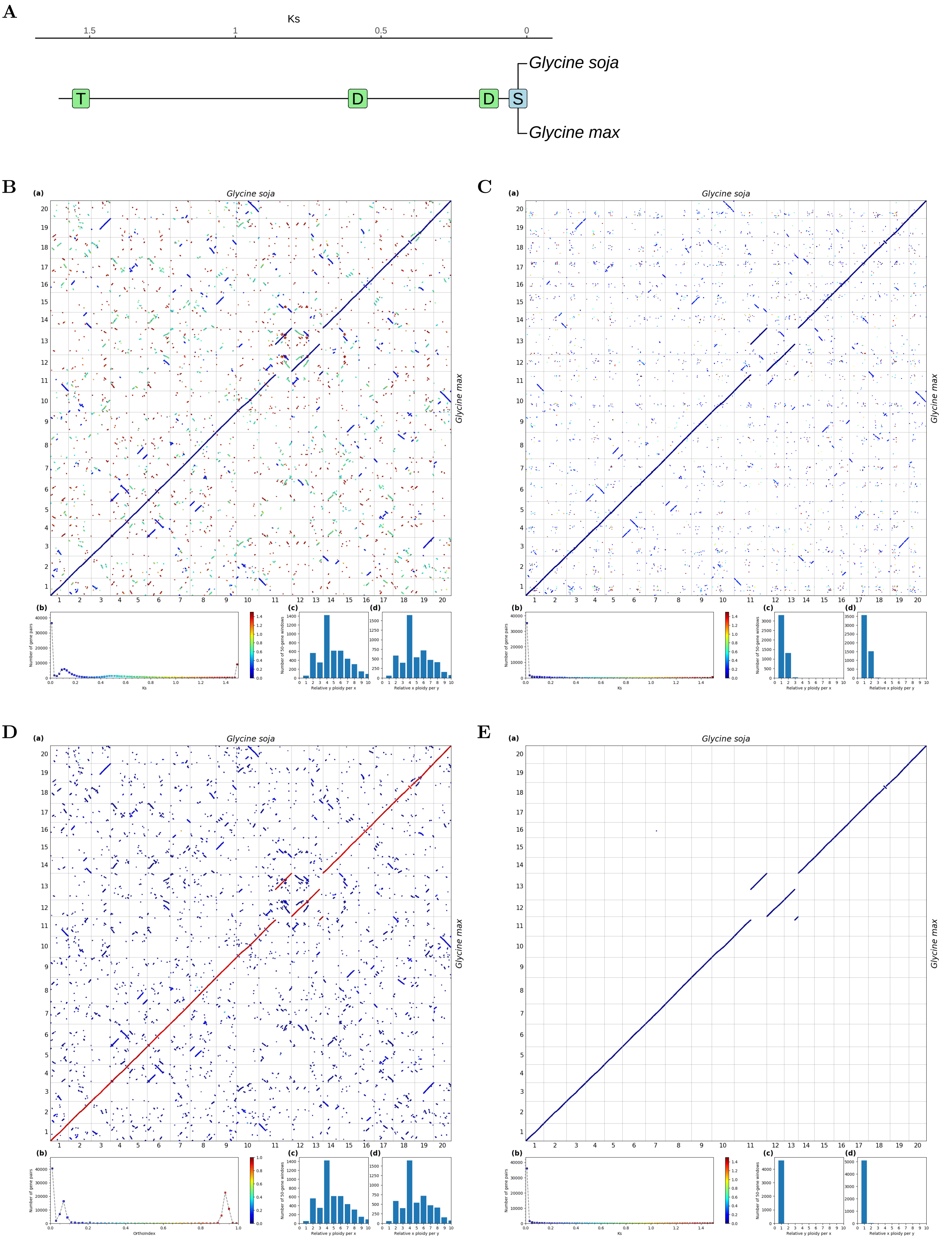


**Fig S61.** ***Orthology Index*** **in the identification of orthologous synteny in** ***Glycine max*** **and** ***Glycine soja*****.** Refer to **Fig.1** for detailed descriptions.

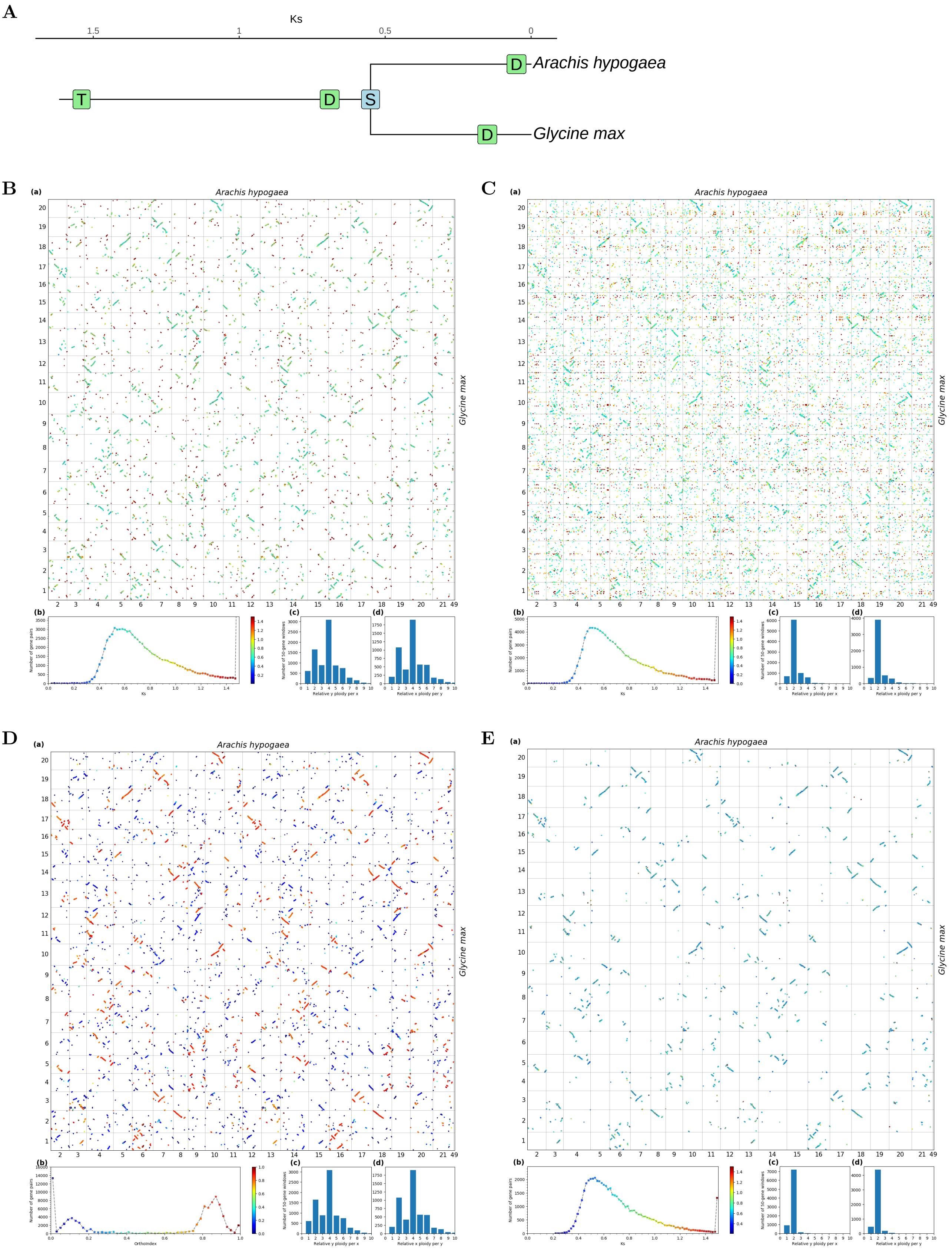


**Fig S62.** ***Orthology Index*** **in the identification of orthologous synteny in** ***Glycine max*** **and** ***Arachis hypogaea.*** Refer to **Fig.1** for detailed descriptions.

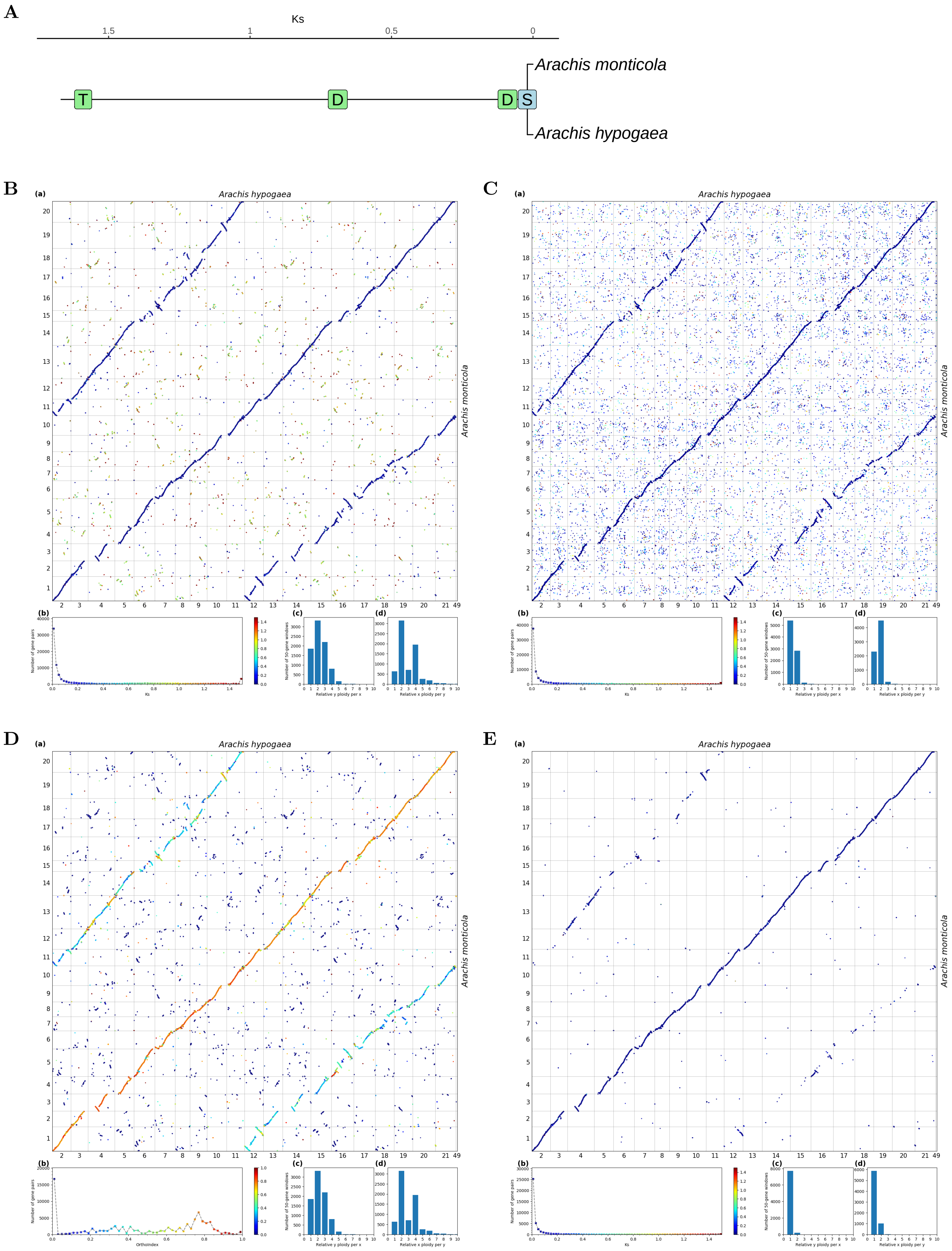


**Fig S63.** ***Orthology Index*** **in the identification of orthologous synteny in** ***Arachis hypogaea*** **and** ***Arachis monticola.*** Refer to **Fig.1** for detailed descriptions.

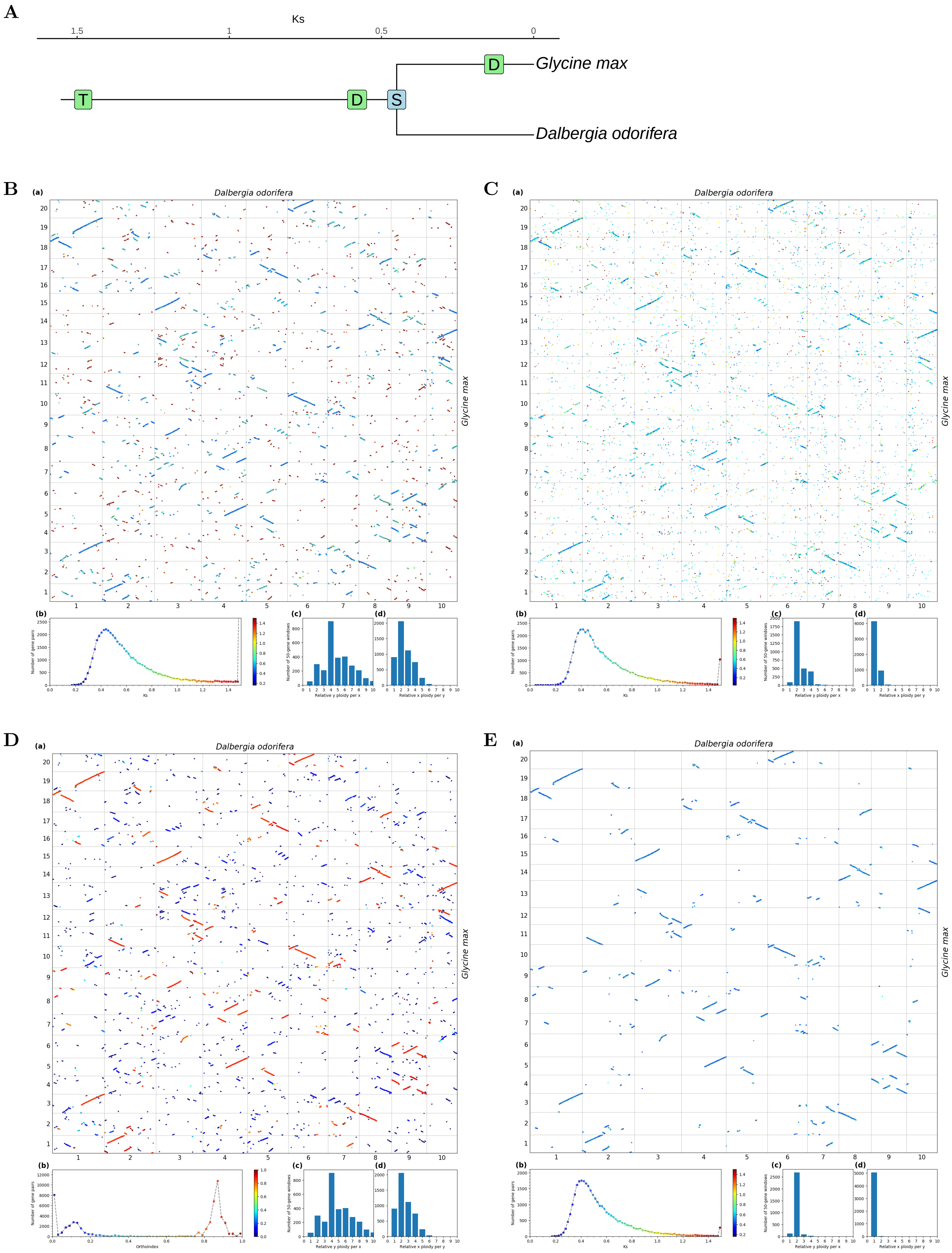


**Fig S64.** ***Orthology Index*** **in the identification of orthologous synteny in** ***Dalbergia odorifera*** **and** ***Glycine max.*** Refer to **Fig.1** for detailed descriptions.

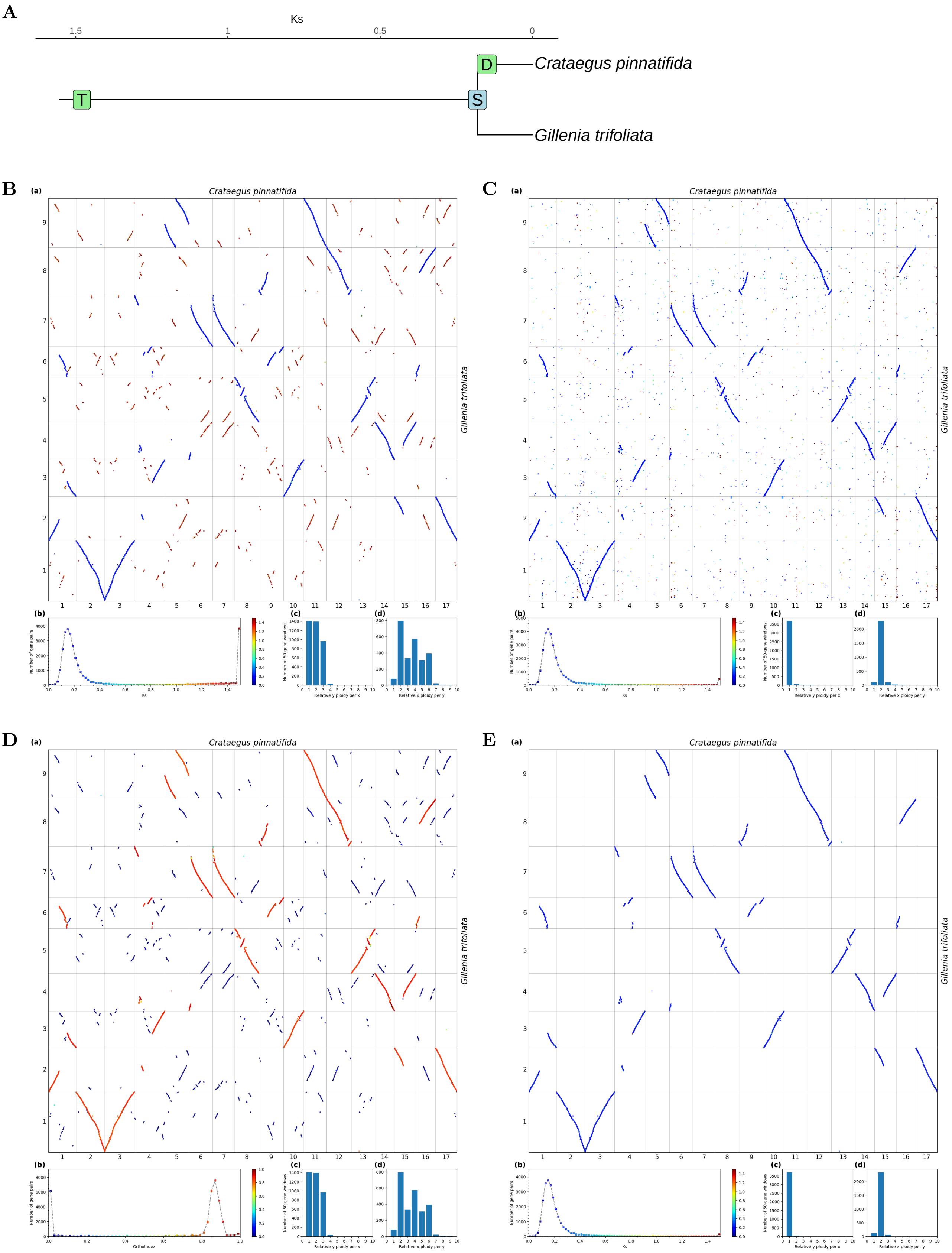


**Fig S65.** ***Orthology Index*** **in the identification of orthologous synteny in** ***Gillenia trifoliata*** **and** ***Crataegus pinnatifida.*** Refer to **Fig.1** for detailed descriptions.

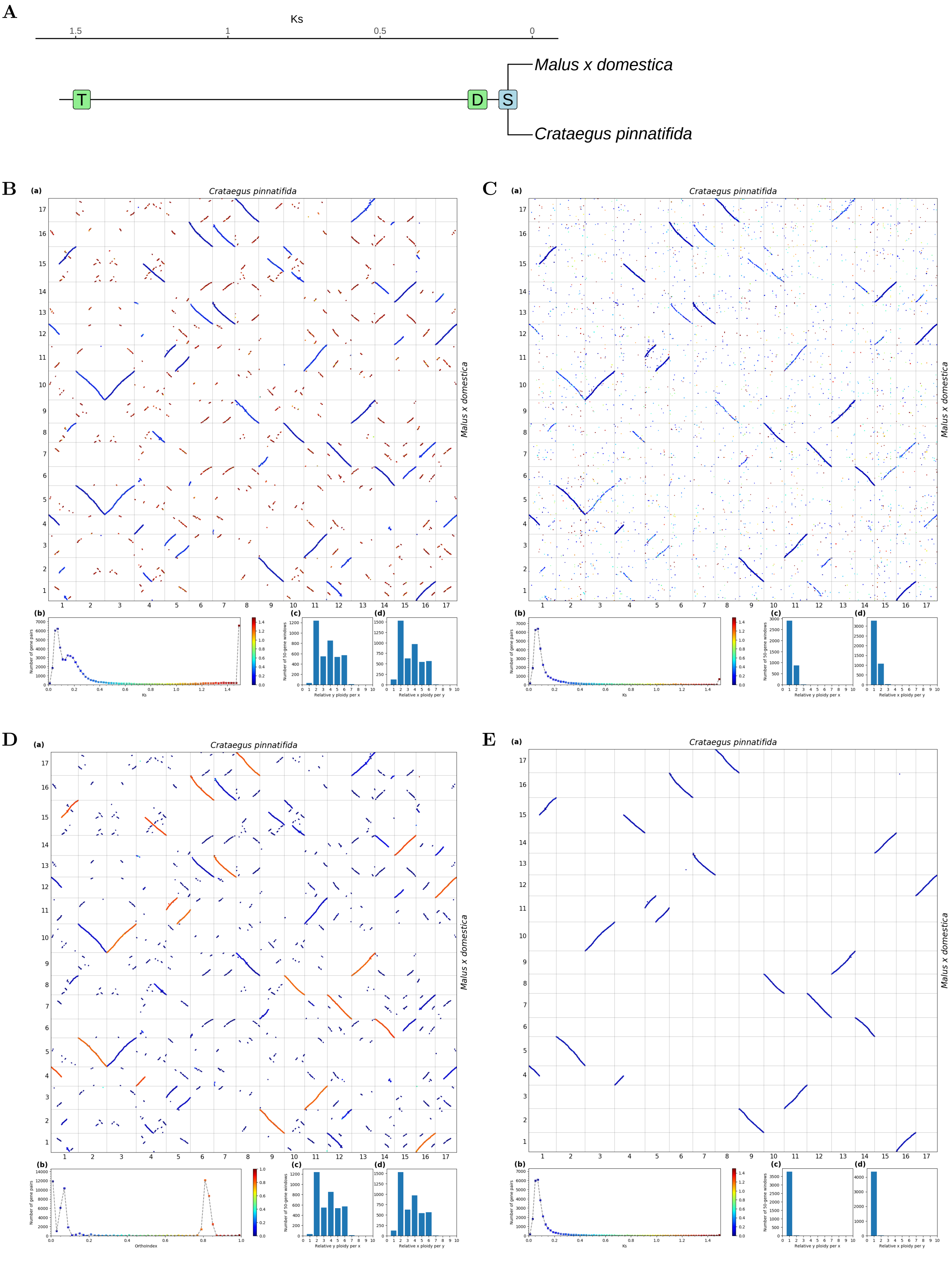


**Fig S66.** ***Orthology Index*** **in the identification of orthologous synteny in** ***Crataegus pinnatifida*** **and** ***Malus x domestica.*** Refer to **Fig.1** for detailed descriptions.

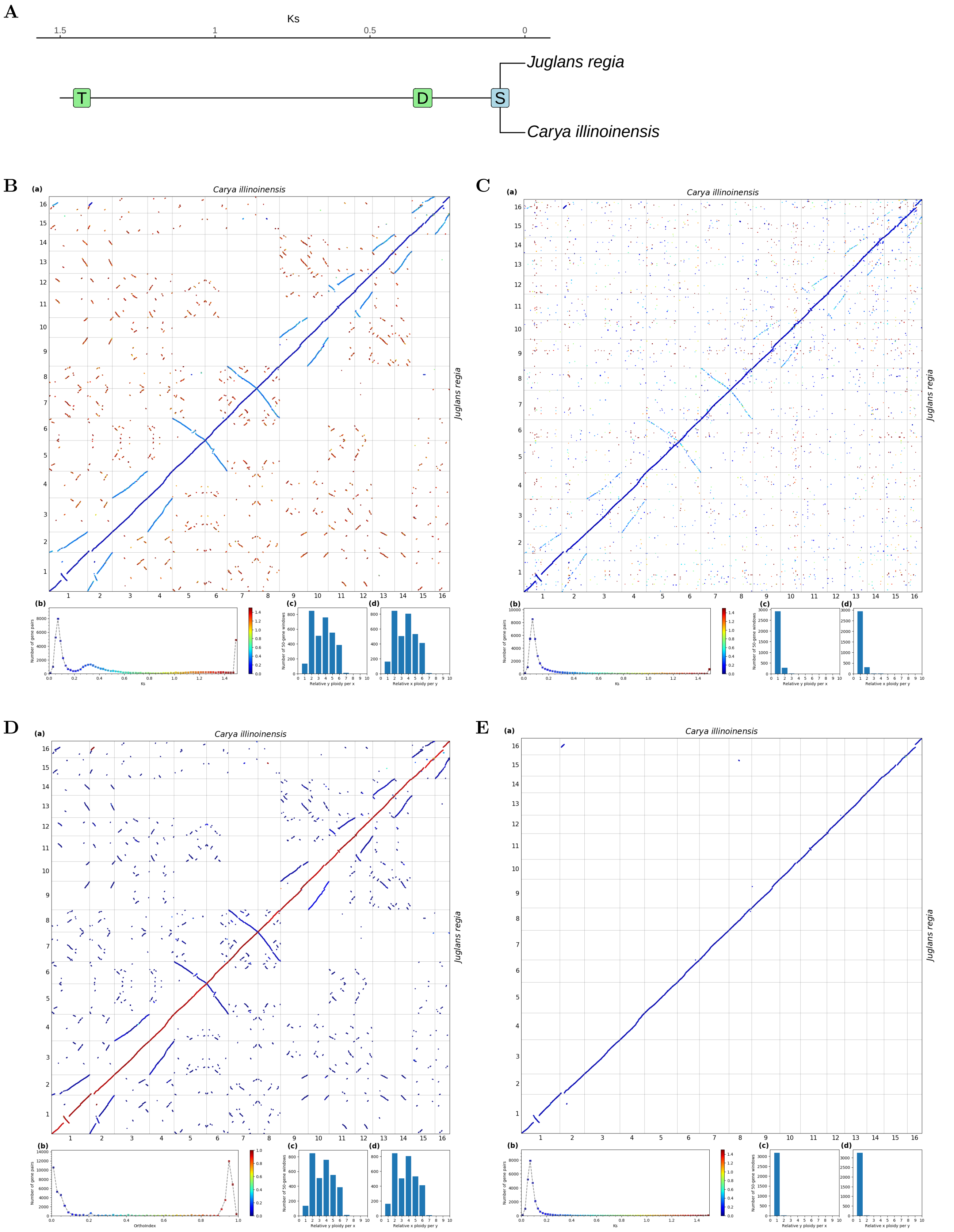


**Fig S67.** ***Orthology Index*** **in the identification of orthologous synteny in** ***Carya illinoinensis*** **and** ***Juglans regia.*** Refer to **Fig.1** for detailed descriptions.

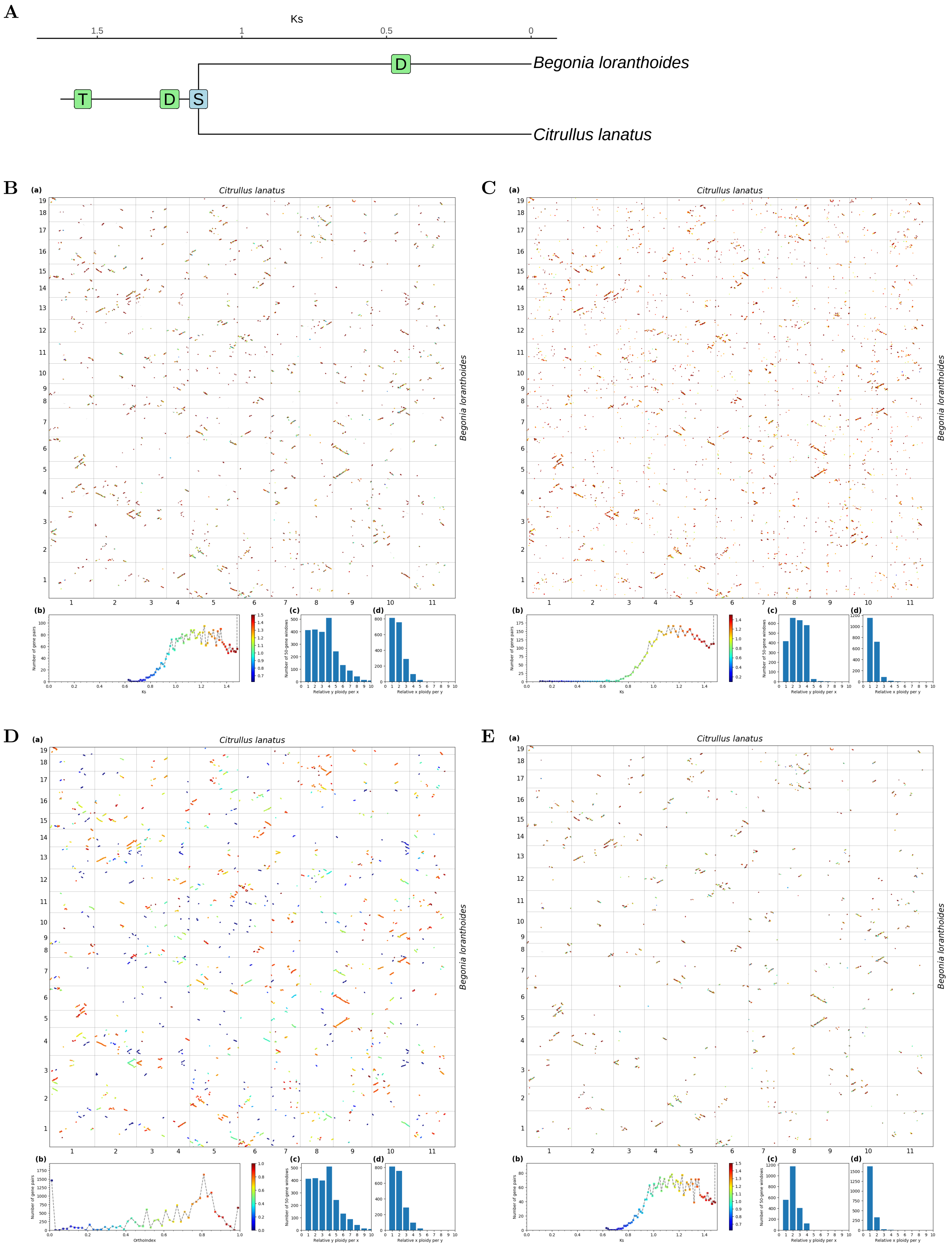


**Fig S68.** ***Orthology Index* in the identification of orthologous synteny in *Citrullus lanatus* and *Begonia loranthoides*.** Refer to **Fig.1** for detailed descriptions.

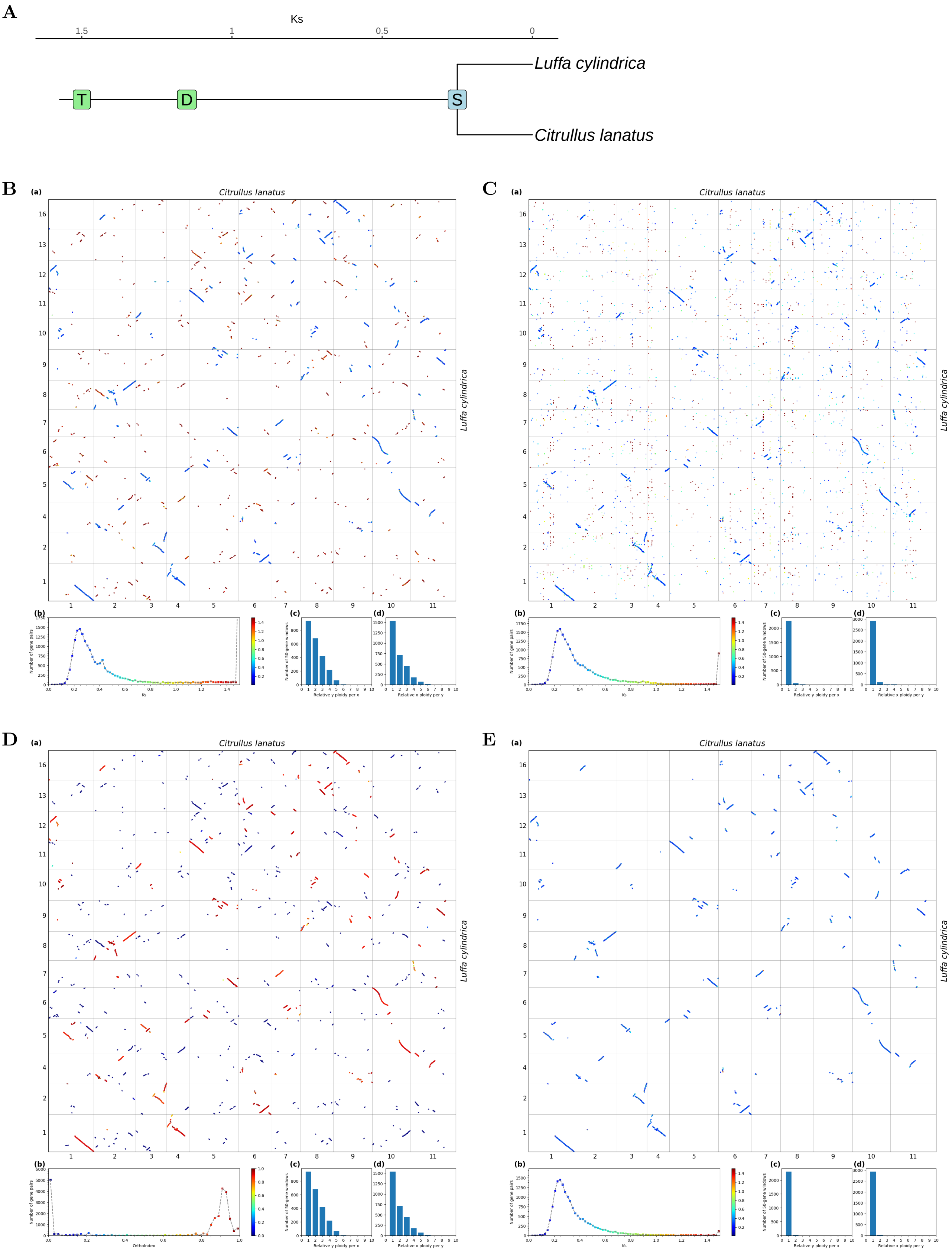


**Fig S69.** ***Orthology Index*** **in the identification of orthologous synteny in** ***Citrullus lanatus*** **and** ***Luffa cylindrica*****.** Refer to **Fig.1** for detailed descriptions.

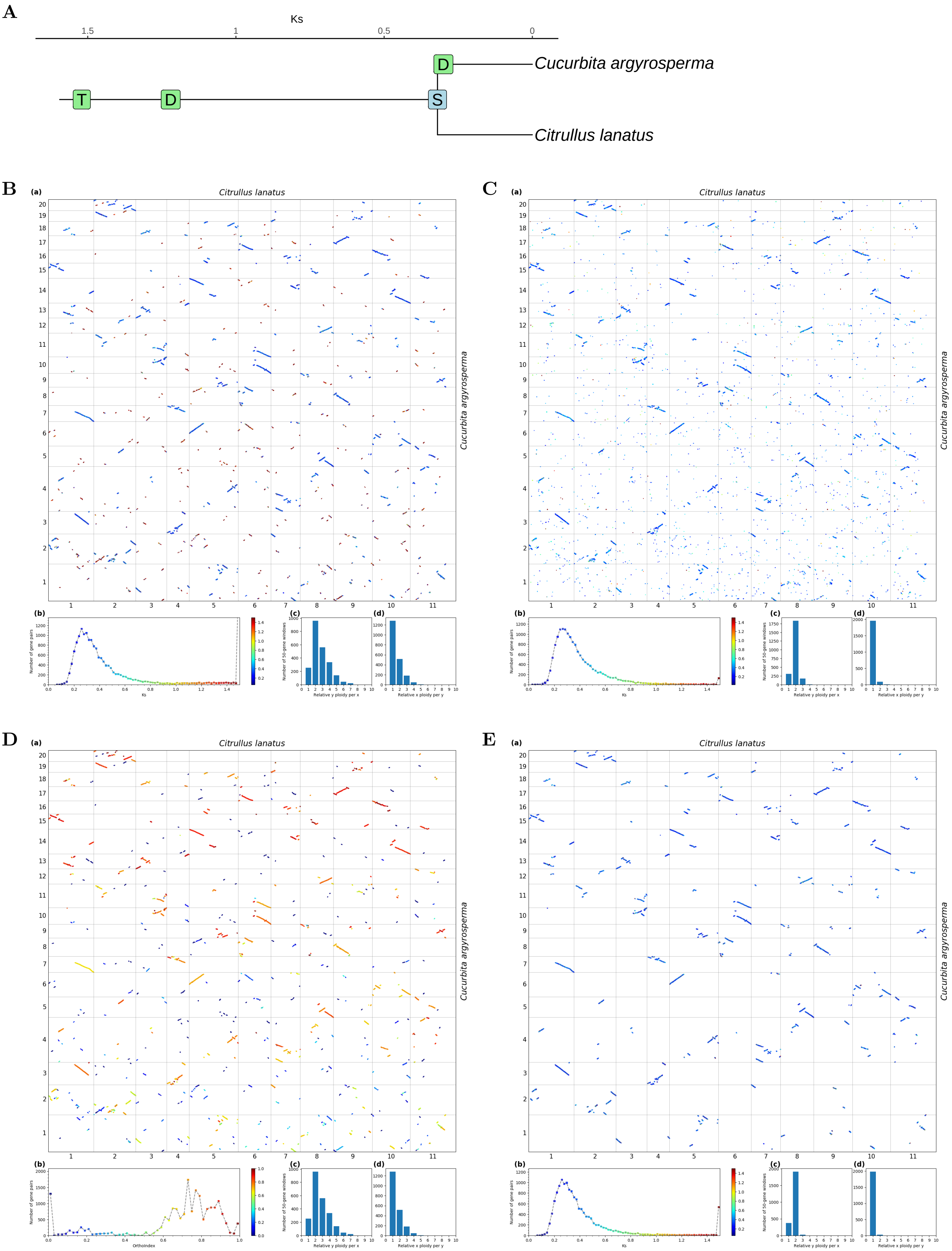


**Fig S70.** ***Orthology Index* in the identification of orthologous synteny in *Citrullus lanatus* and *Cucurbita argyrosperma*.** Refer to **Fig.1** for detailed descriptions.

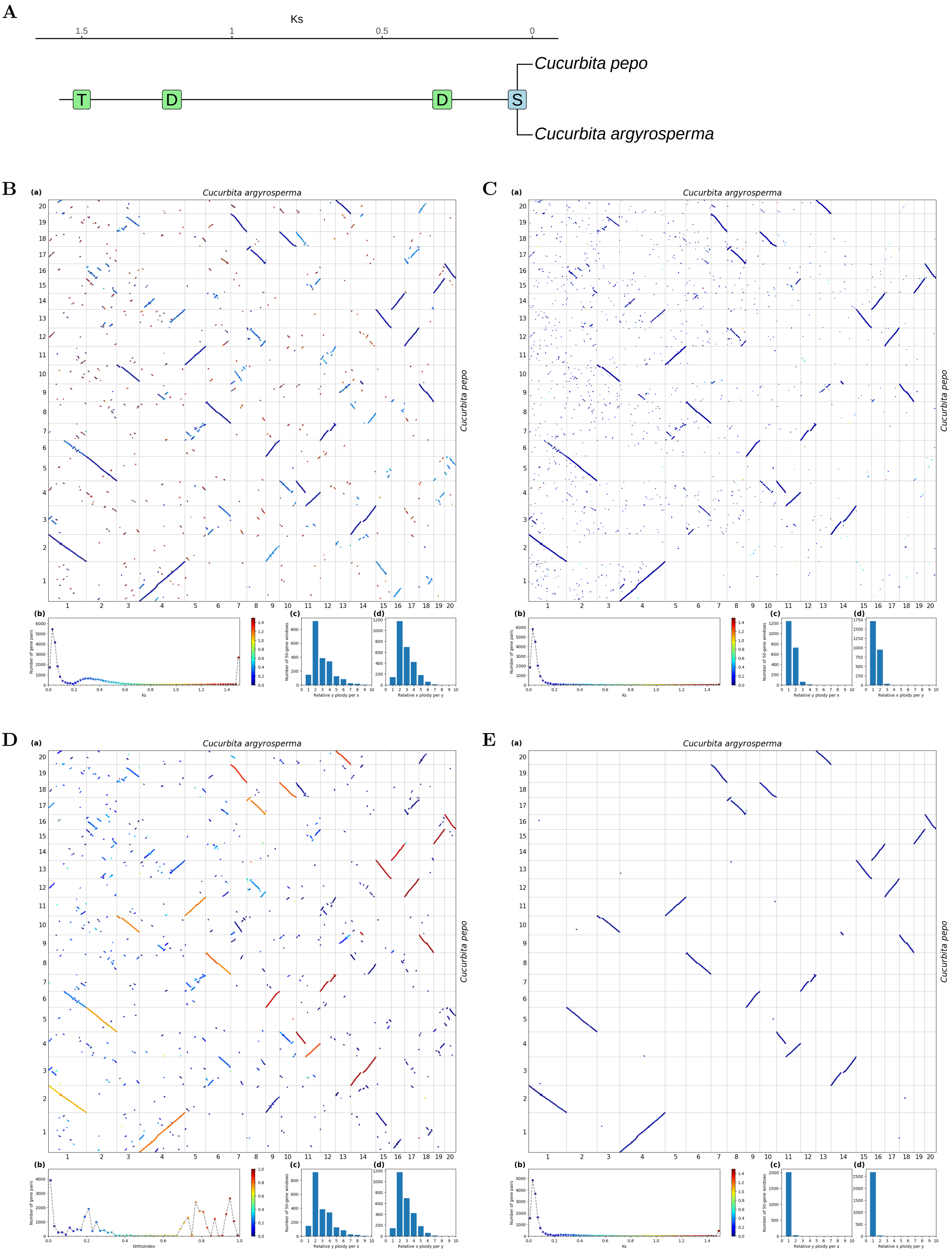


**Fig S71.** ***Orthology Index*** **in the identification of orthologous synteny in** ***Cucurbita argyrosperma*** **and** ***Cucurbita pepo*****.** Refer to **Fig.1** for detailed descriptions.

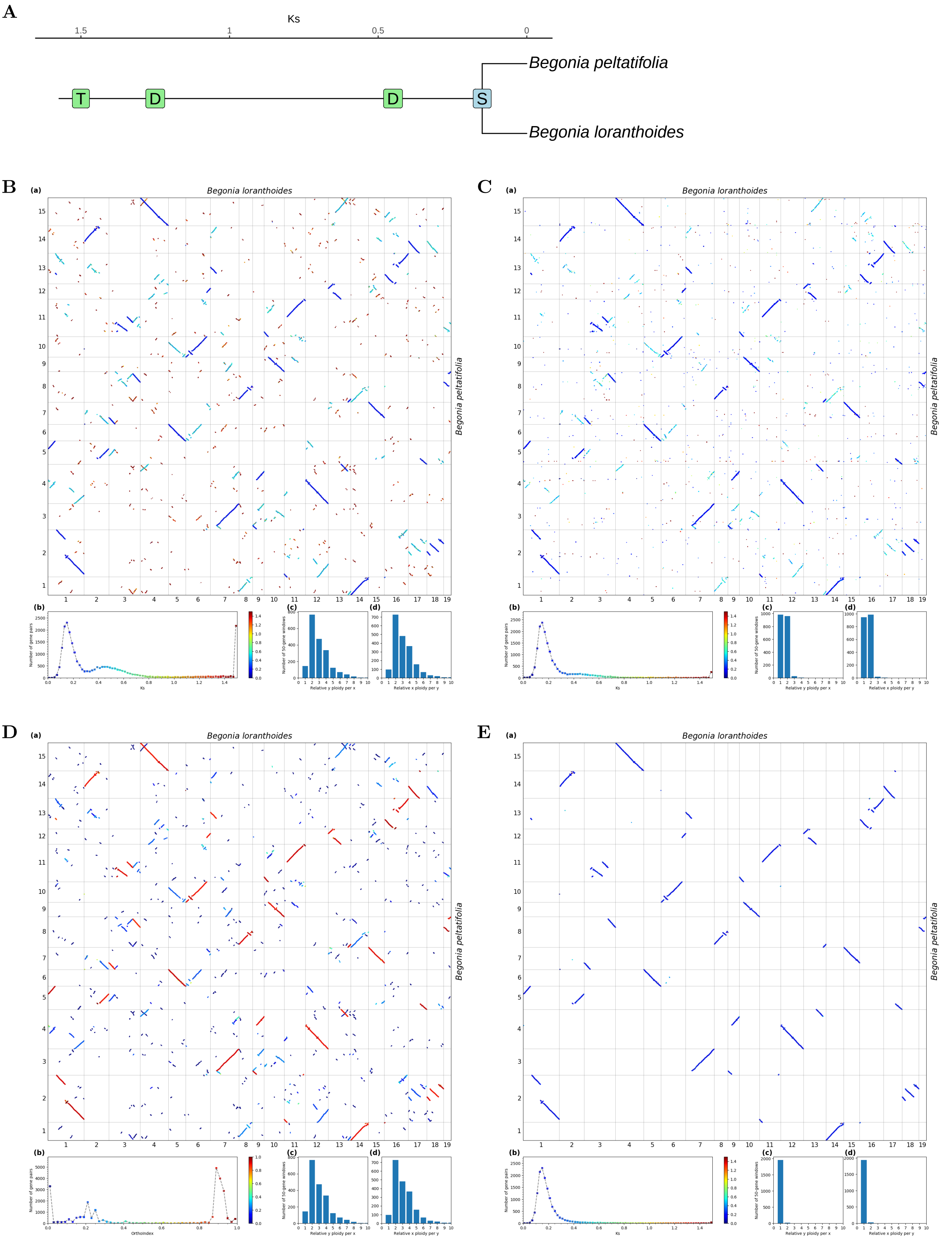


**Fig S72.** ***Orthology Index*** **in the identification of orthologous synteny in** ***Begonia loranthoides*** **and** ***Begonia peltatifolia.*** Refer to **Fig.1** for detailed descriptions.

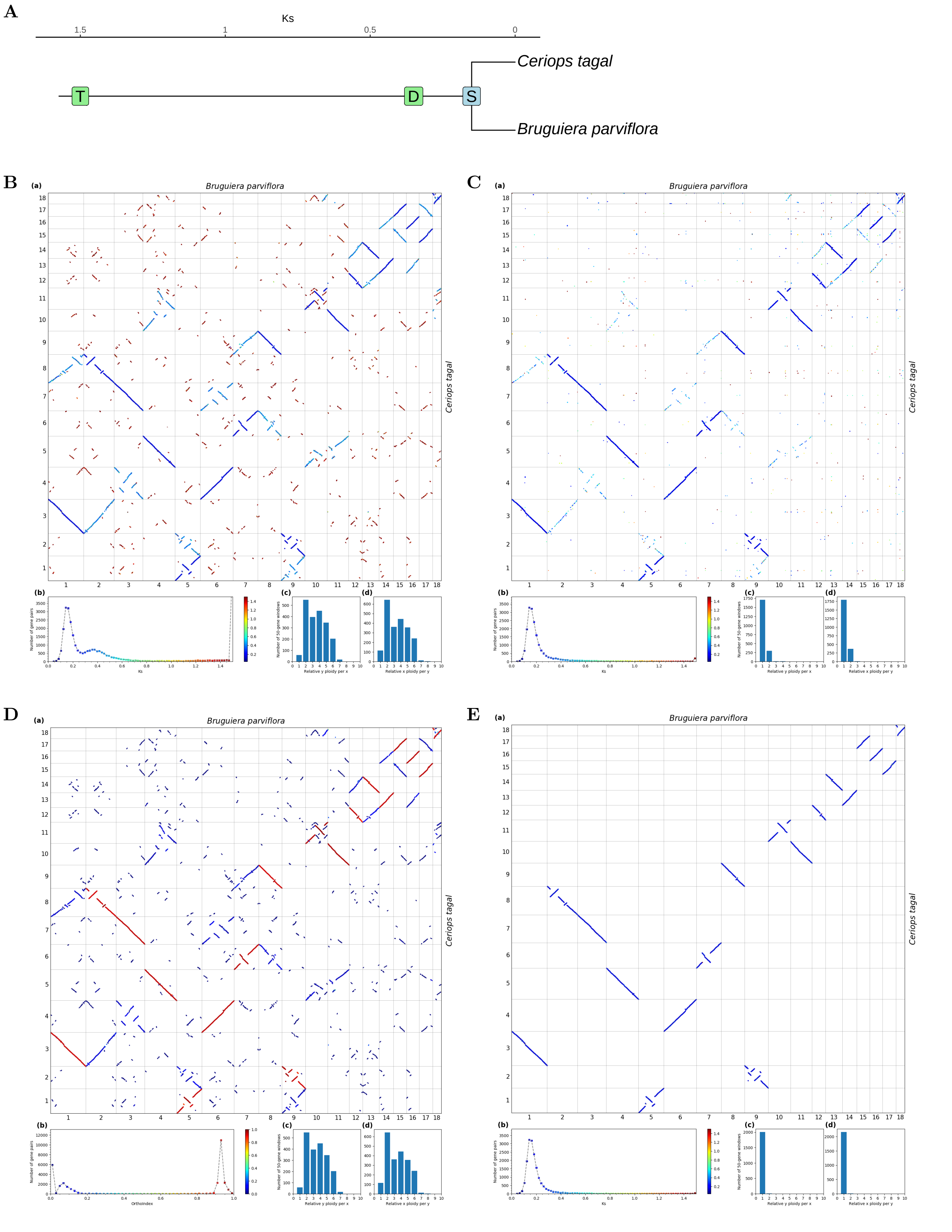


**Fig S73.** ***Orthology Index* in the identification of orthologous synteny in *Bruguiera parviflora* and *Ceriops tagal*.** Refer to **Fig.1** for detailed descriptions.

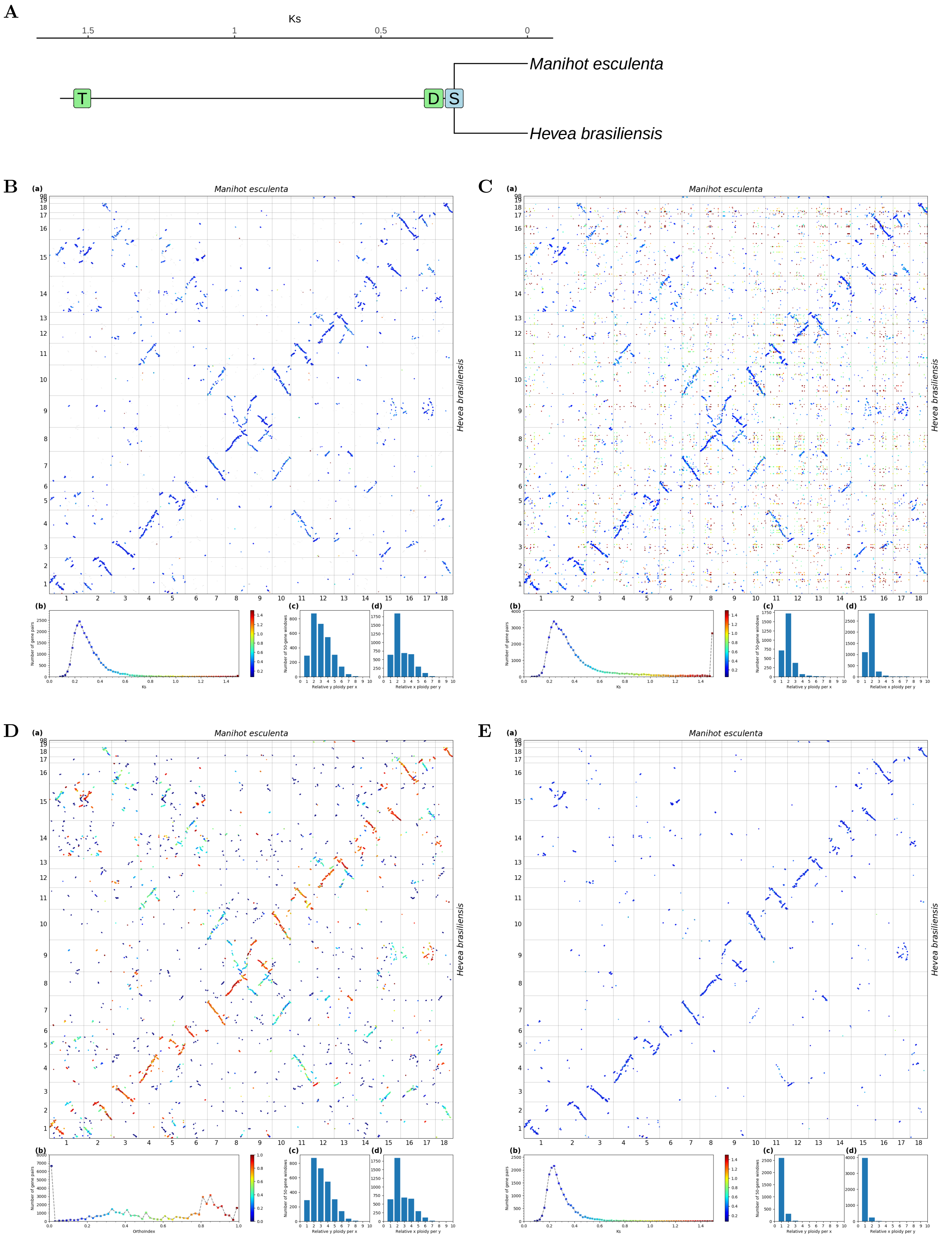


**Fig S74.** ***Orthology Index*** **in the identification of orthologous synteny in** ***Hevea brasiliensis*** **and** ***Manihot esculenta*****.** Refer to **Fig.1** for detailed descriptions.

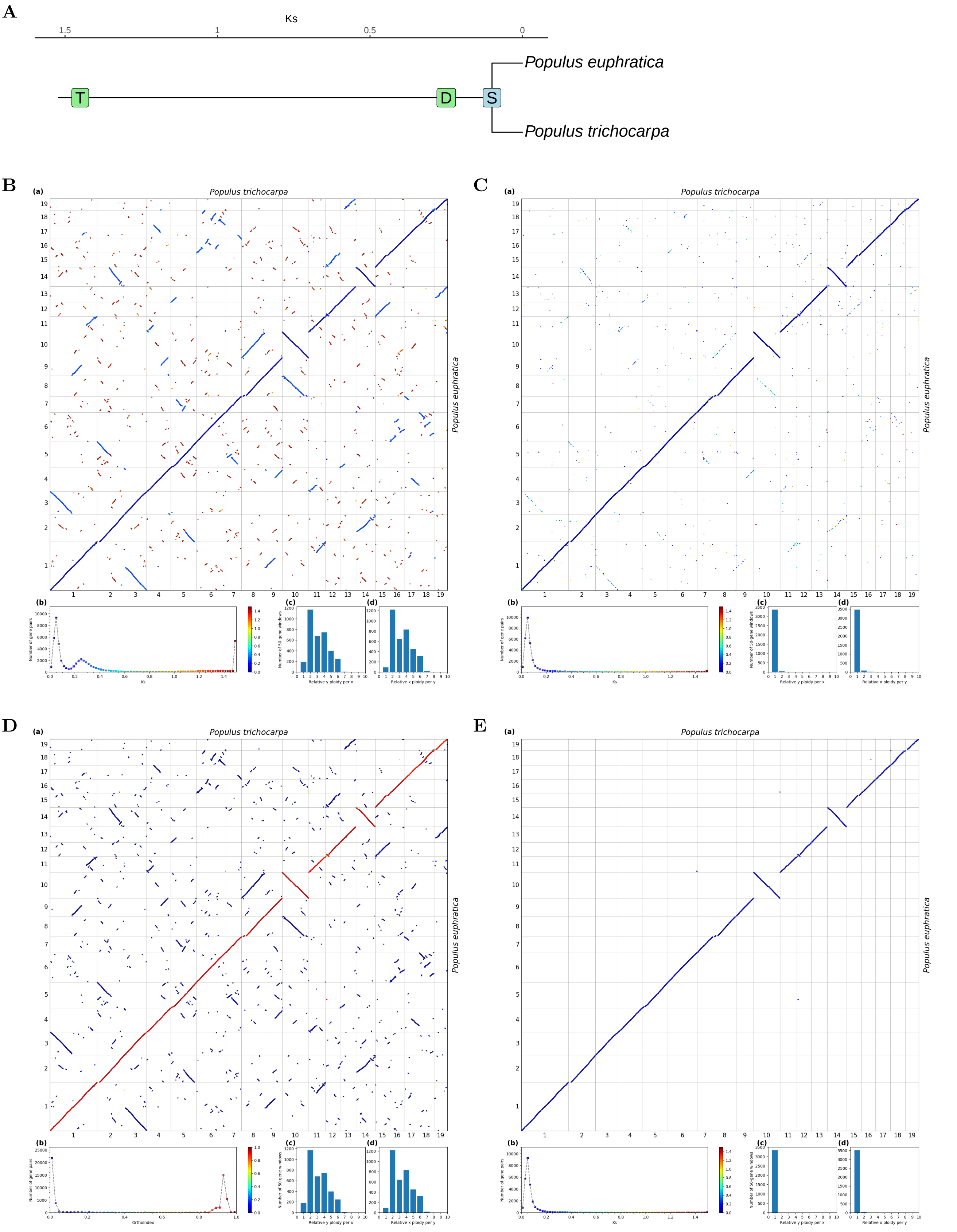

**Fig S75. *Orthology Index* in the identification of orthologous synteny in *Populus trichocarpa* and *Populus euphratica*.** Refer to **Fig.1** for detailed descriptions.

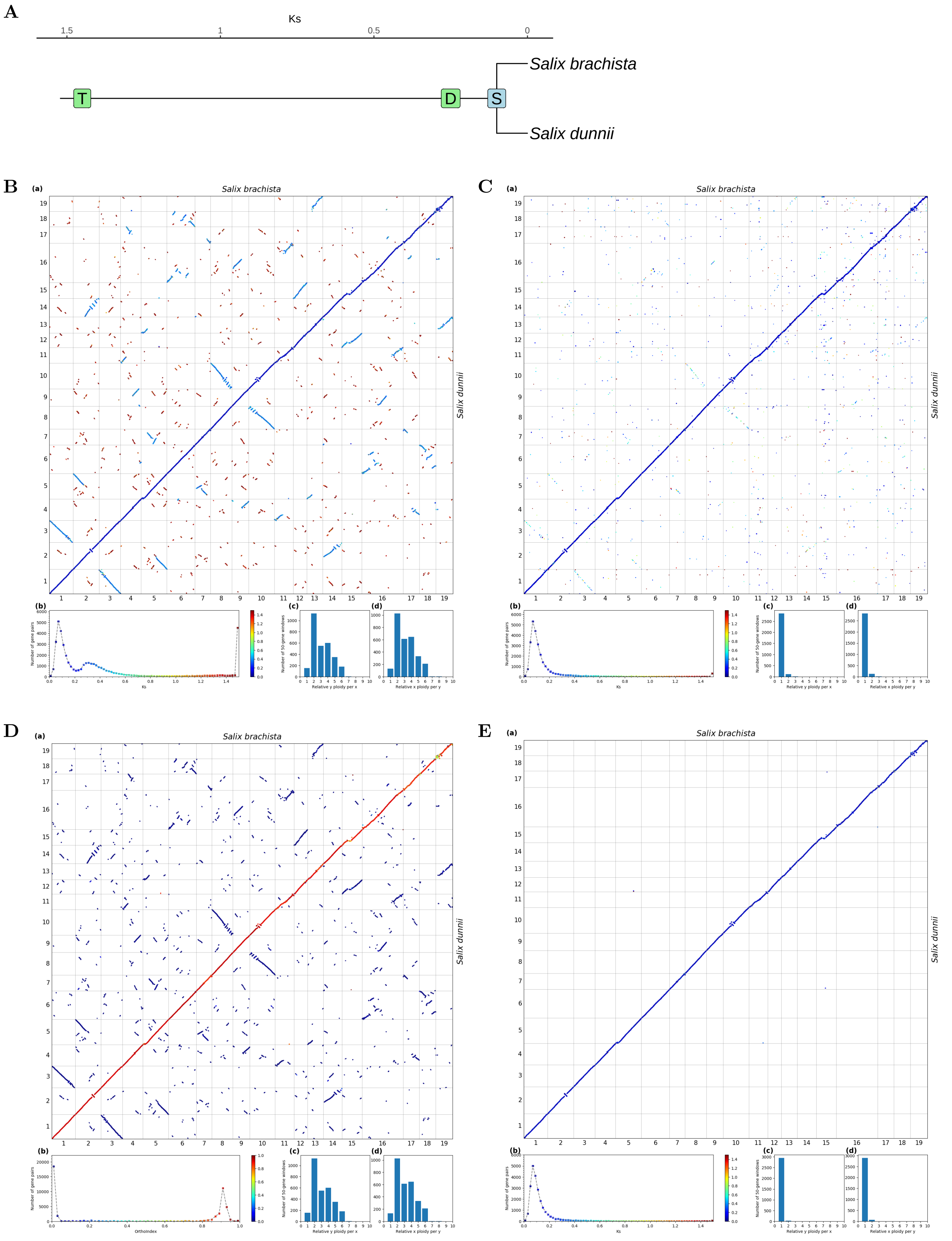


**Fig S76.** ***Orthology Index* in the identification of orthologous synteny in *Salix dunnii* and *Salix brachista*.** Refer to **Fig.1** for detailed descriptions.

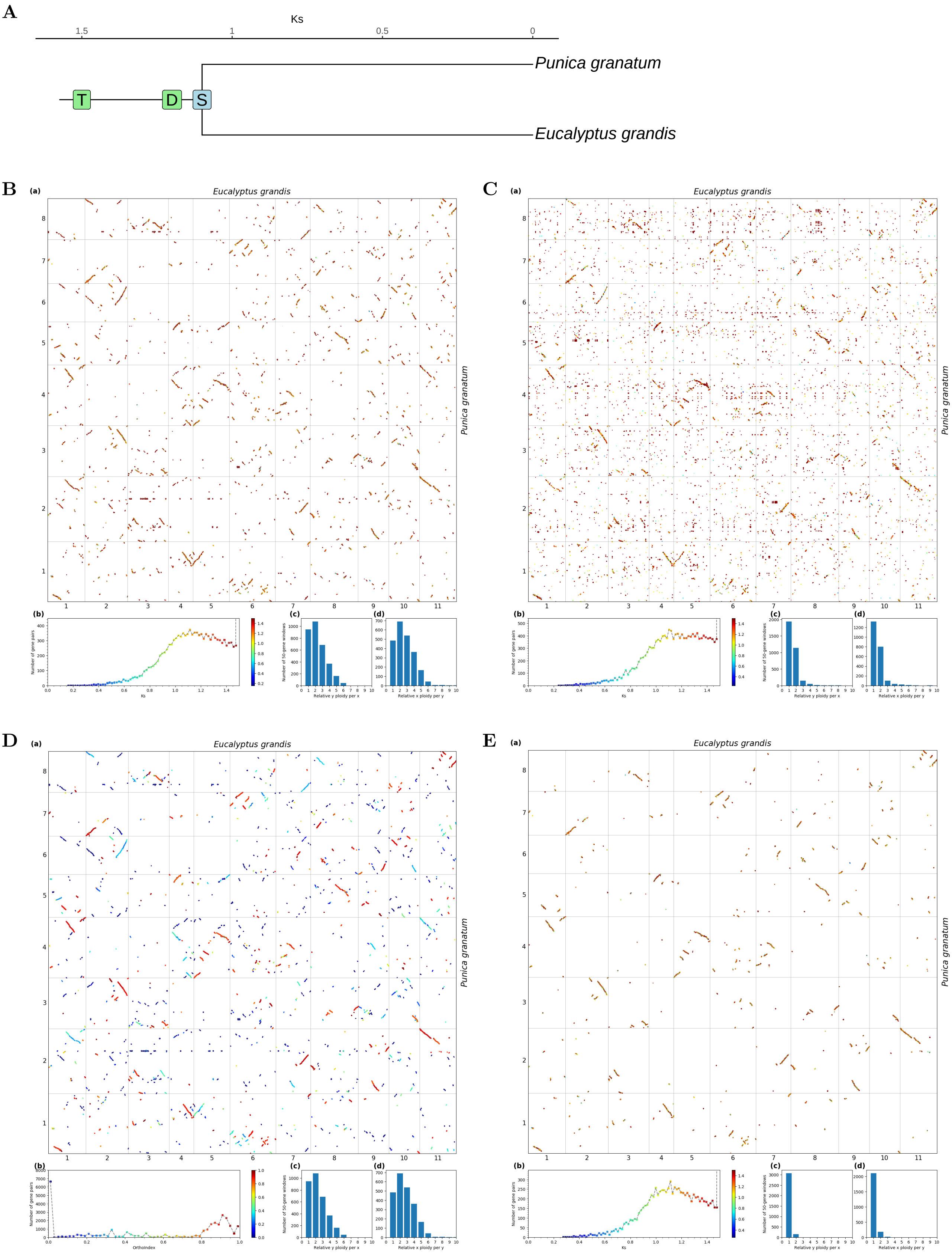


**Fig S77.** ***Orthology Index* in the identification of orthologous synteny in *Eucalyptus grandis* and *Punica granatum*.** Refer to **Fig.1** for detailed descriptions.

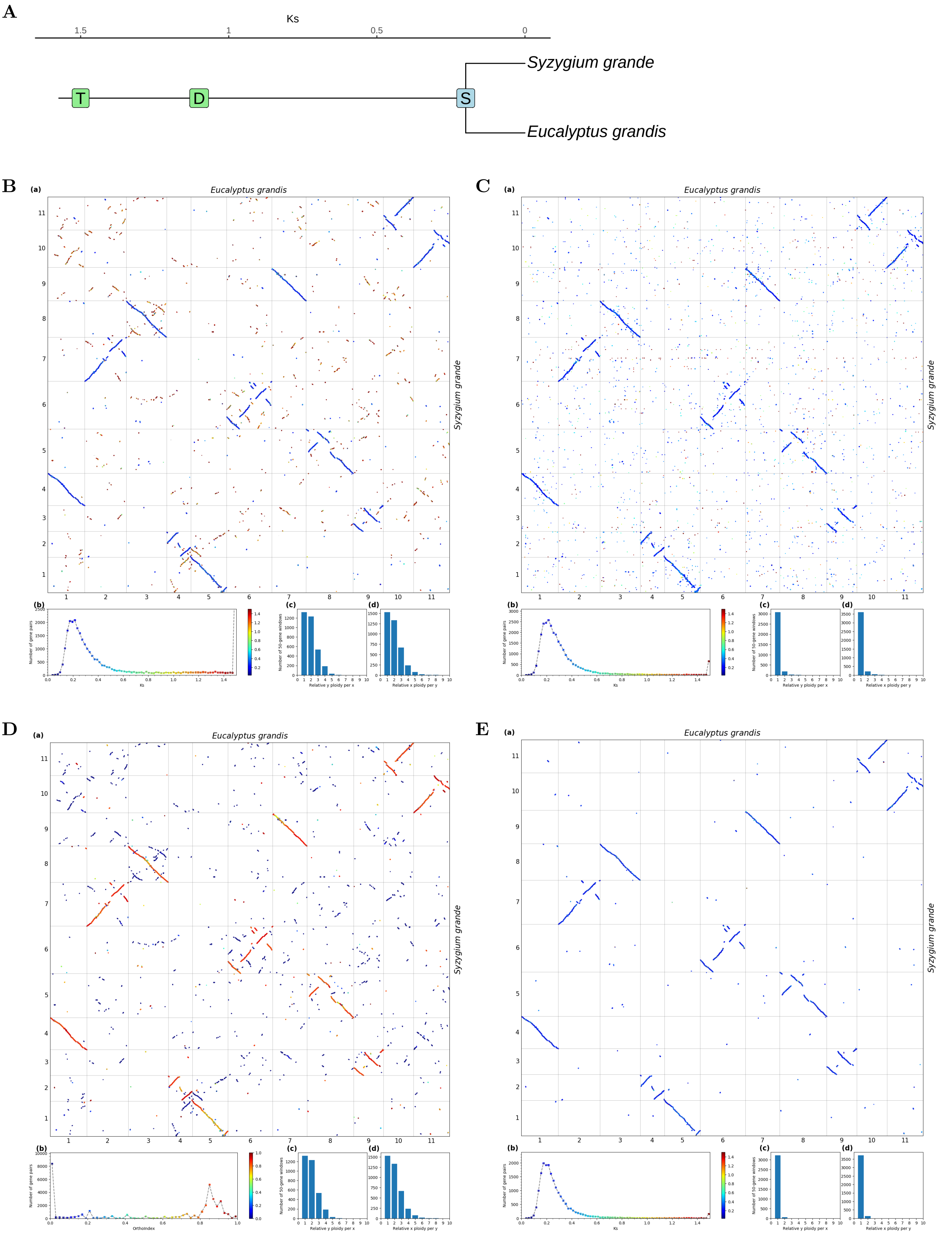


**Fig S78.** ***Orthology Index*** **in the identification of orthologous synteny in** ***Eucalyptus grandis*** **and** ***Syzygium grande*****.** Refer to **Fig.1** for detailed descriptions.

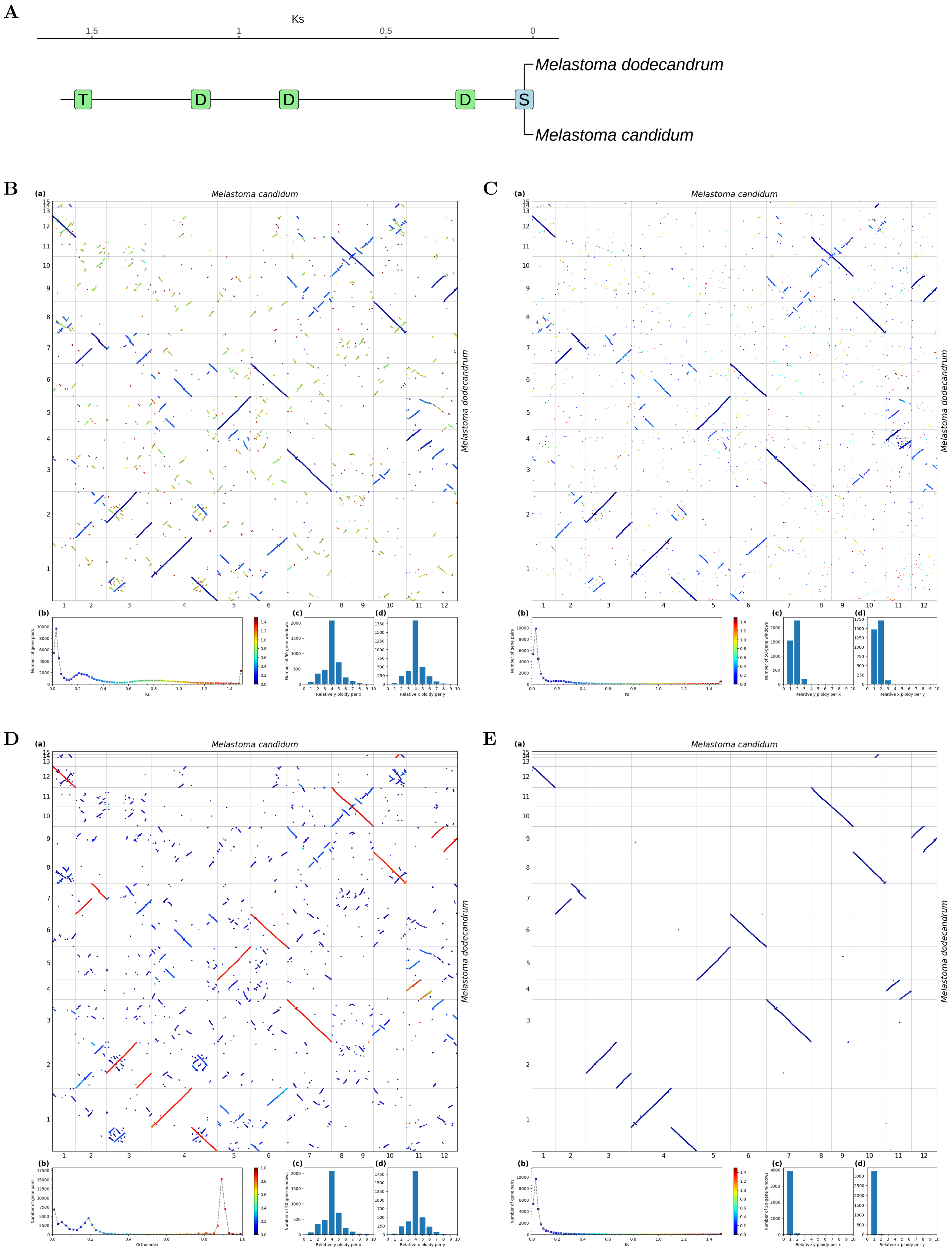


**Fig S79.** ***Orthology Index* in the identification of orthologous synteny in *Melastoma candidum* and *Melastoma dodecandrum*.** Refer to **Fig.1** for detailed descriptions.

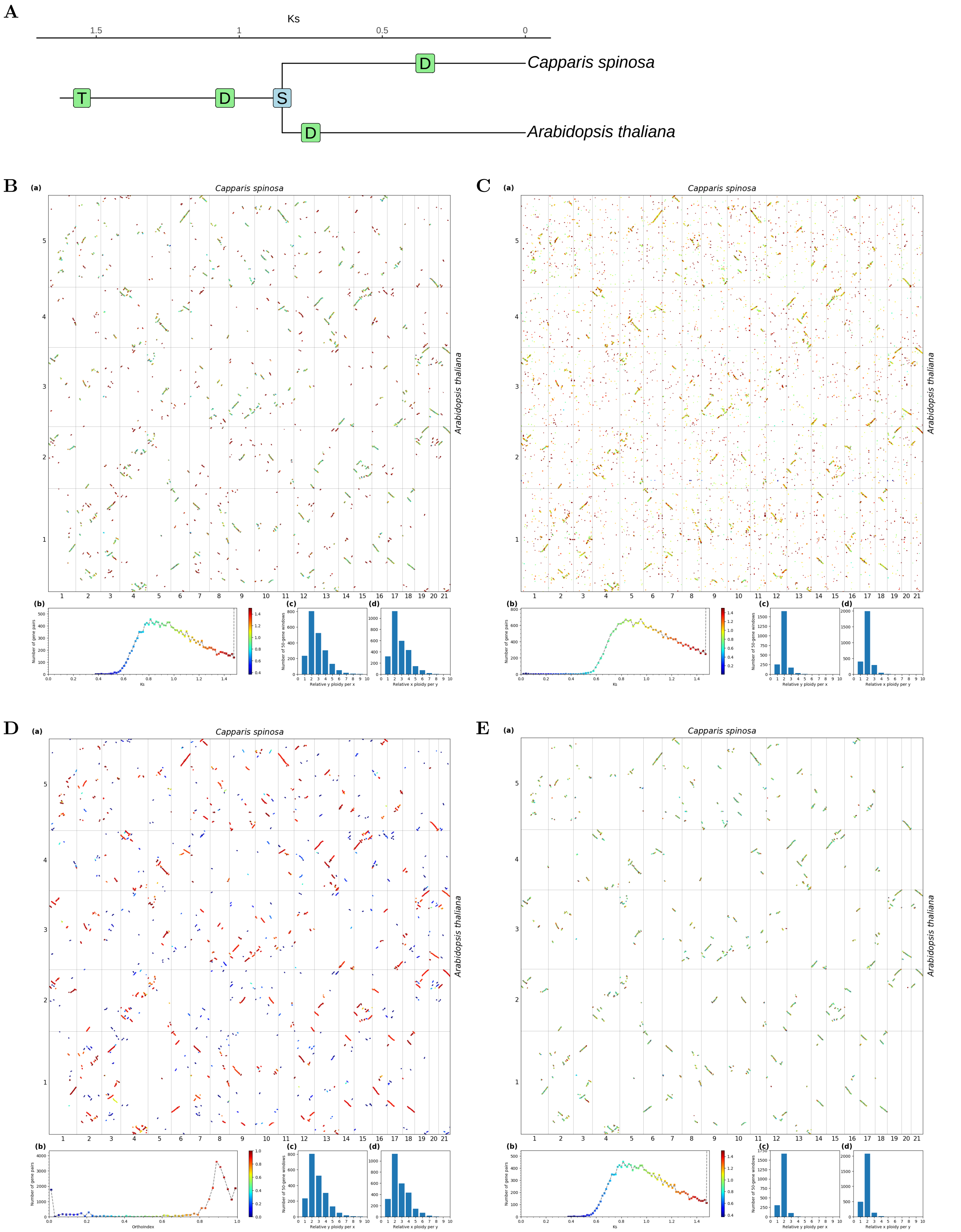


**Fig S80.** ***Orthology Index*** **in the identification of orthologous synteny in** ***Arabidopsis thaliana*** **and** ***Capparis spinosa.*** Refer to **Fig.1** for detailed descriptions.

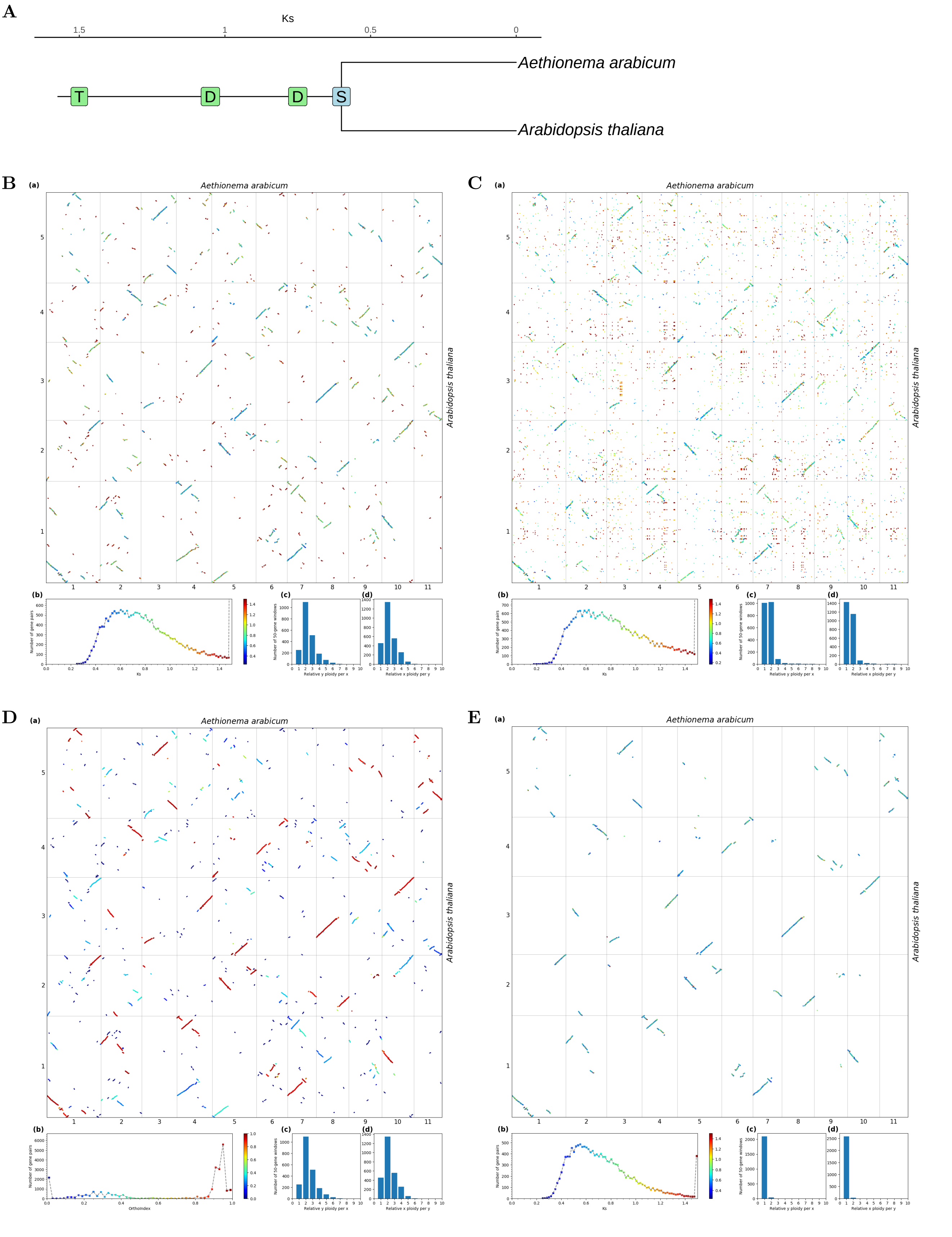


**Fig S81.** ***Orthology Index*** **in the identification of orthologous synteny in** ***Arabidopsis thaliana*** **and** ***Aethionema arabicum.*** Refer to **Fig.1** for detailed descriptions.

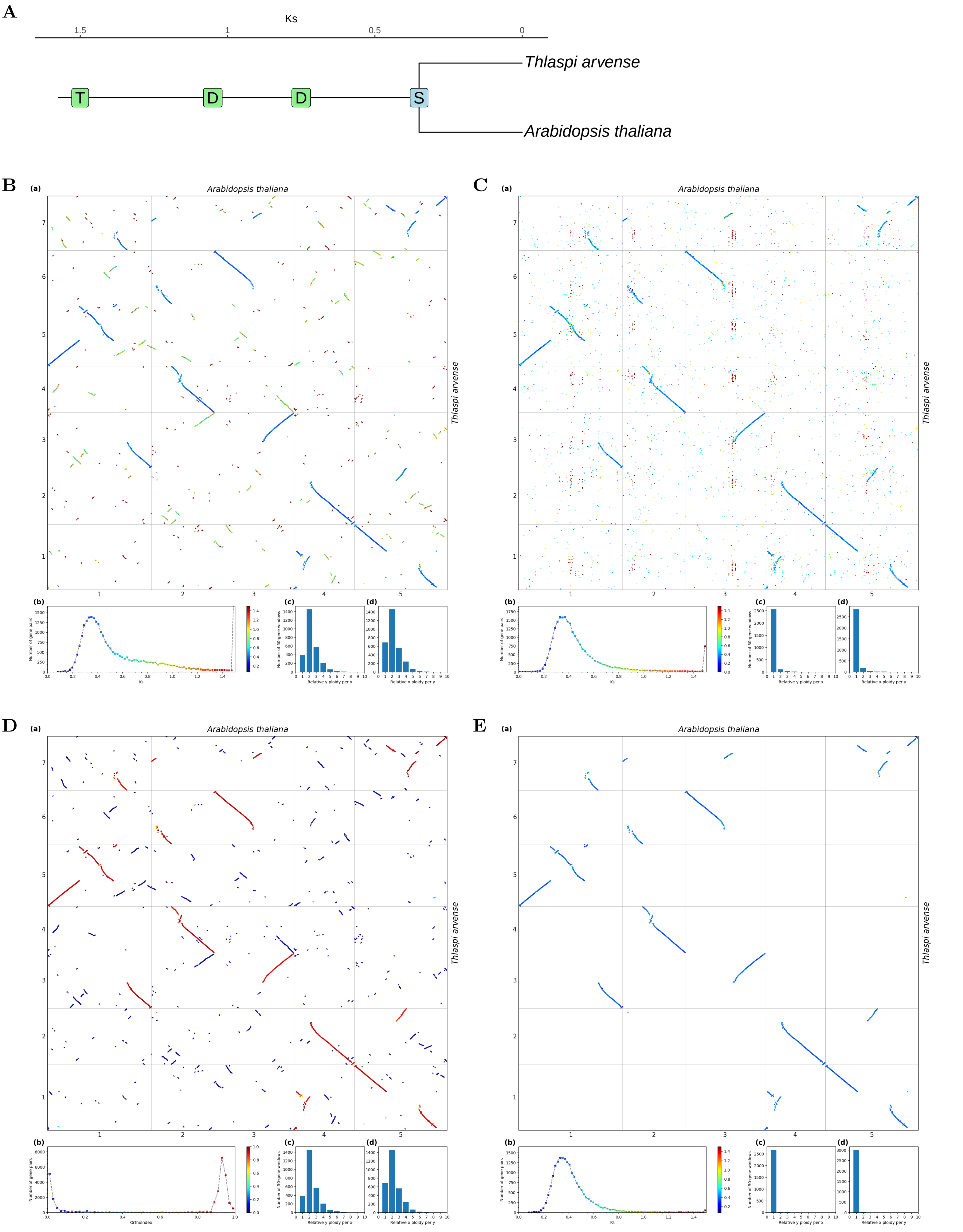


**Fig S82.** ***Orthology Index*** **in the identification of orthologous synteny in** ***Arabidopsis thaliana*** **and** ***Thlaspi arvense.*** Refer to **Fig.1** for detailed descriptions.

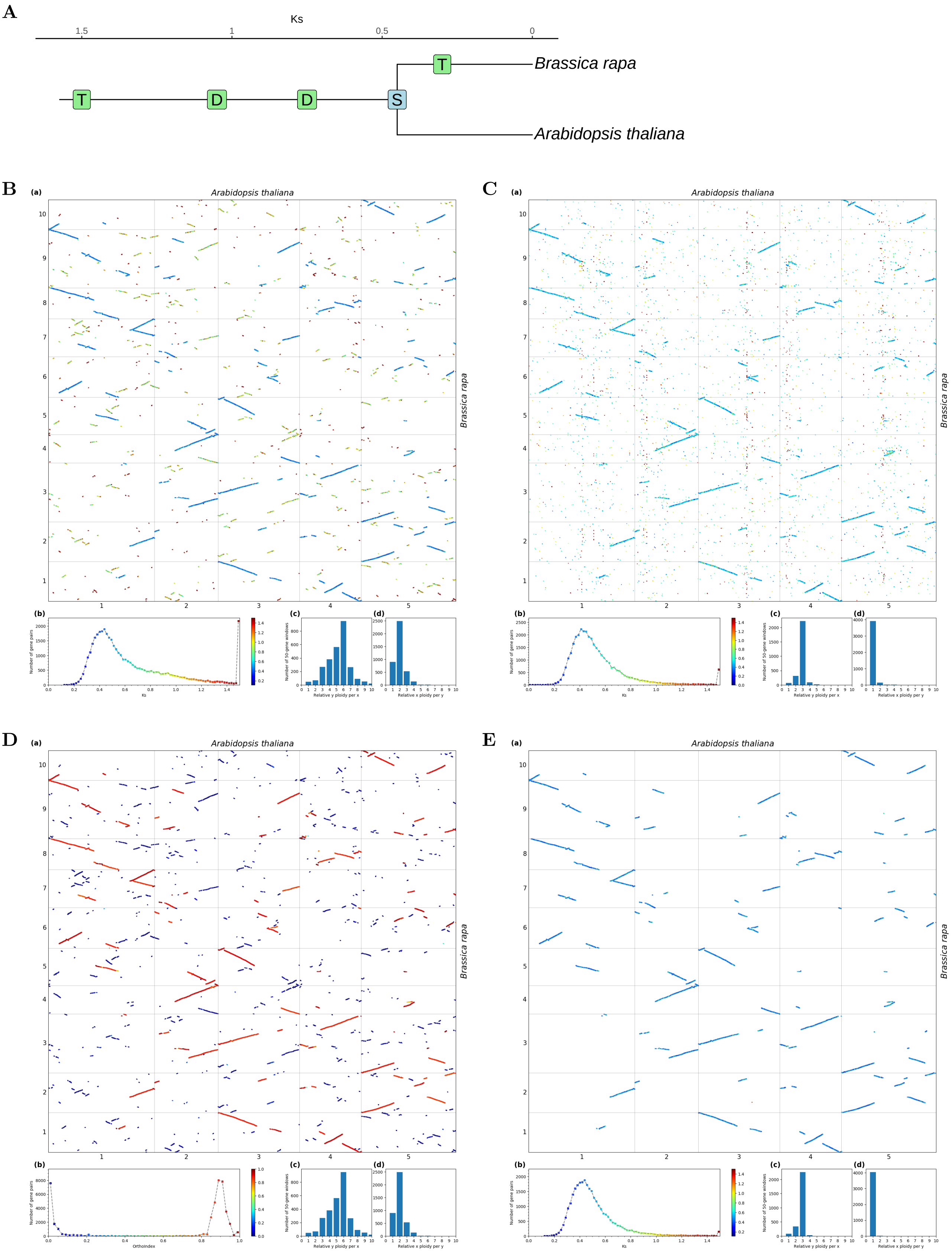


**Fig S83.** ***Orthology Index* in the identification of orthologous synteny in *Arabidopsis thaliana* and *Brassica rapa*.** Refer to **Fig.1** for detailed descriptions.

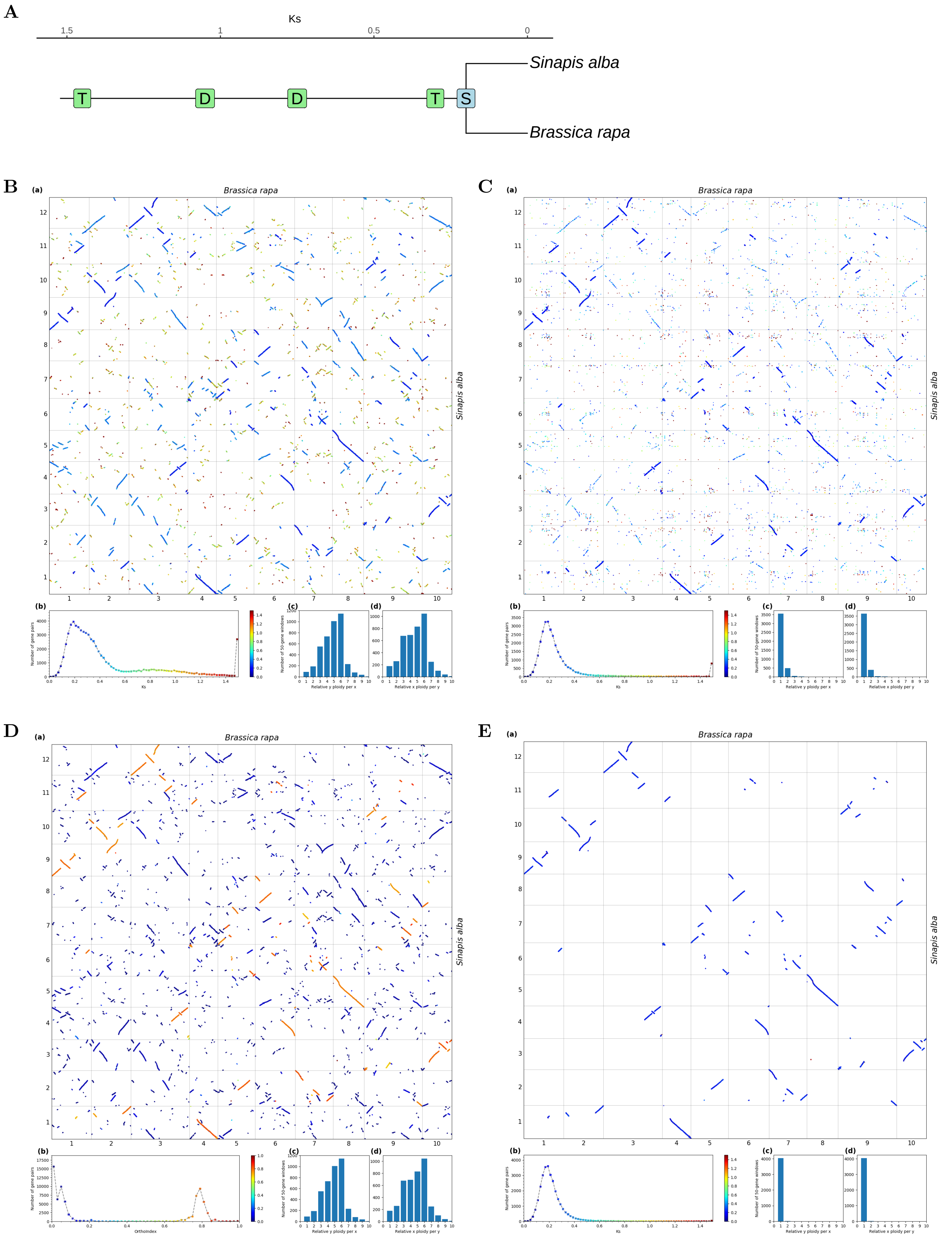


**Fig S84.** ***Orthology Index* in the identification of orthologous synteny in *Brassica rapa* and *Sinapis alba.*** Refer to **Fig.1** for detailed descriptions.

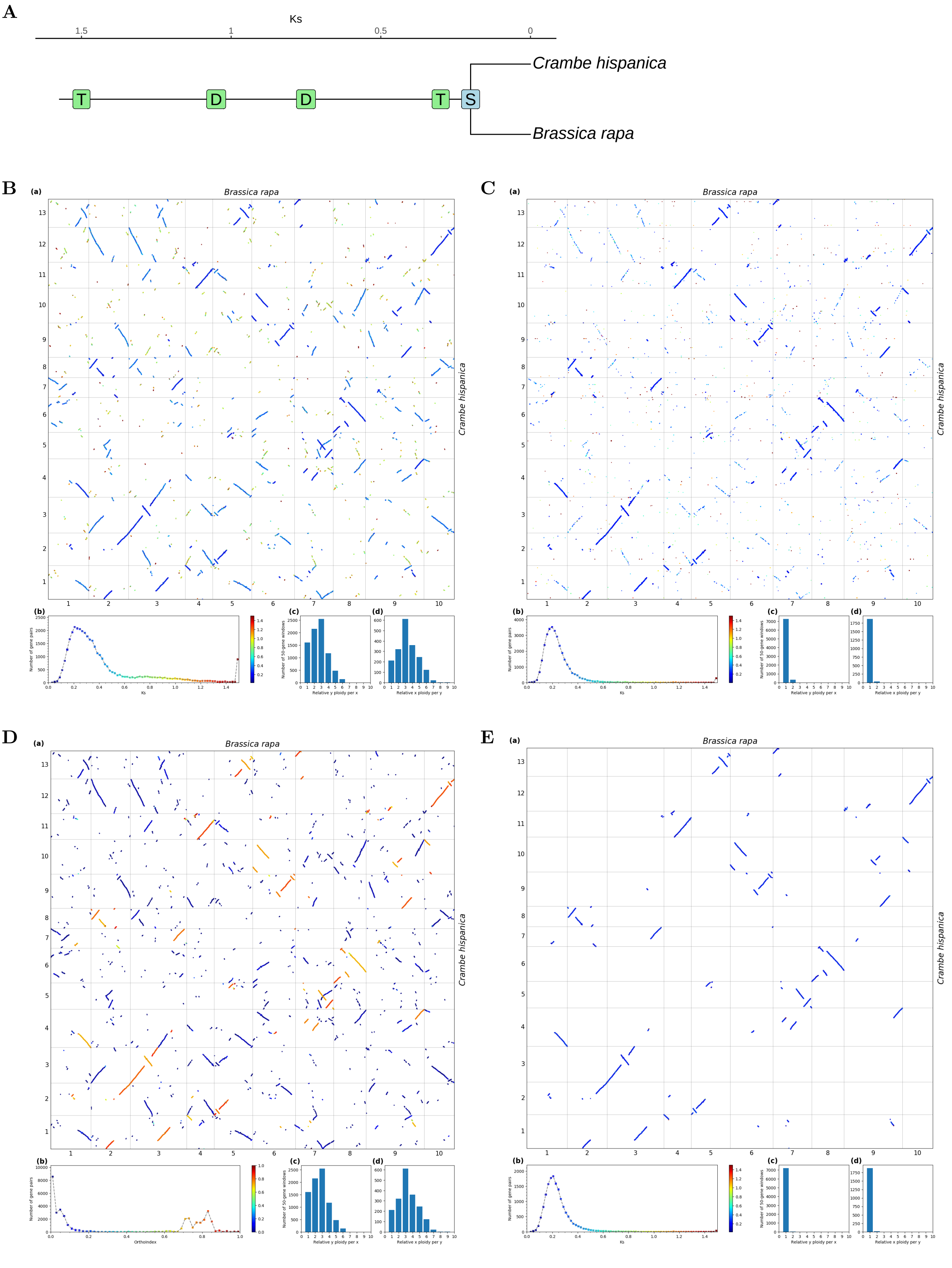


**Fig S85.** ***Orthology Index* in the identification of orthologous synteny in *Brassica rapa* and *Crambe hispanica*.** Refer to **Fig.1** for detailed descriptions.

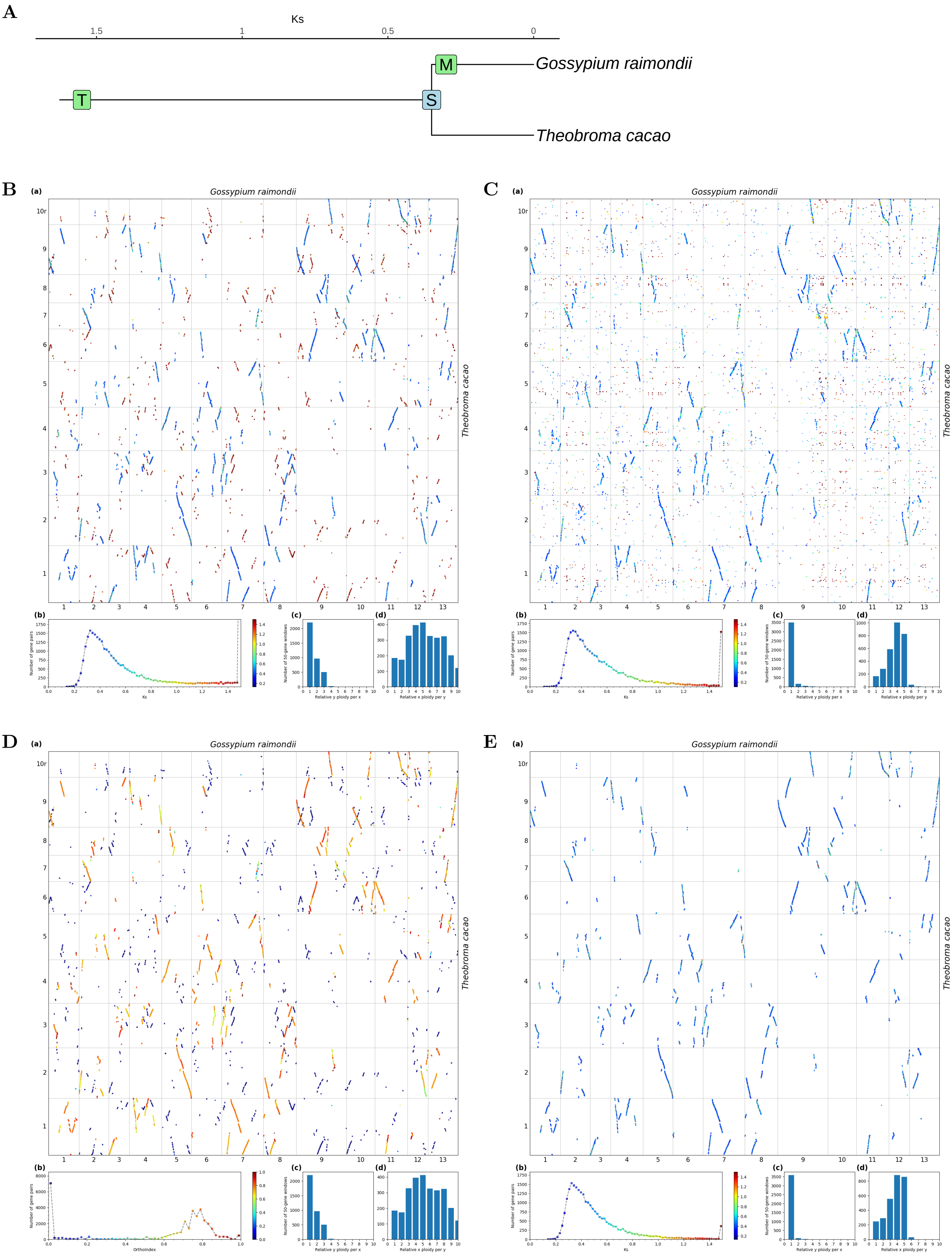


**Fig S86.** ***Orthology Index*** **in the identification of orthologous synteny in** ***Theobroma cacao*** **and** ***Gossypium raimondii.*** Refer to **Fig.1** for detailed descriptions.

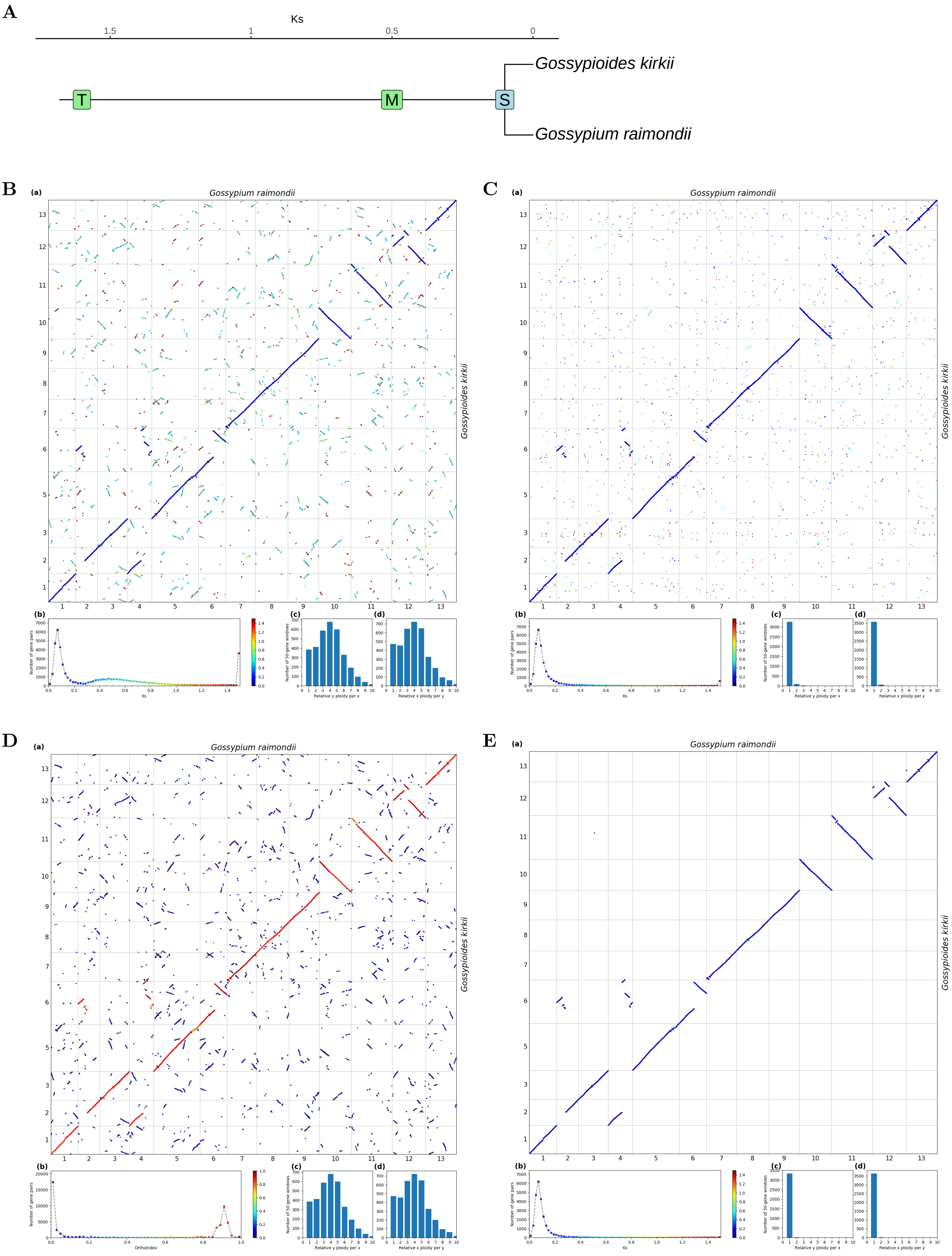


**Fig S87.** ***Orthology Index*** **in the identification of orthologous synteny in** ***Gossypium raimondii*** **and** ***Gossypioides kirkii.*** Refer to **Fig.1** for detailed descriptions.

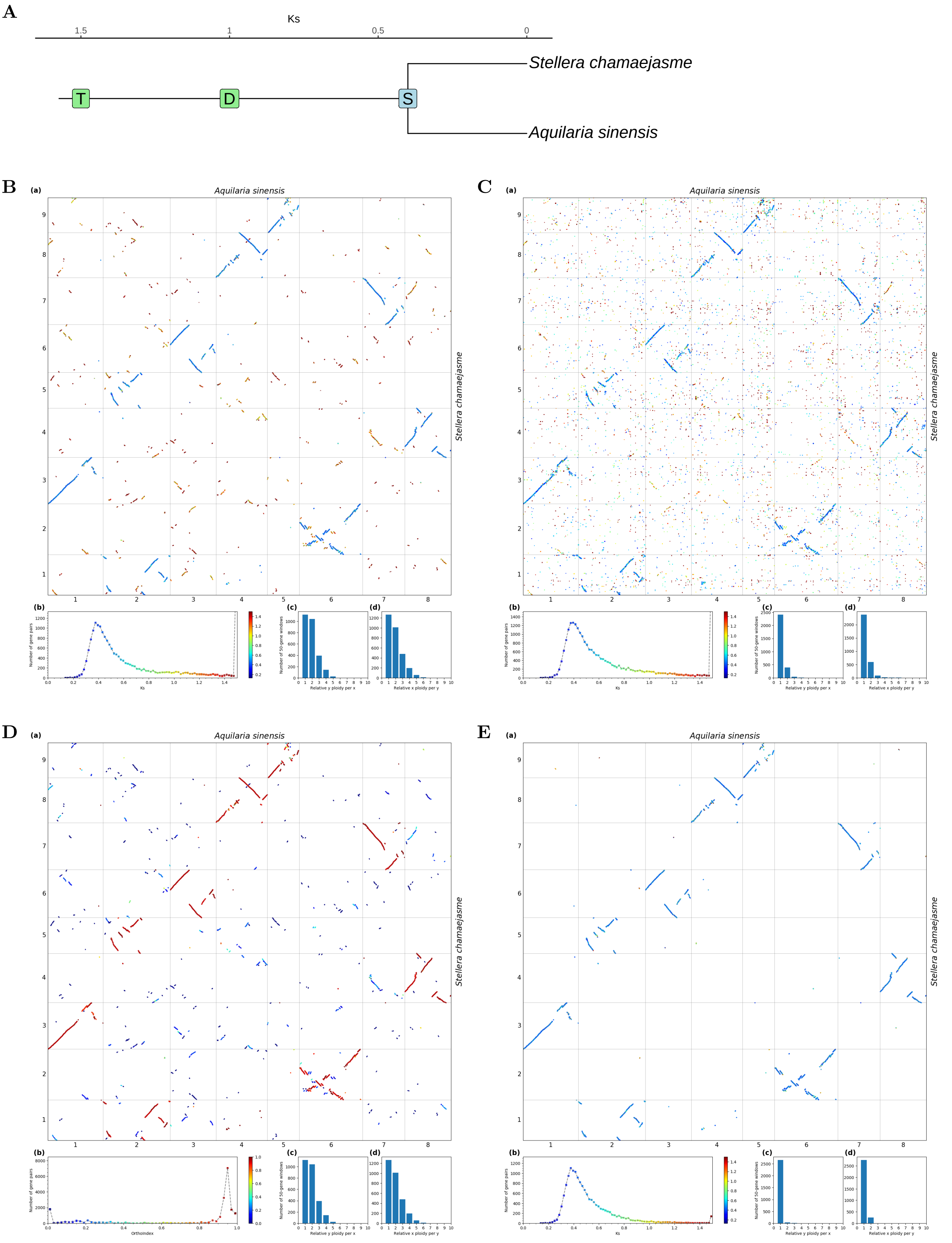


**Fig S88.** ***Orthology Index*** **in the identification of orthologous synteny in** ***Aquilaria sinensis*** **and** ***Stellera chamaejasme.*** Refer to **Fig.1** for detailed descriptions.

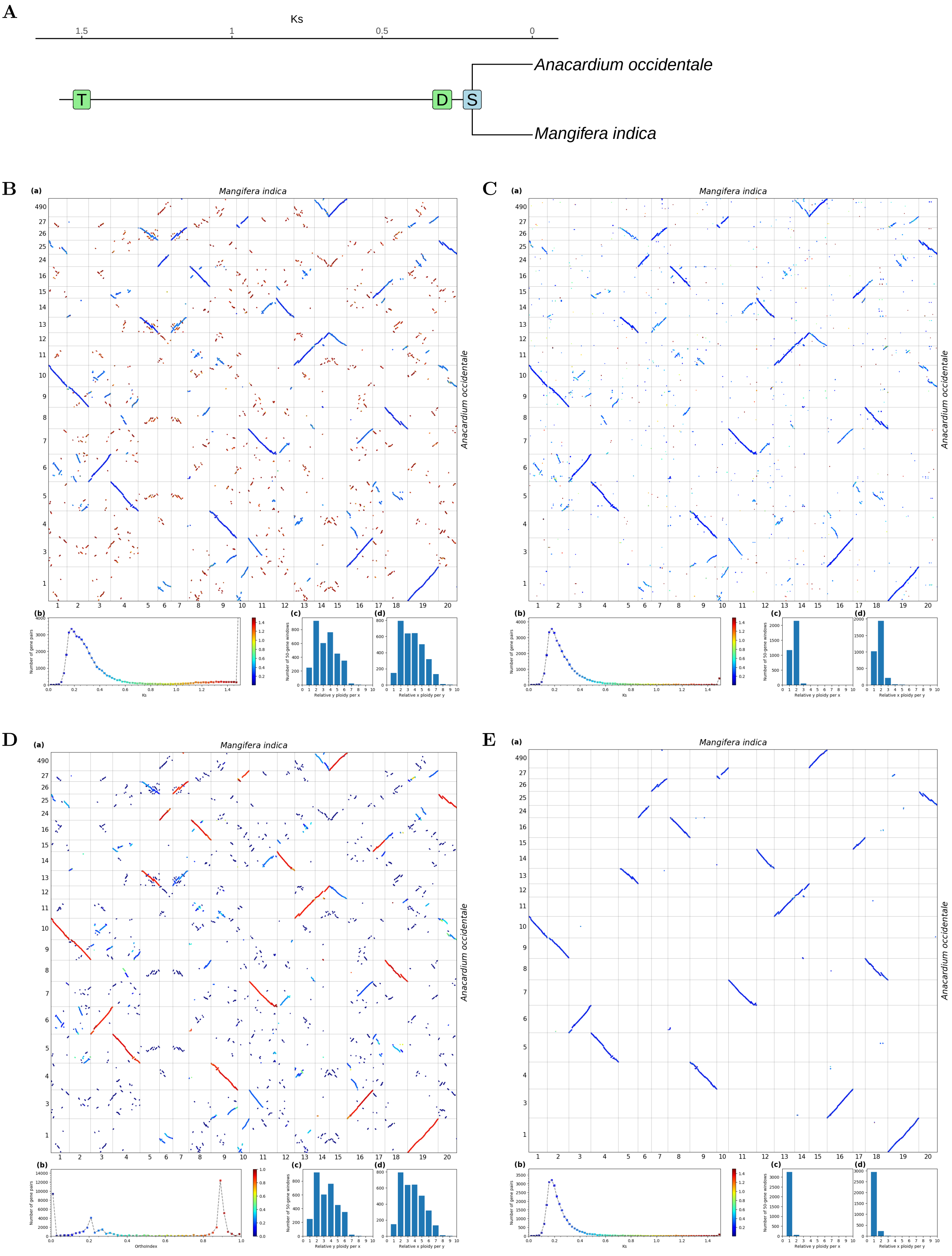


**Fig S89.** ***Orthology Index*** **in the identification of orthologous synteny in** ***Mangifera indica*** **and** ***Anacardium occidentale.*** Refer to **Fig.1** for detailed descriptions.

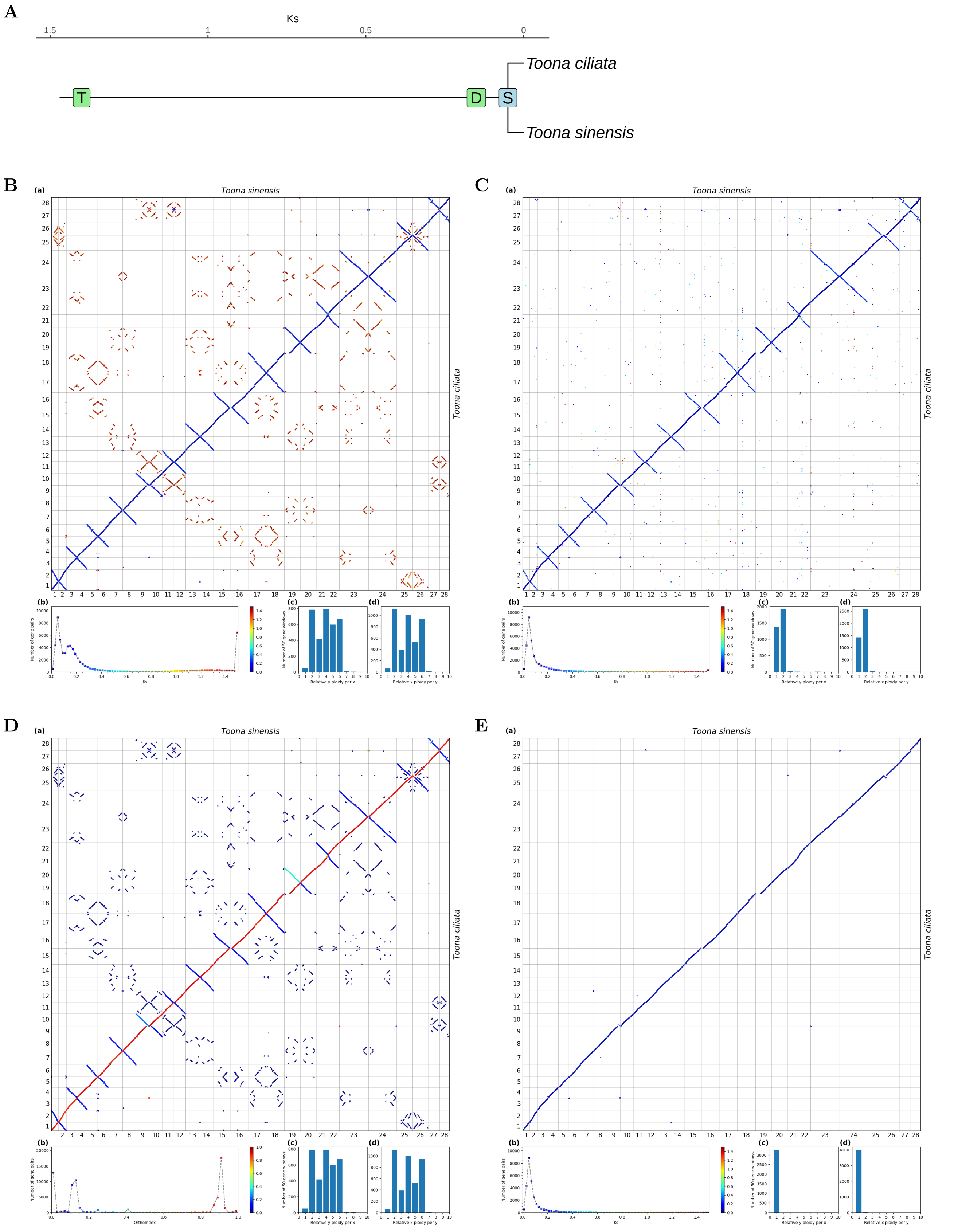


**Fig S90.** ***Orthology Index* in the identification of orthologous synteny in *Toona sinensis* and *Toona ciliata*.** Refer to **Fig.1** for detailed descriptions.

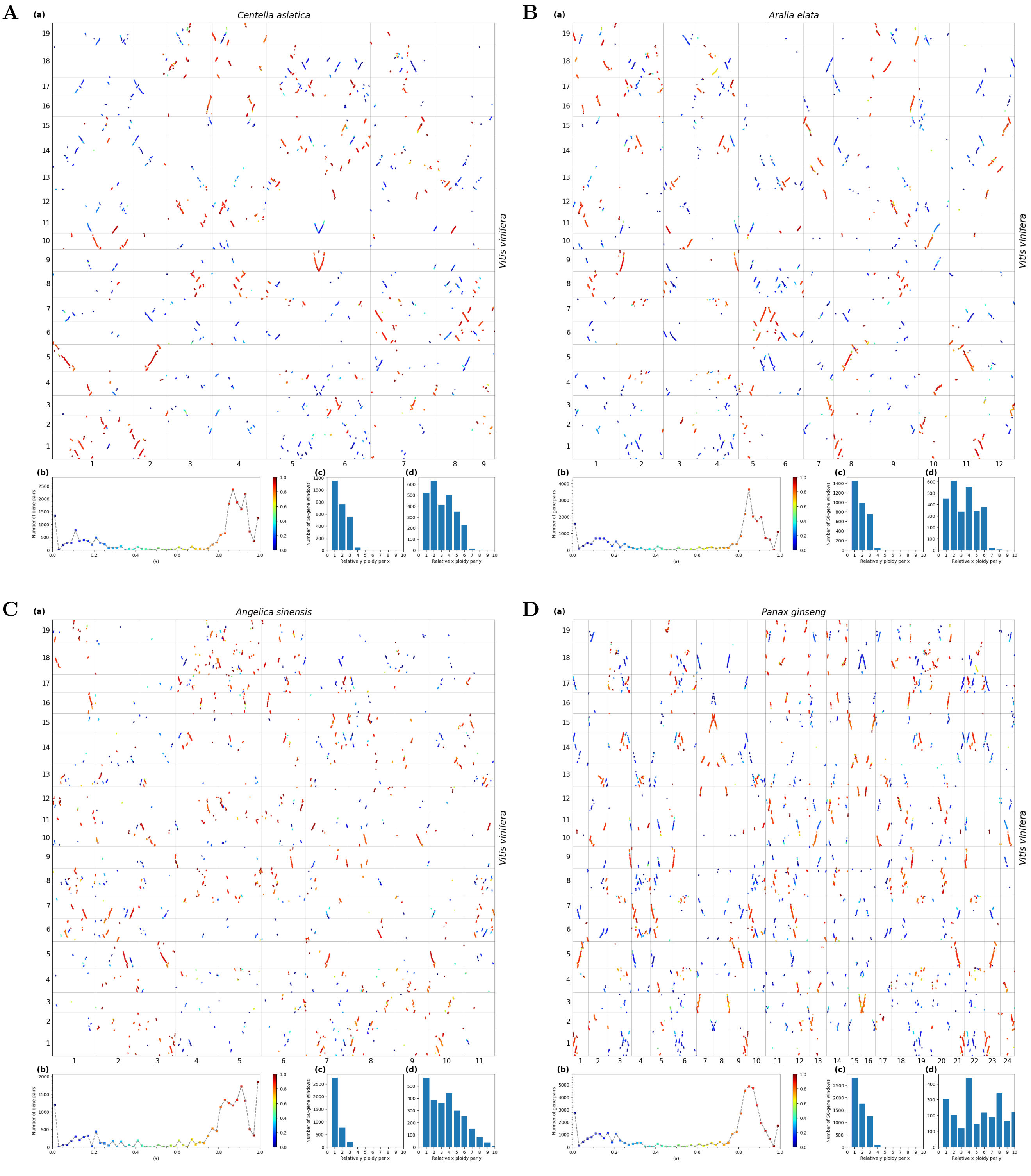


**Fig S91.** ***Orthology Index*-colored dot plots showing orthologous syntenic relationships between *Centella asiatica* : *Vitis vinifera* (2:1 orthologous synteny depth ratio), *Aralia elata* : *Vitis vinifera* (2:1), *Angelica sinensis* : *Vitis vinifera* (4:1), and *Panax ginseng* : *Vitis vinifera* (4:1).**

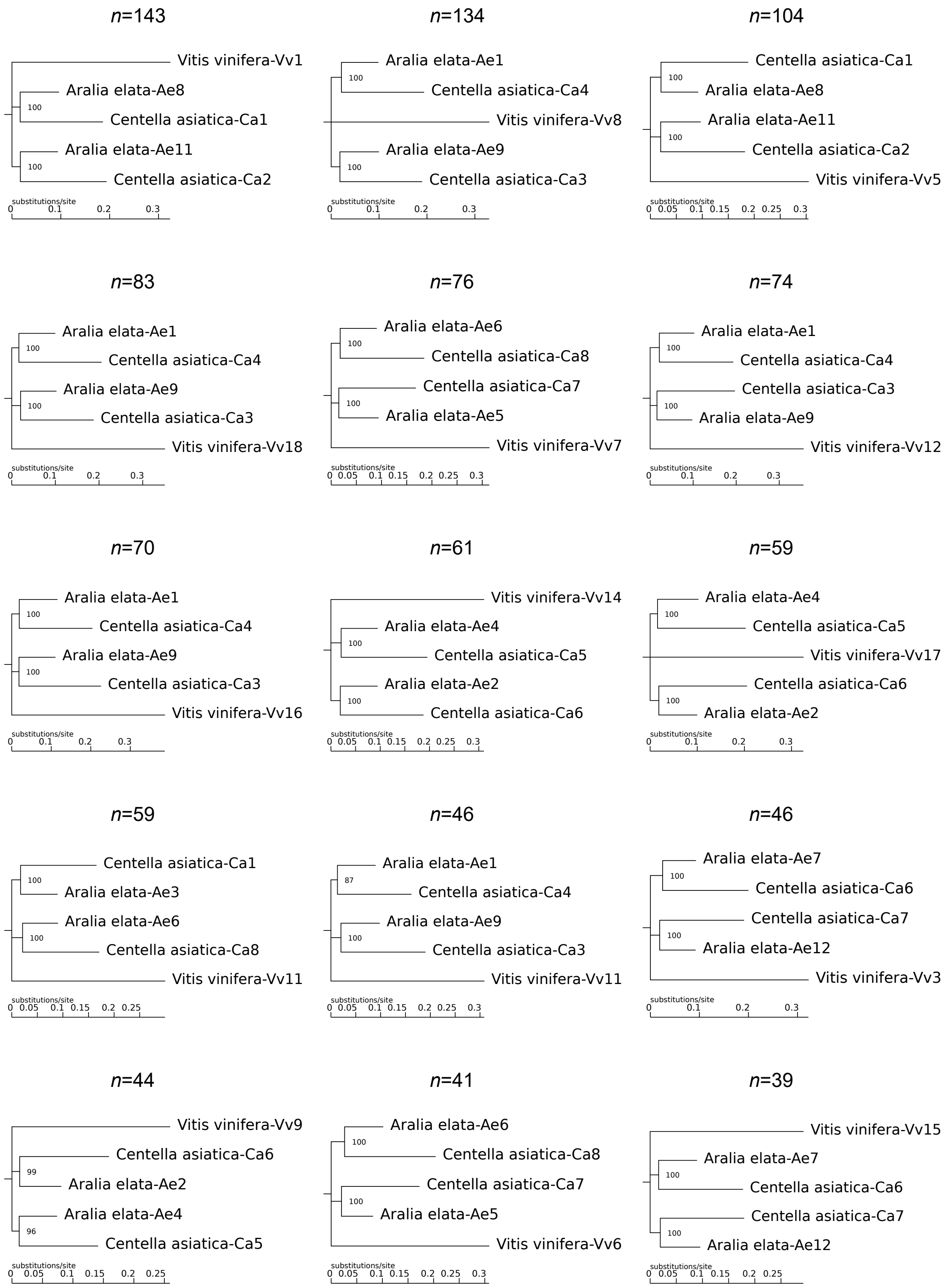


**Fig S92. Macro-synteny phylogenies inferred based on concatenated 1:2:2 (*Vitis vinifera* : *Aralia elata* : *Centella asiatica*) orthologous syntenic genes. The leaf labels on the trees are chromosome numbers. n, number of orthologous syntenic genes. Numbers at nodes represent the bootstrap values (percentage).**

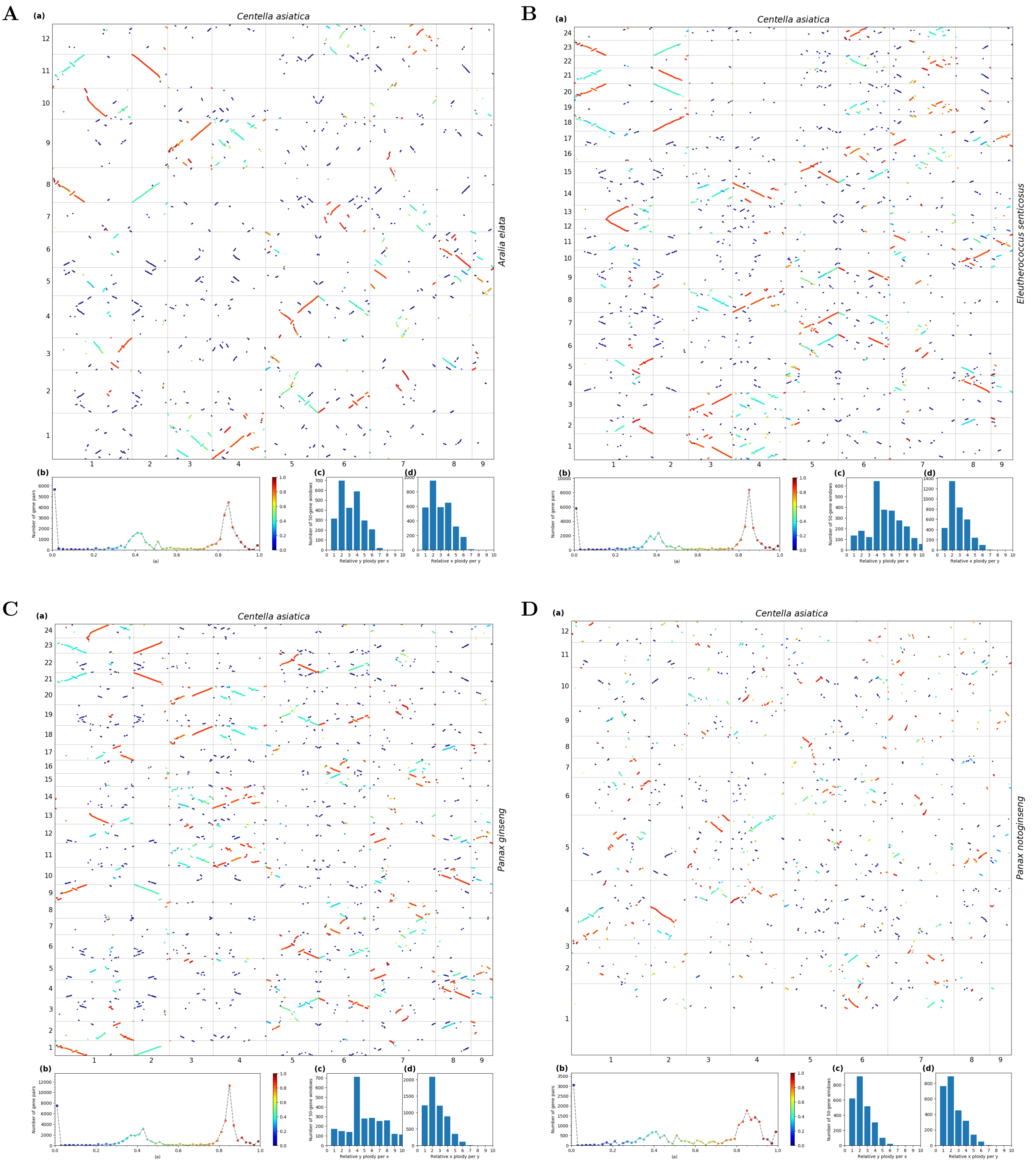


**Fig S93.** ***Orthology Index*-colored dot plots showing orthologous syntenic relationships between *Centella asiatica* and Araliaceae (*Centella asiatica* : *Aralia elata* = 1:1, *Centella asiatica* : *Eleutherococcus senticosus* = 1:2, *Centella asiatica* : *Panax ginseng*=1:2, *Centella asiatica* : *Panax notoginseng*=1:1).**

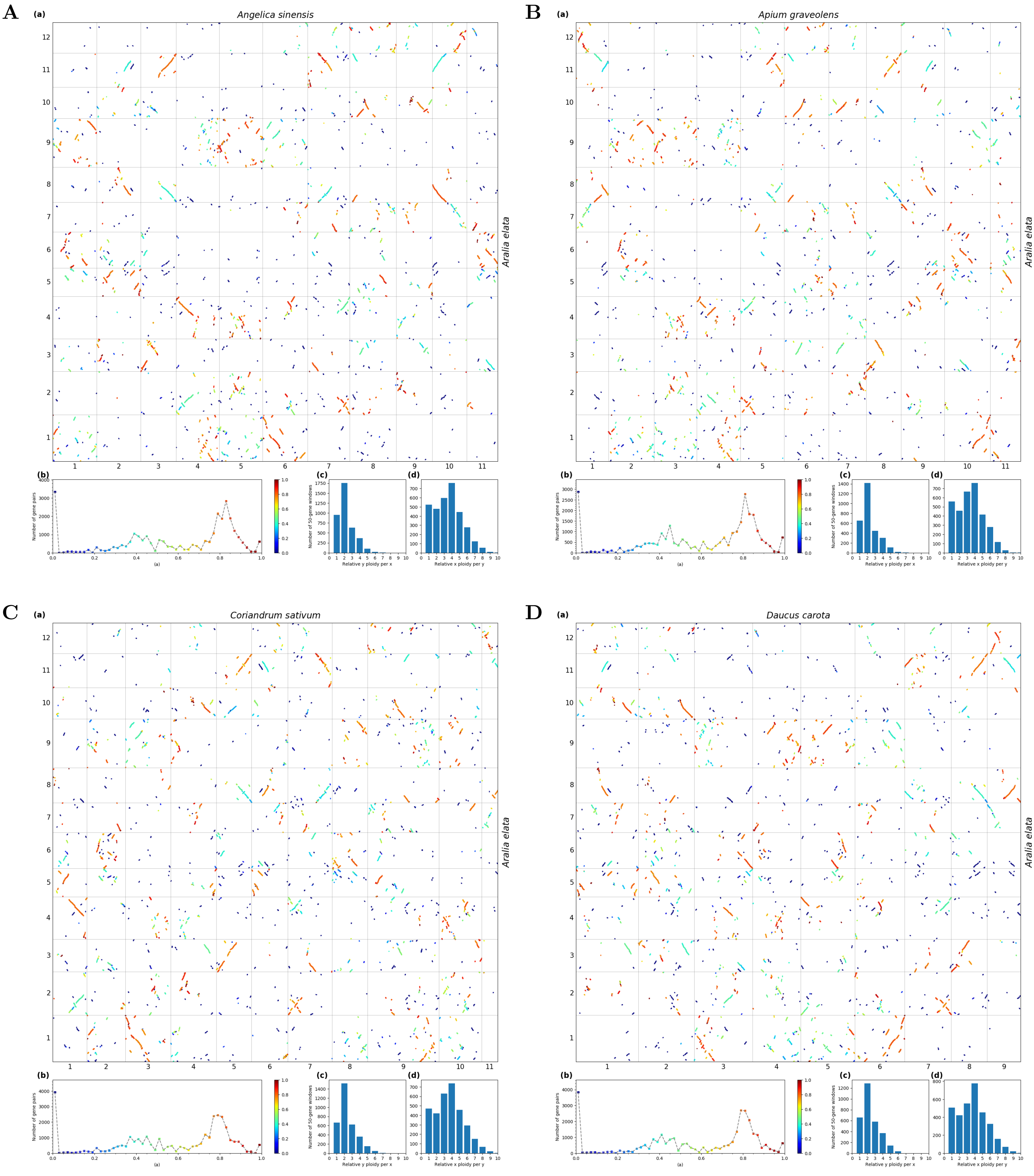


**Fig S94.** ***Orthology Index*-colored dot plots showing orthologous syntenic relationships between *Aralia elata* and Apioideae (all 1:2).**

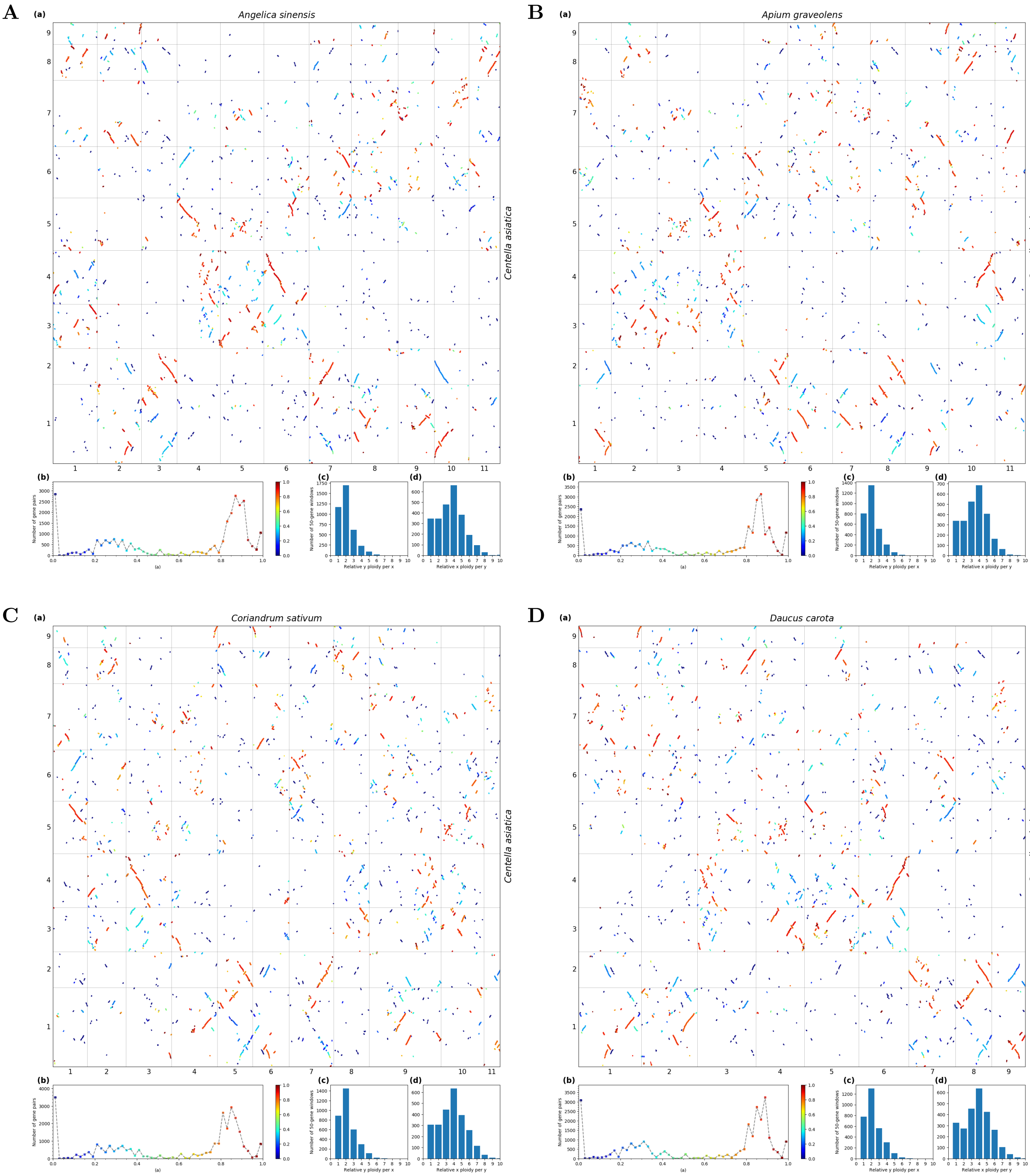


**Fig S95.** *Orthology Index*-colored dot plots showing orthologous syntenic relationships between *Centella asiatica* and Apioideae (all 1:2).

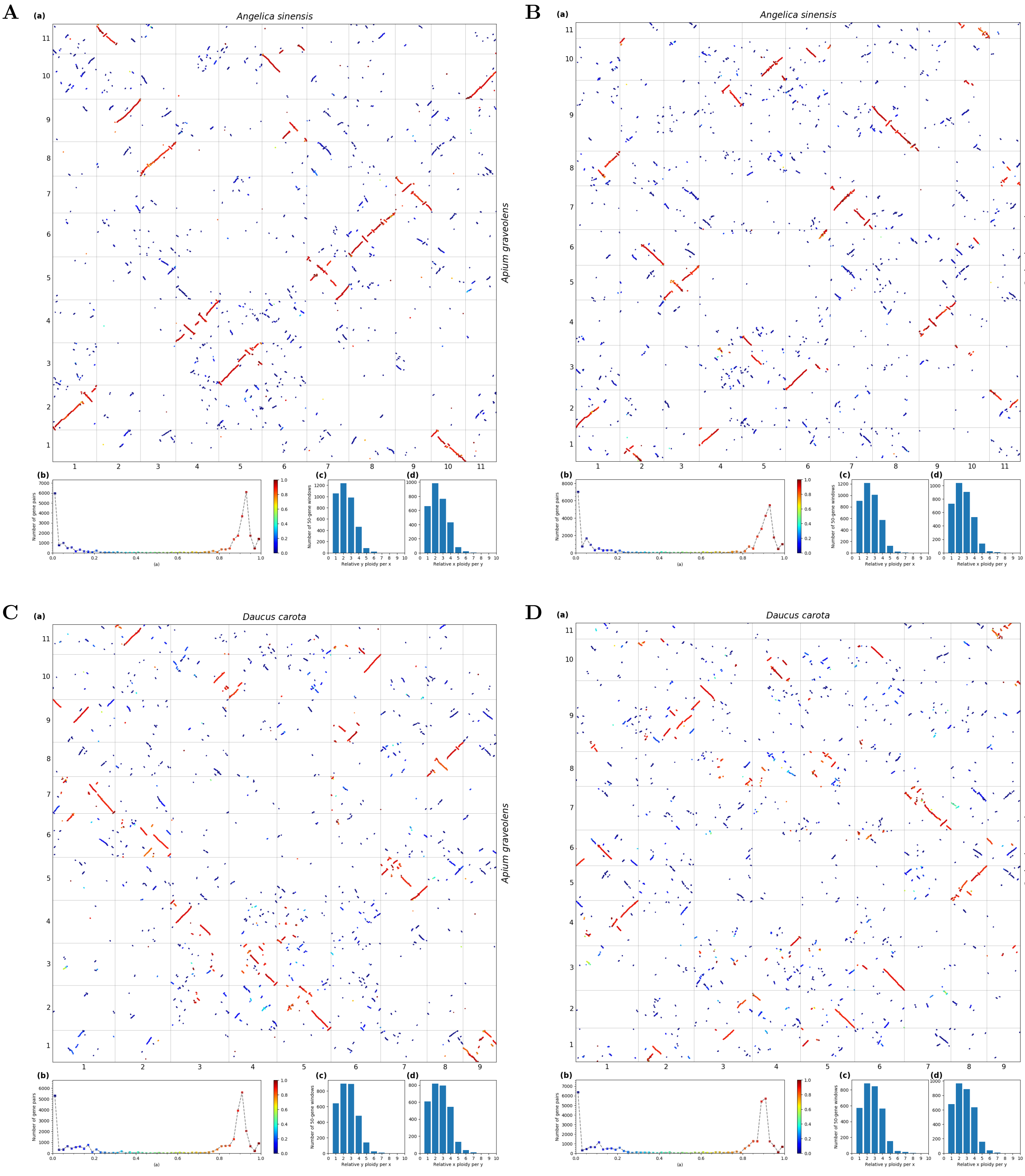


**Fig S96.** ***Orthology Index*-colored dot plots showing orthologous syntenic relationships within Apioideae (all 1:1).**

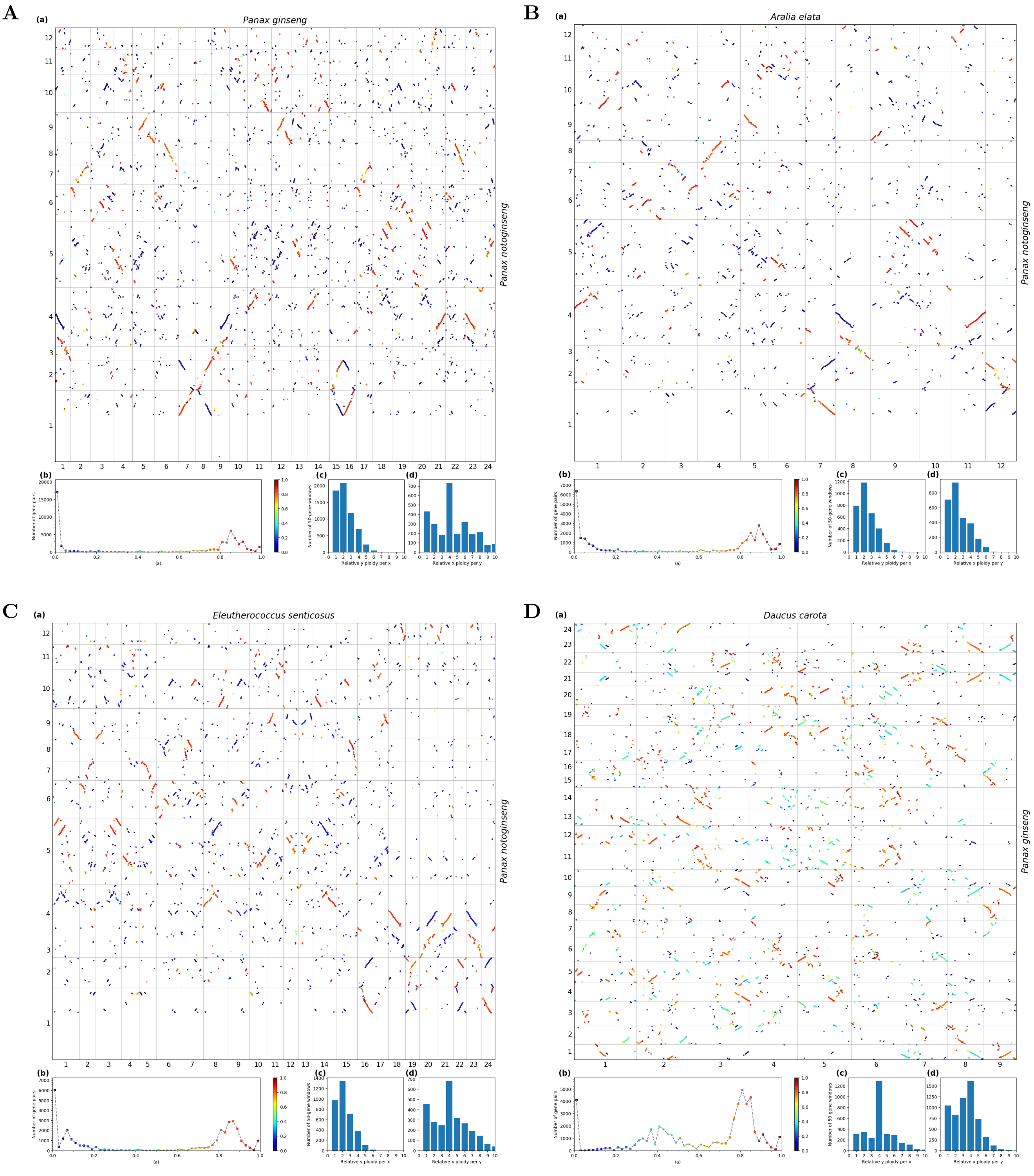


**Fig S97.** ***Orthology Index*-colored dot plots showing orthologous syntenic relationships within Araliaceae (*Panax ginseng* : *Panax notoginseng* = 2:1, *Aralia elata* : *Panax notoginseng* = 1:1, *Eleutherococcus senticosus* : *Panax notoginseng* = 2:1). The other orthologous syntenic ratios (*Aralia elata*: *Eleutherococcus senticosus* = 1:2, *Aralia elata*: *Panax ginseng* = 1:2, *Eleutherococcus senticosus*: *Panax ginseng* = 2:2) can be found in Fig. S57-59.**

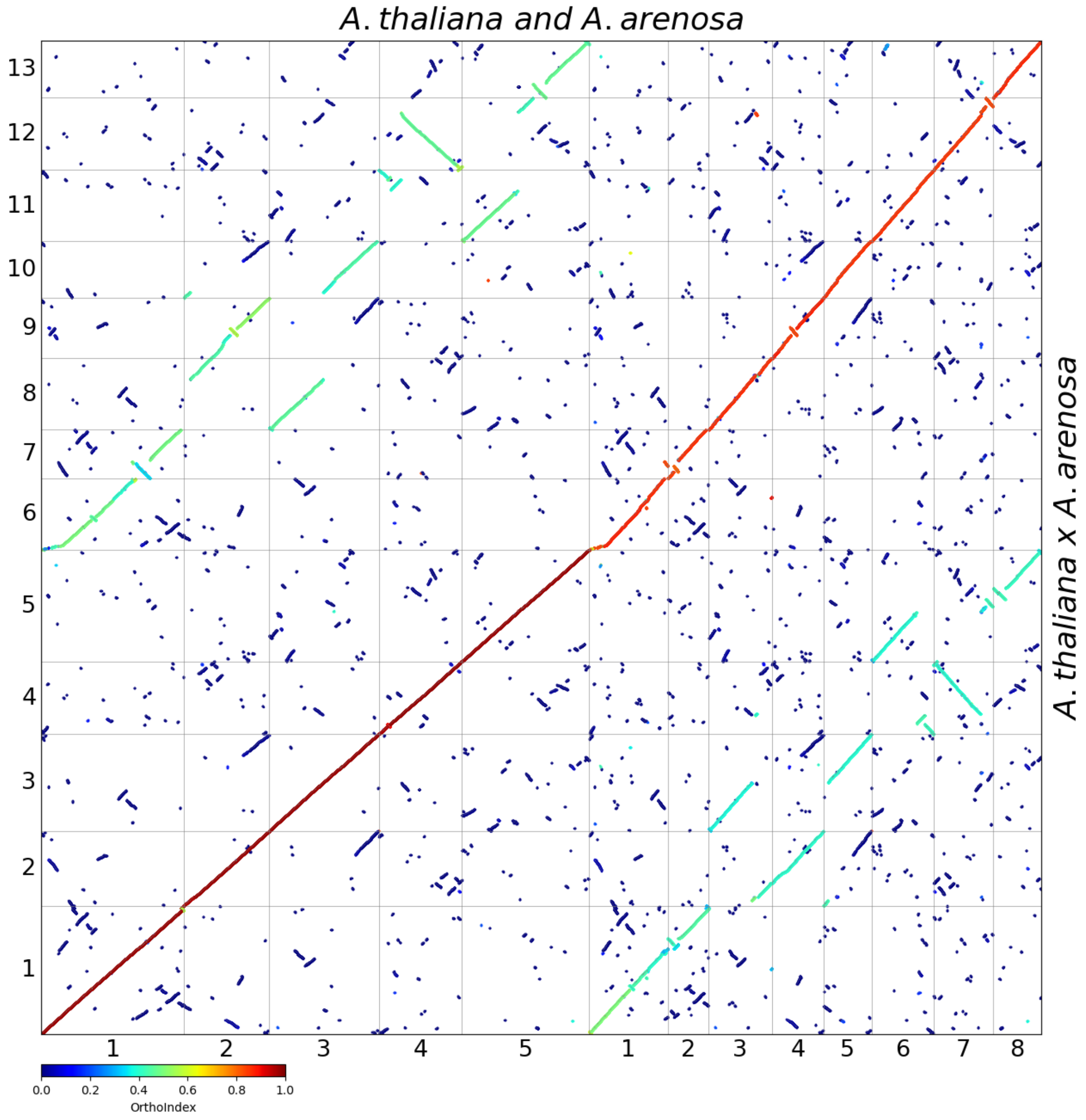


**Fig S98.** ***Orthology Index*-colored dot plots showing orthologous syntenic relationships between *Arabidopsis thaliana* x *A.arenosa* and *A. thaliana* + *A. arenosa*.**

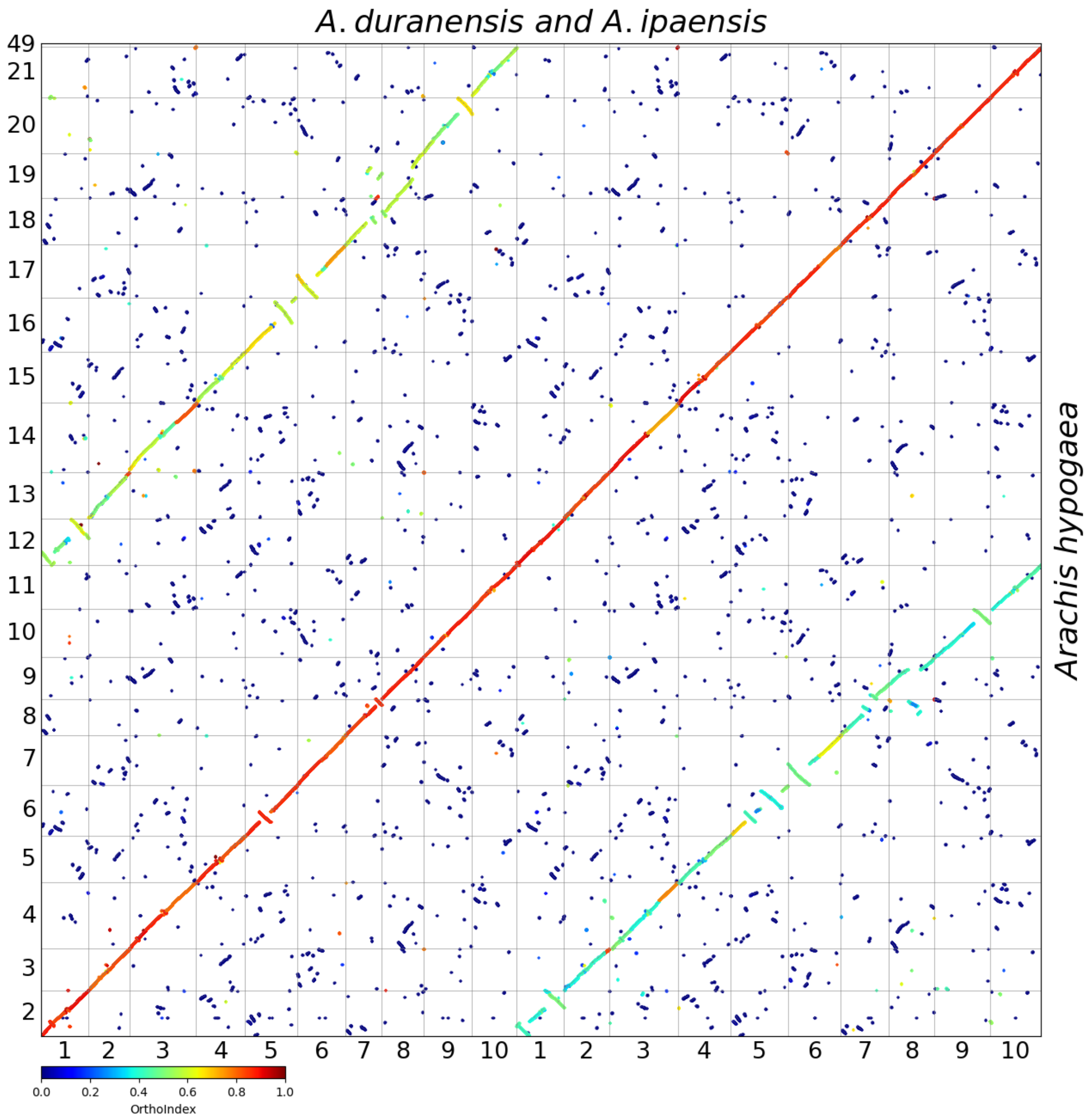


**Fig S99.** ***Orthology Index*-colored dot plots showing orthologous syntenic relationships between *Arachis hypogaea* and *A. duranensis* + *A. ipaensis*.**

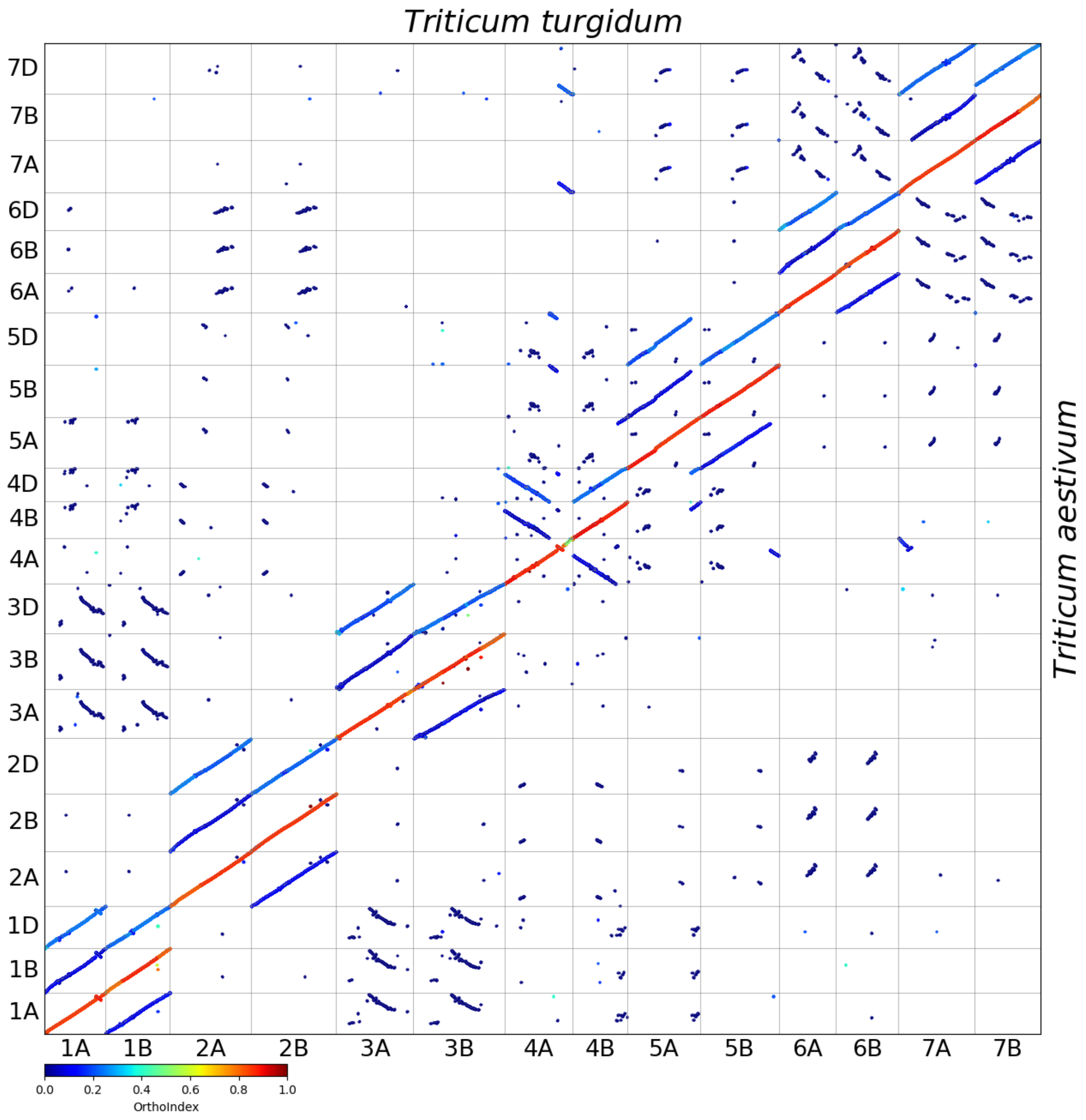


**Fig S100.** ***Orthology Index*-colored dot plots showing orthologous syntenic relationships between *Triticum turgidum* and *T. aestivum.***

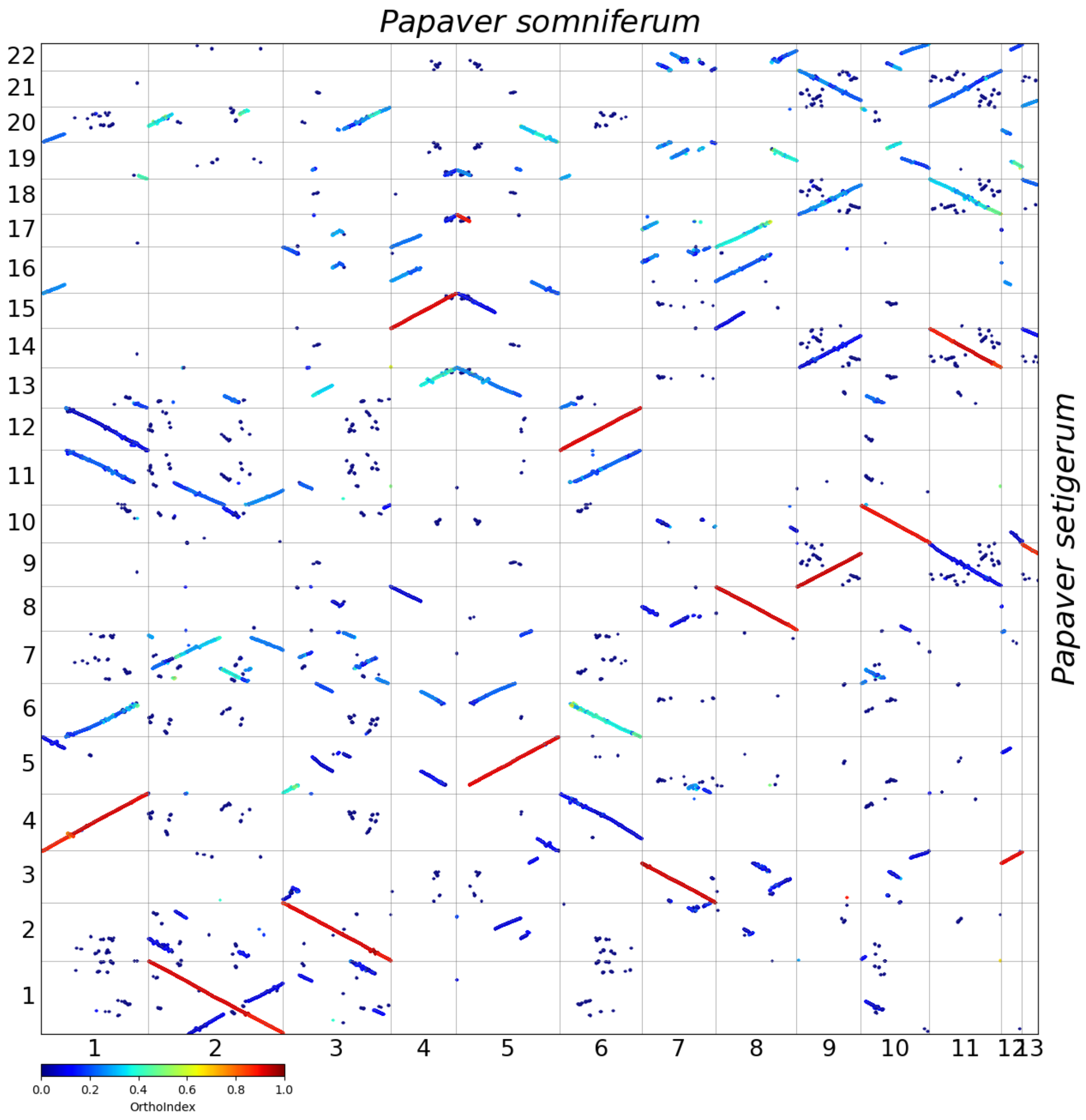


**Fig S101.** ***Orthology Index*-colored dot plots showing orthologous syntenic relationships between *Papaver somniferum* and *P. setigerum*.**

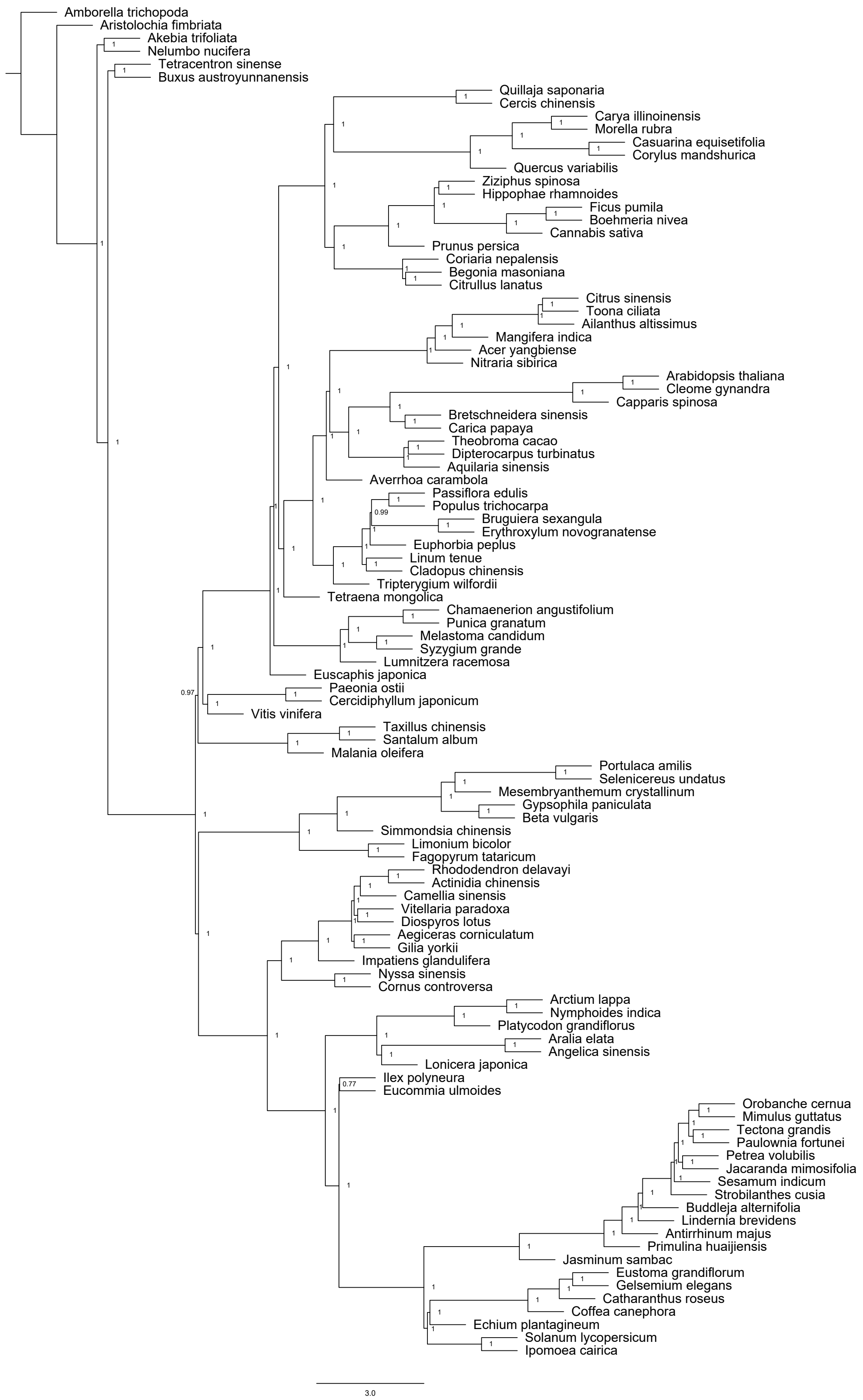


**Fig S102. Phylogenetic relationships within the core eudicots based on 12,277 multi-copy SOGs.** The numbers at the nodes are posterior probabilities from ASTRAL. Bar, 3.0 coalescent units.

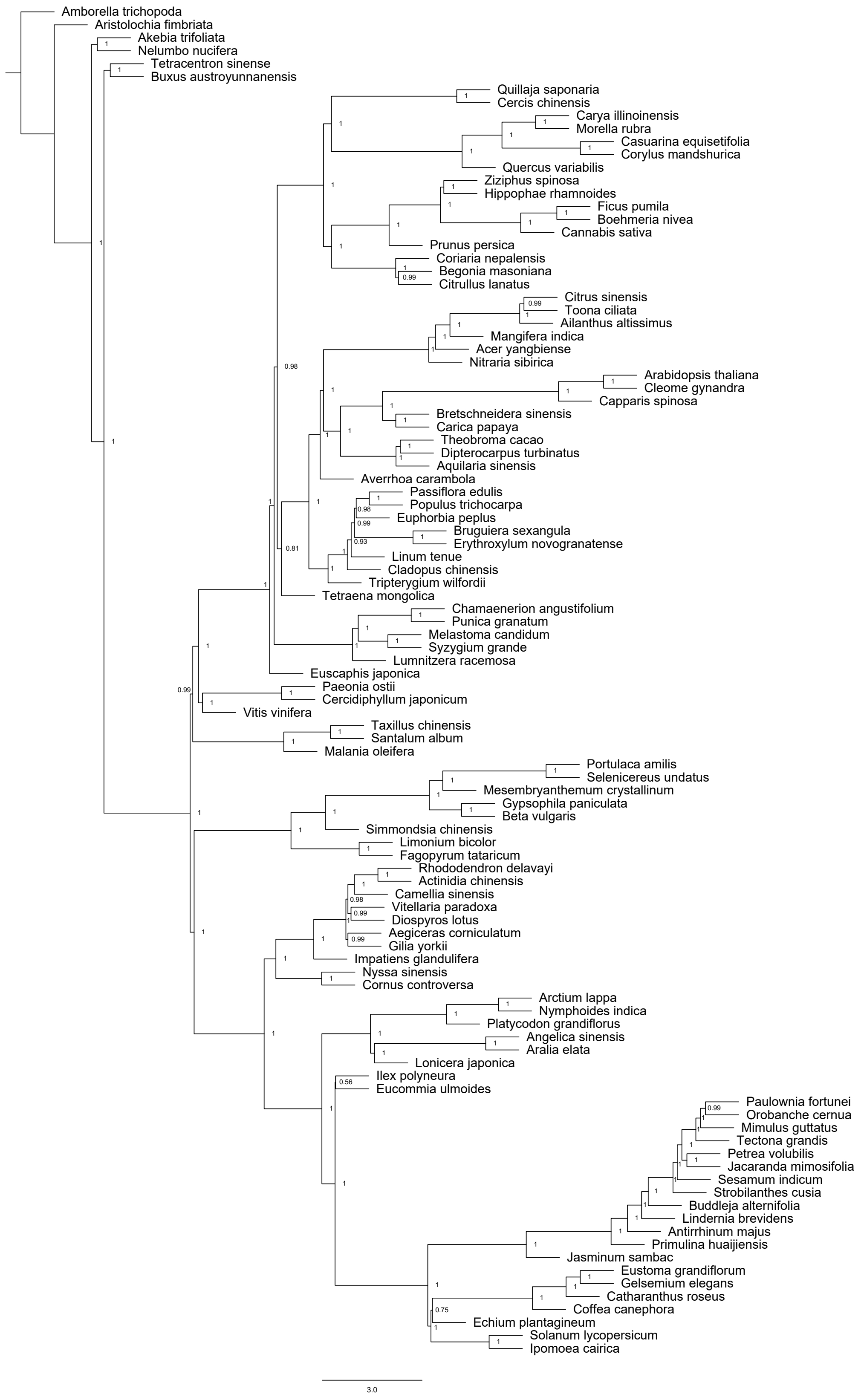


**Fig S103. Phylogenetic relationships within the core eudicots based on 5,154 single-copy SOGs.** The numbers at the nodes are posterior probabilities from ASTRAL. Bar, 3.0 coalescent units.

Table S1. Cases with shared polyploidization event(s) used in this study.

| Orthology | Lineages | Out-paralogy | In-paralogy | References |
|---|---|---|---|---|
| *Liriodendron_chinense-Cinnamomum_kanehirae* | Magnoliidae | 1 WGD | 1 WGD in Cka | [1] |
| *Liriodendron_chinense-Magnolia_biondii* | Magnoliaceae | 1 WGD | None | [2] |
| *Cinnamomum_kanehirae-Chimonanthus_salicifolius* | Laurales | 1 WGD | 1 WGD in Cka<br>1 WGD in Csa | [3] |
| *Cinnamomum_kanehirae-Phoebe_bournei* | Lauraceae | 2 WGDs | None | [4] |
| *Protea_cynaroides-Telopea_speciosissima* | Proteaceae | 1 WGD | None | [5] |
| *Protea_cynaroides-Macadamia_integrifolia* | Proteaceae | 1 WGD | None | [5] |
| *Aquilegia_coerulea-Coptis_chinensis* | Ranunculaceae | 1 WGD | None | [6] |
| *Aquilegia_coerulea-Papaver_somniferum* | Ranunculales | 1 WGD | 1 WGD in Pso | [6] |
| *Corydalis_tomentella-Papaver_somniferum* | Papaveraceae | 1 WGD | 1 WGD in Pso | [7] |
| *Tetracendron_sinense-Trochodendron_aralioides* | Trochodendraceae | 2 WGDs | None | [8] |
| *Vitis_vinifera-Theobroma_cacao* | Pentapetalae | 1 WGT | None | [9] |
| *Vitis_vinifera-Populus_trichocarpa* | Pentapetalae | 1 WGT | 1 WGD in Ptr | [10] |
| *Vitis_vinifera-Arabidopsis_thaliana* | Pentapetalae | 1 WGT | 2 WGDs in Ath | [10] |
| *Vitis_vinifera-Solanum_lycopersicum* | Pentapetalae | 1 WGT | 1 WGT in Sly | [11] |
| *Vitis_vinifera-Malania_oleifera* | Pentapetalae | 1 WGT | None | [12] |
| *Vitis_vinifera-Spinacia_oleracea* | Pentapetalae | 1 WGT | None | [13] |
| *Vitis_vinifera-Camellia_sinensis* | Pentapetalae | 1 WGT | 1 WGD in Csi | [14] |
| *Vitis_vinifera-Coffea_canephora* | Pentapetalae | 1 WGT | None | [15] |
| *Vitis_vinifera-Eucommia_ulmoides* | Pentapetalae | 1 WGT | 1 WGD in Eul | [16] |
| *Vitis_vinifera-Daucus_carota* | Pentapetalae | 1 WGT | 2 WGDs in Dca | [17] |
| *Vitis_vinifera-Cercidiphyllum_japonicum* | Pentapetalae | 1 WGT | None | [18] |
| *Vitis_vinifera-Paeonia_ostii* | Pentapetalae | 1 WGT | None | [19] |
| *Vitis_vinifera-Prunus_persica* | Pentapetalae | 1 WGT | None | [20] |
| *Vitis_vinifera-Tripterygium_wilfordii* | Pentapetalae | 1 WGT | 1 WGT in Twi | [21] |
| *Vitis_vinifera-Betula_pendula* | Pentapetalae | 1 WGT | None | [22] |
| *Vitis_vinifera-Euscaphis_japonica* | Pentapetalae | 1 WGT | 1 WGD in Eja | [23] |
| *Vitis_vinifera-Carica_papaya* | Pentapetalae | 1 WGT | None | [24] |
| *Vitis_vinifera-Acer_yangbiense* | Pentapetalae | 1 WGT | None | [25] |
| *Populus_trichocarpa-Arabidopsis_thaliana* | Rosidae | 1 WGT | 1 WGD in Ptr<br>2 WGDs in Ath | [10] |
| *Amaranthus_tricolor-Amaranthus_hypochondriacus* | Amaranthaceae | 1 WGT, 1 WGD | None | [26] |
| *Fagopyrum_tataricum-Fagopyrum_esculentum* | Polygonaceae | 1 WGT, 2 WGDs | None | [27] |
| *Santalum_album-Santalum_yasi* | Santalaceae | 2 WGTs | None | [28] |
| *Rhododendron_simsii-Vaccinium_corymb* | Ericaceae | 1 WGT, 1 WGD | None | [29] |

| | | | | |
|---|---|---|---|---|
| *osum* | | | | |
| *Rhododendron_simsii-Actinidia_chinensis* | Ericaceae | 1 WGT, 1 WGD | 1 WGD in Ach | [29] |
| *Rhododendron_simsii-Camptotheca_acuminata* | Ericales-Cornales | 1 WGT | 1 WGD in Rsi<br>1 WGD in Cac | [29] |
| *Davidia_involucrata-Camptotheca_acuminata* | Nyssaceae | 1 WGT, 1 WGD | None | [30] |
| *Paulownia_fortunei-Sesamum_indicum* | Lamiales | 1 WGT, 1 WGD | None | [31] |
| *Orobanche_cernua-Mimulus_guttatus* | Lamiales | 1 WGT, 1 WGD | None | [32] |
| *Tectona_grandis-Jacaranda_mimosifolia* | Lamiales | 1 WGT, 1 WGD | None | [33] |
| *Tectona_grandis-Salvia_splendens* | Lamiaceae | 1 WGT, 1 WGD | 2 WGDs in Ssp | [33] |
| *Salvia_splendens-Salvia_hispanica* | Lamiaceae | 1 WGT, 2 WGDs | 1 WGD in Ssp | [34] |
| *Jasminum_sambac-Osmanthus_fragrans* | Oleaceae | 2 WGTs | 1 WGD in Oeu | [35] |
| *Syringa_oblata-Osmanthus_fragrans* | Oleaceae | 2 WGTs, 1 WGD | None | [36] |
| *Coffea_canephora-Marsdenia_tenacissima* | Gentianales | 1 WGT | None | [37] |
| *Coffea_canephora-Ophiorrhiza_pumila* | Rubiaceae | 1 WGT | None | [38] |
| *Ophiorrhiza_pumila-Neolamarckia_cadamba* | Rubiaceae | 1 WGT | 1 WGD in Nca | [39] |
| *Solanum_lycopersicum-Anisodus_acutangulus* | Solanaceae | 2 WGTs | 1 WGD in Aac | [40] |
| *Solanum_lycopersicum-Nicotiana_tabacum* | Solanaceae | 2 WGTs | 1 WGD in Nta | [41] |
| *Solanum_lycopersicum-Lycium_barbarum* | Solanaceae | 2 WGTs | None | [42] |
| *Ipomoea_cairica-Cuscuta_europaea* | Convolvulaceae | 2 WGTs | None | [43] |
| *Arctium_lappa-Taraxacum_mongolicum* | Asteraceae | 2 WGTs | None | [44] |
| *Mikania_micrantha-Helianthus_annuus* | Asteraceae | 2 WGTs, 1 WGD | None | [45] |
| *Mikania_micrantha-Stevia_rebaudiana* | Asteraceae | 2 WGTs, 1 WGD | None | [45] |
| *Mikania_micrantha-Smallanthus_sonchifolius* | Asteraceae | 2 WGTs, 1 WGD | 1 WGD in Sso | [45] |
| *Helianthus_annuus-Smallanthus_sonchifolius* | Asteraceae | 2 WGTs, 1 WGD | 1 WGD in Sso | [45] |
| *Daucus_carota-Angelica_sinensis* | Apiaceae | 1 WGT, 2 WGDs | None | [46] |
| *Aralia_elata-Eleutherococcus_senticosus* | Araliaceae | 1 WGT, 1 WGD | 1 WGD in Ese | [47] |
| *Aralia_elata-Panax_ginseng* | Araliaceae | 1 WGT, 1 WGD | 1 WGD in Pgi | [47] |
| *Panax_ginseng-Eleutherococcus_senticosus* | Araliaceae | 1 WGT, 1 WGD | 1 WGD in Pgi<br>1 WGD in Ese | [47] |
| *Lupinus_albus-Lupinus_angustifolius* | Fabaceae | 1 WGT, 1 WGD, 1 WGT | None | [48] |
| *Glycine_max-Glycine_soja* | Fabaceae | 1 WGT, 2 WGDs | None | [49] |
| *Glycine_max-Arachis_hypogaea* | Fabaceae | 1 WGT, 1 WGD | 1 WGD in Ahy<br>1 WGD in Gma | [50] |
| *Arachis_hypogaea-Arachis_monticola* | Fabaceae | 1 WGT, 2 WGDs | None | [51] |
| *Dalbergia_odorifera-Glycine_max* | Fabaceae | 1 WGT, 1 WGD | 1 WGD in Gma | [52] |

| | | | | |
|---|---|---|---|---|
| *Gillenia_trifoliata-Crataegus_pinnatifida* | Rosaceae | 1 WGT | 1 WGD in Cpi | [53] |
| *Crataegus_pinnatifida-Malus_x_domestica* | Rosaceae | 1 WGT, 1 WGD | None | [53] |
| *Carya_illinoinensis-Juglans_regia* | Juglandaceae | 1 WGT, 1 WGD | None | [54] |
| *Citrullus_lanatus-Begonia_loranthoides* | Cucurbitales | 1 WGT, 1 WGD | 1 WGD in Bma | [55] |
| *Citrullus_lanatus-Luffa_cylindrica* | Cucurbitaceae | 1 WGT, 1 WGD | None | [56] |
| *Citrullus_lanatus-Cucurbita_argyrosperma* | Cucurbitaceae | 1 WGT, 1 WGD | 1 WGD in Car | [55] |
| *Cucurbita_argyrosperma-Cucurbita_pepo* | Cucurbitaceae | 1 WGT, 2 WGDs | None | [55] |
| *Begonia_loranthoides-Begonia_peltatifolia* | Begoniaceae | 1 WGT, 2 WGDs | None | [55] |
| *Bruguiera_parviflora-Ceriops_tagal* | Rhizophoraceae | 1 WGT, 1 WGD | None | [57] |
| *Hevea_brasiliensis-Manihot_esculenta* | Euphorbiaceae | 1 WGT, 1 WGD | None | [58] |
| *Populus_trichocarpa-Populus_euphratica* | Salicaceae | 1 WGT, 1 WGD | None | [59] |
| *Salix_dunnii-Salix_brachista* | Salicaceae | 1 WGT, 1 WGD | None | [59] |
| *Eucalyptus_grandis-Punica_granatum* | Myrtales | 1 WGT, 1 WGD | None | [60] |
| *Eucalyptus_grandis-Syzygium_grande* | Myrtaceae | 1 WGT, 1 WGD | None | [60] |
| *Melastoma_candidum-Melastoma_dodecandrum* | Melastomataceae | 1 WGT, 3 WGDs | None | [61] |
| *Arabidopsis_thaliana-Capparis_spinosa* | Brassicales | 1 WGT, 1 WGD | 1 WGD in Ath<br>1 WGD in Csp | [62] |
| *Arabidopsis_thaliana-Aethionema_arabicum* | Brassicaceae | 1 WGT, 2 WGDs | None | [63] |
| *Arabidopsis_thaliana-Thlaspi_arvense* | Brassicaceae | 1 WGT, 2 WGDs | None | [64] |
| *Arabidopsis_thaliana-Brassica_rapa* | Brassicaceae | 1 WGT, 2 WGDs | 1 WGT in Bra | [65] |
| *Brassica_rapa-Sinapis_alba* | Brassicaceae | 1 WGT, 2 WGDs, 1 WGT | None | [66] |
| *Brassica_rapa-Crambe_hispanica* | Brassicaceae | 1 WGT, 2 WGDs, 1 WGT | None | [67] |
| *Theobroma_cacao-Gossypium_raimondii* | Malvaceae | 1 WGT | 1 WGM in Gra | [68] |
| *Gossypium_raimondii-Gossypioides_kirkii* | Malvaceae | 1 WGT, 1 WGM | None | [69] |
| *Aquilaria_sinensis-Stellera_chamaejasme* | Thymelaeaceae | 1 WGT, 1 WGD | None | [70] |
| *Mangifera_indica-Anacardium_occidentale* | Anacardiaceae | 1 WGT, 1 WGD | None | [71] |
| *Toona_sinensis-Toona_ciliata* | Meliaceae | 1 WGT, 1 WGD | None | [72] |

Note: WGD, whole genome duplication; WGT, whole genome triplication; WGM, whole genome multiplication.

**Table S2. Genomic data used in this study (accessed before 2023-06-29).**

| **Species** | **Order** | **Family** | **Source** |
|---|---|---|---|
| *Amborella_trichopoda* | Amborellales | Amborellaceae | [73] |
| *Angelica_sinensis* | Apiales | Apiaceae | [46] |

| | | | |
|---|---|---|---|
| *Apium_graveolens* | Apiales | Apiaceae | [17] |
| *Centella_asiatica* | Apiales | Apiaceae | [74] |
| *Coriandrum_sativum* | Apiales | Apiaceae | [75] |
| *Daucus_carota* | Apiales | Apiaceae | [76] |
| *Aralia_elata* | Apiales | Araliaceae | [47] |
| *Eleutherococcus_senticosus* | Apiales | Araliaceae | [77] |
| *Panax_ginseng* | Apiales | Araliaceae | [78] |
| *Panax_notoginseng* | Apiales | Araliaceae | [79] |
| *Ilex_polyneura* | Aquifoliales | Aquifoliaceae | [80] |
| *Arctium_lappa* | Asterales | Asteraceae | [45] |
| *Helianthus_annuus* | Asterales | Asteraceae | [81] |
| *Mikania_micrantha* | Asterales | Asteraceae | [82] |
| *Smallanthus_sonchifolius* | Asterales | Asteraceae | [45] |
| *Stevia_rebaudiana* | Asterales | Asteraceae | [83] |
| *Taraxacum_mongolicum* | Asterales | Asteraceae | [84] |
| *Platycodon_grandiflorus* | Asterales | Campanulaceae | [85] |
| *Nymphoides_indica* | Asterales | Menyanthaceae | [86] |
| *Echium_plantagineum* | Boraginales | Boraginaceae | [87] |
| *Bretschneidera_sinensis* | Brassicales | Akaniaceae | [88] |
| *Aethionema_arabicum* | Brassicales | Brassicaceae | [89] |
| *Arabidopsis_arenosa* | Brassicales | Brassicaceae | Genbank: GCA_905216605 |
| *Arabidopsis_thaliana* | Brassicales | Brassicaceae | [90] |
| *Arabidopsis_thaliana_x_Arabidopsis_arenosa* | Brassicales | Brassicaceae | [91] |
| *Brassica_rapa* | Brassicales | Brassicaceae | [92] |
| *Crambe_hispanica* | Brassicales | Brassicaceae | [67] |
| *Sinapis_alba* | Brassicales | Brassicaceae | [66] |
| *Thlaspi_arvense* | Brassicales | Brassicaceae | [64] |
| *Capparis_spinosa* | Brassicales | Capparaceae | [62] |
| *Carica_papaya* | Brassicales | Caricaceae | [93] |
| *Cleome_gynandra* | Brassicales | Cleomaceae | [94] |
| *Buxus_austroyunnanensis* | Buxales | Buxaceae | [95] |
| *Mesembryanthemum_crystallinum* | Caryophyllales | Aizoaceae | [96] |
| *Amaranthus_hypochondriacus* | Caryophyllales | Amaranthaceae | [97] |
| *Amaranthus_tricolor* | Caryophyllales | Amaranthaceae | [26] |
| *Selenicereus_undatus* | Caryophyllales | Cactaceae | [98] |
| *Gypsophila_paniculata* | Caryophyllales | Caryophyllaceae | [99] |
| *Beta_vulgaris* | Caryophyllales | Chenopodiaceae | [100] |
| *Spinacia_oleracea* | Caryophyllales | Chenopodiaceae | [101] |
| *Limonium_bicolor* | Caryophyllales | Plumbaginaceae | [102] |
| *Fagopyrum_esculentum* | Caryophyllales | Polygonaceae | GWH: GWHBJBK00000000 |
| *Fagopyrum_tataricum* | Caryophyllales | Polygonaceae | GWH: GWHBJBL00000000 |
| *Portulaca_amilis* | Caryophyllales | Portulacaceae | [103] |

| | | | |
|---|---|---|---|
| *Simmondsia_chinensis* | Caryophyllales | Simmondsiaceae | 104 |
| *Tripterygium_wilfordii* | Celastrales | Celastraceae | 21 |
| *Cornus_controversa* | Cornales | Cornaceae | 105 |
| *Camptotheca_acuminata* | Cornales | Nyssaceae | 107 |
| *Davidia_involucrata* | Cornales | Nyssaceae | 30 |
| *Nyssa_sinensis* | Cornales | Nyssaceae | 108 |
| *Euscaphis_japonica* | Crossosomatales | Staphyleaceae | 23 |
| *Begonia_loranthoides* | Cucurbitales | Begoniaceae | 109 |
| *Begonia_masoniana* | Cucurbitales | Begoniaceae | 109 |
| *Begonia_peltatifolia* | Cucurbitales | Begoniaceae | 109 |
| *Coriaria_nepalensis* | Cucurbitales | Coriariaceae | 110 |
| *Citrullus_lanatus* | Cucurbitales | Cucurbitaceae | 111 |
| *Cucurbita_argyrosperma* | Cucurbitales | Cucurbitaceae | 112 |
| *Cucurbita_pepo* | Cucurbitales | Cucurbitaceae | 113 |
| *Luffa_cylindrica* | Cucurbitales | Cucurbitaceae | 114 |
| *Lonicera_japonica* | Dipsacales | Caprifoliaceae | 115 |
| *Actinidia_chinensis* | Ericales | Actinidiaceae | 116 |
| *Impatiens_glandulifera* | Ericales | Balsaminaceae | Refseq: GCF_907164915 |
| *Diospyros_lotus* | Ericales | Ebenaceae | 117 |
| *Rhododendron_delavayi* | Ericales | Ericaceae | 118 |
| *Rhododendron_simsii* | Ericales | Ericaceae | 29 |
| *Vaccinium_corymbosum* | Ericales | Ericaceae | 119 |
| *Gilia_yorkii* | Ericales | Polemoniaceae | 120 |
| *Aegiceras_corniculatum* | Ericales | Primulaceae | 121 |
| *Vitellaria_paradoxa* | Ericales | Sapotaceae | 122 |
| *Camellia_sinensis* | Ericales | Theaceae | 123 |
| *Arachis_duranensis* | Fabales | Fabaceae | 124 |
| *Arachis_hypogaea* | Fabales | Fabaceae | 50 |
| *Arachis_ipaensis* | Fabales | Fabaceae | 124 |
| *Arachis_monticola* | Fabales | Fabaceae | 51 |
| *Cercis_chinensis* | Fabales | Fabaceae | 125 |
| *Dalbergia_odorifera* | Fabales | Fabaceae | 52 |
| *Glycine_max* | Fabales | Fabaceae | 126 |
| *Glycine_soja* | Fabales | Fabaceae | 126 |
| *Lupinus_albus* | Fabales | Fabaceae | 127 |
| *Lupinus_angustifolius* | Fabales | Fabaceae | 48 |
| *Quillaja_saponaria* | Fabales | Quillajaceae | 128 |
| *Betula_pendula* | Fagales | Betulaceae | 22 |
| *Corylus_mandshurica* | Fagales | Betulaceae | 129 |
| *Casuarina_equisetifolia* | Fagales | Casuarinaceae | 130 |
| *Quercus_variabilis* | Fagales | Fagaceae | 131 |
| *Carya_illinoinensis* | Fagales | Juglandaceae | 132 |
| *Juglans_regia* | Fagales | Juglandaceae | 133 |

| | | | |
|---|---|---|---|
| *Morella_rubra* | Fagales | Myricaceae | 134 |
| *Eucommia_ulmoides* | Garryales | Eucommiaceae | 16 |
| *Catharanthus_roseus* | Gentianales | Apocynaceae | 135 |
| *Marsdenia_tenacissima* | Gentianales | Apocynaceae | 37 |
| *Gelsemium_elegans* | Gentianales | Gelsemiaceae | 136 |
| *Eustoma_grandiflorum* | Gentianales | Gentianaceae | 137 |
| *Coffea_canephora* | Gentianales | Rubiaceae | 15 |
| *Neolamarckia_cadamba* | Gentianales | Rubiaceae | 39 |
| *Ophiorrhiza_pumila* | Gentianales | Rubiaceae | 38 |
| *Strobilanthes_cusia* | Lamiales | Acanthaceae | 139 |
| *Jacaranda_mimosifolia* | Lamiales | Bignoniaceae | 140 |
| *Primulina_huaijiensis* | Lamiales | Gesneriaceae | 141 |
| *Salvia_hispanica* | Lamiales | Lamiaceae | 34 |
| *Salvia_splendens* | Lamiales | Lamiaceae | 33 |
| *Tectona_grandis* | Lamiales | Lamiaceae | 142 |
| *Lindernia_brevidens* | Lamiales | Linderniaceae | 143 |
| *Jasminum_sambac* | Lamiales | Oleaceae | 35 |
| *Osmanthus_fragrans* | Lamiales | Oleaceae | 144 |
| *Syringa_oblata* | Lamiales | Oleaceae | 145 |
| *Orobanche_cernua* | Lamiales | Orobanchaceae | 32 |
| *Paulownia_fortunei* | Lamiales | Paulowniaceae | 31 |
| *Sesamum_indicum* | Lamiales | Pedaliaceae | 146 |
| *Mimulus_guttatus* | Lamiales | Phrymaceae | 147 |
| *Antirrhinum_majus* | Lamiales | Plantaginaceae | 148 |
| *Buddleja_alternifolia* | Lamiales | Scrophulariaceae | 149 |
| *Petrea_volubilis* | Lamiales | Verbenaceae | 150 |
| *Chimonanthus_salicifolius* | Laurales | Calycanthaceae | 3 |
| *Cinnamomum_kanehirae* | Laurales | Lauraceae | 151 |
| *Phoebe_bournei* | Laurales | Lauraceae | 4 |
| *Liriodendron_chinense* | Magnoliales | Magnoliaceae | 152 |
| *Magnolia_biondii* | Magnoliales | Magnoliaceae | 2 |
| *Erythroxylum_novogranatense* | Malpighiales | Erythroxylaceae | 40 |
| *Euphorbia_peplus* | Malpighiales | Euphorbiaceae | 153 |
| *Hevea_brasiliensis* | Malpighiales | Euphorbiaceae | 58 |
| *Manihot_esculenta* | Malpighiales | Euphorbiaceae | 154 |
| *Linum_tenue* | Malpighiales | Linaceae | 155 |
| *Passiflora_edulis* | Malpighiales | Passifloraceae | 156 |
| *Cladopus_chinensis* | Malpighiales | Podostemaceae | 157 |
| *Bruguiera_parviflora* | Malpighiales | Rhizophoraceae | 57 |
| *Bruguiera_sexangula* | Malpighiales | Rhizophoraceae | 158 |
| *Ceriops_tagal* | Malpighiales | Rhizophoraceae | 159 |
| *Populus_euphratica* | Malpighiales | Salicaceae | 160 |
| *Populus_trichocarpa* | Malpighiales | Salicaceae | 161 |

| | | | |
|---|---|---|---|
| *Salix_brachista* | Malpighiales | Salicaceae | 162 |
| *Salix_dunnii* | Malpighiales | Salicaceae | 163 |
| *Dipterocarpus_turbinatus* | Malvales | Dipterocarpaceae | 164 |
| *Gossypioides_kirkii* | Malvales | Malvaceae | 165 |
| *Gossypium_raimondii* | Malvales | Malvaceae | 69 |
| *Theobroma_cacao* | Malvales | Malvaceae | 166 |
| *Aquilaria_sinensis* | Malvales | Thymelaeaceae | 167 |
| *Stellera_chamaejasme* | Malvales | Thymelaeaceae | 70 |
| *Lumnitzera_racemosa* | Myrtales | Combretaceae | 168 |
| *Punica_granatum* | Myrtales | Lythraceae | 169 |
| *Melastoma_candidum* | Myrtales | Melastomataceae | 61 |
| *Melastoma_dodecandrum* | Myrtales | Melastomataceae | 170 |
| *Eucalyptus_grandis* | Myrtales | Myrtaceae | 171 |
| *Syzygium_grande* | Myrtales | Myrtaceae | 60 |
| *Chamaenerion_angustifolium* | Myrtales | Onagraceae | Genbank: GCA_946814005 |
| *Averrhoa_carambola* | Oxalidales | Oxalidaceae | 172 |
| *Aristolochia_fimbriata* | Piperales | Aristolochiaceae | 1 |
| *Triticum_aestivum* | Poales | Poaceae | 173 |
| *Triticum_turgidum* | Poales | Poaceae | 174 |
| *Nelumbo_nucifera* | Proteales | Nelumbonaceae | 175 |
| *Macadamia_integrifolia* | Proteales | Proteaceae | 176 |
| *Protea_cynaroides* | Proteales | Proteaceae | 5 |
| *Telopea_speciosissima* | Proteales | Proteaceae | 177 |
| *Akebia_trifoliata* | Ranunculales | Lardizabalaceae | 178 |
| *Corydalis_tomentella* | Ranunculales | Papaveraceae | 7 |
| *Papaver_setigerum* | Ranunculales | Papaveraceae | 179 |
| *Papaver_somniferum* | Ranunculales | Papaveraceae | 179 |
| *Aquilegia_coerulea* | Ranunculales | Ranunculaceae | 180 |
| *Coptis_chinensis* | Ranunculales | Ranunculaceae | 181 |
| *Cannabis_sativa* | Rosales | Cannabaceae | 182 |
| *Hippophae_rhamnoides* | Rosales | Elaeagnaceae | 183 |
| *Ficus_pumila* | Rosales | Moraceae | 184 |
| *Ziziphus_spinosa* | Rosales | Rhamnaceae | 185 |
| *Crataegus_pinnatifida* | Rosales | Rosaceae | 53 |
| *Fragaria_iinumae* | Rosales | Rosaceae | 186 |
| *Fragaria_nipponica* | Rosales | Rosaceae | 187 |
| *Fragaria_vesca* | Rosales | Rosaceae | 188 |
| *Fragaria_viridis* | Rosales | Rosaceae | 189 |
| *Fragaria_x_ananassa* | Rosales | Rosaceae | 190 |
| *Gillenia_trifoliata* | Rosales | Rosaceae | 191 |
| *Malus_x_domestica* | Rosales | Rosaceae | 192 |
| *Prunus_persica* | Rosales | Rosaceae | 20 |
| *Boehmeria_nivea* | Rosales | Urticaceae | 193 |

| | | | |
|---|---|---|---|
| *Taxillus_chinensis* | Santalales | Loranthaceae | 194 |
| *Santalum_album* | Santalales | Santalaceae | 28 |
| *Santalum_yasi* | Santalales | Santalaceae | 28 |
| *Malania_oleifera* | Santalales | Ximeniaceae | 195 |
| *Anacardium_occidentale* | Sapindales | Anacardiaceae | JGI |
| *Mangifera_indica* | Sapindales | Anacardiaceae | 196 |
| *Toona_ciliata* | Sapindales | Meliaceae | 197 |
| *Toona_sinensis* | Sapindales | Meliaceae | 198 |
| *Nitraria_sibirica* | Sapindales | Nitrariaceae | 199 |
| *Citrus_sinensis* | Sapindales | Rutaceae | 200 |
| *Acer_yangbiense* | Sapindales | Sapindaceae | 25 |
| *Ailanthus_altissimus* | Sapindales | Simaroubaceae | Genbank:GCA_946807835 |
| *Cercidiphyllum_japonicum* | Saxifragales | Cercidiphyllaceae | 18 |
| *Paeonia_ostii* | Saxifragales | Paeoniaceae | 19 |
| *Cuscuta_europaea* | Solanales | Convolvulaceae | 201 |
| *Ipomoea_cairica* | Solanales | Convolvulaceae | 43 |
| *Anisodus_acutangulus* | Solanales | Solanaceae | 40 |
| *Lycium_barbarum* | Solanales | Solanaceae | 202 |
| *Nicotiana_tabacum* | Solanales | Solanaceae | 41 |
| *Solanum_lycopersicum* | Solanales | Solanaceae | 203 |
| *Tetracentron_sinense* | Trochodendrales | Trochodendraceae | 8 |
| *Trochodendron_aralioides* | Trochodendrales | Trochodendraceae | 204 |
| *Vitis_vinifera* | Vitales | Vitaceae | 205 |
| *Tetraena_mongolica* | Zygophyllales | Zygophyllaceae | 206 |

Note: Genbank/Refseq: https://www.ncbi.nlm.nih.gov/assembly/; GWH: https://ngdc.cncb.ac.cn/gwh/; JGI: https://phytozome-next.jgi.doe.gov/.
